# Population genomics of the Viking world

**DOI:** 10.1101/703405

**Authors:** Ashot Margaryan, Daniel Lawson, Martin Sikora, Fernando Racimo, Simon Rasmussen, Ida Moltke, Lara Cassidy, Emil Jørsboe, Andrés Ingason, Mikkel Pedersen, Thorfinn Korneliussen, Helene Wilhelmson, Magdalena Buś, Peter de Barros Damgaard, Rui Martiniano, Gabriel Renaud, Claude Bhérer, J. Víctor Moreno-Mayar, Anna Fotakis, Marie Allen, Martyna Molak, Enrico Cappellini, Gabriele Scorrano, Alexandra Buzhilova, Allison Fox, Anders Albrechtsen, Berit Schütz, Birgitte Skar, Caroline Arcini, Ceri Falys, Charlotte Hedenstierna Jonson, Dariusz Błaszczyk, Denis Pezhemsky, Gordon Turner-Walker, Hildur Gestsdóttir, Inge Lundstrøm, Ingrid Gustin, Ingrid Mainland, Inna Potekhina, Italo Muntoni, Jade Cheng, Jesper Stenderup, Jilong Ma, Julie Gibson, Jüri Peets, Jörgen Gustafsson, Katrine Iversen, Linzi Simpson, Lisa Strand, Louise Loe, Maeve Sikora, Marek Florek, Maria Vretemark, Mark Redknap, Monika Bajka, Tamara Pushkina, Morten Søvsø, Natalia Grigoreva, Tom Christensen, Ole Kastholm, Otto Uldum, Pasquale Favia, Per Holck, Raili Allmäe, Sabine Sten, Símun Arge, Sturla Ellingvåg, Vayacheslav Moiseyev, Wiesław Bogdanowicz, Yvonne Magnusson, Ludovic Orlando, Daniel Bradley, Marie Louise Jørkov, Jette Arneborg, Niels Lynnerup, Neil Price, M. Thomas Gilbert, Morten Allentoft, Jan Bill, Søren Sindbæk, Lotte Hedeager, Kristian Kristiansen, Rasmus Nielsen, Thomas Werge, Eske Willerslev

**Affiliations:** Lundbeck Foundation GeoGenetics Centre, GLOBE Institute, University of Copenhagen, Øster Voldgade 5-7, 1350 Copenhagen K, Denmark; Institute of Molecular Biology, National Academy of Sciences, 7, Hasratian St., 0014, Yerevan, Armenia; Section for Evolutionary Genomics, GLOBE Institute, University of Copenhagen, Øster Voldgade 5-7, 1350 Copenhagen K, Denmark; MRC Integrative Epidemiology Unit, University of Bristol, Bristol, UK; Novo Nordisk Foundation Center for Protein Research, Faculty of Health and Medical Sciences, University of Copenhagen, Blegdamsvej 3B, 2200 Copenhagen, Denmark; Department of Biology, The Bioinformatics Centre, University of Copenhagen, 2200 Copenhagen N, Denmark; Smurfit Institute of Genetics, Trinity College Dublin, Dublin; Historical archaeology, Department of Archaeology and Ancient history, Lund University, PB 192, SE 22100 Lund, Sweden; Sydsvensk arkeologi AB, PB 134, SE 29122 Kristianstad, Sweden; Department of Immunology, Genetics and Pathology, Science for Life Laboratory, Uppsala University, 751 08 Uppsala, Sweden; Department of Genetics, University of Cambridge, Downing Street, Cambridge CB2 3EH, UK; New York Genome Center, 101 Avenue of the Americas, New York, NY, USA, 10013; National Institute of Genomic Medicine (INMEGEN), Periférico Sur 4809, 14610 Mexico City, Mexico; Museum and Institute of Zoology, Polish Academy of Sciences, Wilcza 64, 00-679 Warsaw, Poland; Anuchin Research Institute and Museum of Anthropology, Moscow State University; Manx National Heritage, Kingswood Grove, Douglas, Isle of Man, British Isles IM1 3LY; Upplandsmuseet, Drottninggatan 7, 753 10 Uppsala, Sweden; NTNU University Museum, Department of Archaeology and Cultural History Norway; Arkeologerna; Thames Valley Archaeological Services (TVAS), Reading, UK; Department of Archaeology and Ancient History, Uppsala University, Box 626, 751 26 Uppsala, Sweden; Institute of Archaeology, University of Warsaw, ul. Krakowskie Przedmieście 26/28, 00-927 Warsaw, Poland; Department of Cultural Heritage Conservation, National Yunlin University of Science and Technology, Douliou, Taiwan; Institute of Archaeology, Iceland. Bárugata 3, 101 Reykjavík, Iceland; UHI Archaeology Institute, University of the Highlands and Islands, Orkney College, Kirkwall, Orkney, KW15 1LX; Department of Bioarchaeology, Institute of Archaeology of Natioanal Academy of Sciences of Ukraine, 12 Geroiv Stalingrada Ave. 04210 Kyiv, Ukraine; Soprintendenza Archeologia, Belle Arti e Paesaggio per le Province di Barletta - Andria - Trani e Foggia, Via Alberto Alvarez Valentini, 8 - 71121 Foggia, Italy; Archaeological Research Collection, Tallinn University, Rüütli 10, Tallinn 10130, Estonia; Jönköping county museum, Jönköping, Sweden; Trinity College Dublin; Oxford Archaeology, Janus House, Osney Mead, Oxford OX2 0ES, UK; Heritage Burial Services, Oxford Archaeology, Janus House, Osney Mead, Oxford OX2 0ES, UK; National Museum of Ireland, Kildare Street, Dublin 2, Ireland; Institute of Archaeology, Maria Curie-Sklodowska University in Lublin, Pl. M. Curie-Sklodowska 4, 20-035 Lublin, Poland; Västergötlands museum, Box 253, 532 23 Skara Sweden; National Museum Cardiff; “Trzy Epoki” Archaeological Service, Poland; Museum of Southwest Jutland; Institute for the history of material culture, Russian Academy of Sciences, Dvotsovaya Emb., 18, Saint-Petersburg, Russia, 191186; National Museum of Denmark, Frederiksholms Kanal 12, DK-1220 Copenhagen, Denmark; Roskilde Museum, Museum Organization ROMU, Sankt Ols Stræde 3, DK-4000 Roskilde, Denmark; Langelands Museum, Jens Winthersvej 12. 5900 Rudkøbing, Langeland, Denmark; Department of Humanities, University of Foggia, Via Arpi, 176, 71121 Foggia, Italy; Department of Molecular Medicine, Faculty of Medicine, University of Oslo; Department of Archaeology and Ancient History, Uppsala University Campus Gotland; Tjóðsavnið - Faroe Islands National Museum. Kúrdalsvegur 15. Postboks 1155. FO-110 Tórshavn; Peter the Great Museum of Anthropology and Ethnography (Kunstkamera), Russian Academy of Science, University Emb, 3, SPb, Russia, 199034; Malmö Museum, Box 406, 201 24 Malmö, Sweden; Laboratoire d’Anthropobiologie Moléculaire et d’Imagerie de Synthèse, CNRS UMR 5288, Université de Toulouse, Université Paul Sabatier, 31000 Toulouse, France; Department of Forensic Medicine, University of Copenhagen, Frederik V’s vej 11, 2100 Copenhagen; Department of Natural History, NTNU; Museum of Cultural History, University of Oslo, P.O. Box 6762 St. Olavs plass, 0160 Oslo, Norway; Centre for Urban Network Evolutions (UrbNet), Aarhus University, School of Culture and Society, Moesgård Allé 20, building 4215, DK-8270 Højbjerg, Denmark; Institute of Archaeology, Conservation and History, Pb. 1019 Blindern, 0315 Oslo, Norway; Department of Historical Studies, University of Gothenburg; Departments of Integrative Biology and Statistics, UC Berkeley, Berkeley, CA 94720, USA; Department of Clinical Medicine, University of Copenhagen, Copenhagen, Denmark; Institute of Biological Psychiatry, Mental Health Services Copenhagen, Copenhagen, Denmark; The Lundbeck Foundation Initiative for Integrative Psychiatric Research, iPSYCH, Denmark; Department of Zoology, University of Cambridge, UK; The Danish Institute for Advanced Study, University of Southern Denmark; The Wellcome Trust Sanger Institute, Cambridge, UK; School of GeoSciences, University of Edinburgh; Department of Health Technology, Section for Bioinformatics, Technical University of Denmark, DTU, 2800 Kgs. Lyngby, Denmark

**Author notes:** These authors contributed equally to this work (Ashot Margaryan, Daniel Lawson, Martin Sikora, Fernando Racimo).

## Abstract

The Viking maritime expansion from Scandinavia (Denmark, Norway, and Sweden) marks one of the swiftest and most far-flung cultural transformations in global history. During this time (c. 750 to 1050 CE), the Vikings reached most of western Eurasia, Greenland, and North America, and left a cultural legacy that persists till today. To understand the genetic structure and influence of the Viking expansion, we sequenced the genomes of 442 ancient humans from across Europe and Greenland ranging from the Bronze Age (c. 2400 BC) to the early Modern period (c. 1600 CE), with particular emphasis on the Viking Age. We find that the period preceding the Viking Age was accompanied by foreign gene flow into Scandinavia from the south and east: spreading from Denmark and eastern Sweden to the rest of Scandinavia. Despite the close linguistic similarities of modern Scandinavian languages, we observe genetic structure within Scandinavia, suggesting that regional population differences were already present 1,000 years ago. We find evidence for a majority of Danish Viking presence in England, Swedish Viking presence in the Baltic, and Norwegian Viking presence in Ireland, Iceland, and Greenland. Additionally, we see substantial foreign European ancestry entering Scandinavia during the Viking Age. We also find that several of the members of the only archaeologically well-attested Viking expedition were close family members. By comparing Viking Scandinavian genomes with present-day Scandinavian genomes, we find that pigmentation-associated loci have undergone strong population differentiation during the last millennia. Finally, we are able to trace the allele frequency dynamics of positively selected loci with unprecedented detail, including the lactase persistence allele and various alleles associated with the immune response. We conclude that the Viking diaspora was characterized by substantial foreign engagement: distinct Viking populations influenced the genomic makeup of different regions of Europe, while Scandinavia also experienced increased contact with the rest of the continent.

## Introduction

Three centuries from approximately 750 to 1050 CE mark a pivotal change for the peoples of Scandinavia. The maritime transformation commonly known as the Viking Age (VA) altered the political, cultural and demographic map of Europe in ways that are evident even today. The Vikings established systems of trade and settlement that stretched from the eastern American seaboard to the Asian steppe^1^. They also exported new ideas, technologies, language, beliefs and practices to these lands. In the process, they gradually developed new socio-political structures, assimilated cultural influences, and adopted the Christian faith^2^.

Currently, most of our understanding of the VA is based on written sources and archaeological evidence. The VA as a historical period has been framed by the first clearly documented raid on Lindisfarne in 793 CE, and the defeat of a Norwegian army at Stamford Bridge in 1066 CE. More recent perspectives emphasize long-term, multi-causal social processes with after-effects that varied greatly by region^3–5^. Similarly, the notion of a Viking ‘expansion’, implying deliberate drive and purpose, has been supplemented by the more fluid concept of a ‘diaspora’ that developed over time^2^. Under this framework, however, the role of demographic dynamics has remained unclear, as has the question of whether VA Scandinavia was genetically structured or represented a homogenous population. Similarly, we still do not know to what extent Vikings mixed with local populations they encountered and how much foreign ancestry was brought back to Scandinavia.

In order to explore the genomic history of the Viking era, we shotgun sequenced 442 ancient human remains, from the Bronze Age c. 2400 BC to the Medieval Age c. 1600 AD (Fig. 1). The majority of these individuals (n=376) were sequenced to between 0.1 and 11X average depth of coverage. The dataset includes Bronze Age (n=2) and Iron Age (n=10) individuals from Scandinavia; Early Viking Age (n=43) individuals from Estonia (n=34), Denmark (n=6) and Sweden (n=3); ancient individuals associated with Norse culture from Greenland (n=23), VA individuals from Denmark (n=78), Faroe Islands (n=1), Iceland (n=17), Ireland (n=4), Norway (n=29), Poland (n=8), Russia (n=33), Sweden (n=118), UK (n=42), Ukraine (n=3) as well as medieval individuals from Faroe Islands (n=16), Italy (n=5), Norway (n=7), Poland (n=2) and Ukraine (n=1). The VA individuals were supplemented with additional published genomes (n=21) from Sigtuna, in Sweden^6^. The skeletons originate from major archaeological sites of VA Scandinavian settlements and activities from Europe to Greenland (Supplementary Table 1). The data from the ancient individuals were analyzed together with previously published data from a total of 3,855 present-day individuals across two reference panels, and data from 922 individuals of ancient origin (Supplementary Note 6).

**Fig. 1:**
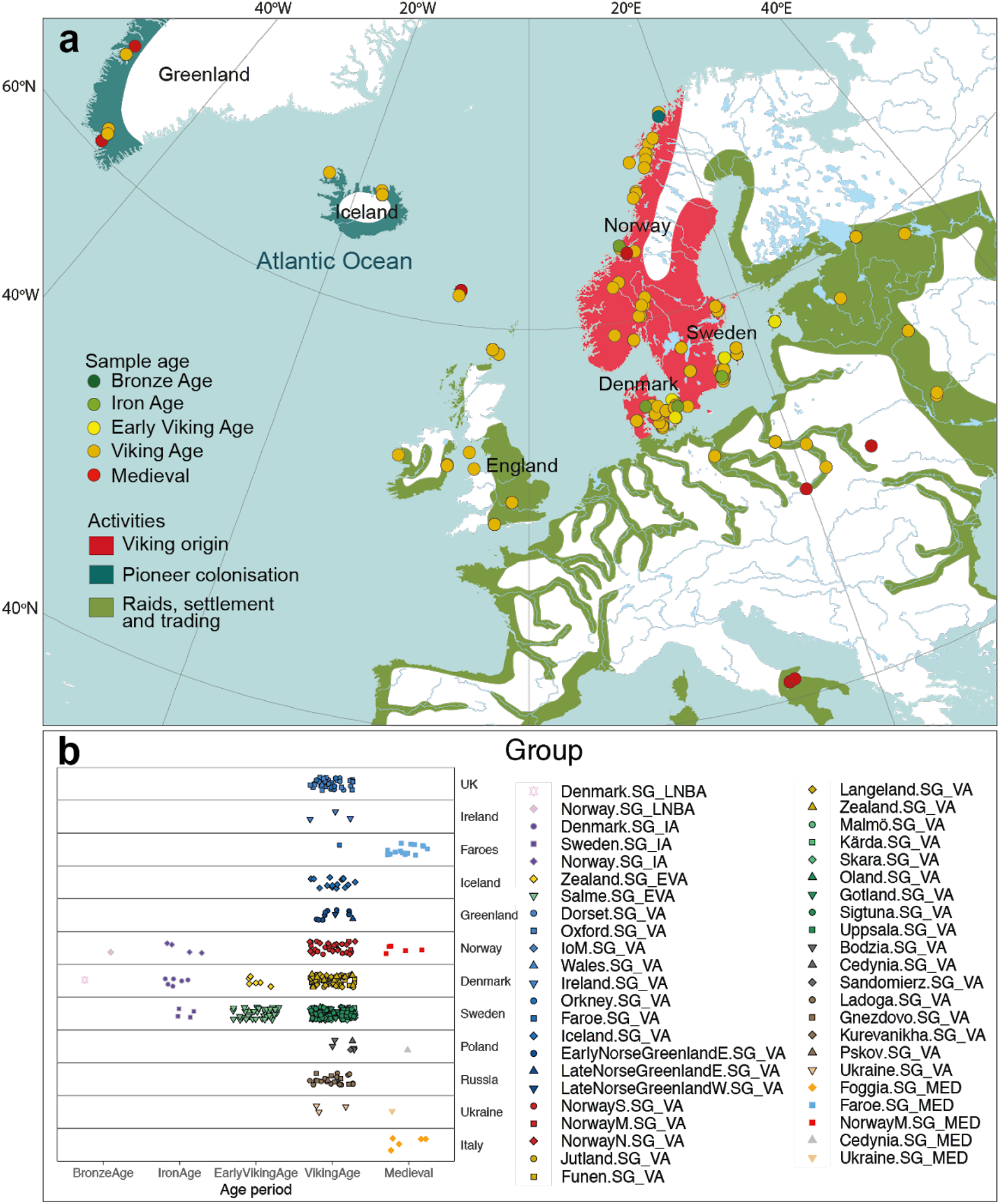
Map of the “Viking World” from 8^th^ till 11^th^ centuries. Different symbols on the map (**a**) correspond to ancient sites of a specific age/culture. The ancient samples are divided into the following five broad categories: Bronze Age (BA) - c. 2500 BC - 900 BC; Iron Age (IA) - c. 900 BC to 700 CE; Early Viking Age (EVA) - c. 700 to 800 CE; VA - c. 800 to 1100 CE; Medieval - c. 1100 to 1600 CE. **b**, All ancient individuals from this study (n=442) and published VA samples (n=21) from Sigtuna^6^ are categorized based on their spatio-temporal origin.

### Scandinavian genetic ancestry and the beginnings of the Viking era

Although VA Scandinavians shared a common cultural, linguistic and material background, there was no common word for Scandinavian identity at that time^1^. The word ‘Viking’ is used in contemporary sources to mean a ‘pirate’ or ‘sea warrior’^2^. As such, there is no single ‘Viking world’, but a coalescence of ‘Viking worlds’ marked by rapidly growing maritime exploration, trade, war and colonization, following the adoption of deep-sea navigation among the coastal populations of Scandinavia and the Baltic Sea area^7,8^. Thus, it is unclear whether the Viking-phenomenon refers to people with a recently shared genetic background and if foreign influence initiated or accompanied the transition from the Scandinavian Iron Age into the Viking era.

To assess the genetic relationship of the VA Scandinavians with that of earlier European peoples, we performed genetic clustering using multi-dimensional scaling (MDS) on a pairwise identity-by-state (IBS) sharing matrix, as well as latent mixed-ancestry models (Admixture)^9^. We find that the majority of our samples broadly cluster within the range of European Bronze Age (BA) and Iron Age (IA) populations, characterized by an ancestry component that is related to pastoralist populations from the Pontic-Caspian steppe (Fig. 2a and Extended Data Fig. 2) entering Europe around 5000 BP^10,11^. A different dimensionality reduction technique using uniform manifold approximation and projection (UMAP) revealed additional fine-scale genetic structure. European individuals from the Bronze Age and onwards are generally distributed within a broad area anchored by four ancestry clusters across the two UMAP dimensions: Early BA individuals from the Steppe; pre-BA Neolithic Europeans; Baltic BA individuals; and Scandinavian IA and early VA individuals (Fig. 2b). We observe a wide range of distributions for VA individuals within this broad area, with notable differences between geographic regions (Fig. S8.10), indicating complex fine-scale structure among the different groups. Modelling Scandinavian groups from the BA and onwards as mixtures of three ancestral components (Mesolithic hunter-gatherers; Anatolian Neolithic; Steppe early BA), again revealed subtle differences in their composition. We find that the transition from the BA to the IA is accompanied by a reduction in Neolithic farmer ancestry, with a corresponding increase in both Steppe-like ancestry and hunter-gatherer ancestry (Extended Data Fig. 6). While most groups show a slight recovery of farmer ancestry during the VA, there is considerable variation in ancestry across Scandinavia. In particular, we observe a wide range of ancestry compositions among individuals from Sweden, with some groups in southern Sweden showing some of the highest farmer ancestry proportions (40% or more in individuals from Malmö, Kärda or Öland). Ancestry proportions in Norway and Denmark on the other hand appear more uniform (Extended Data Fig. 6). Finally, we detect an influx of low levels of “eastern” ancestry starting in the early VA, mostly constrained among groups from eastern and central Sweden as well as some Norwegian groups (Extended Data Fig. 6). Testing of putative source groups for this “eastern” ancestry revealed differing patterns among the Viking Age target groups, with contributions of either East Asian- or Caucasus-related ancestry (Supplementary Note 10).

**Fig. 2:**
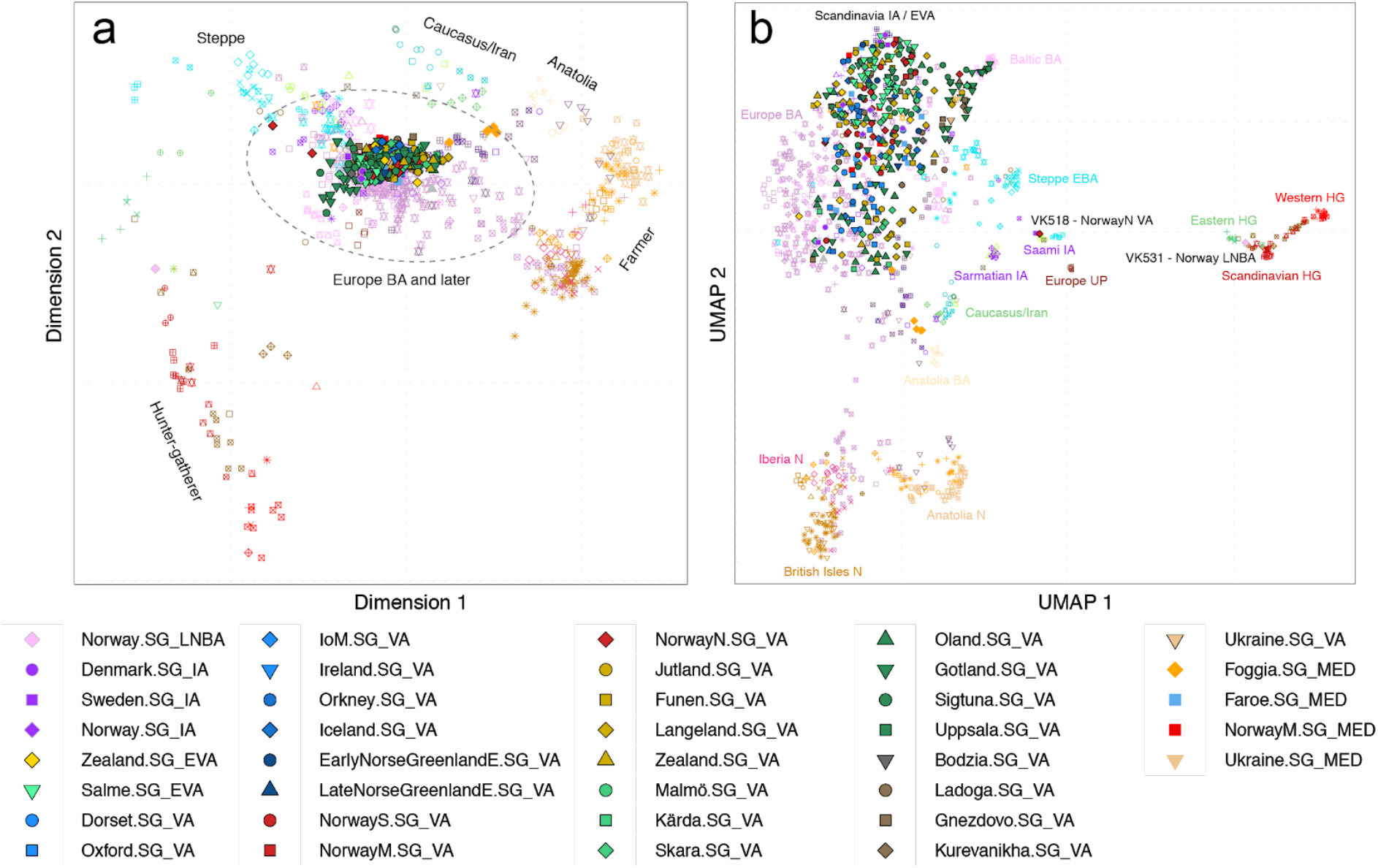
Genetic structure of VA samples. **a**, Multidimensional scaling (MDS) plot based on a pairwise identity-by-state (IBS) sharing matrix of the VA and other ancient samples (Supplementary Table 3). **b**, Uniform manifold approximation and projection (UMAP) analysis of the same dataset as in plot (**a)**.

Overall, our findings suggest that the genetic makeup of VA Scandinavia derives from mixtures of three earlier sources: Mesolithic hunter-gatherers, Neolithic farmers, and Bronze Age pastoralists. Intriguingly, our results also indicate ongoing gene flow from the south and east into Iron Age Scandinavia. Thus, these observations are consistent with archaeological claims of wide-ranging demographic turmoil in the aftermath of the Roman Empire with consequences for the Scandinavian populations during the late Iron Age^12,13^. We caution, however, that our sampling for the periods preceding the VA is still sparse, and hence do not provide a full picture of the genetic diversity across Scandinavia during that period.

### Genetic structure within Viking-Age Scandinavia

By the end of the Iron Age in the 8^th^ century CE, Scandinavia formed a patchwork of conflicting and competing kingdoms with a shared cultural background. For centuries, a political economy based on raiding, trading and gifts had been common^5^. However, the cause for the development of this economic and political system into the more organized maritime society of the Viking era remains debated^5^. It is commonly argued that seafaring^8,14^ contributed to create a densely interlinked Scandinavia during the Viking era^2,15,16^.

To disentangle the fine-scale population structure within VA Scandinavia, we performed genotype imputation on a subset of 300 individuals with sufficient coverage (>0.5X) and inferred genomic segments shared via identity-by-descent (IBD) within the context of a reference panel of 1,464 present-day Europeans, using IBDseq. We find that VA Scandinavians on average cluster into three groups according to their geographic origin, shifted towards their respective present-day counterparts in Denmark, Sweden and Norway (Fig. 3a). Closer inspection of the distributions for the different groups reveals additional complexity in their genetic structure (Fig. S10.1). We find that the ‘Norwegian’ cluster includes Norwegian IA individuals, who are distinct from both Swedish and Danish IA individuals which cluster together with the majority of central and eastern Swedish VA individuals. Many individuals from southwestern Sweden (e.g. Skara) cluster with Danish present-day individuals from the eastern islands (Funen, Zealand), skewing towards the ‘Swedish’ cluster with respect to early and more western Danish VA individuals (Jutland). Some individuals have strong affinity with Eastern Europeans, particularly those from the island of Gotland in eastern Sweden. The latter likely reflects individuals with Baltic ancestry, as clustering with Baltic BA individuals is evident in the IBS-UMAP analysis (Fig. 2b) and through f_4_-statistics (Extended Data Fig. 4).

**Fig. 3:**
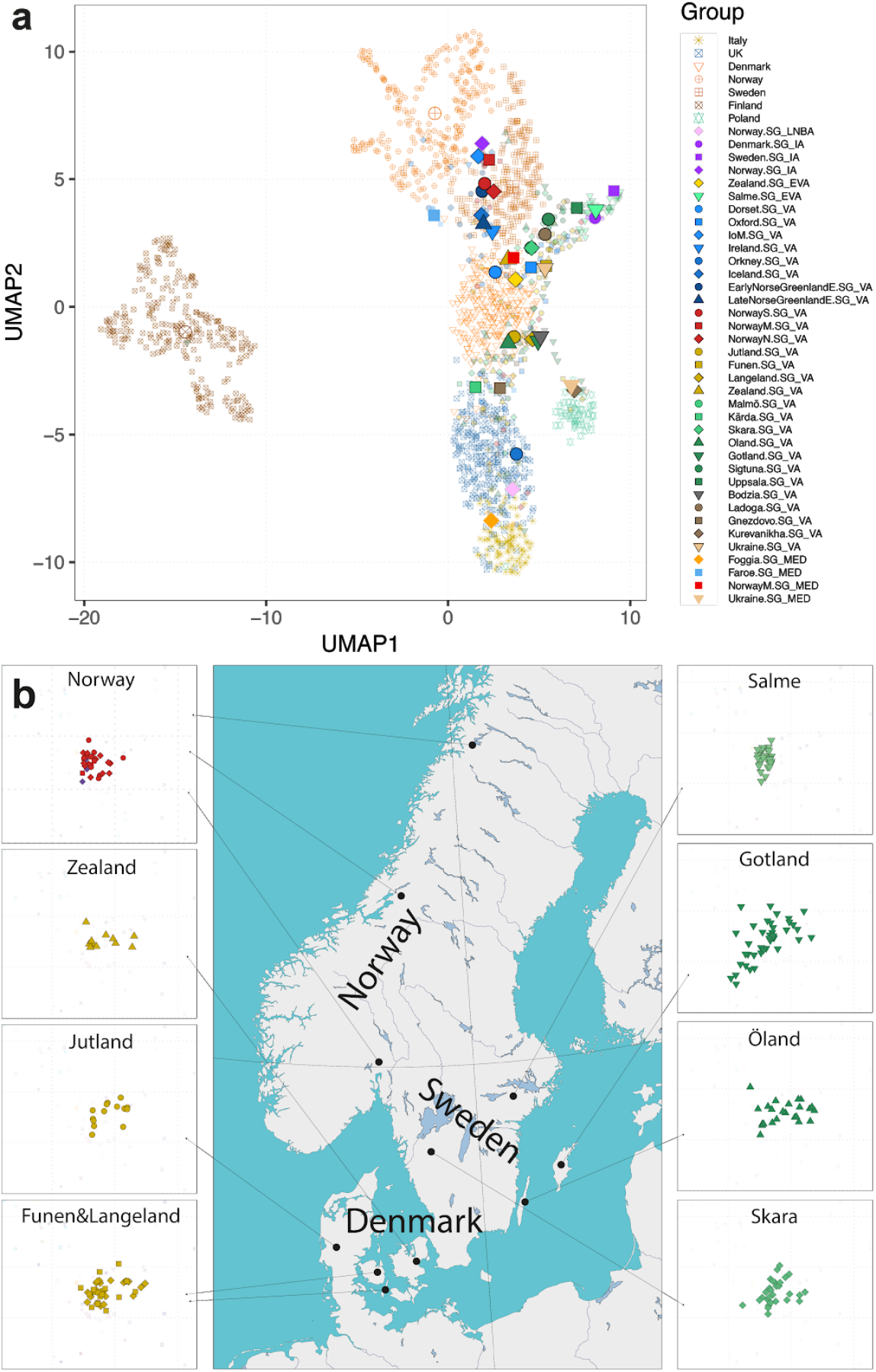
Genetic structure and diversity of ancient samples. **a**, Uniform manifold approximation and projection (UMAP) analysis of the ancient and modern Scandinavian individuals based on the first 10 dimensions of MDS using identity-by-descent (IBD) segments of imputed individuals. Large symbols indicate median coordinates for each group. **b**, Genetic diversity in major Scandinavian VA populations. Plots next to the map show MDS analysis based on a pairwise IBS sharing matrix. Here “Norway” represents all the sites from Norway. The scale is identical for all the plots.

To further quantify the within-Scandinavia population structure, we used ChromoPainter^17^ to identify long, shared haplotypes among sequenced individuals using a reference panel enriched with Scandinavian populations (n=1,675 individuals, see Supplementary Notes 6 and 11). Our approach detects subtle population structure present during the VA in Scandinavia. Supplementary Figures S11.1-10 and Supplementary Note 11 describe the supervised method that we used to obtain power to robustly identify local ancestry variation in the presence of sequencing rate variation. We find at least four major ancestry components in Scandinavia, each with affinities with a present-day population (Fig. S11.11): a Danish-like, a Swedish-like, a Norwegian-like and a British-like component. Henceforth, we call this latter component ‘North Atlantic’, and we suspect it may reflect originally Celtic individuals that occupied the British Isles and were brought into Scandinavia. We refer to the first three ancestries as ‘Danish-like’, ‘Swedish-like’ and ‘Norwegian-like’, though we emphasize that the correspondence between these ancestries and present-day inhabitants of the respective Scandinavian countries is by no means exact or exclusive. During the VA, we mostly find high levels of Norwegian-like and Swedish-like components in Norway and Sweden, respectively, while Danish-like and ‘North Atlantic’ components are more widespread within Scandinavia (Fig. S11.12 and Supplementary Table 6). Notably, the ‘Swedish-like’ component is higher in Salme, Estonia, than in Sweden, because our sampling scheme included several individuals from the famous Salme Viking ship burial, of which archaeological and isotopic data suggest a Scandinavian origin^18,19^. While in general individuals from most of the Scandinavian VA settlements show mixed (Danish, Norwegian and Swedish) genetic ancestries, VA individuals from Jutland (Denmark) do not have significant Swedish-like or Norwegian-like genetic components. Furthermore, gene flow within Scandinavia appears to be broadly northwards, dominated by Danish Vikings moving into what are now Norway and Sweden (Table S11.2; see Supplementary Note 11).

Although the majority of the Viking genomes within Scandinavia and abroad show affinities to Danish, Norwegian, Swedish or British populations, there are some notable exceptions. We identified two ancient individuals (VK518 and VK519) originating from northern regions of Norway (Nordland), which have affinities to present-day Saami. This signal is weaker for VK519, indicating that he might have also had Norwegian-like ancestors. Given the geographic provenance of these samples, it was not unexpected to find individuals with Saami-like ancestry among the VA samples. However, as VK519 is indeed an admixed individual with both Norwegian-like and Saami-like ancestries, it appears that genetic contacts between these groups were already underway in VA Norway.

Importantly, present-day country boundaries are not always well reflected in the genetic data. Thus, the south-western part of Sweden in the VA is genetically more similar to Danish VA populations than the eastern regions of mainland Sweden (i.e. the area around the Mälaren Valley), likely due to geographic barriers that prevented gene flow in Sweden.

We quantified genetic diversity in our samples using two measures: conditional nucleotide diversity (Supplementary Note 9) and variation in inferred ancestry (Supplementary Note 11; Extended Data Fig. 5 and Fig. S11.13). We find overall high nucleotide diversity among most Viking-Age groups, with diversity values exceeding those of earlier Neolithic or BA groups, and only slightly lower than the highly diverse IA individuals from the British Isles (Fig. S9.1). Both measures of diversity vary significantly across locations. Denmark and Gotland in Sweden have the highest genetic diversity in the region, suggesting that these regions may have been centers of interaction and trade during this time. They also possess high diversity in inferred ancestries. North Norway also has high diversity in inferred ancestry due to its mixture of ‘North Atlantic’ and ‘Norwegian-like’ ancestry.

Interestingly, on Gotland, there are much more Danish-like, British-like and Finnish-like genetic components than Swedish-like components, supporting the notion that the island may have been marked by extensive maritime contacts during the VA. Our two Gotland sampling sites, Fröjel and Kopparsvik, have traditionally been argued to contain non-local individuals^20^, but recent Sr-isotope analyses have suggested otherwise^21,22^.

On Öland in Sweden, we observe high genetic diversity and the most variable patterns of recent ancestry (Extended Data Fig. 5) in Scandinavia. Sr and O isotope variation in these samples, and more contemporary samples from Öland have concluded that there is: (i) a high proportion (68%) of non-locals, (ii) high diversity in geographical origins and (iii) long distance migration^23^. Thus, the genetic diversity observed for Öland in the VA fits well with all of these results.

In conclusion, the results for Gotland and Öland agree with the archaeological record, suggesting that Öland and Gotland were important trading posts from the Roman period onwards^24,25^. A similar pattern is observed at a few archaeological sites from the central Danish islands, such as Langeland, although at a lower scale. Interestingly, genetic diversity here increases from the early to the late VA, suggesting increasing interregional interaction.

Our findings do not agree with the view of an overall highly connected population in Viking Scandinavia ^2,8,14–16^. Rather, we find clear genetic population structure within Scandinavia. We see evidence of a few cosmopolitan centers to the south, in southern Sweden and Denmark, where we see higher diversity of ancestries than in the rest of Scandinavia. These patterns are consistent with a restricted number of sea routes between the different Scandinavian areas and beyond.

### Viking migrations

Viking society is particularly famous for its ship technology, allowing for fast transport of large numbers of individuals in a single vessel^26^. These vessels enabled the Vikings not only to carry out lucrative raids and extended trade routes across Western Eurasia, but also to reach and settle lands in the North Atlantic^27–30^. Based on historical and archaeological data, Viking presence extended into both western and eastern Europe, reaching perhaps as far as the Pontic Steppe and the Middle East^31,32^. It is commonly believed that the westward migrations and raids were mainly carried out by people from what are now Norway and Denmark in the 9^th^ and 10^th^ centuries CE. In contrast to western movements, eastward expansions are commonly believed to have been carried out by Swedish Vikings, trading along navigable river systems and overland caravan routes^32^. Swedish Vikings (the ‘Rus’) are also credited for being active in the formation of the first Russian state^33,34^. Overall, our fine-scale ancestry analysis based on genomic data largely support the Viking expansion patterns inferred from archaeology (Figs. 3, 4 and S11.12). The eastward movements mainly involved individuals with Swedish-like ancestry, while the Viking individuals with Norwegian-like ancestry travelled to Iceland, Greenland, Ireland and the Isle of Man. A Danish-like ancestry component is more pronounced in present-day England, which is also in accordance with historical records^35^ and still visible in place-names^34^, and modern genetics^36,37^. Importantly, however, it is currently impossible for us to distinguish Danish-like ancestry in the British Isles from that of the Angles and Saxons, who migrated in the 5^th^-to-6^th^ centuries CE from Jutland and Northern Germany.

**Fig. 4:**
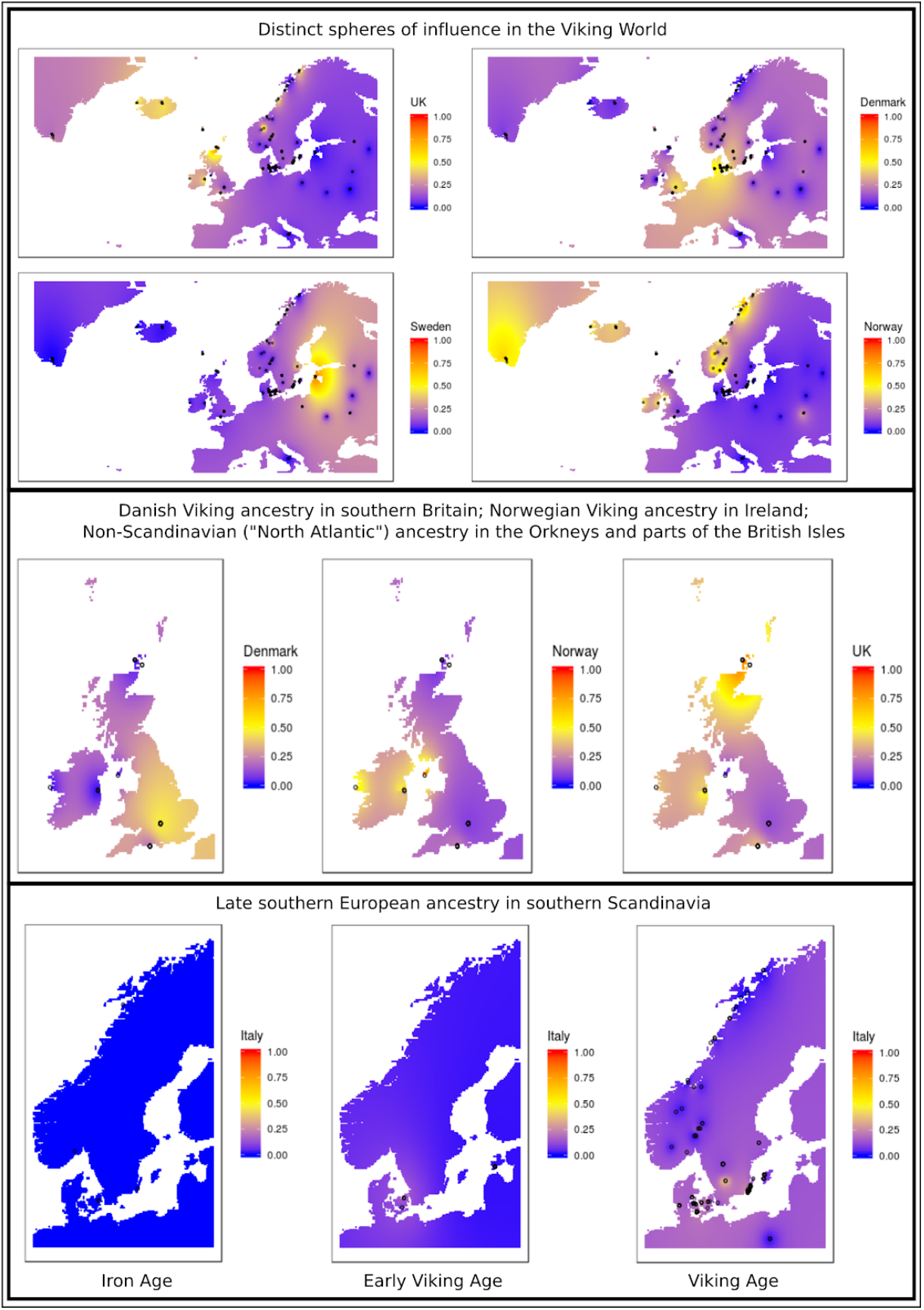
Spatiotemporal patterns of Viking and non-Viking ancestry in Europe during the IA, EVA and VA. UK = ‘British-like’ / ‘North Atlantic’ ancient ancestry component. Sweden = ‘Swedish-like’ ancient ancestry component. Denmark = ‘Danish-like’ ancient ancestry component. Norway = ‘Norwegian-like’ ancient ancestry component. Italy = ‘Southern European-like’ ancestry component. See Table S11.2 for statistical tests. The ‘Swedish-like’ ancestry is the highest in present-day Estonia due to the ancient samples from the Salme ship burial, which originated from the Mälaren Valley of Sweden, according to archaeological sources.

**Fig. 5:**
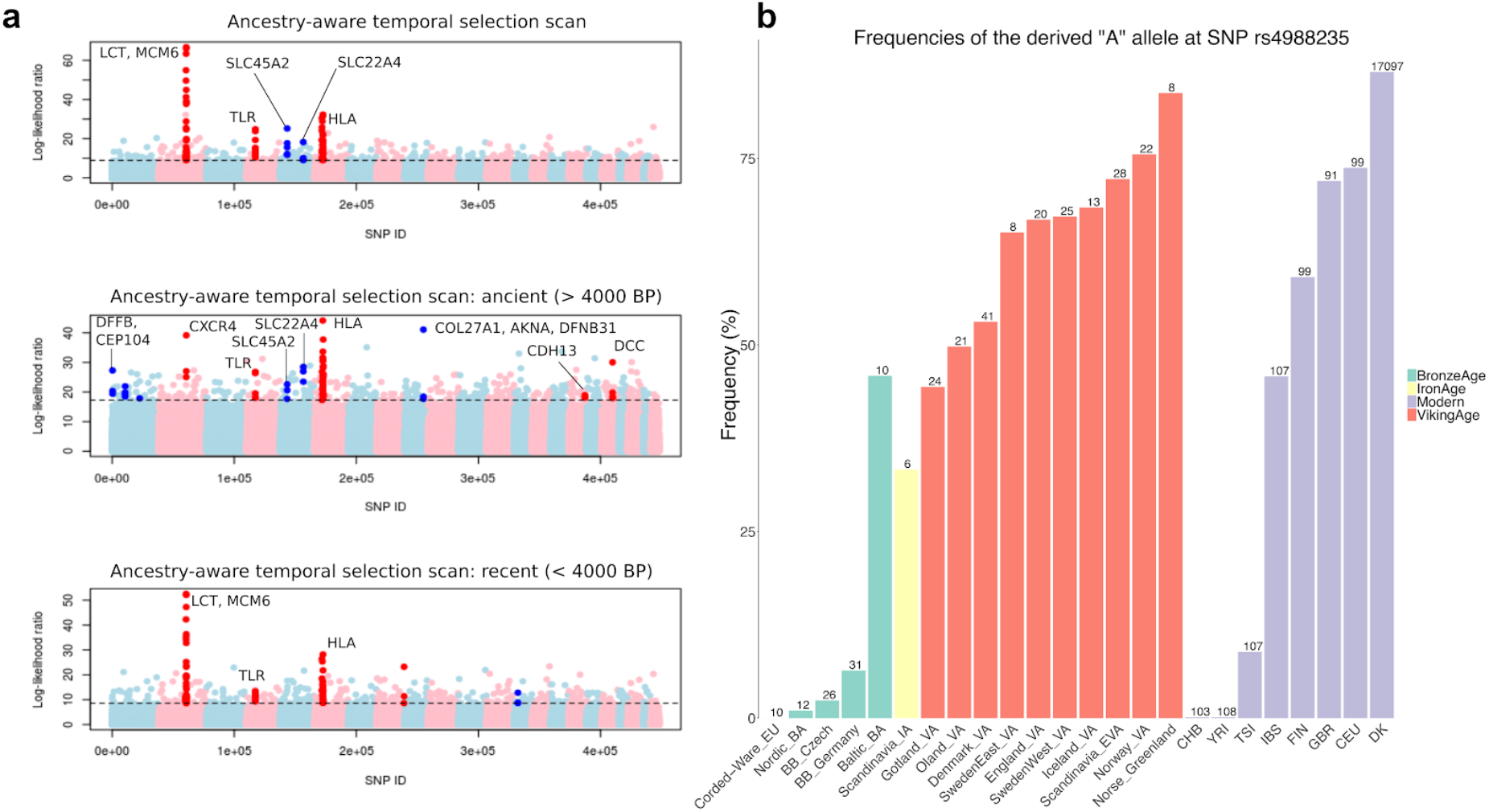
Positive selection in Europe. **a**, Manhattan plots of the likelihood ratio scores in favor of selection looking at the entire 10,000-year period (top, “general” scan), the period up to 4,000 BP (middle, “ancient” scan) and the period from 4,000 BP up to the present (bottom, “recent” scan). The highlighted SNPs have a score larger than the 99.9% quantile of the empirical distribution of log-likelihood ratios, and have at least two neighboring SNPs (+/− 500kb) with a score larger than the same quantile. **b**, Frequencies of the derived ‘A’ allele rs4988235 SNP responsible for lactase persistence in humans for different Viking-Age groups, present-day populations from the 1000 Genomes Project as well as relevant Bronze Age population panels. The numbers at the top of the bars denote the sample size on which the allele frequency estimates are based.

Interestingly, the ancient individuals from two execution sites in England (Dorset and Oxford) have significant local ‘North Atlantic’ ancestry, as well as Danish-like and Norwegian-like ancestries. If these represent Viking raiding parties coming to grief, as has been suggested^38,39^, this implies such forces were composed of individuals from different places of origin. This pattern is also suggested by isotopic data from the warrior cemetery in Trelleborg, Denmark^40^. Similarly, the presence of Danish-like ancestry in an ancient sample from Gnezdovo (Eastern Europe) indicates that the eastern migrations were not entirely composed of Vikings from Sweden.

However, in some cases, localities seem to have taken up Viking culture while incorporating little to no Scandinavian ancestry components, suggesting that the “Viking” identity was not always necessarily associated with Scandinavian genetic ancestry. Archaeological evidence indicates that the six higher coverage VA individuals from three different archaeological sites in the Orkney Islands have Scandinavian cultural links^41,42^. However, four (VK201, VK202, VK203 and VK207) of these samples have over 85% “UK” ancestry and are genetically similar to present-day Irish and Scottish populations (Figs 3a and S10.1, Supplementary Table 6), which is in contrast to the isotopic evidence^43^. Haplotype-based analyses corroborate that four of these samples possessed local genetic ancestries, with little Scandinavian contribution. Only two individuals - VK204 and VK205 - displayed c. 50% Norwegian and Danish ancestries (Supplementary Table 6), respectively, which may indicate admixture between the locals and Scandinavians on the Orkney Islands during the VA. The four ancient genomes of Orkney individuals with little Scandinavian ancestry may be the first ones of Pictish people published to date (Supplementary Note 12). Yet a similar (>80% “UK” ancestry) individual was found in Ireland (VK545) and five in Scandinavia, implying that Pictish populations were integrated into Scandinavian culture by the Viking Age.

### Gene flow into Scandinavia during the Viking era

Archaeological findings and the written sources support the hypothesis that Viking back migrations and interaction between the newly settled areas and Scandinavia occurred as part of the process^44^. Presumably, if these migrations took place, native ancestry from these areas must have also been introduced into Scandinavia. We therefore aimed to assess the levels of non-Scandinavian ancestry emerging in Scandinavia during the VA.

Using fineStructure^17^, we find that the levels of non-Scandinavian ancestry in the Danish, Norwegian and Swedish Vikings agree with known trading routes (Supplementary Notes 11 and 12). The most obvious genetic signals are from Finnish and Baltic sources into the area of what is now modern Sweden, including Gotland. These ancestries are present at considerably lower levels or are completely absent in most individuals from Denmark and Norway. A substantial interaction across the Baltic Sea is also suggested by objects from Finland found in graves in Middle Sweden, albeit recent Sr-isotope analyses are inconclusive regarding the origin of the buried individuals ^45,46^. In comparison, western regions of Scandinavia have much higher levels of ancestry from the British Isles, in comparison with the eastern regions of Sweden (Supplementary Notes 11 and 12). We also observe several individuals (Supplementary Table 6) with large amounts of South European ancestry in Denmark and southwest Sweden during the Viking period (Fig. 4). No such individuals are found among our Scandinavian Iron Age samples, though we stress that our sampling for this period is more limited than for the other two. The discovery of individuals with ancestry from Southern Europe and the British Isles is the first direct evidence for movement into Scandinavia from these regions. The directions of interaction marked by these individuals is consistent with the major directions of gene flows outwards from Scandinavia also seen in the data.

Surprisingly, three individuals from the Kärda site show much higher genetic similarity to Late Neolithic/Early Bronze Age Danish individuals than to all other VA individuals in the dataset. The site is located far inland, in south-west Sweden. This similarity is quite unexpected, given that the samples are AMS-dated to the middle of the VA, and consistent with the presence of Caucasus-related ancestry inferred in the qpAdm ancestry modelling. Studies of VA burial customs suggest that the Småland area was characterized by locally confined cultural groups^47^. The genetic data suggest that this pattern of cultural isolation was sustained in marked contrast to contemporary coastal and island communities. Consistent with this hypothesis we find that the individuals from Kärda show a marked reduction in nucleotide diversity compared to other VA groups (Fig. S9.1), although they also have high amounts of Southern European ancestry.

### Disappearance of the Greenlandic Norse

From around 980 to 1440 CE South-west Greenland was settled by peoples of Scandinavian (Norse) descent. They likely originated from Icelandic Vikings who established a colony there at the end of the 9^th^ century CE^29,48^. It is believed that the Norse also reached Labrador, North America, from Greenland around 1000, although no permanent settlement was established^30^. The fate of the Norse in Greenland remains debated, but probable causes of their disappearance are social or economic processes in Europe (e.g. political relations within Scandinavia and changed trading systems) and natural processes, like climatic changes^29,49,50^.

We see no evidence of long-term inbreeding in the Greenlandic Norse genomes, though we note that we only have one high-coverage genome from the later period of occupation of Greenland (Supplementary Note 10; Figs. S10.2 and S10.3). This suggests a depopulation scenario over approximately 100 years which would be in line with previous demographic models^51^, as well as the archaeology. Indeed, the latter indicates that marginal farms in the Western Settlement and the northern and southern parts of the Eastern Settlement were abandoned from about 1200 CE, with no converse intensification of settlement in the central areas.

We also find no evidence of ancestry from local populations from the Western Atlantic (Paleo Eskimo, Inuit or Native American) in the Norse genomes. This is in accordance with previous physical anthropological studies of the skeletal remains^51^. This suggests that either sexual interactions did not take place or that, if they did, then on a very small and incidental scale with the children remaining in the native communities. In terms of genetic ancestry of the Greenlandic Norse, we find evidence of admixture between Scandinavians (mostly from Norway) and individuals from the British Isles, similar to the first settlers of Iceland^52^, which supports the archaeological and historical links between the Greenlandic Norse and the Icelandic Vikings.

### Genetic composition of the earliest Viking expedition and kinship findings

Maritime raiding has been a constant of seafaring cultures for millennia. However, the VA is unusual in that it is partly defined by such activity^53^. Despite the historical importance of Viking raiding, the exact nature and composition of these war parties is unknown^5^. Only one Viking raiding or diplomatic expedition has left direct archaeological traces, at Salme in Estonia, where 41 Swedish Vikings who died violently were buried in two boats accompanied by high-status weaponry^18,19^. Importantly, the Salme boat-burial predates the first textually documented raid (in Lindisfarne in 793) by nearly half a century.

Comparing the genomes of 34 individuals from the Salme burial using kinship analyses, we find that these elite warriors included four brothers buried side by side and a 3rd degree relative of one of the four brothers (Supplementary Note 4). In addition, members of the Salme group had very similar ancestry profiles, in comparison to the profiles of other Viking burials (Supplementary Notes 10 and 11). This suggests that this raid was conducted by genetically homogeneous people of high status, including close kin. Isotope analyses indicate that the crew descended from the Mälaren area in Eastern Sweden^19^ thus confirming that the Baltic-Mid-Swedish interaction took place early in the VA.

Intriguingly, we identified several additional pairs of kin among the other Viking genomes. One is a pair of 2^nd^ degree male relatives (i.e. half-brothers, nephew-uncle or grandson-grandfather) from two locations separated by the North Sea: one of the samples (VK279) was excavated in Denmark (Galgedil site on Funen; this cemetery was also analyzed for strontium with a group of non-locals there) while the other individual (VK144) was found in the UK (Oxford site). Another pair of individuals with 3^rd^ or 4^th^ degree relatedness (e.g. cousins) was discovered in Sweden, namely a male sample excavated on the island of Öland (VK342) and a female individual from Skämsta, Uppsala (VK527), some 300-400 kilometers apart. Interestingly, the female from Uppsala (VK527) also had a brother (VK517), and both siblings display a rare genetic disorder of abnormal skeletal development: spondyloepiphyseal dysplasia. Given the very low frequency of this disorder, the close family ties between these individuals were expected by the archaeologists^22^. Such long-distance relationships in our dataset underscore the degree of individual-level mobility during the Viking era.

### Positive selection in Europe in the last 10,000 years

The availability of hundreds of genomes from the IA and VA - in combination with previously published Mesolithic, Neolithic and Bronze Age genomes^10,11,54,55^ - permit us to directly investigate the role of positive selection using time series of allele frequencies from the last ten millennia of European history. We looked for SNPs whose allele frequencies changed significantly in the last 10,000 years, using a newly developed method called “neoscan” that is implemented in the Ohana software package^56,57^, and that can detect strong allele frequency shifts in time that cannot be explained by temporal changes in genome-wide genetic ancestry alone (Supplementary Note 14). Figure 5a shows the resulting likelihood ratio scores in favor of selection looking at the entire 10,000-year period (top, “general” scan), the period up to 4,000 BP (middle, “ancient” scan) and the period from 4,000 BP up to the present (bottom, “recent” scan). The strongest candidate for selection - especially in the “recent” scan - is a cluster of SNPs near the LCT gene - a signal that has been extensively characterized in the past^58,59^. The rise in frequency of the lactase persistence allele to its present-day levels in Northern Europe is, however, poorly understood. We know that this rise must have occurred after the Bronze Age, a time at which this allele was still segregating at low frequencies^10,54^. Based on the archaeological record, we also know that VA Scandinavians used a variety of dairy products as an essential part of their daily food intake. Our dataset allows us, for the first time, to directly assess the frequency of the lactase persistence allele (at SNP rs4988235, upstream of the LCT gene) in Scandinavia during the Iron Age and VA, and trace its evolution since the Bronze Age.

Figure 5b shows that VA groups had very similar allele frequencies at the LCT lactase persistence SNP to those found in present-day northern European populations. In contrast, the persistence allele was at low frequencies in Bronze Age Scandinavians, as well as Corded Ware and Bell Beaker cultures from central Europe, even though there is evidence for milk consumption in these regions by that time. The allele frequency in Iron Age samples is at intermediate levels (c. 37.5%), suggesting this rise in frequency must indeed have occurred during the Iron Age (c. 1500-2500 years ago), but was largely complete at the onset of the VA. Interestingly, the allele frequency of the allele is much higher (c. 40%) in the Bronze Age population from the neighboring Baltic Sea region than in Bronze Age Scandinavia. Given the geographic and cultural proximities between Scandinavia and the Baltic region, this may suggest gene flow between the two regions resulting in increased frequency of lactase persistence in Scandinavia during the Iron Age.

Other candidates for selection include previously identified regions like the TLR1/TLR6/TLR10 region, the HLA region, SLC45A2 and SLC22A4^54^. We also find several new candidate regions for selection in the “ancient” scan, some of which contain SNPs where the selected allele rose in frequency early in the Holocene but then decreased later on (Supplementary Note 14). These candidate regions include a region overlapping the DCC gene, which has been implicated in colorectal cancer^60^ and another overlapping the AKNA gene, which is involved in the secondary immune response by regulating CD40 and its ligand^61^. This highlights the utility of using ancient DNA to detect signatures of selection that may have been erased by recent selective dynamics.

### Pigmentation-associated SNPs

Exploring twenty-two SNPs with large effect associated with eye color and hair pigmentation, we observe that their frequencies are very similar to those of present-day Scandinavians (Supplementary Note 13). This suggests that pigmentation phenotype in VA Scandinavians may not have differed much from the present-day occupants of the region (although see section on complex traits below for an analysis including alleles of small effect). Nevertheless, it is important to stress that there is quite a lot of variation in the genotypes of these SNPs across the sequenced samples, and that there is therefore not a single ‘Viking phenotype’. For example, two of the ancient samples with the highest coverage have different pigmentation phenotypes: VA individual VK42 from Skara, Sweden has alleles associated with brown eyes and darker hair coloration while VK1 from Greenland was likely to have had blue eyes and lighter hair.

### Evolution of complex traits in Scandinavia

To search for signals of recent population differentiation of complex traits, we compared genotypes of Viking age samples with those of a present-day Scandinavian population for a range of trait-associated SNP markers. We selected 16 traits for which summary statistics from well-powered genome-wide association studies (GWAS) were available through the GWAS ATLAS (https://atlas.ctglab.nl)^62^. For comparison with the Viking age samples we used a random population subset of the IPSYCH case-cohort study of individuals born in Denmark between 1981-2011^63^. We derived polygenic risk scores (PRS) for the 16 traits, based on independent genome-wide significant allelic effects and tested for a difference in the distribution of polygenic scores between the two groups, correcting for sex and ancestry-sensitive principal components (Supplementary note S15). We observed a significant difference between the polygenic scores of VA samples and current Danish population samples for three traits; black hair color (P = 0.00089), standing height (P = 0.019) and schizophrenia (P = 0.0096) (Extended Data Fig. 5). For all three traits, the polygenic score was higher in the VA group than in the present-day Danish group. The observed difference in PRS for height and schizophrenia between the groups did however not remain significant after taking into account the number of tests. A binomial test of the number of black hair color risk alleles found in higher frequency in the VA sample and the present-day sample, also returned a significant difference (65/41; P = 0.025), which suggests that the signal is not entirely driven by a few large-effect loci.

Thus, we only find evidence for systematic changes in combined frequencies of alleles affecting hair color (and possibly also height and schizophrenia), among all the anthropometric traits and complex disorders we tested. Also, we cannot conclude whether the observed difference in allele frequencies are due to selection acting on these alleles between the Viking Age and the present time or to some other factors (such as more ethnic diversity in the VA sample), nor can we conclude whether a similar change has occurred in other Nordic populations than the Danish.

### Genetic legacy of the Vikings in present-day populations

To test whether present-day Scandinavians share increased ancestry with their respective ancient Viking counterparts, we first inferred D-statistics of the form D(YRI, ancient; present-day X, present-day DK), which contrast allele sharing of a test ancient individual with a present-day test population X and present-day Danes. We find subtle but noticeable shifts of ancient individuals towards their present-day counterparts in the distributions of these statistics (Extended Data Fig. 3). We further examined variation in present-day populations using fineSTRUCTURE, and then described these present-day groups by their ancestry from ancient populations (Fig. S11.14).

We find that within Scandinavia, present-day populations are still structured according to the ancient Viking population groups. The component that we associated as Norwegian-like is present at 45-65% in present-day Norway. Similarly, the ancient Swedish-like ancestry is present at 15-30% within Sweden. Of the four Swedish clusters, one is more related to the ancient Finnish than the Swedish-like ancestry, and a second is more related to Danes and Norwegians. Danish-like ancestry is now high across the whole region.

Outside of Scandinavia, the genetic legacy of the Vikings is consistent, though limited. A small component is present in Poland (up to 5%) and the south of Europe. Within the British Isles, it is difficult to assess how much of the Danish-like ancestry is due to pre-existing Anglo-Saxon ancestry; however, the Norwegian-like ancestry is consistently around 4%. The Danish-like contribution is likely to be similar in magnitude and is certainly not larger than 16% as found in Scotland and Ireland. The lack of strong variation in ancestry from Scandinavia makes sense if the Vikings did not maintain a diaspora identity over time but instead integrated into the respective societies in which they settled. The genetic impacts are stronger in the other direction. The ‘British-like’ populations of Orkney became ‘Scandinavian’ culturally, whilst other British populations found themselves in Iceland and Norway, and beyond. Present-day Norwegians vary between 12 and 25% in their ‘British-like’ ancestry, whilst it is still (a more uniform) 10% in Sweden. Separating the VA signals from more recent population movements is difficult, but these numbers are consistent with our VA estimates.

## Discussion

Until now, our main understanding of the VA was largely based on a combination of historical sources and archaeological evidence. These often characterize the VA as a period of high mobility and interaction between peoples. Networks of trade were established, connecting distant regions within Scandinavia through established waterways with significant movement between regions. It has also been viewed as a time where links were created to regions outside Europe, from the Pontic Steppe in the east to North America in the west.

Our genomic analyses add complex layers of nuance to this simple picture. We largely reconstruct the long-argued movements of Vikings outside Scandinavia: Danish Vikings going to Britain, Norwegian Vikings moving to Ireland, Iceland, and Greenland, and Swedish Vikings sailing east towards the Baltic and beyond. However, we also see evidence of individuals with ancient Swedish and Finnish ancestry in the westernmost fringes of Europe, whilst Danish-like ancestry is also found in the east, defying our modern notions of historical groupings. It is likely that many such individuals were from communities with mixtures of ancestries, likely thrown together by complex trading, raiding and settling networks that crossed cultures and the continent.

Our observations also suggest that the different parts of Scandinavia were not as evenly connected, as has often been assumed. Despite relatively fast and easy communication between the coastal regions of Denmark, Norway, and Sweden, we find that clear genetic structure was present in Viking-Age Scandinavia. In fact, our data indicate that Viking Scandinavia consisted of a limited number of transport zones and maritime enclaves^64^ where contact was made with Europe, while the remaining regions had limited external gene flow with the rest of the Scandinavian continent. Some Viking-Age Scandinavian locations are relatively homogeneous both in terms of genetic diversity and patterns of ancestry; particularly mid-Norway, Jutland, and the Atlantic settlements, which contain predominantly Norwegian-like and ‘North Atlantic’ (including pre-Anglo Saxon British) ancestry. Indeed, one of the clearest vectors of contrast observed in this study is between the strong genetic variation seen in relatively populous coastal trading communities such as in the islands Gotland and Öland, and the reduced diversity in less populated (mostly inland) areas in Scandinavia. Such high genetic heterogeneity, which was likely due to increased population size, extends the urbanization model of Late Viking Age city of Sigtuna proposed by Krzewińska et al.^6^ both spatially and further back in time.

Interestingly, our findings correspond with paleodemographic studies based on place-name evidence and archaeological distributions suggesting population density was higher in Denmark than elsewhere in Viking-Age Scandinavia^65^. Gene flow from Denmark to the north is also paralleled by the linguistic affinities of the medieval Scandinavian languages: The 12th-century Icelandic law text Grágás states that the common language of Swedes, Norwegians, Icelanders, and Danes was dǫnsk tunga (‘Danish tongue’)^66^. It appears that the formation of large-scale trading and cultural networks that spread people, goods and warfare took time to affect the heartlands of Scandinavia, which received much more restricted gene flow, retaining pre-existing genetic differences between Scandinavian populations. This pattern of behavior seems to prevail from the beginning of the Viking diaspora to its end at the beginning of the medieval period.

Our findings also show that Vikings are not simply a direct continuation of the Scandinavian Iron Age groups. Rather than simple continuity, we observe foreign gene flow from the south and east into Scandinavia, starting in the Iron Age, and continuing throughout the duration of the Viking period from an increasing number of sources. Our findings also contradict the myth of the Vikings as peoples of pure local Scandinavian ancestry. In fact, we found many Viking Age individuals with high levels of foreign ancestry, both within and outside Scandinavia, suggesting ongoing gene flow with different peoples across Europe. Indeed, it appears that some foreign peoples contributed more genetic ancestry to Scandinavia during this period than the Vikings contributed to them which could partially be due to smaller effective population size of the VA Scandinavians as opposed to their continental and British neighbors.

## Acknowledgements

This work was supported by the Mærsk Foundation, the Lundbeck Foundation, the Novo Foundation, the Danish National Research Foundation, KU2016, and the Wellcome Trust (grant nos. WT104125MA). The authors thank the iPSYCH Initiative, funded by the Lundbeck Foundation (grant nos. R102-A9118 and R155-2014-1724), for supplying SNP frequency estimates from the present-day Danish population for comparison with Viking Age samples. EW would like to thank St. John’s College, Cambridge for providing an excellent environment for scientific thoughts and collaborations. SR was supported by the Novo Nordisk Foundation (NNF14CC0001). FR was supported by a Villum Young Investigator Award (project no. 000253000). The authors are grateful to Marisa Corrente for providing access to the skeletal remains from Cancarro and Nunzia M. Mangialardi and Marco Maruotti for the useful suggestion. G.S. and E.C. were supported by a Marie Skłodowska-Curie Individual Fellowship “PALAEO-ENEO”, a project funded by the European Union EU Framework Programme for Research and Innovation Horizon 2020 (Grant Agreement number 751349). RM was supported by an EMBO Long-Term Fellowship (ALTF 133-2017). We thank Mattias Jakobsson and Anders Götherström for providing preliminary access to the sequencing data of 23 Viking Age samples from Sigtuna; L. Vinner, A. Seguin-Orlando, K. Magnussen, L. Petersen and C. Mortensen at the Danish National Sequencing Centre for producing the analysed sequences; P.V. Olsen and T. Brand for technical assistance in the laboratories. We thank Richard M. Durbin and James H. Barrett for comments and suggestions.

## Extended Data Figures

**Extended Data Fig. 1:**
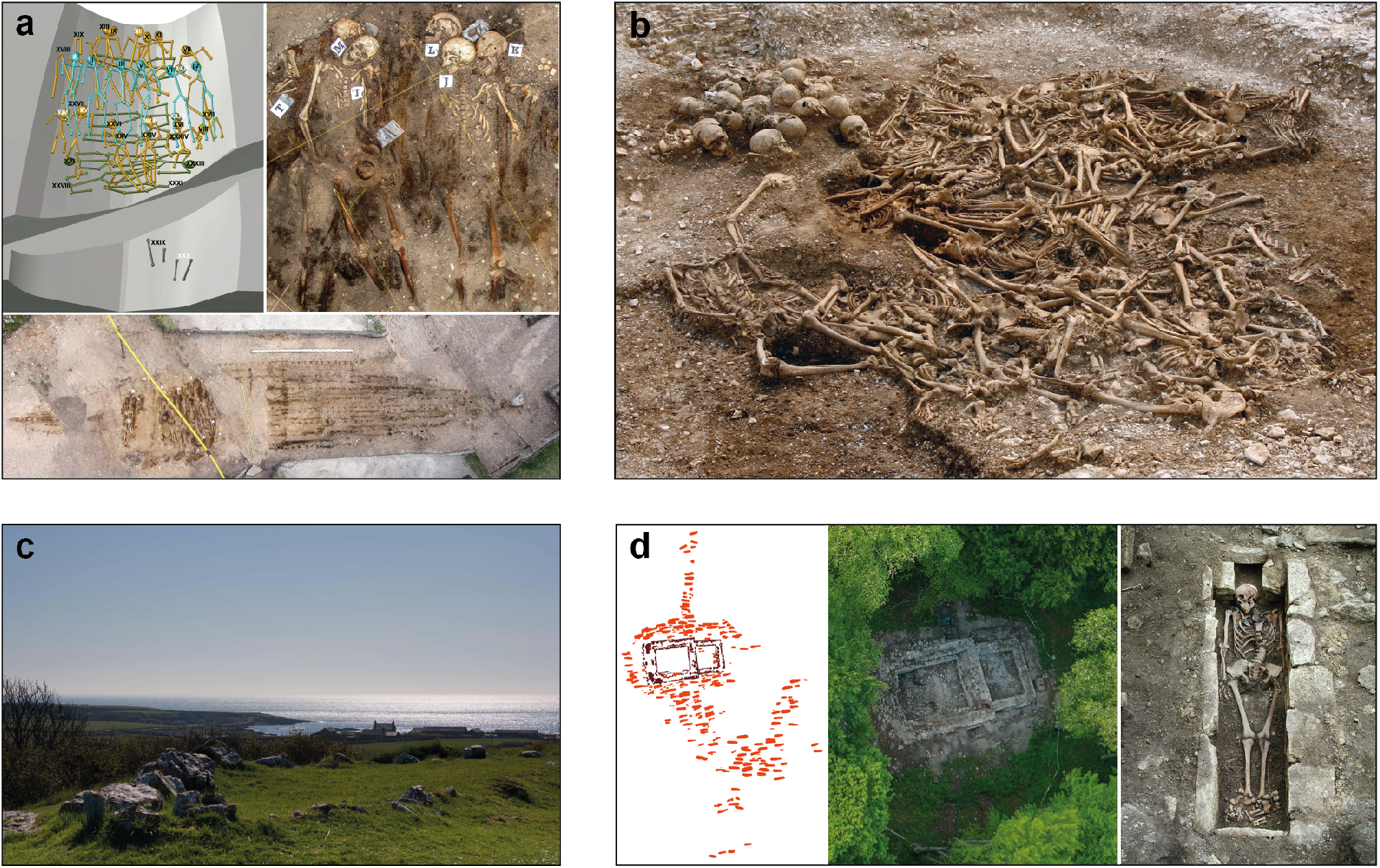
Viking Age archaeological sites. Examples of a few archaeological Viking Age sites and samples used in this study. **a,** Salme II ship burial site of Early Viking Age excavated in present-day Estonia: schematic representation of skeletons (upper left-hand corner image) and aerial images of skeletons (upper right-hand corner and lower images). **b,** Ridgeway Hill mass grave dated to the 10th or 11th century, located on the crest of Ridgeway Hill, near Weymouth, on the South coast of England. Around 50 predominantly young adult male individuals were excavated. **c,** The site of Balladoole: around AD 900, a Viking was buried in an oak ship at Balladoole, Arbory in the south east of the Isle of Man. **d,** Viking Age archaeological site in Varnhem, Sweden: Schematic map of the church foundation (left) and the excavated graves (red markings) at the early Christian cemetery in Varnhem; foundations of the Viking Age stone church in Varnhem (middle) and the remains of a 182 cm long male individual (no. 17) buried in a lime stone coffin close to the church foundations (right).

**Extended Data Fig. 2:**
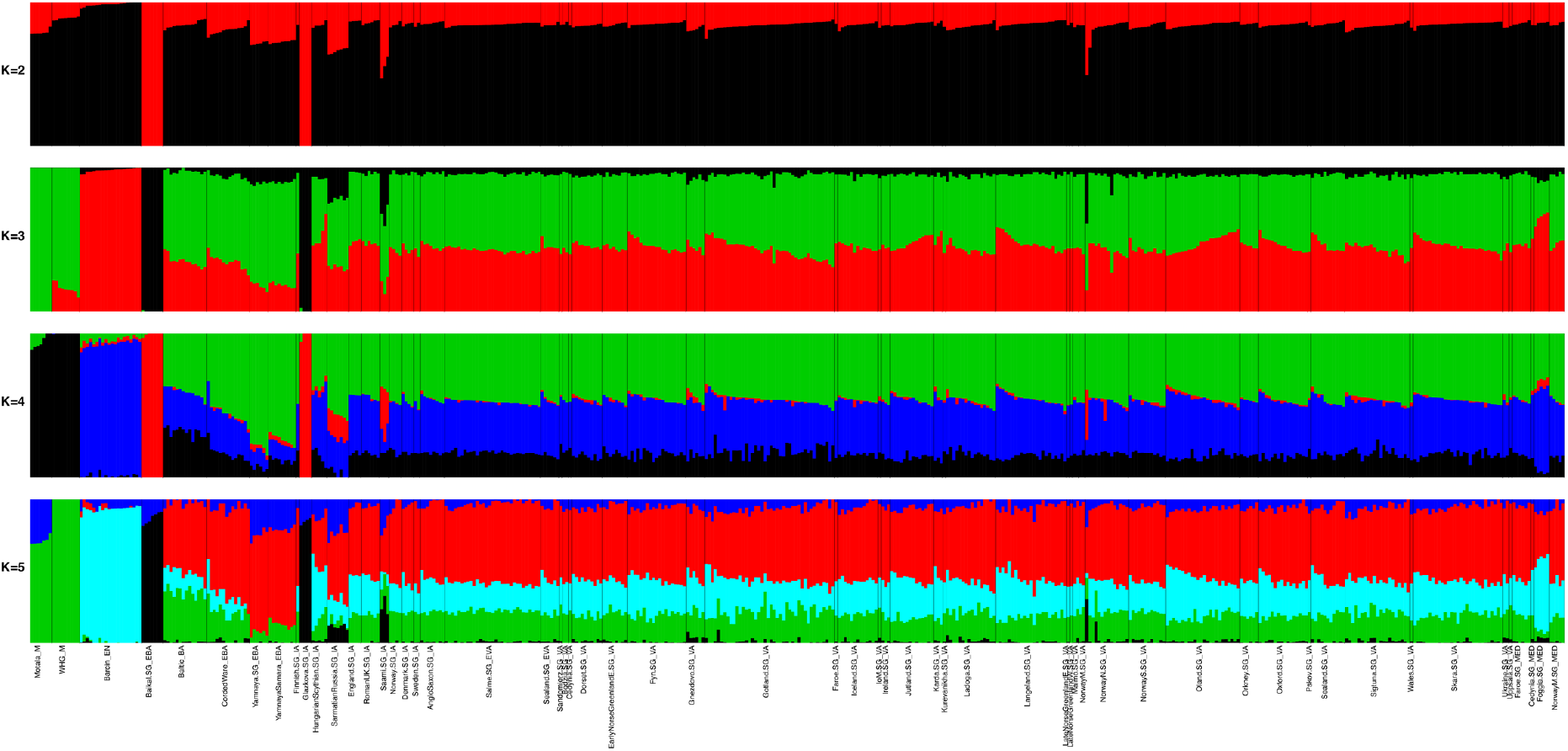
Model-based clustering analysis. Admixture plot (K=2 to K=5) for 517 ancient individuals spanning 60 different populations. This figure is a subset of most relevant individuals and populations from Figure S7.2, see Supplementary Note 7 for details.

**Extended Data Fig. 3:**
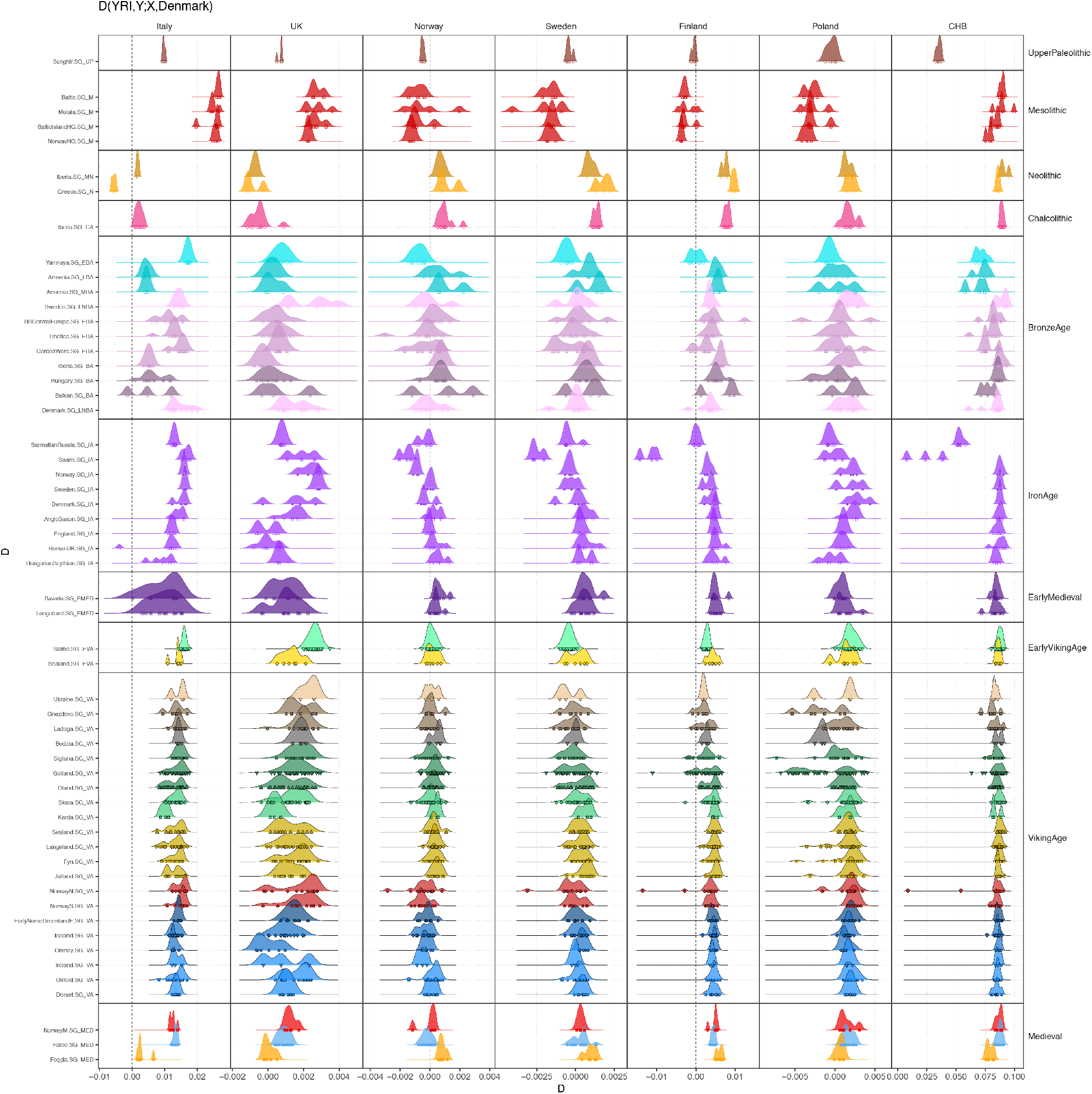
Symmetry tests of genetic affinity of ancient individuals with contemporary populations. Panels show D-statistics of the form D(YRI,Y; X,Denmark), which contrast allele sharing of an ancient individual Y with either contemporary population X or Denmark. Plot symbols show point estimates, and density plots distributions across all individuals per analysis group.

**Extended Data Fig. 4:**
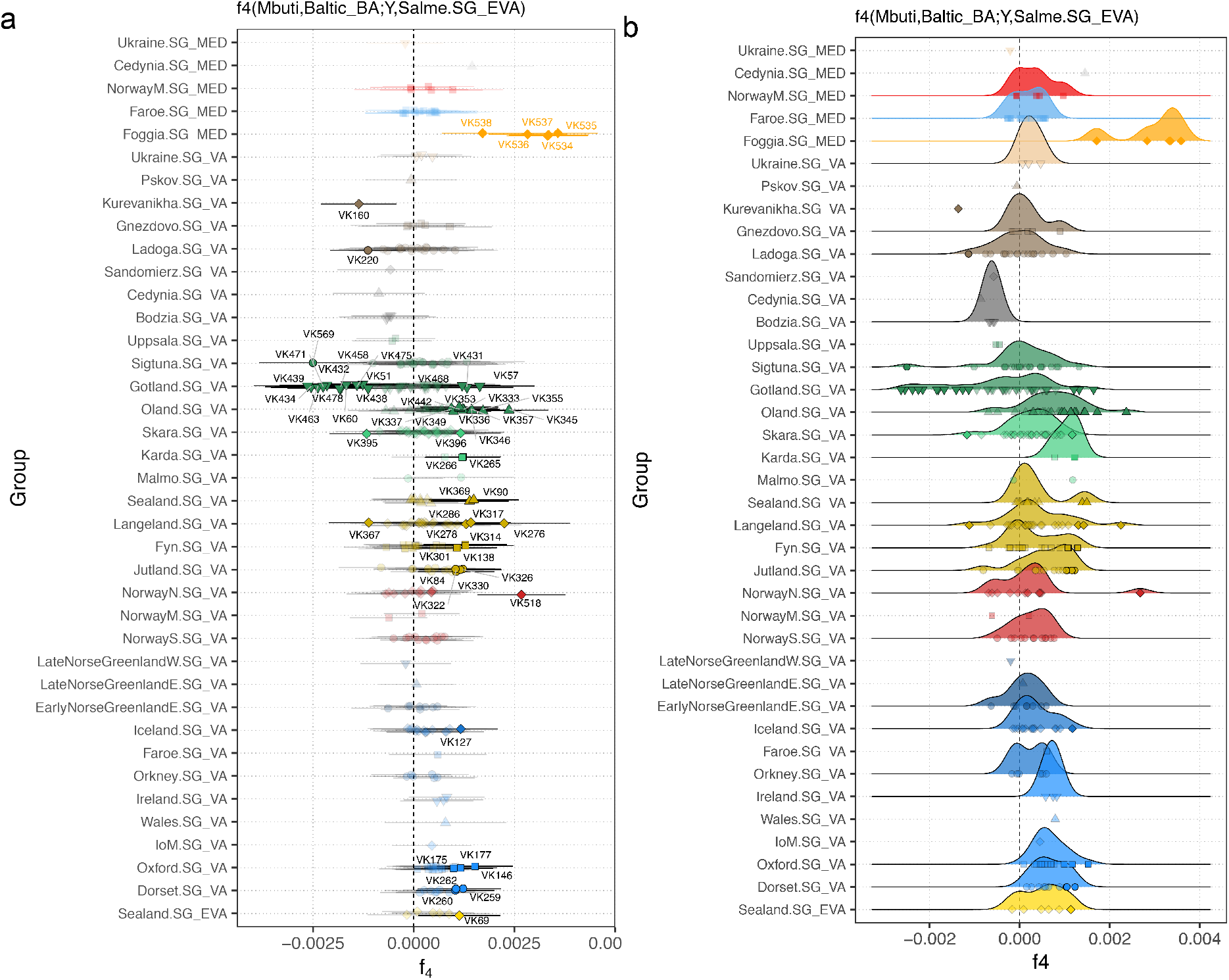
Symmetry tests for genetic affinity with Baltic Bronze Age. Panels show *f*_4_-statistics of the form *f*_4_(Mbuti,Baltic_BA;Y, Salme.SG_EVA), which contrast allele sharing of Baltic_BA with either a test individual Y or Salme.SG_EVA. **a,** point estimates and error bars (± 3 standard errors) for each target individual, aggregated by analysis group. Individuals with significant *f*_4_-statistics (|Z| ≥ 3) are indicated without transparency and respective sample IDs. **b,** as in **(a)**, with density plot for distributions across all individuals per analysis group.

**Extended Data Fig. 5:**
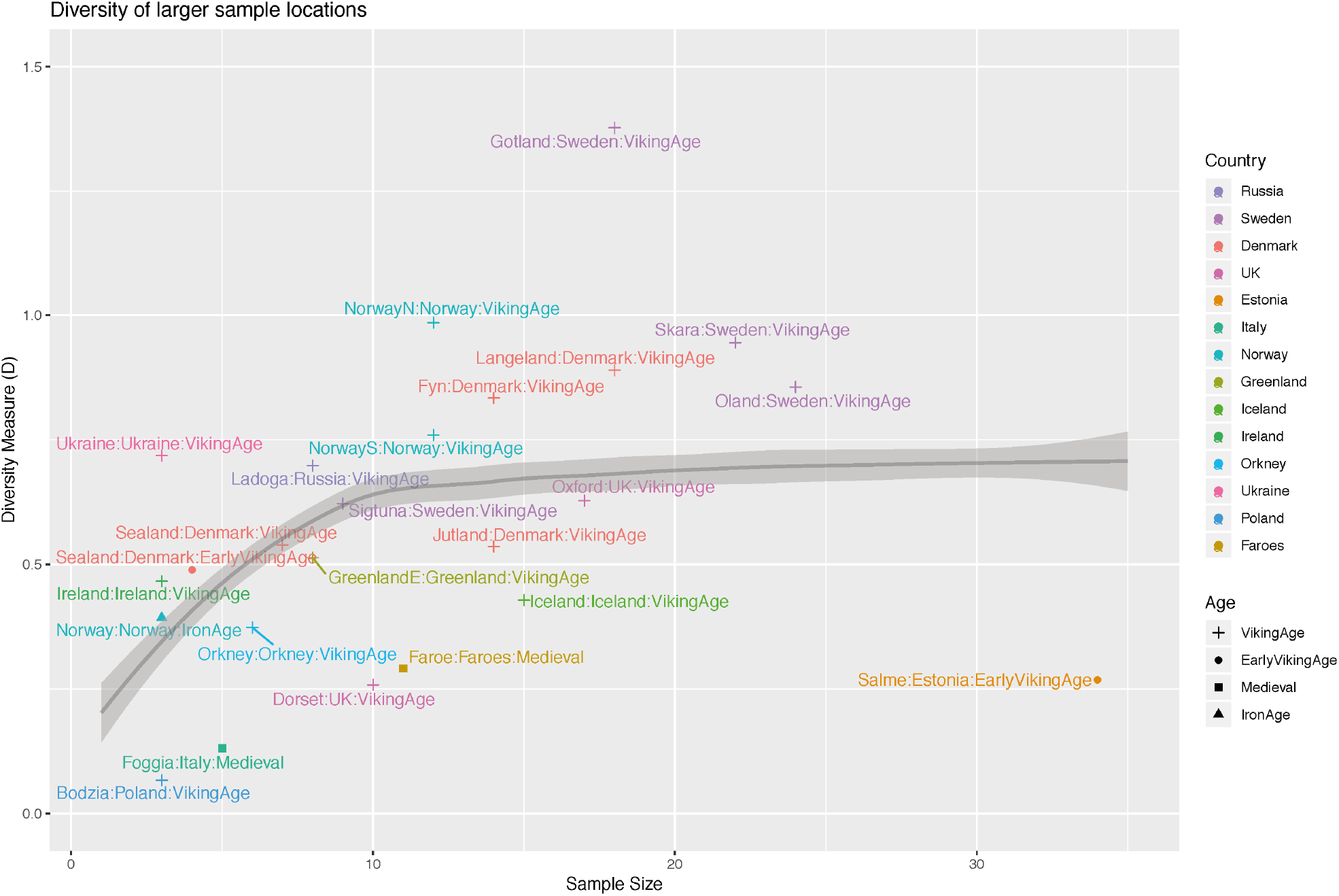
Ancestry diversity of different population groups. Diversity of different labels (i.e. sample locations combined with historical age) are shown as a function of their sample size. The Diversity measure is the Kullback-Leibler divergence from the label means, capturing the diversity of a group with respect to the average of that group; see text for details. Larger values are more diverse, though a dependence on sample size is expected. The simulation expectation for the best-fit to the data (0=0.2) is shown.

**Extended Data Fig. 6:**
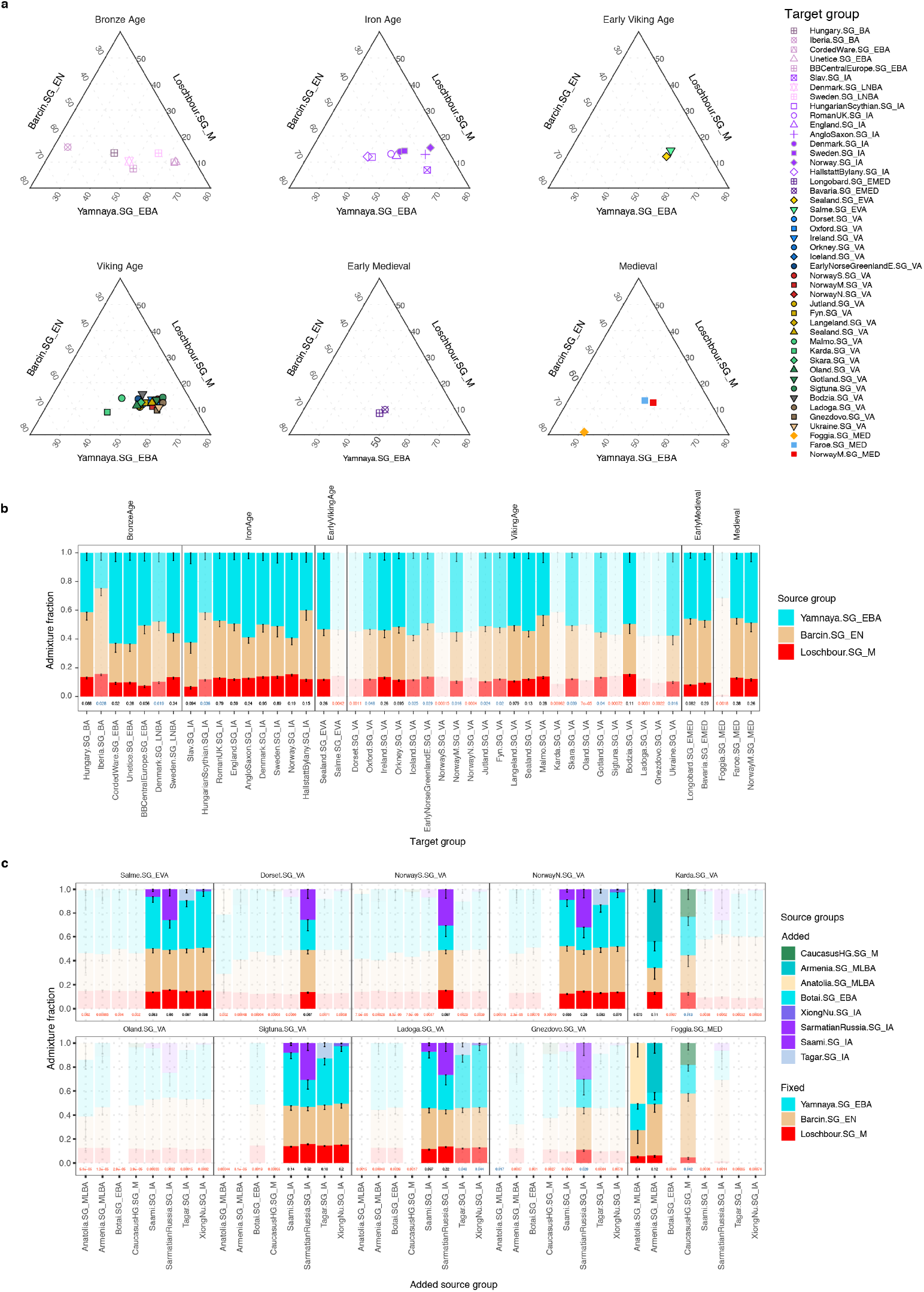
Ancestry modelling using *qpAdm*. **a,** Ternary plots of ancestry proportions for a three-way model of Mesolithic hunter-gatherer (Loschbour.SG_M), Neolithic farmer (Barcin.SG_EN) and Bronze Age Steppe herders (Yamnaya.SG_EBA). **b,** Bar plots with ancestry proportions as in **(a)**, with error bars indicating standard errors and transparency/text colors indicating p-value for model fit (no transparency/black: p ≥ 0.05; light transparency/blue: 0.05 > p ≥ 0.01; strong transparency/red: p ≤ 0.01). **c,** Ancestry proportions of four-way models including additional putative source groups for target groups for which three-way fit was rejected (p ≤ 0.01); transparency/text colors as in **(b)**

**Extended Data Fig. 7:**
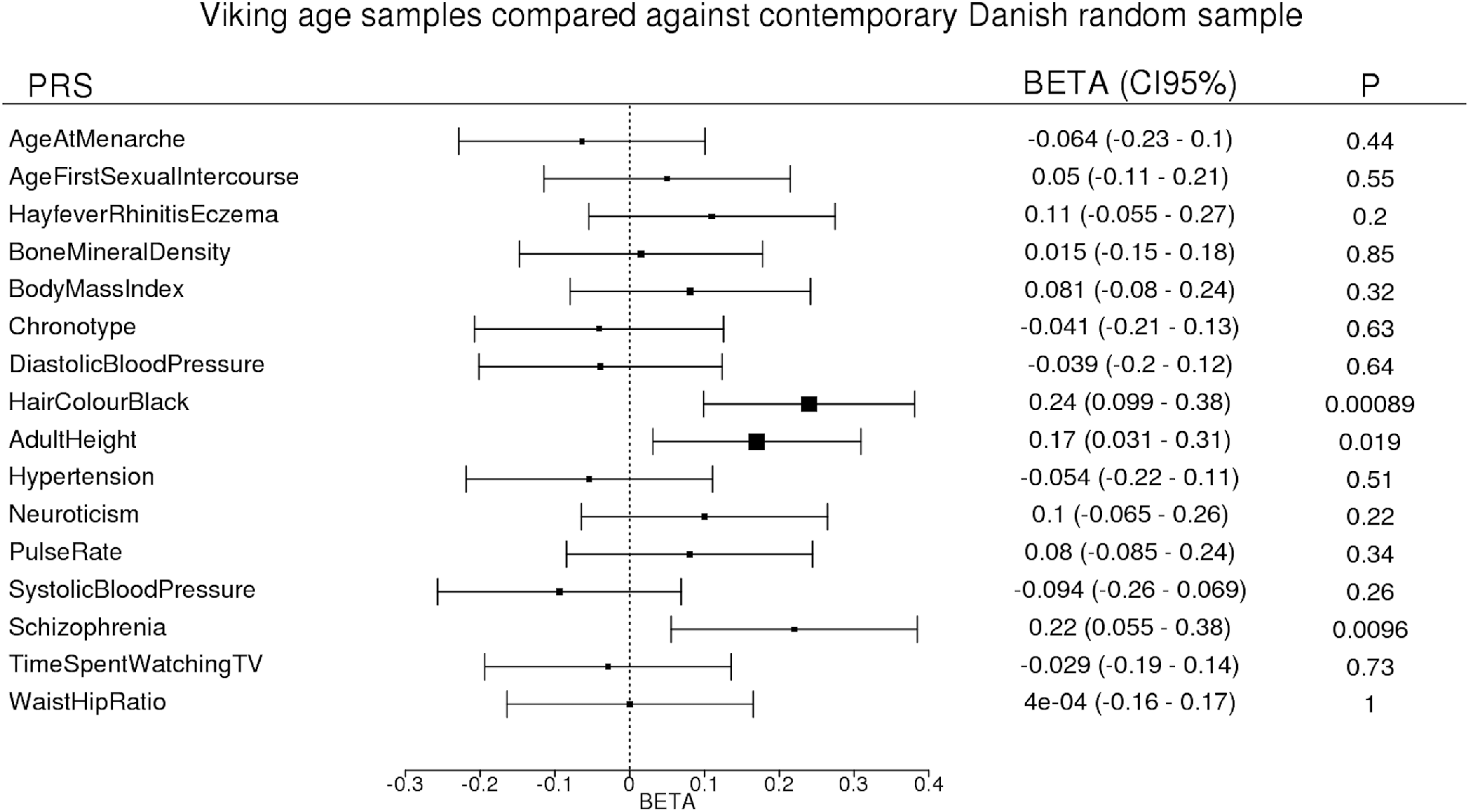
Polygenic risk scores. Polygenic risk scores (PRS) for 16 complex human traits in Viking Age samples from Denmark, Sweden and Norway compared against a reference sample of >20,000 Danish-ancestry individuals randomly drawn from all individuals born in Denmark in 1981-2011. The PRS is in each case based on allelic effects for >100 independent genome-wide significant SNPs from recent GWAS of the respective traits. Only PRS for black hair colour is significantly different between the groups after taking account of multiple testing, although PRS for height and schizophrenia are considerably elevated as well in the Viking Age samples.

**Extended Data Fig. 8:**
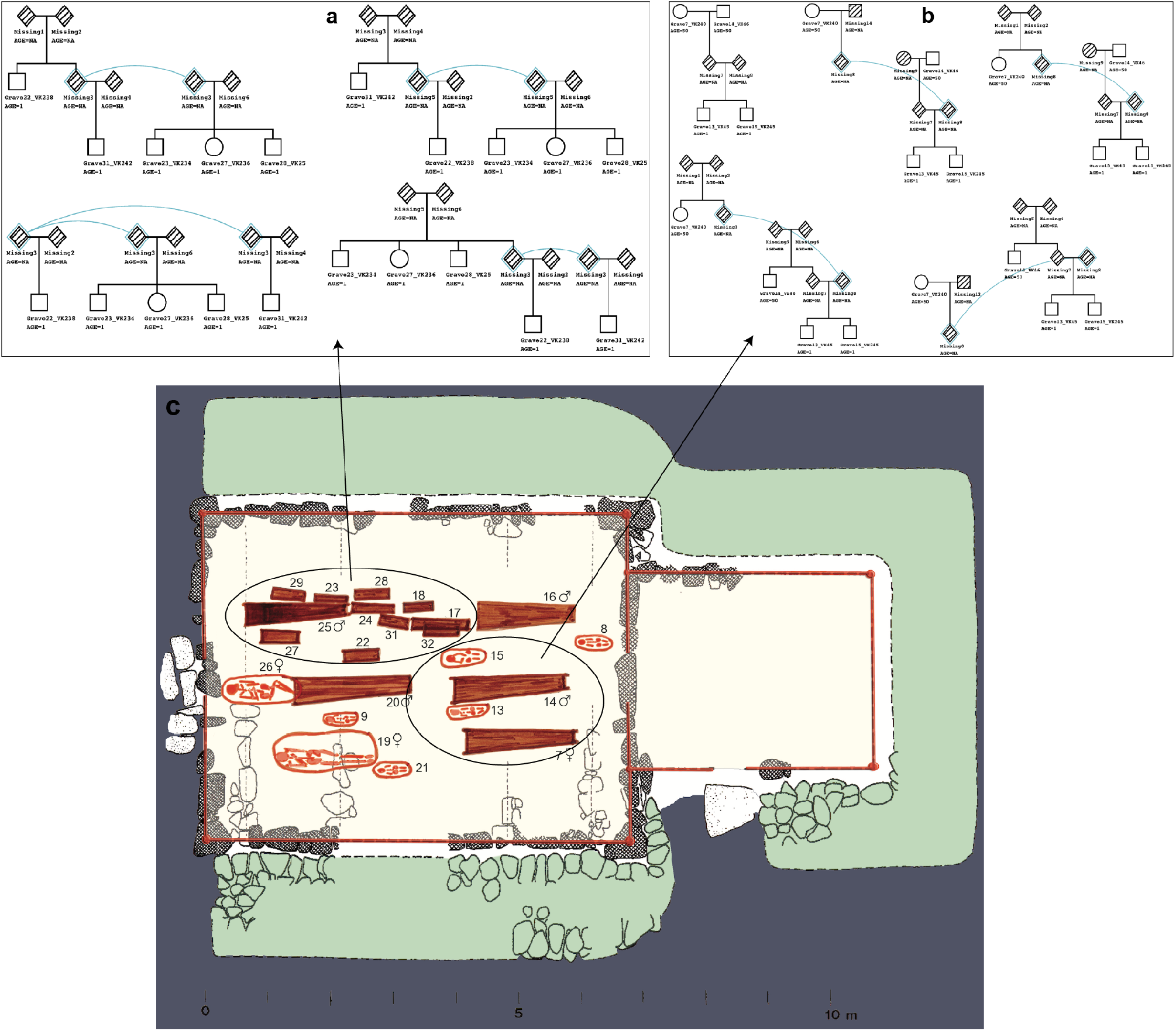
Kinship analysis of ancient samples from Sandoy Church 2 site in Faroe Islands. **a,** Reconstruction of four most likely pedigree networks for one (Family-1) of the three families in Sandoy Church 2 site in Faroe Islands. **b,** Five most likely pedigree networks for the Family-2: the most “parsimonious” network (top left) is likely to represent the true family relationship between the individuals (i.e. grandparents and grandsons) based on the burial pattern of the graves as shown at the bottom image (**c**). Ages of the individuals are approximate to help pedigree reconstructions. Blue diamond shapes and lines in each possible pedigree reconstruction represent the same individual.

