## Supplementary for "Population genomics of the Viking world"

### SUPPLEMENTARY INFORMATION

#### Supplementary Note 1. Ancient samples and archaeological background

##### The Viking Age: A definition

The Viking Age is understood as referring to the period c. 800-1050 CE in Scandinavia, the British Isles, the North Atlantic and the Baltic Sea Area. Like most other historical time periods (e.g. the "Middle Ages", "Age of Crusades", "Reformation Period", or "Age of Enlightenment"), the term "Viking Age" does not refer to a naturally bounded time slice. It is commonly defined as a historical durée marked by the widespread and politically significant operation of sea-born warriors and colonists of Scandinavian descent in regions outside Scandinavia[1].

Albeit the definition and naming of the period is debated[2], most researchers agree that it is a phase of rapid and profound economic and cultural changes[3,4]. Many of these are related to the expansion and diversification of deep-sea seafaring, with a number of knock-on effects for societies. These include: closer interactions and confrontations between groups around the Baltic and North Seas and further afield[5,6]; an acceleration of cultural change and exchange, leading, for example, to the adoption of monetary systems of exchange and the spread of Christian religion in Scandinavian kingdoms[7–9]; an expansion of commercial exchange, focused on emerging maritime trading towns[10–12]; and colonization of the North Atlantic islands including the Faroese, Iceland, Greenland, with brief ventures into Newfoundland[13,14]. The demographic conditions and consequences of these events are widely discussed[4,15,16], but many basic facts concerning both the size and development of populations and their interactions are poorly known from existing sources.

##### Populations and connectivity in first millennium CE Scandinavia

The Viking Age does not by any measure mark a break of previous isolation, and any genetic developments in the period must be considered on the basis of a long and changing history of interactions in Northern Europe. Scandinavian societies were exposed to an increased cultural and probably also genetic exchange already in the late Roman period (c. 200-375 CE) with evidence of intense interactions between Scandinavian societies and the Roman Empire, especially its active Rhine boundary[17]. In the Migration period (c. 375-550 CE) written sources and archaeological evidence point to large scale migrations from Northern Europe including parts of Scandinavia into

Western Europe and the Mediterranean area[18]. This includes the settlement of Anglo-Saxons from present-day Northern Germany and Denmark in England, who left a very considerable cultural and genetic impact[19,20].

The period around c. 530-550 CE has recently attracted particular attention, as evidence accumulated for a severe climatic event, the "Late Antique Little Ice Age", when several major volcanic eruptions caused extreme global weather events in the Northern hemisphere[21]. This lead to widespread famine and pandemic, the "Justinian Plague", which is recorded across wide areas of Europe, probably including Scandinavia[22]. The effects in Scandinavia are suggested to include a demographic drop through widespread death and migrations[23]. While little evidence currently exists to qualify the detailed pattern of developments, it is possible that regions were affected differently, whence changes in population genetics could have occurred as a consequence.

The period c. 550-700 CE shows reduced archaeological evidence of migrations from and supra-regional interaction between Scandinavia and other regions[24], albeit punctuated by finds such as the c. 630 CE Sutton Hoo boat burial in East Anglia, which shows marked Scandinavian affinities in material culture and burial ritual[7,25].

As a historical dynamic, the "Viking phenomenon" is considered to emerge gradually in the course of the 700s CE. This is suggested by various lines of evidence including: the chronology of objects, which were likely brought into Scandinavia through the earliest raid and maritime trade[26–28]; evidence for the evolution of boat-building technology[29]; or direct evidence of maritime warfare, such as the boat burial of warriors from a mid-eighth century raiding party found in Salme, Estonia[30]. In particular, the chronology is charted by the spread of maritime trading towns or *emporia* in the Baltic Sea area (e.g. Ribe in Western Denmark c. 700 CE; Åhus in South East Sweden and Truso in Eastern Poland c. 725 CE; Birka in Middle Sweden and Staraja Ladoga in North West Russia after c. 750 CE [10,12]).

A first wave of raids, trade and colonization culminated in the period c. 840-880 CE, when large Viking armies operated in France and the British Isles, while the earliest colonists settled in Iceland, and trading expeditions reached the Black Sea, the Atlantic coast of Spain and Portugal, and with occasional ventures into the Mediterranean[31]. This is followed by a period with less evidence of Viking activity in Western Europe. Eventually this gave way to what is sometimes referred to as the "second Viking Age" in c. 970-1020 CE, when raids resumed in the west, leading to the amalgamation of a North Sea empire under king Cnut, who came to rule England, Denmark, Norway and part of Sweden between c. 1016 and his death in 1035[32]. While there is evidence for large-scale movement

and migrations across sea in both these episodes, it is not clear how they will have contributed relatively to the demography and population genetics of the period.
The end of the Viking Age is conventionally set to the mid eleventh century CE, when Scandinavian seaborne armies cease to be a primary political force in Western Europe. However, in many regions this date is acknowledged to be an arbitrary convention, as the pattern of maritime activities and connectivities, which mark out the Viking Age, continue to evolve in Scandinavia, the Baltic Sea area, Atlantic Britain and the North Atlantic well into the following centuries[1].

#### The Viking Age burial record and DNA retrieval

Burial customs show marked regional and chronological variation in and around Scandinavia in the Viking Age. While most regions see a gradual increase in the use of inhumation burials towards the end of the period, the early Viking Age is represented mainly by cremation burials in many parts of Scandinavia. This implies that there is a limited record of relevant bone remains available for sampling in much of Sweden, for example[33,34]. This problem is accentuated for the centuries immediately before the Viking Age (600-800 CE), where cremation burial is even more prevalent, leaving few options to sample for human DNA in most regions of Scandinavia.
The adoption of Christian burial practices in Scandinavia and the Baltic Sea area from the 10th century onwards implied that virtually all members of society from infants to adults were buried in designated burial areas around churches, leading to a very comprehensive archaeological record. For all previous periods, the record of individuals given an archaeologically visible burial is highly selective and skewed towards social elites. This must be borne in mind when assessing evidence for mobility and diversity, as there is much to suggest that elites maintained wider networks and enjoyed higher degrees of mobility compared to the average population[33].

For this study, we have shotgun sequenced 528 ancient samples most of which were excavated in the major areas of Viking expansion during the late 8th until late 11th centuries. After removing poorly preserved and contaminated samples the number of samples was reduced to 442. The approximate locations of the archaeological sites are presented in Figure 1 in the main text. The details of sampled individuals with their unique IDs used throughout this study is presented in Supplementary Table 1. The vast majority of skeletal remains used for ancient DNA extraction were teeth and petrous bones. The samples originate from over 80 archaeological sites, covering large parts of Northern Europe and Greenland which is now represented by multiple countries including: Greenland (Eastern and Western settlements, n=23 individuals), Iceland (n=17), Faroe Islands (n=17), Ireland (n=4), UK (n=42), Norway (n=41), Denmark (n=89), Sweden (n=123), Poland (n=10), Estonia (n=34), Russia (n=33), Ukraine (n=4) and Italy (n=5). The approximate locations of the main archaeological sites are presented in Figure 1 in the main text.

Most samples have been archaeologically dated to the Viking age (8-12$^{th}$ centuries CE) or had direct relevance to the Viking period such as the ones from later Norse settlements in Greenland, medieval samples from Faroe Island or noble family members of Ukrainian rulers of 12-13$^{th}$ centuries CE as well as earlier pre-Viking (from Estonia) and Iron Age samples. Two samples that were initially sampled as Viking Age individuals were identified as Late Neolithic/Bronze Age after radiocarbon dating.

This appendix contains a brief description of most archaeological sites from which we have ancient samples used for this study.

### DENMARK

**Catalogue of Danish skeletons: Introduction**

The catalogue present information on the skeletons from Danish collections included in the study. This comprises find circumstances, burial type, a brief report of objects found with the skeleton and their dating, plus reports on osteological, pathological and biochemical analyses that have been carried out prior to the current project. The catalogue is based on information in the national database "Fund og Fortidsminder", together with digests from published sources. The catalogue is organized geographically after the National Sites and Monuments Numbers ("Stednummer"). First, the museum and the museum number of the find is given. After this, the site is indicated by place name, parish ("sogn"), shire ("herred") and county ("amt"). On the next line, the National Sites and Monuments number is given together with the year of finding. The project-specific "VK"-number(s) for Viking Genomics is/are listed after the description of the sites/graves.

**Bakkendrup**

**National Museum of Denmark RAS 45/83, Bakkendrup, Bakkendrup Sogn, Løve Herred, Holbæk Amt.**

Nat. Sites and Monuments no.: 03.02.01-52 (sb. 52.) Year of finding: 1979

A cemetery with flat inhumation graves near Bakkendrup Bro. It was excavated in 1979 by the National Museum. Dated to the second half of the 9th century AD.

Samples used for DNA analysis:

VK294 Denmark_Bakkendrup losfund-2, conc.5

VK315 Denmark_Bakkendrup Grav 16

VK369 Denmark_Bakkendrup losfund-2, conc.1

**Bårse**

**Museum Southeast Denmark SMV 28/86, Bårse, Tyvestensagern, Bårse Sogn, Bårse Herred, Præstø Amt.**

Nat. Sites and Monuments no.: 05.02.03-67. Year of finding: 1986

Excavated by Museum Southeast Denmark in 1986. A small Viking Age burial ground with 11 inhumations located on a small hill. Grave A was the deepest and was covered with three large stones. Grave goods: a strap-end mount, a buckle from a belt (same as the one found in Rantzausminde) dated to the 10th century AD, whetstones and a pumice with abrasion marks. Cremation graves were found underneath the inhumations (not known if these were of animals).

Samples used for DNA analysis
VK281 Denmark_Barse Grav A

**Besser**
**National Museum of Denmark N01 353/41, C24368-83, Besser S, Samsø Herred, Holbæk Amt.**
Nat. Sites and Monuments no.: 03.05.01-45 Year of finding: 1941
Excavated by the National Museum in 1941. Six graves were found under a compact layer of stone on a small hill. The graves were originally constructed to fit wooden coffins, but fragments from a wood coffin were only recovered in one grave (Grave 1). Grave 1 was also the richest of the six burials. It contained a trefoil silver brooch, two oval brooches, a spindle whorl, four glass beats, a key, an iron knife and a hemispherical vessel. Grave 4 contained a shield boss, a whet stone, a knife, a ring needle of bronze and some undefined iron. The other graves only contained an iron knife. Grave 3 was the only inhumation grave. Grave 1, 5 and 6 were cremation graves but in man-length coffins. There were no human remains recovered in Grave 2 and 4. Has been dated to be from 750-1066 AD.

Samples used for DNA analysis
VK298 Denmark_Besser Grav III

**Bødkergarden**
**Langelands Museum LMR 13372, Bødkgårds mark/Bødkergård, Tryggelev Sogn, Langelands Søndre Herred, Svendborg Amt.**
Nat. Sites and Monuments no.: 09.03.07-40. Year of finding: 1997
Excavated by Langelands Museum in 1997. A small Viking-Age burial ground on a small hill containing five preserved inhumations. Soil is gravel and clay. Three graves are oriented E-W and each had an iron knife as grave goods. Two adjacent graves in a N-S direction. One contained a small clay-pot. Remains of settlement from early Iron Age with postholes and pits. Burial D (Male, adult), SW-NE oriented. Skull separated from body and found adjacent to lower part of torso. Mandible

found upside down in "original head end". Buried in supine position. Robust. Not young. Knife was found underneath the skull. Grave H (Male, adult) Right side hocker position in N-S direction with head N. Arms and legs bended. A small clay pot (14 cm diameter) next to torso. Charcoal and burned bone (animal) found in fill. Entire cemetery dates broadly to 9th century AD based on the finding of buckles/pins (type P 42 after Ingmar Jansson 1985)[35]. Grave D: Younger than grave C. Difficult to date more precisely. Grave H did not have any stratigraphy that could be related to the other graves. It was buried with a clay jar. It has also been dated to 9th century AD.

Samples used for DNA analysis
VK289 Denmark_Bodkergarden Grav H, sk 1
VK291 Denmark_Bodkergarden Grav D, sk 1

**Bogøvej**

**Langelands Museum LMR 12077, Bogøvej 21, Lindelse Sogn, Langelands Sønder Herred, Svenborg Amt.**

Nat. sites and monuments no.: 09.03.04-28. Year of finding: 1987, 1988, 1989

Excavated in 1987, 1988 and 1989 by Langelands Museum. Burial ground from Viking period. The cemetery is located on small drumlins of very mixed moraine material with a high chalk content; there were thus varying, but often good, conditions for the preservation of skeletons[36]. The burial ground is located on a small hill 500 m south of Lidelse Nor in an elongated area (70m x 30 m in a NE-SW orientation). The burial ground was discovered in 1920, when four inhumation graves were uncovered. A total of 49 graves have been excavated of which three were double graves. Excavations in 1987 and 1988 also revealed activity dated to the late Bronze Age including 17 pits of which some contained flint tools, large amount of pottery, small fragments of bronze and some tools made of antler. The majority of the individuals had been buried in a supine position and a few in hocker. Some of the inhumation graves contained grave goods: knife- , whetstones, buckles, glass beads. Two stone lined tomb graves (Grave BA) had carriages (minus wheels) as coffins ("Vognfadning"), and a female burial with the remains of a wooden shrine containing an intact comb. A male burial contained an axe dated to the late Viking Age period, a strike-a-light and an Arabic silver coin. One skeleton was decapitated (Grave T). Most graves were oriented E-W. Of the sexed individuals: 15 were males and 22 females. Only few subadults. Dated to 10th century, possibly with few graves dated to the early 11th century AD (unknown which ones).

Samples used for DNA analysis

VK286 Denmark_Bogovej Grav BJ
VK288 Denmark_Bogovej Grav BA
VK292 Denmark_Bogovej Grav A.D.
VK320 Denmark_Bogovej Grav S
VK338 Denmark_Bogovej Grav BV
VK361 Denmark_Bogovej BX
VK362 Denmark_Bogovej LMR 12077
VK363 Denmark_Bogovej BT
VK364 Denmark_Bogovej BN
VK365 Denmark_Bogovej BS
VK366 Denmark_Bogovej BY
VK367 Denmark_Bogovej D
VK368 Denmark_Bogovej T

**Galgedil**

**Odense City Museums OBM 4520, Galgedil, Otterup Sogn, Lunde Herred, Odense Amt.**

Nat. Sites and Monuments no.: 080306 -08. Year of finding: 1999–2005

Excavated by Odense City Museums in 1999–2005. Galgedil is a Viking-Age cemetery (c. 800-1050AD) located in the northern part of the Danish island of Funen. The site revealed 54 graves containing 59 inhumations and 2 cremation burials. Previous study of the remains to date has included light isotopes of carbon and nitrogen in collagen (10 samples) and the radiocarbon determination of the age of eight samples. In addition, aDNA was investigated in 10 samples from the cemetery by Melchior et al.[37] and Strontium analysis was carried out by Price et al.[38] The cemetery is situated on the top and down the southern and western slope of a small hill in a rolling, moraine landscape 5 km from the former seacoast. The investigated skeletal material from Galgedil consisted of the human remains from 57 inhumations with varying degrees of preservation. The number of male and female skeletons was almost the same. There were 24 males (48%) and 19 females (38%). Sex could not be determined in six of the adult skeletons (14%) or for the eight subadult individuals of various ages. Several individuals seem to be non-locals based on strontium isotopes: OU, UD.

Samples used for DNA analysis

VK133 Denmark_Galgedil KO
VK134 Denmark_Galgedil ALZ
VK135 Denmark_Galgedil ALY
VK138 Denmark_Galgedil AQQ
VK139 Denmark_Galgedil ANG
VK140 Denmark_Galgedil PT
VK141 Denmark_Galgedil OMB/BFQ
VK278 Denmark_Galgedil TQ
VK279 Denmark_Galgedil AXE
VK280 Denmark_Galgedil UO
VK370 Denmark_Galgedil ANO
VK371 Denmark_Galgedil UD-Vest
VK372 Denmark_Galgedil KM
VK373 Denmark_Galgedil BER
VK411 Denmark_Galgedil TT
VK446 Denmark_Galgedil LS

**Gl. Lejre**

**Roskilde Museum ROM 641/85, Mysselhøjgård, Allerslev Sogn, Volborg Herred, København Amt**

Nat. Sites and Monuments no.: 02.06.01-115. Year of finding: 2009

Excavated by Roskilde Museum in 2009. The archaeological site of Gl. Lejre. is located on Sealand ca. 35 km west of Copenhagen. Seven inhumation graves or deposit finds were excavated at the hall area of the aristocratic residence of Gl. Lejre[39]. One of the graves, A1636 was placed on the top of a hill in the centre of the residence while a group of four burials, A1859, A1860, A1861 and A1896 were found 14 meters south of the it on a small slope. Grave A1880 was found in a disturbed area north of grave A1636 and 13 meters to the west in the North-Western corner of the residence another disturbed grave, A1697, was found.

All but one skeleton was lying in supine position with head in the west and in a long oval shaped graves and with no evidence of coffin, except for grave A1636. Grave A1896 was lying in hocker position. A total of five individuals could be sexed: four males and one female. Everybody were middle aged adults or older. The only grave good that was found consisted of a brass finger ring placed on the left fourth finger of the female A1861).

Samples used for DNA analysis

VK94 Denmark_GI Lejre-A1861

VK445 Denmark_Gl Lejre-A1896

**Hesselbjerg**

**Moesgaard Museum FHM 1379, Hesselbjerg, Randlev Sogn, Hads Herred, Århus Amt**

Nat. Sites and Monuments no.: 15.02.11-12. Year of finding: 1963–70, 1997–2001

The cemetery has a distinct location on a narrow sand and gravel hill stretching 3-400 m in a north-south direction. The surrounding landscape is flat with fertile agricultural land (UTM 574947 / 6200145). The cemetery is located approximately 1 km south-east of the parish village Over Randlev and 3.8 km from the coast of Kattegat. The graves were positioned on the hilltop and on the east side of the hill. The graves were excavated by Moesgaard museum in 1963–70 and 1997-2001[40–42]. A total of 104 graves were recovered of which 84 were inhumations and 20 were cremations. The preservation of the skeletons varied. More or less intact skeletons were recovered from 69 graves. Over 80% of those interred in the cemetery were women. There were only limited graves and these were mostly personal equipment such as knives, iron belt buckles, whetstones, pottery, and a few pieces of jewelry. Only few graves contained elaborate artefacts. A single pit contained numerous glass and amber beads and an elaborately decorated bronze gilt belt buckle. The dating based on grave goods is mid- 9$^{th}$ to 10$^{th}$ century AD. An associated building was excavated c. 200 m north-west of the cemetery. A well has been dated through dendrochronology to about 900 AD. A pollen analysis of the bottom shows that the nearby area consisted of fields for grassing animals.

Samples used for DNA analysis

VK84 Denmark_Hesselbjerg Grav 3

VK86 Denmark_Hesselbjerg Grav 13

VK87 Denmark_Hesselbjerg Grav 41b, sk PC

VK300 Denmark_Hesselbjerg Grav 22, sk IR

VK339 Denmark_Hesselbjerg Grav 16, sk GO

VK340 Denmark_Hesselbjerg Grav 5, sk V

VK383 Denmark_Hesselbjerg Grav 11, sk DT
VK384 Denmark_Hesselbjerg Grav 14, sk EU

**Hesselbjergmarken**

**Langelands Museum LMR 11163, Hesselbjergmarken, Magleby Sogn, Langelands Søndre Herred, Svendborg Amt.**

Nat. Sites and Monuments no.: 09.03.06-35. Year of finding: 1860, 1981

Exacavated by Langelands Museum in 1982[36]. First burials discovered in 1860 and in 1981. Viking-Age burial ground. Grave B was buried with iron knife, whetstone and a silver coin (Museum no. 11163:6a) Samanide coin dated to 903-913 AD. Buried in supine position with head in east end with face turned north and feet in the west.

Samples used for DNA analysis
VK318 Denmark_Hesselbjergmarken Grav B

**Hessum**

**Odense City Museums/Nationalmuseet? [ML1] NM 818/50, Hessum, Skeby Sogn, Lunde Herred, Odense Amt.**

Year of finding: 1950

The cemetery was recovered during gravel digging and four of the graves were excavated and analysed by Odense City Museums and the National Museum of Denmark in 1950. Two of the graves were relatively well preserved. Only limited grave goods in the form of an iron knife with wooden handle. It is unknown which grave it belonged to.

Samples used for DNA analysis
VK295 Denmark_Hessum sk 1
VK316 Denmark_Hessum sk II

**Hundstrup Mose**

**National Museum of Denmark N01 673/47, C26290-91, C28241-51, Hundstrup Mose, Hammer Sogn, Hammer Herred, Præstø Amt.**

Nat. Sites and Monuments no.: 05.04.01-12 (sb. 12) Year of finding: 1947

Sacrificed victims buried in the bog of Hundstrup. In total two adults, two children and two infants were recovered. The two children (7-11 and 11-13 years) were found lying together. They had been covered with branches and had been buried with an iron knife. The preservation was very good, but none of the skeletons was complete. There are no signs of disease or trauma on the remains. Strontium analyses have been done on all of the children and adults and show that Sk. 2 (VK297) was non-local. The graves date to Germanic period rather than Viking Age[43].

Samples used for DNA analysis
VK296 Denmark_Hundrup Mose sk 1
VK297 Denmark_Hundrup Mose sk 2

**Kaagården**

**Langelands Museum LMR 11563, Kaagården, Lindelse Sogn, Langelands Sønder Herred, Svendborg Amt.**

Nat. Sites and Monuments no.: 09.03.04-60. Year of finding: 1984–87

Excavated by Langelands Museum from 1984–87[36]. Viking-Age cemetery (date to entire Viking Age period until 1000 AD. Grave BF possibly to 10$^{th}$ century AD) on a small hill with 70 graves (from late Germanic period to late Viking Age) but most of graves date to late Viking Age. The richest graves placed on the top of the hill. Contained graves of adults and subadults. Subadults were buried with iron knives. The presence of sand and the shells of marine molluscs in some of these graves at Kaagården, strongly indicates that the human skeletons were cremated on the nearby beach.

Samples used for DNA analysis
VK274 Denmark_Kaargarden 391
VK275 Denmark_Kaargarden 217
VK276 Denmark_Kaargarden BH
VK285 Denmark_Kaargarden Grav BZ
VK287 Denmark_Kaargarden Grav BS
VK317 Denmark_Kaargarden Grav BF99

**Kumle høje**

**Langelands Museum LMR 12845, Kumle høje, Lindelse Sogn, Langelands Sønder Herred, Svendborg Amt.**

Nat. Sites and Monuments no.: 09.03.04-41. Year of finding: 1998

Excavated by Langelands Museum in 1998. Viking-Age cemetery (date to 10$^{th}$ century AD ) with min. 11 adult individuals buried (males and females). A rare double grave with two males on top of each other: one on its back and the other facing down. Some cultivation post-mortem damage. Another male grave with healed cranial trauma, a female grave with a 17cm long iron knife, one male grave with broken legs (perimortem), a female grave covered with large stone slabs. The richest grave was of a female in supine position with iron artefacts and amber, part of a bronze belt buckle. Several other male graves showed trauma.

Samples used for DNA analysis

VK290 Denmark_Kumle Høje Grav O

**Ladby**

**National Museum of Denmark NM 182/34, Ladby, Kølstrup Sogn, Bjerge Herred, Odense Amt.**

Nat. Sites and Monuments no.: 080106-06 Year of finding: 134–35, 1938

The burial ground of Ladby was uncovered west of Nymarksgård near the ship-grave of Ladby, south of Kerteminde Fjord. It was found during the course of gravel digging. It was excavated by the National Museum in 1934-35 and 1938. Ten flat inhumation graves were recovered. Traces of wooden coffin was found in Grave 2 (VK319). The grave was oriented NW-SE. Grave 4 (VK301) was buried in a supine position with head resting on a stone. The grave is oriented WN-ES. An iron knife was found in the grave. It has been difficult to determine the earliest and latest date for the use of the burial ground, since most of the graves were badly damaged and did not contain any objects that could yield specific dating. Most of the finds were iron knives. A C-14 dating of grave 4 from 1997 yields a date of 640-890 AD with +/- 1 standard deviation. Conclusively, the burial ground is thought to have been in use from about 700 to sometime into 10$^{th}$ century CE[44–46].

Samples used for DNA analysis

VK301 Denmark_Ladby Grav 4

VK319 Denmark_Ladby Grav 2

**Lejre**

**Roskilde Museum/National Museum of Denmark NM 194/45; NM 152/46, NM C29995-30150, Lejre, Kornerup Sogn, Sømme Herred, Københavns Amt.**

Nat. Sites and Monuments no.: 02.06.01- 7. Year of finding: 1953–68

Viking-Age cemetery comprising 49 graves at the Gl. Lejre ship setting. Excavated by National museum in 1953-68. Four graves were cremation graves. The fill of several graves contained potsherds and cremated bones. No archaeological information is available.

Samples used for DNA analysis

VK90 Denmark_Lejre Grav 902

VK92 Denmark_Lejre Grav 935

VK247 Denmark_Lejre Grav 804

VK385 Denmark_Lejre Grav 321

**Rantzausminde**

**Langelands Museum NM 288/27, Rantzausminde, Egense Sogn, Sunds Herred, Svendborg Amt.**

Nat. Sites and Monuments no.: 09.05.04-66. Year of finding: 1926, 1927

Viking-Age cemetery. Seven inhumations. No further metadata at Antrolab. There is no C-14 dating, but few archaeological artefacts date the graves to late $8^{th}$ to early $9^{th}$ century AD.

Samples used for DNA analysis

VK312 Denmark_Rantzausminde Grav 1

VK313 Denmark_Rantzausminde Grav 2

VK314 Denmark_Rantzausminde Grav 5

**Stengade**

**Langelands Museum LMR C195, Stengade I, Tullebølle Sogn, Langelands Nørre Herred, Svendborg Amt.**

Nat. Sites and Monuments no.: 09.02.09-58. Year of finding: 1905–23, 1963, 1972-74

Excavated by Langelands Museum in 1972-74. A Viking Age cemetery constructed on a small bank and discovered during marl digging. The site comprises four inhumation graves, of which Grave 3 had a rich equipment of weapons and riding gear as well as the skeleton of a horse. Human remains were only preserved in Grave 4, a mature adult male buried with an iron knife. The site also comprised two inhumation graves from the Early Roman Period, four cremation graves from Roman period and three undateable inhumation graves. Skeletal material has not been preserved from these graves. Grave LMR c195 (AS 36/78) may stem from this site[44,47,48].

Samples used for DNA analysis

VK282 Denmark_Stengade I, LMR c195

**Tollemosegård**

**Tollemosen, Tollemosegård. Græse Sogn, Lynge-Frederiksborg Herred, Frederiksborg Amt. MFG 113/97.**

Nat. Sites and Monuments no.: 01.03.03-61. Year of finding: 1997

A cemetery dated to late Germanic and early Viking Age period (700-1000 AD) based on grave goods. A total of 54 graves. Several containing double graves and a few cremations. Double graves in layers. Varying amounts of grave goods: from none, to rich goods including sacrificed animals, knives, glass beads, amber beads, nails, wooden buckets. Strontium analysis has been conducted on BQ (VK65), BT, and EW (VK 70) samples.

Samples used for DNA analysis

VK65 Denmark_Tollemosegard-BQ

VK69 Denmark_Tollemosegard-DS

VK70 Denmark_Tollemosegard-EW

VK71 Denmark_Tollemosegard-BU

**Trekroner**

**Roskilde Museum ROM 2285, Trekroner Øst-Grydehøj, Fløng Sogn, Sømme Herred, Roskilde Amt.**

Nat. Sites and Monuments no.: 02.04.01-17. Year of finding: 2000

Grydehøj is a small hill east of Roskilde. The hill has been used as a burial site for approximately 3000 years. The west side of the hill was used during the early Viking Age period in which 27 inhumations have been recovered. At the time of excavation, the highest point of Trekroner-Grydehøj was approximately 39 m above sea level and thereby the highest point within 1km radius. At the foot of the east side there is a small lake today, but may have been a peat bog at that time.

The burials dated to the Viking Age period represents burials from the very early Viking Age ($8^{th}$ to $9^{th}$ century). Only few graves contained grave goods that could aid in the dating of the graves and all pointed towards this early period. Anthropological examination has shown that there were 14 males, 8 females and 10 unidentified individuals. Grave goods were found in 21 graves. Larger stones were found in the grave A2058 and a stone was found on the left shoulder of the body[49].

Samples used for DNA analysis

VK284 Denmark_Grydehoj A2058

**Ribe**

**Sydvestjyske Museer ASR 13, Ribe, Ribe Domkirke sogn, Ribe Herred, Ribe Amt.**

Nat. Sites and Monuments no.: 19.04.08-81. Year of finding: 2008–11.

Morten Søvsø, Museum of Southwest Jutland, Denmark

Excavations of this site were carried out in 2008-11 at the cathedral in Ribe (Fig. S1.1), Jutland located a Christian cemetery sealed by settlement layers c. 1050 AD[50]. $^{14}$C-dates combined with a large archaeological dataset including stratigraphy, preservation, grave customs and coffin types dates the burials between c. 850 and 1050 AD leaving little doubt, that the find documents the existence of Ansgar's church, founded c. 855 AD. Ribe is Scandinavia's first town from around AD 700. Became a major trading place/emporium with controlled coin economy in the $8^{th}$ C. Decline from late $9^{th}$ C. A Church, the present-day Cathedral, was founded c. AD 860. Samples come from Christian burials at this site predating AD 1050.

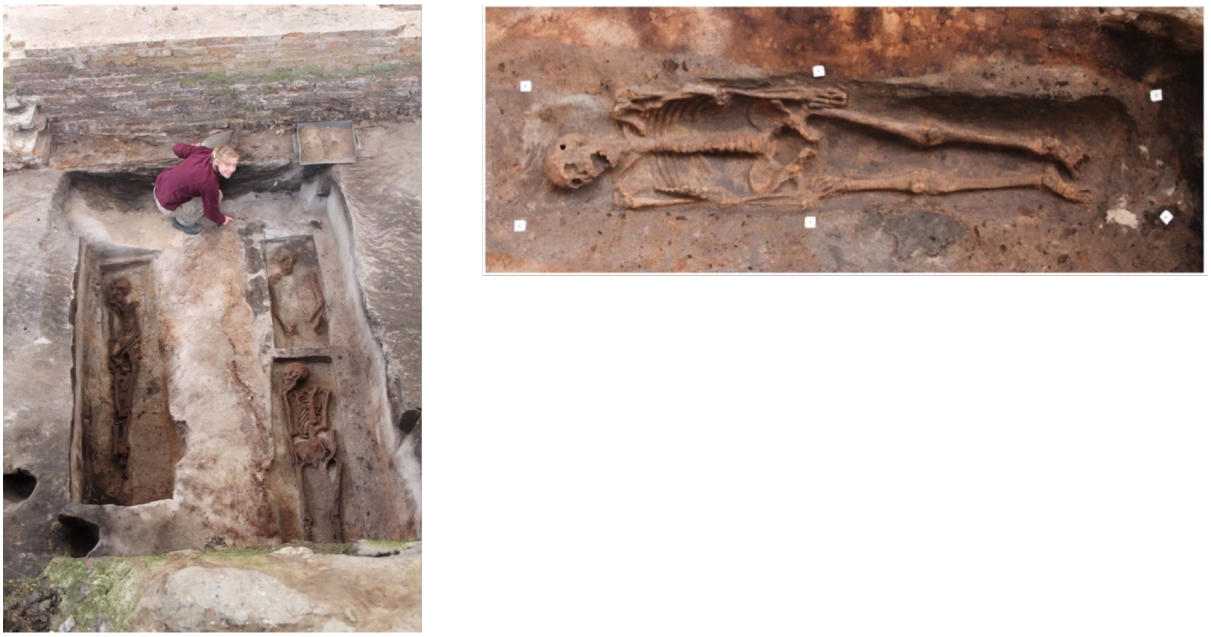

**Fig. S1.1**: The Ribe site during excavation (left) and the skeleton K1586 sampled for aDNA analysis in this study. Photo: Museum of Southwest Jutland.

Samples used for DNA analysis

VK322 K1568
VK323 K1563
VK324 K1552
VK325 K1572
VK326 K1578
VK327 K1586
VK328 K1594
VK329 K1600
VK330 K1582

**Gerdrup**

**Roskilde Museum ROM 191/8, Gerdrup, Kirkerup Sogn, Sømme Herred, Københavns Amt.**

Nat. Sites and Monuments no.: 02.04.08-67. Year of finding: 1981, 1983

Ole Kastholm, Roskilde Museum, Denmark

In the autumn 1981, the find of a Bronze Age sword was reported to Roskilde Museum. The sword came from a field a few km north of Roskilde, an area with many burial mounds, some still standing as visible monuments in the landscape, others more or less destroyed by ages of ploughing and stone harvesting. Archaeologists of the museum visited the find place, and it was evident that the sword originated from a now destroyed burial mound. Furthermore, a number of dark spots were observed in the newly ploughed field, which were thought to be cremation graves. Based on this, a small trial excavation was carried out, which confirmed the existence of such graves as well as an inhumation grave. Two graves were examined in 1981 and the museum returned for a larger campaign in 1983. The excavations unearthed a number of burials dating from Late Neolithic to the Viking Age. In total c. 1800 $m^2$ was excavated.

The 1981-campaign:

Grave A: cremation grave, no artefacts, AMS-dated to c. 400 AD (sample A10) and c. 1800 BC (sample A12).

Grave B: double inhumation grave, male + female, contextually dated to the 9th century AD. The male person (age 35-40 years) was lying with the head and neck in an awkward position, and had probably his neck broken. Furthermore, his ankles were crossed, as if they had been tied together. He was buried with an iron knife. The female person (age 40-50 years) had given birth at least once and was buried with an iron knife, a bone case with iron needles and a spear. The spear head (Jan Petersen's Type E)[51] dates the grave.

Samples used for DNA analysis

VK213 Denmark_Gerdrup-A10

VK214 Denmark_Gerdrup-A12

VK215 Denmark_Gerdrup-B; sk 1

VK216 Denmark_Gerdrup-B; sk 2

**Kragehave Ødetofter**

**Kroppedal Museum TAK 1049, Høje Tåstrup Sogn, Smørum Herred, Københavns Amt.**

Nat. Sites and Monuments no: 020207-115 Year of finding: 2004

Excavated by Kroppedal museum in 2004. Settlements including 42 houses, 11 fence-structures and several wells were found dated to the later part of the Iron Age 3-8th century AD.

Three inhumation graves were recovered. One, a female dated to Early Roman Iron Age (0-200AD), and the other two, a female and a male (x1718) dated to the Late Roman Iron Age (C2/C3) (200-375AD).

The skeleton x1718 was of an old adult male (50+ years) with osteoarthritis in elbow joints, hips and vertebrae. Significant dental wear, AM loss and crowded teeth.

Samples used for DNA analysis

VK532 Kragehave Odetofter XL718

**Brøndsager**

**Kroppedal Museum SØL 941, Torslunde Sogn, Smørum Herred, Københavns Amt**

Nat. Sites and Monuments no: 020213-41. Year of finding: 1997-1998

Excavated by Søllerød Museum in 1997-98. A cemetery containing minimum 20 graves. The graves were found on a small plateau. Settlements were found on the south side of the plateau. Only three skeletons were excavated. They dated to the Late Roman Iron Age (C2) (200-380AD). One grave (grave 900) contained a male (25-30 years) buried in supine position in a wooden coffin with the head to the south. The grave contained rich grave goods. A silver fibula was found next to his right shoulder, a comb of bone and bronze rivets was found next to his hip, and he was wearing a small gold ring on his toe. A wooden bucket with bronze inlays and handle was found by his feet. Two smaller bowls were found inside the bucket as well as a glass. The remains of a slaughtered sheep were found in an upper layer of the grave. He had osteoarthritis in his lower spine.

Another grave contained a sub adult, ca. 14-15 years, possibly male (based on grave goods) buried in a wooden coffin with a wooden bucket with a bronze belt, four clay bowls, two Roman drinking glasses, a play board with red paint, 30 while and 29 black playing pieces, a comb of bone with bronze rivets, a pearl necklace and a gold Aureus with an eyelet. He had a gold coin in his mouth and a gold ornamented ring dated to the Late Roman Iron Age. He was buried with a pig and butchered sheep.

The third grave contained a 4-5-year-old child also buried in a wooden coffin. It was buried with a pearl necklace made of bronze and amber, a comb of bone with bronze rivets, two clays bowls and a pig.

Samples used for DNA analysis

VK521 Sol941 Grav900 Brondsager Torsiinre

**Alken Enge**

**Skanderborg Museum SBM 1028, Torslunde Sogn, Smørum Herred, Københavns Amt**

Nat. Sites and Monuments no: 020213-41. Year of finding: 1997-1998

Excavated by Skanderborg Museum in 2013-2014. The wet meadows of Alken Enge (Alken Meadows) at Lake Mossø are the site of a mass grave with skeletal remains of a minimum estimate of 380 defeated warriors deposited in an original sea basin. The specific bone sample was deposited is in a natural channel between two sea basins. The tooth is from the lower jaw / mandibula on a disarticulated male individual found centrally in the northern main field of the Alken Enge excavation. No abrasion, pathologies or special bone damage have been observed. There were six remaining teeth - the rest has fallen out. In the absence of dating, it must be found attributed to the total dating frame 2 BC-54 AD. Context and fund data etc. is presented in the publication and its supplementary data[52].

Samples used for DNA analysis

VK582 SBM1028 ALKEN ENGE 2013, X2244

#### Catalogue of Norwegian skeletons
#### (Without samples from Trondheim)
**Jan Bill, Museum of Cultural History, Oslo, Norway**

##### Introduction to the catalogue

The catalogue presents information on the Viking Age and earlier skeletons from Norwegian collections included in the study. This comprises find circumstances, burial type, a brief report of objects found with the skeleton and their dating, plus reports on osteological, pathological and biochemical analyses that have been carried out prior to the current project. The catalogue is mainly based on information gathered from published sources, as well as the collection databases of the Norwegian university museums and the database for the Schreiner Collection, where the skeletons are kept. The Schreiner Collection distinguishes between skeletons with crania of 'Nordic', 'Sami', 'Eskimoic', 'Other' or 'Unknown' type in order to avoid conflict with regulations for the sampling and use of Sami human material for research. All the skeletons in the present catalogue have been categorized as having skulls of 'Nordic' type.

The catalogue is geographically organized, starting with skeletons found in SE Norway, and ending with those found in N Norway. First is given the museum number, the starting letters of which indicates to which museum collection the finds belong. The museum number serves as reference to information in the museum archives and registers. "C" is the Museum of Cultural History and University of Oslo; "T" is the University Museum at the Norwegian University of Science and Technology in Trondheim; and "Ts" is the University Museum at The Arctic University of Norway in Tromsø. Next, starting with A, is given the skeleton's number in the Schreiner Collection; the Schreiner Collection is today part of the Institute of Basic Medical Sciences, formerly the Anatomical Institute, and holds the majority of archaeological skeletons from the Viking Age and earlier found in Norway. Next follows the project-specific "VK"-number for Viking Genomics. After that are given the name of the region ("fylke"), the municipality ("kommune") and the farm ("gård"), followed by the farm number, the smallholding number and, if applicable, any local place name associated with the site.

In the next line follows the identification number in the National Register for Archaeological Monuments ("Askeladden"); however, not all finds are registered in this database.

Geographical coordinates are given in EU89-UTM zone 33; if imprecise, the accuracy is stated as within the "smallholding", "farm", or "municipality". Following this, the year of finding is stated.

#### Catalogue

##### Hedmark

**C25552, A4005, VK448, Hedmark, Ringsaker, Berg øvre (619/9 «Breidablikk»)**

Askeladden ID: None

Coordinates: EU89-UTM zone 33 N6746886 E275540 (smallholding) Year of finding: 1933

The skeleton was found in 1933 when a burial mound was being removed by the land user. It was found lying on two stone slabs, and had been covered by two or three further slabs; an accompanying sickle type R. 384 dates the burial to the Viking Age. The site was inspected by Alf Søderholm from the Institute of Anatomy at the University of Oslo, and he concluded that the skeleton, which had been removed from the burial before his arrival, was female.

The Schreiner database describes the skeleton as brownish and from an adult woman with a skull of Nordic type. It also describes the position of the skeleton as 'atypical', and cites a letter from archaeologist Sigurd Grieg, University of Oslo, saying that the head was said to be found lying between the legs, and the body with its upper part to the north.

**C26737, A4304, VK422, Hedmark, Åmot, Arnestad lille (25/1)**

Askeladden ID: 41956-1

Coordinates: EU89-UTM zone 33 N6785034 E303152 Year of finding: 1938

The burial, which was placed in a small mound, was found by a farmer in 1938 when he quarried the mound for stones. Apart from the skeleton, he found the following objects: One double-edged swords of Petersen's M-type, a double-edged sword blade, an axe of Petersen's G-type and an arrowhead type R. 539. The grave was subsequently investigated by an archaeologist, Sverre Marstrander, but apart from another arrowhead, somewhat similar to type R. 538, this did not produce further finds. The human bones in the burial are reported to originate from two individuals, a young, strongly built man in his early twenties, and another person being 50-60Y old. It is also stated that there are animal bones, including horse bones, included in the osteological material.

The M-type swords are dated by Androschchuk[53] to around the mid-10th century, and is a common companion to the G-type axes[51].

The Schreiner database describes the human material as being yellowish and originating from a male and a person of unknown sex. The skull is described as Nordic.

Russ[54] has examined the material and separated it into two individuals, A, and B. It was individual A, which was sampled for DNA. The post-cranial bones were not possible to assign to either of the two individuals. The remains of A consist of a very fragmented cranium, to which two fragments of a mandibula seemingly can be associated. The sex of this individual cannot be determined, but the age determination indicates that he/she was 17-30Y, and most likely in his/her early 20's at the time of death. Porosities on frontal, parietal and occipital bones may indicate anaemia.

Naumann[55] has carried out isotopic analyses on both individuals. A sample from a mandible from individual A produced the following isotope results: $\delta^{13}C$: -20.6, %C: 41.4, $\delta^{15}N$: 10.9, %N: 14.0, C:N: 3.4[55]. A sample from a 2nd molar from the mandible produced those: $\delta^{13}C$: -20.7, %C: 36.2, $\delta^{15}N$: 11.5, %N: 12.6, C:N: 3.3, $^{87}Sr/^{86}Sr$: 0.7159.

**C17564, A2813, VK420, Hedmark, Vang, Tommelstad (6/-)**

Askeladden ID: None

Coordinates: EU89-UTM zone 33 N6747756 E287415 (bruk) Year of finding: *c* 1893

The skeleton originates from a burial mound, where it was found together with three iron knives of a type different from those of the Early Iron Age, an arrowhead of the type R. 539, a fire steel of the common type R. 426 and an iron handle for a pot. The finds only allow a dating to the Viking Age in general.

The information in the Schreiner database is sparse. The skeleton is described as brownish and adult, but no sex determination has been made. The cranium with mandible and a part of the right humerus is preserved, and is characterized as Nordic.

**C25720, A4006, VK393, Hedmark, Ringsaker, Mæhlum (752/1)**

Askeladden ID: 29529

Coordinates: EU89-UTM zone 33 N6751636 E282299 Year of finding: 1933

The skeleton was found during the professional excavation of a burial mound, 7.5 m in diameter, situated on a natural hummock *c* 250 m W of the farm. The skeleton was situated in the middle of the mound, placed E-W. Six arrowheads – three type R. 358, two type R. 359 and one of undetermined design – were placed near the head, while the other finds were made on the northern side of the torso and at the hips. These included an iron tool of unknown function, a possible fragment of an iron buckle, a fire steel type R. 426, and a fragment of a three-layered bone or antler comb with an iron rivet.

The R. 358 arrowheads date the burial to the $10^{th}$ century, a dating not in conflict with the presence of type R. 359 arrowheads[51] and a type R. 426 fire-steel[56].
The Schreiner database describes the skeleton as brownish in color and from an adult male. The skull is characterized as Nordic. Russ[54] also suggests that the individual is male, and based on the teeth she suggest that he was 33-45Y at the time of death, while the cranial sutures indicate a somewhat higher age. The teeth are worn, at least two shows caries, and there is ample calculus and hypoplasia. The man has been strongly built and was possibly above average height. The spine shows lipping age- or wear-related lipping. The cranium shows a large lambdoid ossicle and a parietal notch bone.
Krzewińska, et al. (2015) estimated the mtDNA HVR1 sequence of this individual to represent haplogroup HV4a[57] .

**C27338, A4460, VK394, Hedmark, Åmot, Arnestad store (24/1)**
Askeladden ID: 12709-1
Coordinates: EU89-UTM zone 33 N6784587 E303102 Year of finding: *c* 1943
During the demolition of a one meter high, 5 m wide burial mound of mainly stones, a skeleton and several finds were unprofessionally recovered. An archaeological inspection of the site after excavation did not produce further finds. The finds included a poorly preserved axe resembling type R. 555, equivalent with Petersen's type H, four arrowheads type R. 539 and two R. 540 ones, plus some unidentifiable iron objects.
The R. 540 arrowheads indicate a migration period dating, which is in contrast to the axe of Petersen's type H, datable to the first half of the $10^{th}$ century, perhaps the entire century; an arrowhead type R. 539 does not provide any clarity. The axe may be considered the more important dating element.
The Schreiner database declares the skeleton to be brownish and from an adult of Nordic type, but gives no sex determination.

**Oppland**
**A2808, VK417, Oppland, Gran, Sandeødegården on Nedre Hov (262/- or 263/-, Lindbak)**
Askeladden ID: None
Coordinates: EU89-UTM zone 33 N6699700 E255988 farm Year of finding: 1868
Two inhumation burials in two mounds (no. 1 and 2), excavated in 1868 by Nicolay Nicolaysen. No objects are associated with the burials, and there were no indications of a construction of a grave chamber. Mound no. 1 was app. 6,5 m in diameter and about 2 m high – a 1.5 m high stone was found

on top of the mound, where it has apparently earlier been placed in an upright position. Charcoal was found sparsely in the mound filling, and the skeleton was found, in a somewhat disturbed state, in the middle of the mound, at some distance above the basis of the mound.

Mound 2 was placed next to and NW of mound 1, and was about 1.2 m high and 7.8 m in diameter. Centrally in the mound was found a skeleton, lying on its back in an E-W orientation, with the head towards east.

The graves can only be dated, by their burial type, to within the Late Iron Age.

The Schreiner database describes the two skeletons as brownish and the skull(s) as of Nordic type. It also mentions that there was at least one animal bone in the material turned over to the Schreiner Collection in 1885, but that was discarded along with other bones in 1924. It is unknown, from which mound the skeleton originates.

**C35586, A5305, VK386, Oppland, Lesja, Skålgård søndre (131/1)**

Askeladden ID: 11363

Coordinates: EU89-UTM zone 33 N6898881 E189426 Year of finding: 1981

The grave was found next to a large stone during digging for the construction of a farm road in 1981. An archaeological control dig produced no further information. Apart from the skeleton, the grave contained a sword of Petersen's type L, with remains of a sheath, a whetstone of sandstone, a fragment of an iron knife blade and an iron fragment of unknown use. Androschchuk[53] dates Petersen's L-type from the 870's into the $11^{th}$ century.

The Schreiner database describes the skeleton as being of varying colour and originating from a mature male, around 50Y old at the time of death.

**C24243 and 24297, A3777-3778, VK421 and VK387, Oppland, Jevnaker, Velo nordre (164/6)**

Askeladden ID: None

Coordinates: EU89-UTM zone 33 N6694793 E247284 Year of finding: 1928

Due to a mistake, the two skeletons VK421 and VK387 are recorded on two different farm numbers in the collection, but originate from the same grave on farm 164/6. VK421, *alias* C24243, was found June 1928 during construction of a roadbed from the farmhouse to the public road and sent by the local sheriff to the museum. VK387, *alias* C24297, originates from the resulting archaeological investigation of the find site, carried out by the Anatomical Institute, not the museum of University of Oslo. Because of their different ways into the museum and the findings of ceramics at the archaeological investigation, they were considered two different graves.

The find site is situated in the yard of the farm Velo Nordre, c. 20 SW of the farmhouse. The initial finds were made 25-30 cm below the surface, and consisted of a skeleton oriented E-W, with the head to the W, and a small axe. This is of Petersen's L-type, datable to the mid-10$^{th}$ to 11$^{th}$ century[51]. By the archaeological excavation of the remaining part of the grave some weeks later, remains of a further, female skeleton was found, along with unrecovered remains of two infants. The female skeleton was orientated the same way as the one first found, and the infants were lying at its feet. The excavation also produced a ceramic shard identified to be from a vessel of type R. 361, two undefinable pieces of burned clay and a small, unidentifiable piece of iron. The entire grave, with all four skeletons, had been lined with rather thick timbers, but no iron nails were found. The shard indicates an Early Iron Age date, but may represent an earlier settlement on the site. The Velo farm has produced finds going back to the Bronze Age.

The Schreiner database states about the VK387 skeleton (A3778) that it is brownish, adult, and female. The cranium is classified as Nordic. The VK421 skeleton (A3777) is described as likewise brownish and adult, but male. Russ[54] confirms A3777 to be male, based on the cranial features. The teeth shows strong wear and post-mortal loss of enamel. The age is estimated to +45Y. The post-cranial bones could not be found in the collection. Krzewińska, et al.[57] suggest haplogroup U5a/U5b2 for the individual's mtDNA HVR1 sequence.

**C14690-14692, A1517, VK414, Oppland, Skjåk, Nedre Hjeltar (59/1)**

Askeladden ID: 51891

Coordinates: EU89-UTM zone 33 N6878623 E153537 Year of finding: 1889

The skeleton was found in an E-W oriented burial, with the head towards W, and the grave goods placed at either side of the cranium. The finds consisted of an axe, identified as a R. 555, four arrow heads and some unidentifiable iron objects. The R. 555 type is the same as Petersen's H-type, datable to the second half of the 10$^{th}$ and the first part of the 11$^{th}$ century. The arrowheads cannot be identified. The Schreiner database describes the skeleton as brownish and from an adult male; the cranium is described as Nordic. According to the description by Schreiner[58], substantial parts of the skeleton are preserved. Russ[54] estimates that the individual is male, and +45Y of age, due to closure of cranial sutures. The few remaining teeth show heavy wear, and there are calculus and signs of extreme parodontosis and resorption. The post-cranial skeleton shows extensive wear and examples of eburnation in several joints. Both femurs have an atypical shape, being flat and broad. Krzewińska, et al.[57] find the mtDNA HVR1 sequence of this individual to represent haplogroup H6[57] .

**C21852, A1520, VK415, Oppland, Lunner, Hov**

Coordinates: EU89-UTM zone 33 N6691573 E258795 Year of finding: 1910

The location of this find is slightly unclear, but is here located according to the placename "Hov". This skeleton was found approximately in the centre of a round mound, extended on its back and with the head towards W. The right arm was bent over the breast. The only grave goods found was a mosaic bead of black glass with milk-white and red decorations, datable to the Viking Age.

The Schreiner database describes the skeleton as brownish, adult and from a female; the skull is described as Nordic. A list of the preserved skeletal parts are provided by Schreiner[58].

**Telemark**

**C21794, A1645, VK392, Telemark,Vinje, Særen (23/-)**

Askeladden ID: None

Coordinates: EU89-UTM zone 33 N6626654 E100574 (farm) Year of finding: 1915

The information about the circumstances of this find is vague. It was found by an Olav Nykos during hunting in a scree 2 km N of the Særen farms. Two keys of type R. 459 dates the grave to the Viking Age before the 11th century[56]. No other finds were made.

In the Schreiner database the skeleton is described as greyish, adult and of undetermined sex, but with a Nordic cranium type. Russ[54] estimates the skeleton to be female and, based on the fusion of the cranial sutures and the heavily worn teeth to be +45Y. One caput mandibulae and one femur shows anomalies; some of the long bones are bent, but it is unclear if this is post-mortem or pathological. Krzewińska, et al.[57] find the individual's mtDNA HVR1 sequence to represent haplogroup H*.

**C22242a-d, A1648A and A1648B, VK390 and VK391, Telemark, Skien, Søndre Mæla (4/76 Berg)**

Askeladden ID: None

Coordinates: EU89-UTM zone 33 N6576689 E191488 (GÅRD) Year of finding: 1918

The two skeletons in this grave were found in 1918 during fence construction on the parcels Kveldro and Vesterbø on the smallholding Berg under Mæla søndre. The skeletons were lying close together, and apart from the human remains, only two not cruciform brooches were found. They resemble the brooch in Shetelig[59] and can be dated to the Migration Period (AD 400-550/600).

In the Schreiner database, both skeletons are described together as being brownish in colour and with skulls of Nordic type. Russ[54] has separated the material into individual A (VK390) and B (VK391), but cannot exclude the presence of more individuals. Individual A is a male of 30-50Y; the mandible from this individual shows massive teeth with massive hypoplasia on the canine teeth. Individual B, with the most intact crania, is a female, aging 18-30Y. Naumann et al.[60] notes the following isotopic values from the mandible: $\delta^{13}$C: -20.5, %C: 42.1, $\delta^{15}$N: 11.6, %N: 15.3, C:N: 3.2[55] . A sample from a 3rd molar produced those: $\delta^{13}$C: -21.0, %C: 41.6, $\delta^{15}$N: 9.7, %N: 14.2, C:N: 3.4, $^{87}$Sr/$^{86}$Sr: 0.7137. From individual B, she got the mandible values $\delta^{13}$C: -20.4, %C: 41.1, $\delta^{15}$N: 12.3, %N: 14.5, C:N: 3.3. A sample from a 2nd molar produced those: $\delta^{13}$C: -21.1, %C: 39.6, $\delta^{15}$N: 11.0, %N: 14.6, C:N: 2.9, $^{87}$Sr/$^{86}$Sr: 0.7109.

**C23941a-f, A3697, VK389, Telemark, Skien, Bergsland (60/77 Lagmannsgårdshøyden)**

Topografisk: Schreinerske arkiv: Schreiner 1927:

Askeladden ID: None

Coordinates: EU89-UTM zone 33 N6576540 E192556 Year of finding: 1926

The finds were made during building work at app. 50 cm depth, below a layer of stone slabs. There were no traces or reports of a mound on the site. The finds consisted of skeletal remains plus a spearhead, a frostnail type R. 591 for a horse and a few iron fasteners. The spearhead is of Petersen's type I and dates according to Petersen to the first half of the 10th century[51]; Andrushchuk[53] dates some of the sword types that occur with the I-type spearhead into the second half of the century; the grave is therefore dated to within the 10th century.

In the Schreiner database the skeleton is described as brownish of colour, adult and male.

**Sør-Trøndelag**

**T16298, A4481, VK516, Sør-Trøndelag, Ørlandet, Østråt (Austrått) (82/91)**

Askeladden ID 56213-1

Coordinates: EU89-UTM zone 33 N7074267 E240441 Year of excavation: 1944

This grave find was made during road construction, and was sent in to the Schreiner Collection by Th Petersen from the university museum in Trondheim. No remains of a mound were reported, but the burial was overlooking a small bay, where in modern times harbour structures were established. Two finds are reported: An iron spearhead type R.521, with copper alloy knots, and a highly fragmented shield boss type R.562. These date the find to the 10th century.

In the Schreiner Collection Database the skeleton is described as originating from a burial site, and as being brownish; calvaria and remains of the facial bones are preserved together with a defect lower jaw, a left humerus and fragments of other bones. The individual is estimated to be male and of adult age. According to Scheiner catalogue 11, the bones were found together with a spearhead type R.521, and a shield boss type R.562 from the 10$^{th}$ century. The museum archive at NTNU reports Schreiner to have estimated the age to over 50 Y and the height to 167-168 cm. Schreiner found that the skull was not of the "typical Iron Age type", and he finds it to be a mixture between that and "the Trøndelag Bronze Age type".

**T13363, A3699, VK523, Sør-Trøndelag, Bjugn, Melem (Herstaen) (84/-)**

Askeladden ID: None

Coordinates: EU89-UTM zone 33 N7089516 E239588 Year of excavation: 1926

This find was unprofessionally extracted from a large, partly stone-lined chamber grave, oriented ENE-WSW, inside a cairn. After the extraction, the grave was excavated by a student, B. Irgens Larsen. Several shards of a very thin-walled ceramic vessel of possible Bronze Age type were recovered. The grave also contained an unusually thick-walled cranium, which was broken by the finder. According to the museum archive, Professor Schreiner found that the skull type indicated a date prior to the Migration Period.

The Schreiner database states that the skeletal fragments consisted of a defect calva and minimal additional remains. The colour is described as greyish, and the skeleton is estimated to be from an adult female.

**Nord-Trøndelag**

**T2327, A3705, VK548, Nord-Trøndelag, Stjørdal, Kil søndre (220/-)**

Coordinates: EU89-UTM zone 33 N7040131 E311368 Year of finding: 1927

Askeladden id.: None

The remains were found during the construction of a railroad, in bog soil next to a creek. On the breast of the skeleton were found two well-preserved single shell oval brooches of copper alloy with riveted, plated knots (T-2293), a brown, opaque glass bead (T-2294) and a clay bead (T-2295). The glass bead is by Bjørn[61] said to be of amber. The oval brooches are of the type R. 649[62], which Petersen[63] dates to the 9$^{th}$ century, a dating that Bjørn agrees in.

The Schreiner database states that the skeleton is yellowish and well-preserved but not complete. In

Schreiner[58] are listed 50 bones that are preserved, plus the cranium. The skeleton is from a female; no age is determined. The cranium is described as being of Nordic type. According to Bjørn[61], the height of the female was calculated to 163 cm.

**Nordland**

**TS7659, A5195, VK514, Nordland, Steigen, Vikran Nordre (71/2)**

Coordinates: EU89-UTM zone 33 N7533257 E505595 Year of finding: 1965/66

Askeladden ID: 37739-1

The cranium was found during ploughing a field in 1965, and an archaeological excavation in 1966 produced the remaining parts of a skeleton. The individual was found lying on its left side with flexed legs and the head in SSW. Right arm was extended, with the fingers in front of the knees. A knife was lying at the hand, and another knife plus a composite comb behind the hip. There is no indication that a mound had ever been present over the burial. The composite comb dates the burial to the $6^{th}$ to $10^{th}$ century.

The Schreiner database describes the remains as brownish and well preserved. No sex determination is given, but the age is estimated to 11-13 Y and the body height to 160 cm.

Russ[54] states that sex determination is not possible due to the low age of the individual. The teeth indicate it to be 11Y ± 30M, the rest of the skeleton 10-14Y, and she concludes that the age was presumably 11-12Y. One milk tooth, no. 55, is still in place. Calculus is present on several teeth, but no caries or hypoplasia. Carabellis cusp is observed on tooth 16 and 26. Very weak cribra were present in both eye sockets. Non-metric traits are present on atlas, as two extra bones in the lambdoid suture and a vastus notch on the right patella.

Naumann et al.[60] produced the following isotope results from a sample from a femur: $\delta^{13}C$: -16.8, %C: 42.2, $\delta^{15}N$: 16.3, %N: 16.3, C:N: 3.0[55]. A sample from a $1^{st}$ molar produced those: $\delta^{13}C$: -17.2, %C: 44.7, $\delta^{15}N$: 16.0, %N: 16.3, C:N: 3.2, $^{87}Sr/^{86}Sr$: 0.7104. Krzewińska, et al. determines the mtDNA HVR1 sequence of this individual to represent haplogroup K1a11[57].

**T20545 h:1, A5317, VK526, Nordland, Herøy, Sørherøy (Prestegården, 4/1)**

Coordinates: EU89-UTM zone 33 N7319886 E376306 Year of finding: 1983

Askeladden ID: 36189-1

The find site is situated next to the Herøy Sound, only 80 m from the beach, and 300 m north of the large Romanesque Herøy church from the $12^{th}$ century. A skeleton was found with one oval brooch

10 cm below the surface during construction works in an area that has yielded several other inhumation burials. During a control excavation after the finding, remains of a further skeleton was found. This had been placed in an E-W direction, with the head towards W. A 1.2 x 1.6 m large dark coloring of the sandy subsoil indicates the presence of a wooden coffin or grave lining. It seemed that that the deceased had been laying in a hocker position. There were no indications of a mound present. The finds from the excavation included wooden remains, iron nails and rivets, animal bones, another oval brooch of the same type as found with the first individual, R.647, variant B[63], a knife and textile fragments from the brooches. The two brooches did differ in design, but may indicate that both skeletons came from the same burial, and dates them to the 9th century[63].

According to the osteological investigation following the excavation, both skeletons were from young females. The poorer preserved one (A5316) was by Schreiner estimated to originate from a 17-18 Y old woman, while the better preserved one (A5317) was judged as belonging to a 12-13 year old female, an interpretation also found in the Schreiner database, where the bones are described as yellowish.

Russ[54] found that A5317 belonged to a younger individual early in puberty, with an age of 12-17 Y, most likely between 13-15 Y. Due to young age it is not possible to determine the sex with certainty, but traits on the hip bone indicate female sex. Some cribra orbita, mostly in the left eye socket, and traces of healed cribra in the right one were the only pathologies found. Several non-metric traits were recorded: extra bones in the sutures, especially lambdoid ossicles and an eptiteric bone.

The mtDNA HVR1 sequence of this individual represented haplogroup J1d/J2b[57].

**T5105, A0642, VK529, Nordland, Steigen, Leines (6/-)**

Coordinates: EU89-UTM zone 33 N7513575 E491831 Year of finding: Before 1897

Askeladden ID: None

This skeleton was given by Sheriff Olaf Olsen to Trondheim University Museum with the information that it had been found on the farm Leines in Ledingen parish, Nordland. Together with the skeleton had been found two single-edged swords of the types R. 491 and R. 500, a celt type R. 401, two whetstones, and a possible sickle. The sword type R. 491 is equivalent with Petersens type C, which Androshchuk[53] dates to the early Viking Age, AD 750-870. The celt and the R. 500 sword may indicate an earlier date.

The Schreiner database describes the skeleton as well preserved, brownish, and belonging to an adult man. Schreiner in 1927 lists its preserved parts and describes them as from a male[58].

In 2013, Russ[54] describes the few remains as well preserved, and suggest that they belong to a male of 24-30 Y of age. A slight cribra is present in both eye sockets; the teeth show calculus. The spine shows small osteophytes and the muscle fastenings, especially on the thigh bone.
Naumann[55] measured the following isotope values on a sample from a tibia: $\delta^{13}$C: -18.3, %C: 43.0, $\delta^{15}$N: 15.0, %N: 15.0, C:N: 3.3. A sample from a $2^{nd}$ molar produced those: $\delta^{13}$C: -17.7, %C: 43.6, $\delta^{15}$N: 13.8, %N: 15.9, C:N: 3.2, $^{87}$Sr/$^{86}$Sr: 0.7109. She suggests that marine foods played a significant role in the man's childhood, but that his consumption of freshwater fish and/or terrestrial predators increased in the later part of his life.
Krzewińska, et al.[57] found the mtDNA HVR1 sequence of this individual to represent haplogroup H*.

**Ts5287, A4691B, VK519, Nordland, Steigen, Steigen Mellem (78/- Hagbartsholm)**
Askeladden ID: 7944
Coordinates: EU89-UTM zone 33 N7535488 E498506 Year of finding: 1954
This find is from an excavation of a Migration Period-Viking Age burial site, carried out by H.E. Lund, Tromsø museum, in 1954. The individual is from grave X, and the finds included fragments of a three-layered, composite antler or bone comb with linear ornaments and a five iron nails with adhering wood plus some charcoal. The comb dates the burial to the $6^{th}$-$10^{th}$ century.
The Schreiner database describes the few remains of the skeleton as brownish of colour, adult of age, and of undetermined sex. The skull is categorized as Nordic. A small iron ring is mentioned as grave goods, an object which does not occur in the archaeological record of the find.
Russ[54] estimates the skeleton to be female, based on the crania and the femur joints. Age is determined to 25-30Y, based on limited teeth wear. An enamel hypoplasia is observed on dx maxilla canin, and on the left femur an exostos on the neck of the left femur, behind the head.
Naumann[55] has measured the following isotope values[55]: Femur: $\delta^{13}$C: 19.1, %C: 43.8, $\delta^{15}$N: 13.8, %N: 15.4, C:N: 3.3. A $1^{st}$ molar produced: $\delta^{13}$C: -17.2, %C: 47.4, $\delta^{15}$N: 15.8, %N: 17.9, C:N: 3.1, $^{87}$Sr/$^{86}$Sr: 0.7106.

**Ts4306, A4511 and A4512, VK515 and VK530, Nordland, Bodø, Rønvik nedre (32/100, Rønvik)**
Askeladden ID: None
Coordinates: EU89-UTM zone 33 N7462971 E473589 Year of finding: *c* 1946

These two graves were found about 20 m apart, and *c* 70 m from the beach, and it is unknown which skeleton belonged to each grave. One was found in a very shallow grave, only 20 cm below the turf, and contained, apart from the human remains, a three-layered bone or antler comb and a needle house of bone of type VGJ 492, a miniature sword of bone or wood of a shape like Petersen type V, plus some possible chest mounts of iron. Androshchuk[53] dates the V-type to around AD 1000, which might be a fair guess of the date of the amulet as well. The other burial, which was 180 cm deep and covered with stones, also contained human remains together with a sword of Petersen's Y-type, and a spearhead of Petersens G-type plus a shield boss of the type R. 563. Androshchuk[53] dates the Y-sword to AD 900-975, and Petersen[51] dates the spearhead to the second half of the tenth and the early 11$^{th}$ century. This grave thus can be dated to the middle or second half of the tenth century. Since the burials both date to the second half of the 10$^{th}$ century or perhaps slightly later, both skeletons can also be assigned to this date.

VK 515 (A4512): The Schreiner database states that the skeleton is brownish of colour and originates from a young male of adult age, with a Nordic type skull. Russ[54] also determines the skeleton to be young and male on the basis of DSP of the coxae, less so of the cranium. Remains of milk teeth roots are still present, but all molars have erupted; there are several examples of enamel hypoplasia. On the basis of the teeth and the fusion of the epiphyses, the age is estimated to 18-20Y. The strong development of clavicula and the forearm bones indicates strong physical labour.

Naumann et al.[60] have measured isotope values: A femur gave $\delta^{13}C$: -16.00, %C: 41.2, $\delta^{15}N$: 16.8, %N: 15.0, C:N: 3.2[55]. A sample from a 1$^{st}$ molar produced $\delta^{13}C$: -18.6, %C: 43.5, $\delta^{15}N$: 14.3, %N: 15.6, C:N: 3.3, $^{87}Sr/^{86}Sr$: 0.7130. Krzewińska, et al.[57] found that the mtDNA HVR1 sequence of this individual was estimated to represent haplogroup H*.

VK 530 (A4511) is described in the Schreiner database as brownish and being from an adult male of 35-35Y of age. The skull is said to be of Nordic type. Some animal bones were found together with the human remains. Russ[54] finds that the features of the cranium are indecisive, but that a humerus measurement indicates it to be a female. The lambdoid suture between occipitale and parietale is indented and irregular, and a light porosity can be seen on the occipital. Based on tooth wear she suggest an age of 25-35Y, which is not contradicted by the state of the bones. The four incisors and the canines in the mandible show signs of lack of space. An abscess has caused the loss of premolar dx maxilla and damaged the bone around pm2.

Naumann et al.[60] have measured the isotope values of a 3$^{rd}$ molar: $\delta^{13}C$: -19.8, %C: 42.4, $\delta^{15}N$: 11.3, %N: 14.6, C:N: 3.4[55]. Krzewińska, et al.[57] found the mtDNA HVR1 sequence of this individual to represent haplogroup H*.

**Ts5252, A4689, VK518, Nordland, Værøy, Nordland, (17/- Vågehamn)**

Askeladden ID: 59407

Coordinates: EU89-UTM zone 33 N7510519 E402848 Year of finding: 1954

The site is situated on the N end of the small island Værøy S of Lofoten, facing very rich fishing waters, but also one of the world's strongest whirlpools, Moskstraumen. The human remains originate from a grave with eight different individuals, buried at the same occasion. The grave was situated *c* 50 m from the beach, 4-5 m a.s.l. and was first discovered through the digging of a trench for a water pipe in August 1954. The remaining part of the grave was dug out in October the same year. It was reported that altogether remains of eight individuals were found, but detailed information is only available on the three of them. All remains were found inside a rectangular stone lining, oriented E-W and measuring 3.1 m in length and about one metre wide. It was not marked above ground. In the eastern part of the burial three parallel skeletons were found with their heads towards the east: In the northern side a child was lying facing north with some animal bones at the breast. In the southern side a male individual was lying on his back with his knees towards south and with an axe above his head; and in the middle a richly equipped female was placed with a number of dress accessories. In the western end of the burial another five persons with equipment were reported.

Among the burial goods the following items dates the burial: Two axe heads of Petersen type D and E, which according to Petersen[51] appears with sword types that Androshchuk et al.[53] date to the mid-$9^{th}$ to $10^{th}$ century; further two type C swords, which Androshchuk dates to the early Viking Age, AD 750-870. An oval brooch of type R. 647 is from the $9^{th}$ century [63], and the same date – with an emphasis on the early part of that century – is stipulated for a Berdal A type oval brooch also found in the grave[63]. The burial thus seems to have taken place around the middle of the $9^{th}$ century.

The Schreiner database describes the find under number A4689a, while A4689b is unrelated. It states that post-cranial bones from A4684 – but no cranium – have been intermixed with 4689a. Only one cranium is mentioned, and it is identified as female; this is the one that was sampled for VK518.

Russ has been examining the human remains stored under A4689 in the Schreiner Collection and found the remains of four individual, which probably reflects the three from the October 1954 excavation, plus the A4684 individual[54]. She found one female – only represented through a cranium – and three males, one of these being only 16-20Y. The female was, based on suture and teeth wear status, determined to be 25-40Y old.

**T9366-9379, A3708, VK524, Nordland, Nesna, Tommeidet (98/22)**

Askeladden ID: 16512-1

Coordinates: EU89-UTM zone 33 N7351266 E397879 Year of finding: *c* 1910

This boat burial was found in one of several mounds overlooking a natural harbour on the W tip of the island Tomme. Remains of two individuals were found in the burial, placed side by side in the SW-NE orientated boat. Horse- and cattle bones were found in the NE end of the grave[64]. Among the rather rich grave goods, the following datable items were found: A couple of oval brooches of copper alloy, of the type R. 652, which Petersen dates from the late $9^{th}$ to the late $10^{th}$ century[63]; a trefoil brooch type R. 671, a model which is also elsewhere found with R. 652[63]; four beads of black glass with white bands and white, red and green dots in glass added to the surface, datable to the #?#; a sword described as similar to R. 495, but with a smaller top rather than a regular knob on the upper end – this sounds like a type P or Y sword, dated by Androshchuk[53] to the late $10^{th}$ century; a shield boss type R. 562 dates according to Petersen[51] to 850-950; and an axe type R. 552, which Petersen finds together with swords of the B, C, E and H type; only the H-type of these continues into the $10^{th}$ century. The grave can probably be dated to the middle of the $10^{th}$ century.

The Schreiner database describes the human remains as greyish and the skull type as Nordic. The materials first came to the Anatomical Institute in 1927, probably when Schreiner was doing his survey for the Oseberg publication. The preserved remains from a male and a female are listed in Schreiner[58]. Russ could not identify any remains of the woman[54]. The surviving remains are from a male (cranium and DSP), 35-50Y old. The cranium shows a severe case of sinusitis frontale, which is suggested to have caused the man’s death. Further spina bifida occulta was observed on sacrum, with arcus missing on 4 and 5. Left femur shows evidence of non-malicious bone cancer. Krzewińska, et al.[57] determines the mtDNA HVR1 sequence to represent haplogroup HV0.

**T12578, A3709, VK525, Nordland, Dønna, Hov (19/1, Løkta)**

Askeladden ID: 73294

Coordinates: EU89-UTM zone 33 N7342211 E397137 Year of finding: 1922

This site is situated only 50 m from the current beach, overlooking an open, stony bay facing ENE. The find was made during the digging of a trench for a water pipe to a farm building. The finds from this burial are few, including a knife and a sickle, plus two oval brooches of types R. 647 and R649, the latter datable to the $9^{th}$ century[63].

In the Schreiner database the find is described as brownish and originating from an adult woman with a skull of Nordic type. It is more fully described in Schreiner[58]. Russ[54] finds it to belong to a female, 25-30Y of age, based on tooth wear. The skeleton is described as poorly preserved, but very gracile. Some skeletal parts from the upper breast region are coloured green by copper alloy, demonstrating that at least one of the oval brooches most likely had been positioned here. The following pathological features were recorded: A double facet on the atlas, and a type B deformation of its posterior arch. Calculus was prominently present and caries was found on two teeth. M1, sin, showed an abscess, and there were found indications of enamel hypoplasia.

Naumann et al.[60] got the following isotopic values: A sample from a femur produced the following isotope results: $\delta^{13}$C: -20.4, %C: 43.8, $\delta^{15}$N: 11.0, %N: 15.7, C:N: 3.2[55]. A sample from a 1$^{st}$ molar produced those: $\delta^{13}$C: -20.2, %C: 54.5, $\delta^{15}$N: 11.3, %N: 16.6, C:N: 3.2, $^{87}$Sr/$^{86}$Sr: 0.7092.

**C18558, A253, VK388, Nordland, Lødingen, Ytterstad (17/-)**

Askeladden ID: None

Coordinates: EU89-UTM zone 33 N7580530 E527347 (farm) Year of finding: 1889

The skeleton was found in a small mound at the farm Ytterstad, about 500 m from the beach. The farm is on the N side of a well sheltered bay facing E and overlooking the mouth of Tjeldsunde, a part of the inner sailing route along the Norwegian coast. The only accompanying object was an unidentified, ornamented antler tool. The ornaments indicate a date to the Viking Age or Middle Ages, but the burial type excludes the latter. The skeleton was initially identified as female, but was later determined to be a young male by Schreiner[58] who also notes that several of the metatarsals showed pathological changes.

Russ[54] describes the skeleton as quite complete, but poorly preserved and belonging to a young male age 15-17Y. The rearmost molar had erupted, but most epiphyses in the post-cranial skeleton had not yet fused. Coxae could not be used for sex determination due to young age, but the cranium allowed a certain identification. Unhealed cribra was found in both cranial orbits. Both feet and legs are showing patological possibly from a severe, enduring inflammation which might have caused the individual's death.

Naumann et al.[60] have recorded the following isotope results from a femur: $\delta^{13}$C: -17.8, %C: 45.5, $\delta^{15}$N: 14.9, %N: 16.5, C:N: 3.2[55]. A sample from a 1$^{st}$ molar produced: $\delta^{13}$C: -20.5, %C: 38.5, $\delta^{15}$N: 12.5, %N: 12.4, C:N: 3.6, $^{87}$Sr/$^{86}$Sr: 0.7175.

**C14554-14557, A1522, VK419, Nordland, Bodø, Rønvik Øvre (31/-)**

Askeladden ID: None

Coordinates: EU89-UTM zone 33 N7463888 E475796 Year of finding: 1889

This burial was found in shell sand in what had earlier been a gentle slope towards the nearby beach, but which had been eroded at the time of finding. There was no memory of a mound on the site. The grave itself were found below two stone slabs, 1.0 and 0.7 m long, which were lying in the topsoil. Apart from the skeletal remains, the grave held an axe of Early Iron Age type, but with similarities with Petersen's type C axe without 'beard', a large scythe type R. 386 and a sickle possibly of type R. 385. C-type axes are often found with H-type swords, which Androshchuk[53] dates to up to AD 950. The burial cannot be dated more closely than Migration Period to Early Viking Age.

The Schreiner database states that the skeleton is brownish in colour and from an adult male; its crania is described as being of Nordic type; the preserved bones are listed in Schreiner[58]. Russ[54] suggests that the individual is most likely a young man, 17-25Y, based on sutures, tooth wear and epiphyses[54]. Naumann et al.[60] analysed a sample from a femur which produced the following isotope results: $\delta^{13}$C: -17.7, %C: 43.0, $\delta^{15}$N: 14.5, %N: 15.1, C:N: 3.3[55]. A sample from a 2$^{nd}$ molar produced those: $\delta^{13}$C: -19.0, %C: 47.8, $\delta^{15}$N: 12.8, %N: 16.9, C:N: 3.3, $^{87}$Sr/$^{86}$Sr: 0.7109. Krzewińska, et al.[57] estimated the mtDNA HVR1 sequence to represent haplogroup U1/U5a.

**C18035-18039, A1502, VK418, Nordland, Steigen, Steigen øvre (77/1)**

Askeladden ID: None

Coordinates: EU89-UTM zone 33 N7535744 E499177 Year of finding: c. 1894

The find was made during drainage work on the present-day cemetery of Steigen, only about 300 m from the rather marshy coast. There are several Iron Age cemeteries in the surrounding area. It produced a weapon equipment, which has been argued to be of eastern origin[65]. A shield boss type R. 221, two spearheads of Rygh's type 206 and 211, and a sword with a decorated scabbard mounting dates this find firmly in the Early Iron Age, according to Sjøvold in the late 4$^{th}$ century.

The Schreiner database describes the skeleton as brownish and from an adult male with a cranium of Nordic type. The extent of the remains are described in Schreiner[58].

**Ts5656, A4727, VK547, Nordland, Tjelsund, Stokke (79/1)**

Coordinates: EU89-UTM zone 33 N7604398 E557131 sogn Lødingen Year of finding: 1957

Askeladden id.: 47377-1

The burial was found through the digging of a cable ditch, straight downhill 15 m N of the main building of the farm and 1 m N of the road between Sandnes and Kjærstad. The site is situated *c* 150 m E of Hol church, at Tjelsundet. A professional excavation of the remaining part of the burial was conducted by curator Simonsen from Tromsø Museum.

The grave was not marked on the surface, and the burial pit, dug in shell sand, was without stones and in the shape of a trough. It measured 30-50 cm in depth, and was 60 cm wide and 130 cm long. It was oriented E-W, and the body was lying on its side, with the head towards E, facing N, and with flexed legs. The sparse grave goods were found at the back of the legs, in the grave's SW corner. They consisted of an auger of iron and a possible awl of iron with a wooden handle. Further three iron tacks with wooden remains comes from an unidentified object. The finds can be dated no closer than to the Viking Age.

In the Schreiner database, the skeleton is described as yellowish and well-preserved. The age of the individual, who is determined to be male, has been estimated to 45-55 Y, and the body height to 180 cm. The cranium is described as being of Nordic type.

**Troms**

**Ts3639, A4184, VK520, Troms, Tromsø, Tussøy, (188/1)**

Askeladden ID: None

Coordinates: EU89-UTM zone 33 N7729512 E621341 Year of excavation: 1935

The smallholder Hjalmar Brox and his son Albert discovered this grave when digging in a small mound located *c.* 7 m a.s.l., *c.* 100 m W of the farmstead. On the behalf of Tromsø Museum a student, H. Haldorsen, excavated a 7 x 4 m large section of the mound and found evidence of three individuals, of which A4184 is individual III. Individual I and II had apparently been buried simultaneously, and from the grave equipment (weapons and tools) they were identified as male and buried in the 8$^{th}$ century; no human remains were reported salvaged. A4184 was in the grave first discovered by Brox, who had disturbed it severely, but the body had apparently been lying with the head towards west. The cranium and a few other bones were recovered from the burial together with two oval brooches of the types Rygh 652 and Rygh 654[62], found in the breast region. The grave dates according to Gjessing to the 10$^{th}$ century[66].

In the Schreiner Collection database the skeleton no. A4184 is described as well-preserved but incomplete and brownish in color. It is stated that it consists of a "very defect and deformed calva with associated right, defect upper jaw with cheek bone, plus a few further bones". The skeleton is judged to be from an adult female with a skull of Nordic type.

Russ[54] describes the skeleton as poorly preserved and with only few skeletal parts present. Age determination based on the $3^{rd}$ molar is 11-17 Y, while the remaining material indicates 12-19Y, most likely 16-18Y. Due to the young age, sex determination is uncertain, but features at the cranium indicate female. Apart from some calculus and a possibly caries in one canine tooth, no pathological features were observed.

**Ts3525, A4049, VK528, Troms, Tromsø, Tussøy, Trygstad i Bø (188/10)**

Askeladden ID: 27358-1

Coordinates: EU89-UTM zone 33 N7730580 E622625 Year of excavation: 1933

A4049 was handed in to Tromsø museum by a local from Tussøy, Tryggve Sletten, and he appears to be the source for the contextual information. The find was made 40-50 cm below the surface, at a place near a stream, 5-6 m a.s.l. The distance to the sea shore is c 50 m. The find included a skeleton in a supine position with the head towards W. It had a sword of Petersen's type C[51] on its right hand side and an axe on the left; supposed animal bones were also found at its side. Further finds from the grave were two arrowheads, a large, four-sided hone of slate, fragments of a knife, an iron staple and a decorated bone comb of type Rygh 447[62]. At the head and foot end of the grave, stone heaps were found, and 5-10 m from the grave, towards the river, charcoal and other unspecified material. Gjessing dates the grave to AD 800-850[66], but Androshchuk dates the C-type swords to the wider range AD 750-870[53] and the comb would fall into Ashby's Type 5, which broadly dates from the 8th to the mid-10th century[67]; Gjessing's narrow dating interval may thus be expanded to AD 750-870.

In the Schreiner Collection Database A4049 is described as a yellowish, well-preserved, incomplete skeleton with a defect cranium. The skeleton is estimated to be male and senilis, around 60 Y of age and with a skull of Nordic type. The body height is, following Trotter and Gleser, estimated to 179 cm.

In her examination of the rather well preserved skeleton in 2013, Russ estimated A4049 to be a male, 40-60 years of age. Of pathological features, she recorded several severe examples of lipping on the vertebras, especially in the lower back. A 28 mm long and 7-8 mm deep cut in the breast bone may be peri- or postmortem. Non-metric traits were three ossicles at lambda and several more in the lambdoid suture, and vastus notch in both patella[54].

A sample from a femur produced the following isotope results: $\delta^{13}C$: -16.0, %C: 44.4, $\delta^{15}N$: 17.3, %N: 15.5, C:N: 3.3. A sample from a $1^{st}$ molar produced those: $\delta^{13}C$: -19.9, %C: 44.8, $\delta^{15}N$: 12.3,

%N: 15.3, C:N: 3.4, $^{87}$Sr/$^{86}$Sr: 0.7118[55]. Naumann et al.[60] points out that the $^{87}$Sr/$^{86}$Sr value differs from known local values, indicating that the individual grew up at a different place than Tussøy. They further suggest a very significant change in diet from terrestrial to highly marine food in later part of life. Among 33 Merovingian and Viking age individuals from Northern Norway, analysed by Naumann et al.[60], A4049 was one of those digesting the most marine diet in his late years, while his childhood diet was among the most terrestrially dominated in the study.
Krzewinska et al.[57] estimate the mtDNA HVR1 sequence of this individual to represent haplogroup K*.

**Ts---- , A5001a, VK531, Troms, Lenvik, Skarsvåg (94/2)**
Askeladden ID: 48876-1
Coordinates: EU89-UTM zone 33 N7709381 E617444 Year of excavation: 1964
No museum number is recorded for this skeleton, but information can be found in the archives of Tromsø museum, file 214/64, and under accession number 1964/104. The grave was situated above the natural harbour Skarsvågen at the NW end of the island Senja, overlooking the inner sailing route along the Norwegian coast. It was archaeologically excavated by P. Simonsen in 1964, who found no objects in the grave and no indications of a mound above. From the grave form, Simonsen dated it to Viking Age, or perhaps a little earlier. However, recent $^{14}$C dating results showed that the sample was much older than initially thought ($^{14}$C date: 3918+/-36).
In the Schreiner Collection Database, the skeleton is described as well-preserved but incomplete, with a skull. It is said to be of varying color and is estimated to be of a male, senilis 55-65 Y of age. The skull is said to be of Nordic type.

#### Ukraine

**Inna Potekhina, Institute of Archaeology of the National Academy of Sciences of Ukraine**

**Chernihiv city, Chernihiv oblast (region), Ukraine**

Relevance for the Vikings: Rurik dynasty.

Gleb Svyatoslavich, «Gleb, son of Sviatoslav» (in Ukrainian and Russian languages endings -ovych, -evych after the name have a meaning «son of») was the 11th century prince of Tmutarakan/Novgorod, aged 25-35 according to anthropologist analysis.

Sources of information: Skull was found in 1967 in a tomb (stone sarcophagus) near the Saviour's Chernihiv cathedral (Fig. S1.3). It was an accident finding during construction works on the territory of State historical and cultural reserve «Chernihiv Starodavniy» (Ancient Chernihiv).

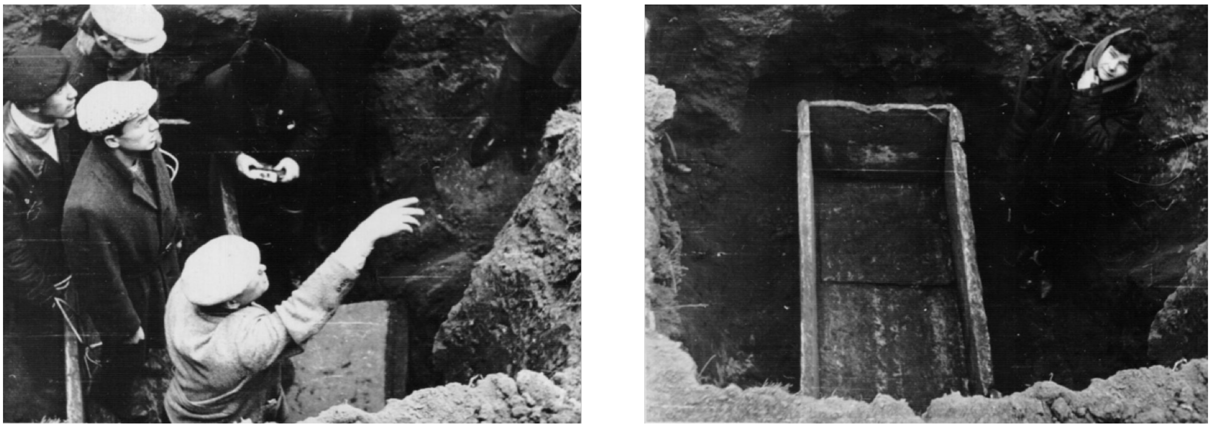

**Fig. S1.2**: Photos from scientific archives of State historical and cultural reserve «Chernihiv Starodavniy».

Prior to the date of finding there was no institution in Chernihiv (USSR) which could professionally make all the archeological data and stratigraphy. Therefore, there are no archeological reports of these findings in archives. The skull was sent to Moscow and returned back in a wooden box. For many years, it laid in this box in the archives and was rediscovered for science only in 2016.

Coordinates: WGS84: 51° 29′ 20.45″ N, 31° 18′ 28″ E; 51.489014°, 31.307778°

Dating (archeological only): The sarcophagus was found in the backyard of the cathedral. The depth of the upper plate of the sarcophagus is 1.9-2.0 meters from modern surface (Fig. S1.2). Stratigraphy is slightly damaged by the later breach of the soil: water supply pipes are laid 0.2 m above the headboard of sarcophagus. Archeologists date it to 11$^{th}$ century.

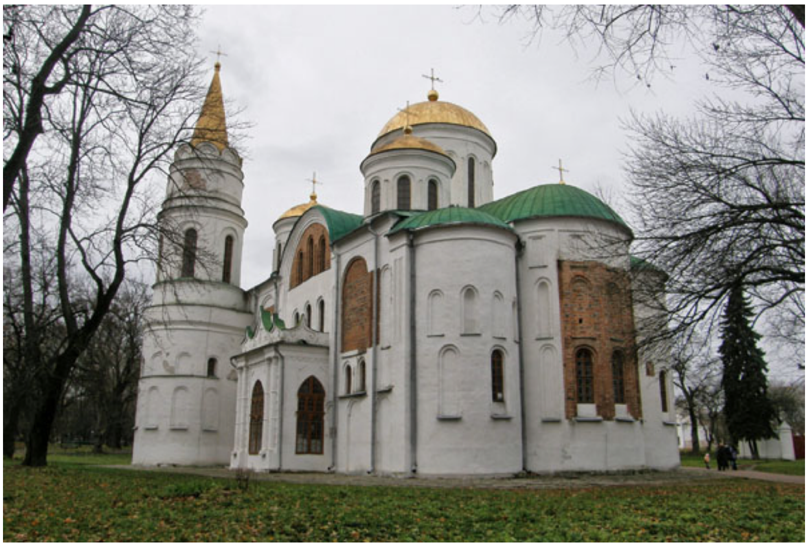

**Fig. S1.3**: Saviour's Chernihiv cathedral (11$^{th}$ century). Modern view of the cathedral and a backyard where sarcophagus was found.

Anthropological data: aged 25-35 years, traces of sword wounds are preserved on the skull. All anthropological measurements and indexes are in the article[68].

Samples used for DNA analysis
VK542 Ukraine_Chernigov

**Lutsk city, Volyn oblast (region), Ukraine**
Relevance for the Vikings: Rurik dynasty
Izjaslav Ingvarevych, «Izjaslav, son of Ingvar» (in Ukrainian and Russian languages endings -ovych, -evych after the name have a meaning «son of») was 13$^{th}$ century prince of Dorogobuzh, Principality of Volhynia/Galicia, aged 30-40 according to anthropologist analysis.
Sources of information: Skull was found in 1989 in a tomb on the territory of Lutsk castle (Figs. S1.4 and S1.5).
Archeology report is located in scientific archive of Lutsk State historical and cultural reserve (Научный архив Государственного историко-культурного заповедника города Луцк). Archive number: 203, 51 pages. Coordinates: WGS84: 50° 44′ 20″ N, 25° 19′ 23″ E; 50.738889°, 25.323056°

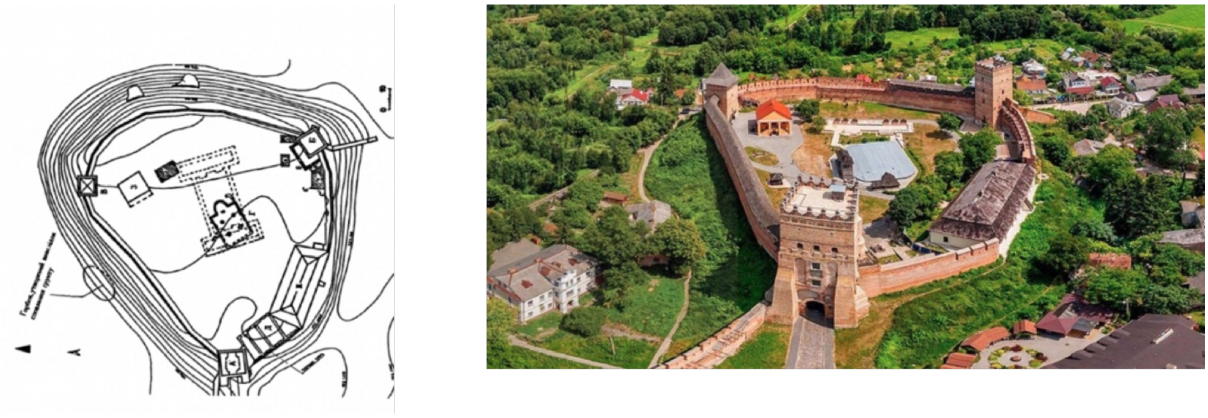

**Fig. S1.4**: Lubart's Castle, ruins of old Rus church (12th -13th century) on the territory of the castle. Today the ruins of the church and tomb are covered with metal construction (in the center of the castle) with inner access.

Dating (archeological only): given to the same church building material used for the construction of the burial chamber and stratigraphy data, burial can be dated to the end of XII - beginning of XIII century. Anthropological data: aged 30-40, traces of stab wounds are preserved on the skeleton, at the moment of finding in 1989 archeologists observed an arrowhead stuck in the skull. All anthropological measurements and indexes are in the article[69].

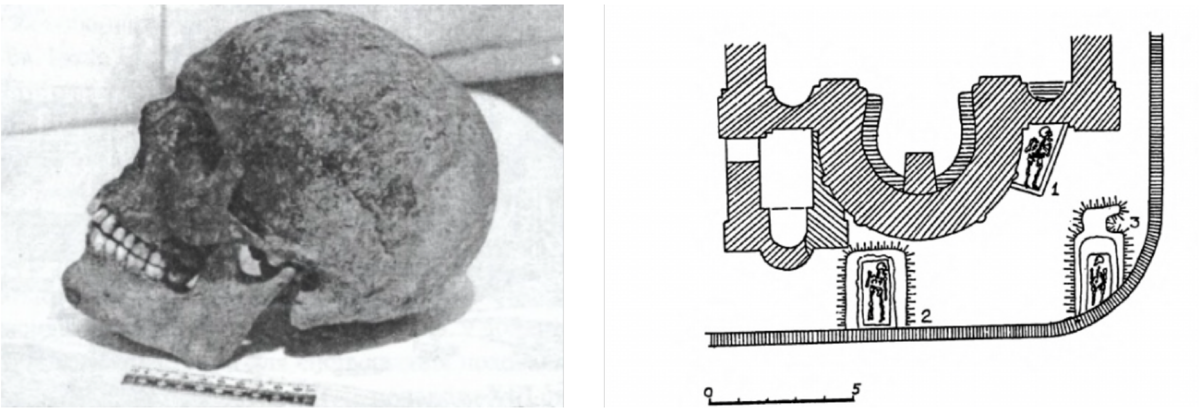


**Fig. S1.5**: Photo of the skull, year 1989 (left) and the schematic drawing of the tombs (right). No: 2 is Izjaslav's tomb.

Samples used for DNA analysis

VK541 Ukraine_Lutsk

**Shestovitsa, Ukraine**

The cemetery Shestovitsa refers to the archaeological complex of the IX-XII century, which consists of a hillfort and numerous burial mounds. It is located in the Korovel tract, near the village of Shestovitsa, on the right bank of the Desna river (the tributary of the Dnieper river), 12 km south-west from the city of Chernigov[70]. Coordinates are 51 ° 21'59.17 " N, 31 ° 10'46.04 " E, the altitude above the sea level is 116 m.

The first excavations of Shestovitsa were carried out in 1925-1927 by P.I.Smolichev. Since 1948 the research of the hillfort and the cemetery was resumed. The cemetery consists of several hundred burial mounds with a varied funerary rite. By now, more than 160 burial mounds have been excavated, including more than 60 with the cremation rite, more than 50 - with the inhumation rite, and the same number with the cenotaphs. Most of the burial mounds date back to the $10^{th}$ -$11^{th}$ centuries, and only a few - the $12^{th}$ century. In many cases, collective burials containing from 2 to 9 skeletons of men, women and children were found. Among the buried there are warriors with weapons and fighting horses. Rich burial inventory often included things of Scandinavian origin: weapons (battle axes, swords, spearheads, daggers, quivers with arrows); ornaments and household items (knives, combs, stucco and pottery); remnants of clothing, fasteners, brooches; Arabic and Byzantine coins. In Shestovitsa, apparently, the squad of the Kiev prince was deployed, in which Varangian soldiers were also members. Since the 1990s Shestovitsa archaeological complex is recognized by historians as one of the largest settlements of Vikings in Europe.

The question of the anthropological composition of the buried in Shestovitsa is debated. Earlier, the researchers attributed them to the Slavs, emphasizing a slight eastern component on the basis of flattening of the face in some male skulls. However, the weak horizontal profiling of the face here is not accompanied by other signs of eastern origin, which allows to deny the presence of a Mongoloid craniological complex and suggests the influence of the Finnish or laponoid component. The presence of immigrants from the Finno-Ugric lands in this region is consistent with the chronicle and archaeological data. It was also suggested that the Norman (Germanic) morphological component is mixed here with the Slavic. Some researchers refer the series from Shestovitsa to the Scandinavian cluster and find it remote parallels in the synchronous population of Sweden and Britain.

Despite a certain multi-vector anthropological connections of the population of Shestovitsa, Slavic and Norman should probably be considered the dominant components in its composition. The real ratio is difficult to determine due to the low representativeness of the series, but there is every reason to believe that the role of each of them was different for the male and female population, and women evidently represented a more homogeneous group of Slavic origin.

For genetic analysis, samples of bone tissue from 12 burials excavated by D.I. Bliefeld in 1956-58 from various mounds of Shestovitsa were transferred. Unfortunately, materials from only two burials proved to be suitable for DNA isolation:

Ukraine_Shestovitsa-8870-97 = kurgan 32(23)/ burial 2, pit Б, male, 25-30.

Ukraine_Shestovitsa-8871-96 = kurgan 32(23)/ burial 1, female, 35-40.

Samples used for DNA analysis

VK539 Ukraine_Shestovitsa-8870-97

VK540 Ukraine_Shestovitsa-8871-96

#### St John's College, Oxford, UK

**Ceri Falys, Thames Valley Archaeological Services**

In 2008, archaeological excavations in advance of development in the grounds of St John's College, Oxford, UK, revealed an unexpected sequence of deposits that shed new light on Oxford's prehistoric ritual past, a grisly episode in its history from the late Anglo-Saxon period (evidence of a massacre dating to around AD 1000), and more mundane but still interesting development from medieval occupation in three tenement plots, to a post-medieval farm and then the arrival of the college.

Of interest to this study, the excavation led to the discovery of a mass grave containing a minimum of 35 human skeletons. The majority of individuals were articulated and relatively complete, although several disarticulated limbs and a skull were also recovered. The remains were not deposited within a purposefully dug grave, but rather had been placed in a pre-existing depression in the landscape, resulting from the incomplete infilling of a portion of a monumental Neolithic henge ditch. The skeletons were deposited in the grave in a disorganized manner, and seemingly with little care or respect. There was no pattern to the positioning or orientation of individuals within the grave. Frequent intermingling of elements between individuals strongly suggest all skeletons had been deposited in a single episode. At the deepest portions of the grave, the bodies were piled four deep.

Osteological analysis found that all individuals, with the exception of two adolescents, were tall and robust adult males. All of them had met a violent death, as every man displayed evidence of perimortem sharp-force trauma (blade and puncture wounds), which was extensive and excessive in most cases. Many of the wounds were inflicted from behind, and few defensive wounds were observed. Most unusually, several skeletal elements had been partially burned. As a whole, the individuals in this assemblage appear to have an unusually high rate of non-fatal pathologies as well, both congenital and acquired through their lifestyle, and minimal evidence of antemortem trauma. It is concluded that this group, all robust adult males (bar the teenagers), mostly of the same generation, taller than average, sharing unusual pathologies, limited antemortem trauma, and extensive perimortem wounds, disposed of in this disorganized way, might well have been a physically distinct sub-group of the population, and had been massacred in a single event.

It is proposed that this is not a battle grave for fresh Vikings newly landed, but Danes who had been settled in Oxford for some years, possibly some of the younger ones may even be second generation. The unusual occurrence of charring of some skeletal elements examined in context with historical documentation points strongly towards these men being the Danish victims of King Aethelred's decree ordering their extermination in AD 1002, during the event known as the 'St Brice's Day

Massacre'.

It was hoped that radiocarbon dating would have confirmed the possibility of these men being the victims of the known massacre, unfortunately, these results have been inconclusive and somewhat puzzling. The dating initially appeared to preclude associating all of the skeletons with the St Brice's Day massacre, with three of the radiocarbon results indicating a perfect fit with a date of AD 1002. However, subsequent testing by a second laboratory returned three dates that appear much too early, which is puzzling and remains unresolved. Archaeologically, however, there is no possibility that the commingled human remains in the top of the Neolithic henge ditch could be interpreted as anything other than a single mass grave.

Isotopic results suggested the possibility that several individuals, including three of the men with the earliest radiocarbon dates, had an unusually high seafood diet for the inland location. It has been suggested that this marine reservoir offset is a contributing factor for the apparent discrepancy in the dating. Other isotope results indicated the majority of men likely originated either in the British Isles or the nearer continent, including western Denmark and Germany. One man suggests an origin in a colder average climate than the rest of the group (e.g. Denmark or southern Scandinavia), two have values well within the limits of the modern UK distribution, but could also be attributed to this northern region, and a further three men have values which would appear to be from somewhere warmer than the UK.

It is highly unusual for archaeology to capture evidence of a historically documented event, and archaeologists have usually been reluctant to make such connections, but it is here contended that, although there are some unresolved problems, the overwhelming probability is that this site presents just such an occasion. Of course other possibilities exist, however, the nature of the extensive perimortem trauma (suggestive of attacks on defenseless victims rather than battle casualties), charring of severed limbs, unusual shared congenital defects, a diet unusually high in fish proteins for the local population, the assemblage comprised of solely tall, robust, hard-working males, with limited previous history of violence, the jumbled nature of their burial and the location of the burial out of town in a prehistoric (pagan) monument all strongly support the hypothesis that these men were the Oxford victims of the St. Brice's Day Massacre[71].

Samples used for DNA analysis

VK163 UK_Oxford_sk 1870
VK146 UK_Oxford_sk 1785
VK147 UK_Oxford_sk 1864
VK148 UK_Oxford_sk 1787
VK149 UK_Oxford_sk 1852
VK150 UK_Oxford_sk 1866
VK151 UK_Oxford_sk 1963
VK172 UK_Oxford_sk 1968
VK173 UK_Oxford_sk 1978
VK174 UK_Oxford_sk 1984
VK175 UK_Oxford_sk 1996
VK164 UK_Oxford_sk 1872
VK176 UK_Oxford_sk 1990
VK177 UK_Oxford_sk 2056
VK178 UK_Oxford_sk 2057
VK165 UK_Oxford_sk 1876
VK166 UK_Oxford_sk 1891
VK167 UK_Oxford_sk 1898
VK168 UK_Oxford_sk 1899
VK143 UK_Oxford_sk 1951
VK144 UK_Oxford_sk 1756
VK145 UK_Oxford_sk 1783

#### Ridgeway Hill Mass Grave, Dorset, UK

**Louise Loe, Oxford Archaeology, UK** **Ridgeway Hill Mass Grave, Dorset, UK**

The Ridgeway Hill mass grave was a multiple burial of executed Vikings, which had been made sometime in the $10^{th}$ or $11^{th}$ century on the crest of Ridgeway Hill, near Weymouth, on the South coast of England. The grave was discovered in 2009 by Oxford Archaeology during the construction of the Weymouth Relief Road. The skeletons, around 50 in total, were predominantly young adult males, all of whom had been decapitated: heads had been deposited in a pile located at the southern edge of the grave, while the beheaded bodies had apparently been thrown in with little care.

The burial had been a one-off occurrence and had taken place at the time of, or shortly following, the men's execution, which had probably been performed at the graveside. The grave itself was a disused Roman quarry, used by the executioners for convenience rather than specifically dug for the purpose. It was an irregular ovoid feature, which by the time it was used as a grave, had partially silted up, the individuals buried to a maximum depth of less than one meter. In keeping with other Anglo-Saxon execution cemeteries, the proximity of the grave to a major road, a parish boundary and prehistoric monuments, would all seem to have been important. It is conceivable that the executions had been performed in front of a large crowd of spectators and as a formal event.

The men had been buried in a variety of positions and occupied no particular orientation (Extended Data Fig. 1b). Most were lying prone (face down) on a West-East alignment, but there was a lot of variation and entanglement giving the overall impression of a jumble of bodies which had been thrown in with little care, with no consideration to how they lay and no attempt to bury them with their heads. This strongly suggests that they had been buried by their executioners and that, possibly, there had been more than one executioner. Site plans and digital survey data were scrutinized to explore the order in which the men had been buried, but no pattern could be discerned suggesting they may have been thrown into the grave from all sides.

It would seem that the men had been stripped prior to their execution, because no dress fastenings or other remains of evidence for clothing were found. There was also no evidence to suggest that the men's hands had been tied, or that any other form of restraint had been used. The probable Icelandic Jomsvikings saga describes the execution by beheading of a number of warriors who did not have their hands tied, but were roped together. Perhaps this had been the same here.

Analysis was unable to determine exactly how many men were executed and buried in the grave, because of some commingling and loss of bones as a result of modern disturbance. By employing counting and pair-matching exercises, it was estimated that between 47 and 52 individuals were present. There were more beheaded skeletons than skulls and this could mean that some heads had been taken as trophies following the executions (Extended Data Fig. 1b).

All of the men had suffered horrific ends, their executions an ugly affair involving excessive violence. The decapitations, probably performed with a sword, were evidenced by wounds that were concentrated in the region of the neck indicating that, in most cases, it had taken several attempts, from a variety of angles, to remove their heads. Blows intended to decapitate had been delivered from as high up as the back of the head to as low down as the shoulder blades, suggesting they had not been very well performed or organized. Approximately 188 wounds were observed on all of the skeletons, that is an average number of almost four wounds per individual. In addition, sharp force lesions were present on the forearms and tops of skulls of some individuals and may have been defense injuries and incapacitating injuries.

Isotope analysis performed on a number of skeletons[72], suggests they were a disparate group of people in terms of their origins, migratory histories and dietary habits, although a general emphasis on Arctic and sub-Arctic areas of Scandinavia, northern Iceland, the Baltic States, Belarus and Russia, and on terrestrial food sources, are suggested. It would appear that the majority were not living in the British Isles in the years leading to their deaths.

Although most of the men were young adults (18-25 years old) when they died the youngest was in their early or mid-teens and the oldest, over 50. They possessed features, particularly those relating to height and facial appearance, that were similar to Scandinavian populations of similar date. At least one individual had filed his teeth, seen as horizontal grooves on his front, central, upper incisors, possibly as a status symbol or a marker of occupation. In addition, evidence for infection and physical impairment was frequent for a group of predominantly young individuals who had died in their prime of life, although none of the skeletons showed convincing evidence for previous war wounds[73].

There are no historical records that directly link the mass grave with an event, but there are a number of possibilities, such as the ravaging of Portland in 982, or Viking attacks in Dorset in 998, 1015 and 1016. Although it seems very likely that these were Vikings executed by the English, the possibility that they were a group of mercenaries fighting for the English and executed by Vikings cannot be entirely ruled out. Other possible scenarios are that the men were merchants or recent settlers in England who were sentenced to judicial execution by the English authorities, were victims of the St Brice's Day massacre (1002) or were hostages or combatants engaging in reprisals against previous enemies during the reign of Cnut (1016-35).

In total, 10 skulls were sampled for DNA analysis for the present project. The skull with the filed teeth – 3736 – is among them.

Samples used for DNA analysis

VK256 UK_Dorset-3722
VK257 UK_Dorset-3723
VK258 UK_Dorset-3733
VK259 UK_Dorset-3734
VK260 UK_Dorset-3735
VK261 UK_Dorset-3736
VK262 UK_Dorset-3739
VK263 UK_Dorset-3742
VK264 UK_Dorset-3744
VK449 UK_Dorset-3746

**Acknowledgements**

We thank Richard Breward and Jon Murden from the Dorset County Museum for allowing access to the assemblage for DNA sampling.

#### Västergötlands museum IM16-107025, Varnhem RAÄ 60, Västergötland, Sweden

**Maria Vretemark, Västergötlands museum**

Varnhem in the central part of Västergötland is well known for its large church with the adjacent ruins of a Cistercian abbey. Less attention has been paid to the history of Varnhem before the abbey was founded in the middle of the twelfth century. In order to learn more about this earlier period, the Museum of Västergötland started an archaeological research project in 2005 named '*Varnhem – innan munkarna kom'* (Varnhem – Before the monks arrived). The archaeological excavations revealed a large settlement area and a Viking Age church with a surrounding churchyard. Thick cultural layers, foundations of buildings, a church ruin, and hundreds of Christian Viking Age graves were discovered. All of this presents an image of a prominent farmstead with roots going far back into the Iron Age.

Varnhem is situated in an area with a great density of prehistoric sites. Graves and settlements dating from all periods are found here. Fertile soil, rich pasture and meadowland for harvesting and grazing, together with woodlands and lakes provided the right conditions for the emergence of a strong wealthy community. Among the rich archaeological finds discovered in the area, the large silver treasure found in 1873 deserves particular mention. This Viking Age hoard consists of 476 silver coins from the early 11$^{th}$ century. Most of the coins are Anglo-Saxon and they point to contact with the West. Rune stones in the region bear witness of men killed in England. Several rune inscriptions also mention '*thegnar*' - a title of a follower of the Danish kings Sven Forkbeard and Cnut the Great, and as such a member of the English/Danish royal forces in England after the conquest in 1015 and the subsequent Danish occupation. Some of these *thegnar* obviously came from Västergötland, a region that had long been part of the Danish sphere of influence. The most successful Viking soldiers might have received a share of the taxes known as the *Danegeld*, which was paid in silver coins. They returned home after their service ended, and this could explain how the large number of Anglo Saxon silver coins ended up in Varnhem.

The presence of a large Iron Age farmstead was confirmed through the discovery of the remains of house constructions, hearths, trenches, pits and postholes, along with pottery, animal bones and other artefacts. A series of radiocarbon dates indicated that the site had been continuously settled for a thousand years, from the Roman Iron Age to Early Medieval times. This was most likely an aristocratic manor. A church, built at the expense of the landowner, was included as one of the buildings on the prominent farm.

In past decades, remains of early churches and early Christian burial grounds have emerged in several places in central parts of Västergötland. The oldest churches in this region have been dated to the period around the year 1000. They were privately built farm churches, predating the centralised church organisation of the twelfth century with its system of parish churches based on a territorial division of the landscape. The foundations of this private church were excavated as part of the archaeological project at Varnhem (Extended Data Fig. 1d). The first church in Varnhem was built in the late $10^{th}$ century. It was a small wooden church. Sometime during the period 1030-1050 AD the wooden church was replaced by a larger church built of locally quarried limestone. This church was probably one of the first stone buildings in Sweden.

Surrounding the foundations of the church in Varnhem, there is an extensive, nearly 4000 $m^2$ Christian burial place containing at least 2000 graves, and perhaps as many as 3000. Approximately 350 graves have been excavated so far and a well-preserved assemblage of human bones has been recovered for osteological analysis. The rest of the graves remain untouched under the grass of the park. The graves at Varnhem exhibit signs of a socially stratified society. Members of the family that owned the magnate's farm were buried closest to the church. This is indicated by the presence of limestone coffins. Further away from the church, the dead had been interred in wooden coffins or in simpler graves without a coffin. The social division of the churchyard is also reflected in the state of health that can be observed in the skeletons. There was also a division by gender. Men were buried to the south of the church and women to the north. The dating of the graves is extremely interesting. The burial ground at Varnhem was used continuously for Christian burials from the first half of the tenth century to the end of the twelfth century, a period of 250-300 years. Christianity was obviously established in the area already by the middle of the tenth century at the latest[74].

Samples used for DNA analysis

VK29 Sweden_Skara 17

VK30 Sweden_Skara 105

VK31 Sweden_Skara 194

VK33 Sweden_Skara 175

VK34 Sweden_Skara 135

VK35 Sweden_Skara 118

VK39 Sweden_Skara 181

VK40 Sweden_Skara 106
VK42 Sweden_Skara 62
VK303 Sweden_Skara 27
VK304 Sweden_Skara 36
VK306 Sweden_Skara 33
VK308 Sweden_Skara 101
VK309 Sweden_Skara 53
VK395 Sweden_Skara 275
VK396 Sweden_Skara 166
VK397 Sweden_Skara 237
VK398 Sweden_Skara 231
VK399 Sweden_Skara 276
VK400 Sweden_Skara 236
VK401 Sweden_Skara 229
VK402 Sweden_Skara 38
VK403 Sweden_Skara 217
VK404 Sweden_Skara 277
VK405 Sweden_Skara 83
VK406 Sweden_Skara 203
VK407 Sweden_Skara 274
VK424 Sweden_Skara 273
VK425 Sweden_Skara 44
VK426 Sweden_Skara 216
VK427 Sweden_Skara 209

#### Skämsta, Sweden

**Caroline Ahlström Arcini, National Historical Museums, Lund**

Two Dwarfs: Upplandsmuseet, inventory number - UM36031_621 and UM36031_623b

In the year 1994 parts of a burial site were investigated in the small hamlet Skämsta north of Uppsala in Sweden. Six inhumation graves containing skeletons from seven individuals were excavated. In addition to the more intact graves, a pit of mixed bones was found in the northern part of the cemetery. These bones came from a damaged grave. The excavated graves were not visible above ground. Four of the graves however were constructed of stone slabs of various size. Five of the individuals were orientated west-east and one was buried east-west. All of them were buried lying on their backs. They were all single graves but in one grave bones from a newborn was found.

Different types of artefacts were found in the graves such as combs, pearls, a ring for the hair. Regarding the ring there is one parallel found in Poland. The graves have, based on the finds, been dated to the later part of the Viking age and beginning of the medieval period (AD 1000-1100). Estimation of age and sex has been done according to standards[75] and the results indicate that five of the buried were adults, one was sub-adult 12-14 years of age and there was also a newborn. As for the adults, two died when they were 20-40 years of age and four between 40-60 years. Regarding the sex of the individuals three of them were men, three were women and the sub-adult was a boy.

Already during the excavation, the archaeologists noticed that there was something unusual with one of the graves (grave 41850, a.k.a. UM36031_623b). It contained a fully grown individual, however the stature was very small, about 130 cm. The osteological analysis of the skeletal material revealed that the individual was a dwarf. Further investigation of the rest of the skeletons showed that this was not the only dwarf. The individual in grave 33124 (UM36031_621), buried just north of the dwarf in grave 41850 was also abnormally short. A closer examination of the bones of these two individuals showed that except for short deformed limbs and joints their vertebras were very flat resulting in a short trunk[76]. To determine the underlying cause to why the individuals were so short and why their limbs were so deformed, a radiologist was consulted at the X-ray department in Lund. X-rays were taken of several parts of the skeletons and the diagnose was that they have suffered from a disease called *Spondyloepifyseal dysplasia (SED)*[77]. Except for the short stature and deformed limbs, the individuals affected by SED could have a flat face, occasionally a cleft palate and/or club foot. Approximately 50 per cent have myopia and/or retinal detachment.

*Spondyloepifyseal dysplasia* congenita is usually inherited as an autosomal dominant condition. It could either arise due to a mutation or appear in a child of an affected parent. If one parent is affected, the risk of inheritance is 50 percent and if both parents are affected the risk is 75 percent. There are three possible family relationships for the dwarfs at Skämsta. They are either brother and sister, mother and son or father and daughter.

Samples used for DNA analysis

VK517 Sweden_Uppsala_UM36031_623b

VK527 Sweden_Uppsala_UM36031_621

**Ljungbacka (Bronsdolken 7, Lockarp parish, Malmö, Sweden)**

**Ingrid Gustin, Dept. of Archaeology and Ancient History, Lund university, Sweden**

**Malmö Museum**

The burial ground of Ljungbacka is situated in south-west Scania, a few kilometres east of the coast. In the 1970s and 80s parts of a major cemetery were excavated close to a Bronze Age burial mound. The total number of excavated burials reached 191; 31 were inhumations and 160 cremations. The cremations were dated to Late Iron Age, while the inhumations were dated to the Late Iron Age, most likely Viking Age. Six of the inhumation burials contained two or more individuals. The upper-lying individuals were often placed in opposite direction versus the underlying[78].

| | | |
|---|---|---|
| New grave number | MHM 6031: 26A | MHM 6031: 28 |
| Old grave number | 4A | 0.250694444 |
| Date | Viking Age | Late Iron Age, probably Viking Age |
| Grave type | inhumation | inhumation |
| Position | southwest-northeast | northeast-southwest |
| Age | 55-75* | 35-40 |
| Sex | female* | male |
| Stature (cm) | 167 | 170 |
| Artefacts | keys, knife, whetstone | knife, whetstone, pottery |
| General comments | Individual placed in prone position, head facing south. Found above a coffin with a supine individual in east- west orientation. | The lower of two burials. This individual had supine position with head facing north. The filling of the grave contained unburned skeletal remains of a young man |

Samples used for DNA analysis

VK108 Grav4A_MHM 6031: 26A

VK217 Grave6_ MHM 6031: 28

#### Öland, Sweden

**Helene Wilhelmson, Sydsvensk arkeologi AB, Kristianstad, Sweden / Archaeology, Dept of archaeology and ancient history, Lund University, Sweden**

##### Öland in the Late Iron Age (AD400-1050)

In the island of Öland in the Baltic Sea, many human remains from the Late Iron Age have been excavated from burials and other contexts. Throughout the period the burials are both cremations as well as inhumations. There are considerable variations in inhumation burial form (pit coffin, stone cist etc) during the Viking age[79–83]. The uncremated human remains from burials (and other contexts) were recently studied using an interdisciplinary bioarchaeological perspective[83] integrating new radiocarbon dates of many graves. The individuals studied for aDNA here are the majority of the late Iron age population discussed in that study.

The most recent dietary isotope analysis of human remains, show a great individual variation in diet[5] supporting the archaeozoological finds and point towards a population with highly varied subsistence strategies. First generation migration to Öland was investigated through $^{87}Sr/^{86}Sr$ and $\delta^{18}O$ isotopes and the results were interpreted to show extensive immigration to the island with 68% non-local individuals in the Late Iron Age. The immigrants appear to be both regional and interregional. The greater variation in individual diet could not be concluded to correlate to provenance of an individual. The society and people living in Öland has therefore been interpreted as population of mixed provenance resulting in a creolized society with a mixing of different non-local and local traditions for burial and subsistence practice[83].

##### The Viking Age burials sampled for this study

These 29 individuals were included in the study of Wilhelmson[83] and consist of all types of burials. They are from 20 sites in Öland, excavated on separate occasions between the years 1931-1975. About half of the individuals (15) are dated by $^{14}C$ and the rest are dated by typology. The burials are inhumations of varied type. They have different orientation (EW, NS or SN), include different architecture (lime stone cists, pits, coffins, full boat burial) and single as well as multiple burials in one grave. Two more burials in this study (id 1099, 1052) are from the Early Iron Age. A table overview below presents the selected individuals details.

#### Samples used for DNA analysis

| Sample | Id | Parish | SHM (grave field) | Grave id | 14C calib. | Year of excavation | local/non-local? (from provenance isotopes) | Age of death | Sex |
|---|---|---|---|---|---|---|---|---|---|
| VK332 | Oland_1088 | Smedby | 23267 | 3 | 858 ±68 AD | 1944 | local | 45-60 | Male |
| VK333 | Oland_1028 | Vickleby | 22486 | | 885 ± 69 AD | 1939 | non local | Mature | Male |
| VK334 | Oland_1058 | Gärdslösa | 28364 | 134 | 1049 ± 58 AD | 1964-66 | gray | Mature | Female |
| VK335 | Oland_1068 | Långlöt | 29352 | 18 | | 1968 | non local | Young-mature | Male |
| VK336 | Oland_1075 | Gärdslösa | 28364 | 164 | 853 ± 67 AD | 1965 | local | Mature | Male |
| VK337 | Oland_1064 | Hulterstad | 22394 | | 858 ± 68 AD | 1939 | local | Mature | Male |
| VK342 | Oland_1016 | Hulterstad | 25096 | | | 1954 | non local | 70+ | female? |
| VK343 | Oland_1021 | Sandby | 26454 | 3 | | 1959 | local | Young/mature | female? |
| VK 344 | Oland_1030 | Södra Möckleby | 25657 | | | 1956 | non local | Mature-old | Male? |
| VK345 | Oland_1045 | Långlöt | 29352 | 24 | | 1968-1969 | non local | 60+ | Male |
| VK346 | Oland_1057 | Långlöt | 29352 | 13 | | 1968 | local | Old | Male |
| VK347 | Oland_1065 | Köpinge | 6393/75 | 3 | | 1975 | non local | Mature-old | Female? |
| VK348 | Oland_1067 | Köpinge | 6393/75 | 20 | | 1975 | non local | Old | Male |
| VK349 | Oland_1073 | Kastlösa | 27771 | 1 | 829 ± 57 AD | 1960 | non local | 55-65 | Male |
| VK350 | Oland_1086 | Långlöt | 29352 | 25 | 799 ± 68 AD (1210+-45 uncal BP) | 1968-73 | non local | Ca 38 | Female |
| VK352 | Oland_1012 | Runsten | 28549 | | | 1966 | non local | 35-45 | Male |
| VK353 | Oland_1024 | Smedby | 25129 | | 1049 ± 58 AD | 1954 | non local | 16 | Female |
| VK354 | Oland_1026 | Hulterstad | 19726 | 1 | 986 ± 38 AD | 1931 | gray | 60 | Male |
| VK355 | Oland_1046 | Ventlinge | 22291 | - | 847 ± 65 AD | 1939 | local | Ca 20 | Male |
| VK357 | Oland_1097 | Kastlösa | 22763 | - | 1053 ± 60 AD | 1941 | non local | mature | Female? |
| VK358 | Oland_1105 | Gärdslösa | 28364 | 108:I:169 | 853 ± 71 AD | 1965 | non local | Young | Male? |
| VK359 | Oland_1130 | Böda | 22231 | A7 | 700-800 AD | 1937 | NA | Over 15 | Undet. |
| VK379 | Oland_1077 | Böda | 22231 | A8 | 700-800 AD | 1937 | non local | juvenile-young | Undet. |
| VK380 | Oland_1078 | Böda | 21367 | A5 | | 1935 | gray | 12-15y | Undet. |
| VK382 | Oland_1132 | Böda | 22231 | A7 | 700-800 AD | 1937 | NA | Over 15 | Undet. |
| VK442 | Oland_1008 | Smedby | 24542 | I undre (=lower) | 847 ±-65 AD | 1951 | non local | 38 ± 10 | Female |
| VK443 | Oland_1101 | Böda | 21367 | 24 | | 1935 | non local | Ca 23 | Male |
| VK444 | Oland_1059 | Smedby | 23494 | 20 | 847 ± 65 AD | 1945 | gray | 19 | Male |
| VK522 | Oland 1052 | Stenåsa | 24846 | 1 | 386 ± 80 AD | 1953 | local | 19 | Male |
| VK533 | Oland 1076 | Gärdslösa | 28364 | 136 | Viking age | 1965 | non local | 50-60 | Female |
| VK579 | Oland 1099 | Mörbylånga | 1785/67 | 5 | AD 200-400 | 1966-67 | local | 18-19 | Female |

#### Kärda, Småland, Sweden

**Jörgen Gustafsson, Jönköpings Läns Museum**

Near the village of Nästa in Kärda parish, around 10 kilometers west of Värnamo in central Småland, several graves from the Late Iron Age have been investigated on a number of occasions. In the surrounding area, there are other grave fields with a total of 100 mounds and 40 circular stone settings. There are also graves (mounds and large stone settings) from the Bronze Age and Early Iron Age in the area, but not at all to the same extent. The local ancient monument environment including the grave fields has been described from around the year 1800 and onwards. In addition to these reports, sections of the grave field Kärda 42b were partially investigated in connection with the building and widening of Road 27 on three occasions: 1936–1937, 1990 and 2015.

Most ancient monuments in the parish date back to the Late Iron Age[84]. The parish, just as the region as a whole, is characterized by a large number of mounds and burial mound fields. Within the parish, there are 21 known grave fields comprising a total of 471 mounds and 158 circular stone settings.

When Road 27 was built between Värnamo and Smålandsstenar in November 1936, human bones were found, which led to an archaeological investigation of the site. During the excavations, 13 graves were found, of which nine held skeletons. Four were cremation graves, of which three were found in a mound with a central cairn dating back to the Early Bronze Age. The age of the primary grave was determined by the find of a dagger-shaped fire-striking flint-stone. The grave field was located on a marked elevation along the old road. The southern part of the grave field was investigated and removed in 1936–1937. At the excavations in 1990, when the road was widened, the remaining parts of the grave field were removed on the southern and south-western slope of the elevation, and a further nine graves containing skeletons were discovered. In 2015, in connection with the laying of a water supply pipe on the south-westernmost part of the grave field, a further grave with skeletal remains were found and investigated.

The graves consisted of slightly rounded mounds with shallow ditches around them and inner stone packing covering the buried skeletons. In some cases, it was found that the dead were buried in coffins. All individuals were adults, with the exception of a boy of 7 to 11 years old. The graves contained few artifacts, including knives, a sandstone whetstone, a gold foil bead and a bronze buckle from the 11th century. Only three $^{14}$C-datings have been made, of which one showed Middle Viking Age and two Late Viking Age, around the year 1000. It was possible to determine that the graves containing skeletons were oriented in an east-west direction and constructed according to Christian tradition. It is quite likely that they are from the era of overlap between pagan and Christian traditions.

In Smålandsstenar, only some 30 kilometers from Kärda, investigated tumuli of approximately the same age were found to be cremation graves of the pagan tradition, which indicates that there were local differences when it came to the transition to Christian burial practices[85].
The samples are taken from the graves that were investigated in 1990.

Samples used for DNA analysis
VK265 Sweden_Karda 17
VK266 Sweden_Karda 19
VK267 Sweden_Karda 21
VK268 Sweden_Karda 22
VK269 Sweden_Karda 24
VK270 Sweden_Karda 25

#### The Swedish History Museum in Stockholm SHM 16098/Gotland's Museum Inv.nr. GFC12675, Kopparsvik, Gotland, Sweden

**Sabine Sten, Department of Archaeology and Ancient History Campus Gotland, Uppsala University, Sweden**

**Kopparsvik**

A large number of graves dated to the Viking Age, 800-1150 AD, have been located at the coast of the Swedish island of Gotland which has always held an important geographic central position as an island in the Baltic Sea. Archaeological finds show artefacts that are not locally manufactured on the island, from as early as the Stone Age, 7000-1800 AD, when the island was visited by foreign peoples who brought their own merchandise. The Viking Age was a very intense historical period for Gotland. Archaeological finds such as jewelry and coins, mainly manufactured by silver, but also gold, show traces of trade and exchange contacts with countries mainly from the east, south, and west. Valuable shiploads also contributed to commotion. Perhaps visiting merchants who never returned home may have been buried on the island, and Gotlandic merchants may never have returned home from their travels.

One of the larger grave fields along the coast, just south of the Visby harbour, is Kopparsvik, Visby 76:1, (The Swedish History Museum in Stockholm SHM 16098, and Gotland's Museum Invnr. GFC12675). An area that due to several forms of exploitations has been excavated a number of times under Oscar Vilhelm Wennersten, during the years 1917 and 1918, Greta Arwidsson in 1956, Erik Nylén in 1963 and Hilkka Pettersson during the period of 1964-1966. The grave field, that has been fully excavated, is estimated to have been approximately 120 x 60m and divided into two areas, a northern and a southern region (Fig. S1.6). Skeletal remains and archaeological artefacts have been collected from 330 excavated and examined graves. It has, based on the approximately 1000 archaeological artefacts, been possible to date the grave field to 900-1050 AD. The archaeological artefacts illustrate both Gotlandic manufacturing and traces of long distance trade. Several papers of the archaeological studies e.g. by Pettersson[86], Westholm[87] and Toplak[88] as well as osteological examinations by Arcini[89,90] have been written about Kopparsvik.

Out of the approximately 330 graves excavated at the Kopparsvik grave field in the 1900's, 174 have been handpicked for an osteological analysis between the years 2013 to 2015, in the research project *Vikingatida Fröjel and Kopparsvik- hälsa, gravläggning och härkomst* (Andersson 2015 unpubl.) at Uppsala University Campus Gotland.

No archaeological trace of settlements has been found in the vicinity of Kopparsvik, and the environment does not allow for farming. Kopparsvik has, therefore, been interpreted as a place of commerce. In correlation to a research project concerning health in archaeological skeletal materials isotopic investigations were performed on fifteen handpicked individuals from Kopparsvik, at the Archaeological Research Laboratory, Stockholm University. The measuring of coal and nitrogen isotopes allowed us an insight of how the diet of the Kopparsvik population may have looked. Were they fishermen who lived mainly on a marine diet, such as fish and seal, or were they farmers who consumed terrestrial diet, such as cattle, sheep, pigs, or terrestrial wildlife. The isotopic analysis results showed an exclusively terrestrial diet (oral report from Professor Kerstin Lidén, Stockholm University).

Before the DNA examination was performed, detailed dental documentation and dental x-ray was performed. The dental examinations were performed at the Osteological laboratory, department of Archaeology and Ancient History, Uppsala University Campus Gotland in cooperation with the Department of Cardiology, Institute of Odontology, Sahlgrenska Academy at University of Gothenburg. The dental x-ray examinations were performed at the dental practice *Tandläkarpraktiken*, Adelsgatan in Visby.

The skeletal remains are generally very well preserved, which contributes to the good documentation conditions for sex- and age assessments, stature estimations, and notes of pathological changes (Andersson 2015 unpubl.).

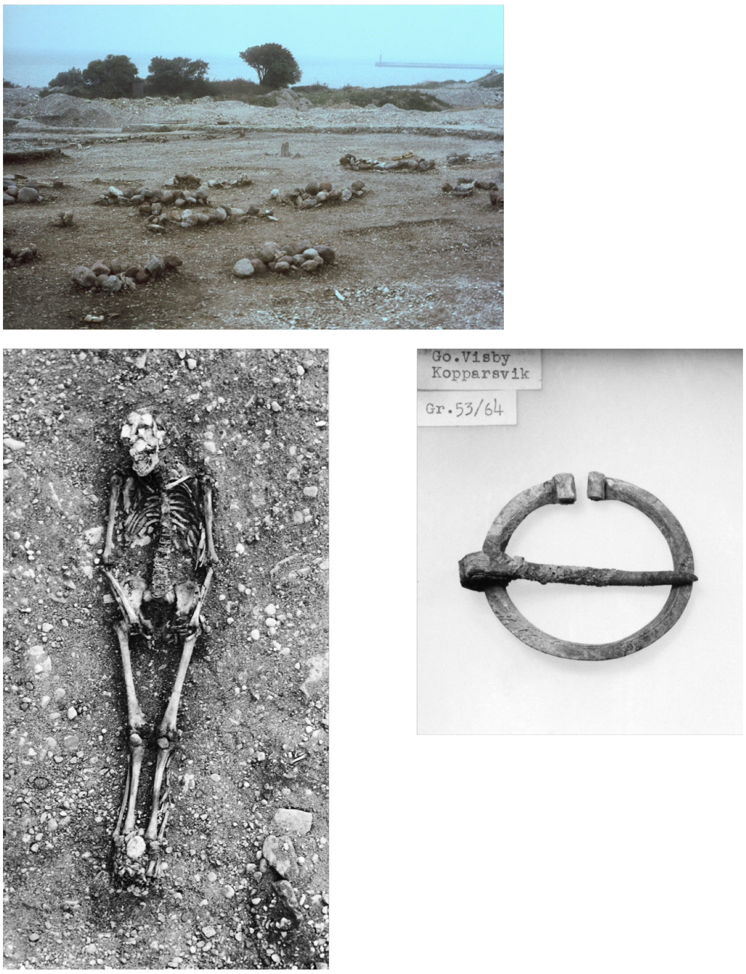


**Fig. S1.6**: An overview of the site Kopparsvik (top); skeletal remains (lower left) and a Viking age artefact (lower right) excavated in grave 53 in the burial site Kopparsvik.

Samples used for DNA analysis

VK48 Gotland_Kopparsvik-212/65

VK50 Gotland_Kopparsvik-53.64

VK51 Gotland_Kopparsvik-88/64

VK53 Gotland_Kopparsvik-161/65

VK232 Gotland_Kopparsvik-240.65

VK251 Gotland_Kopparsvik-30.64

VK450 Gotland_Kopparsvik-35

VK452 Gotland_Kopparsvik-111

VK453 Gotland_Kopparsvik-134

VK454 Gotland_Kopparsvik-140

VK467 Gotland_Kopparsvik-181
VK468 Gotland_Kopparsvik-235
VK469 Gotland_Kopparsvik-260
VK471 Gotland_Kopparsvik-63
VK472 Gotland_Kopparsvik-112
VK473 Gotland_Kopparsvik-126
VK474 Gotland_Kopparsvik-137
VK475 Gotland_Kopparsvik-187
VK476 Gotland_Kopparsvik-225
VK477 Gotland_Kopparsvik-228
VK478 Gotland_Kopparsvik-271
VK479 Gotland_Kopparsvik-272

**Fröjel**

Gotland's position in the Baltic Sea, with mile long coasts (800 km) and the vicinity water has been of great importance to the people on the island for centuries, both regarding fishing, seal-and bird hunting and marine commerce. A large number of fishing hamlets and larger harbors has been located along the coasts. Fröjel, Bottarve/Nymans, placed on the western coast of the island of Gotland, was one of the largest and most important harbors and commerce sites during the Viking Age and the early Middle Ages. Other important contemporary places of commerce, in the Baltic Sea area, was Birka in Uppland, Sweden, Hedeby in northern Germany, Wolin in northern Poland, and Grobin in south west Lithuania.

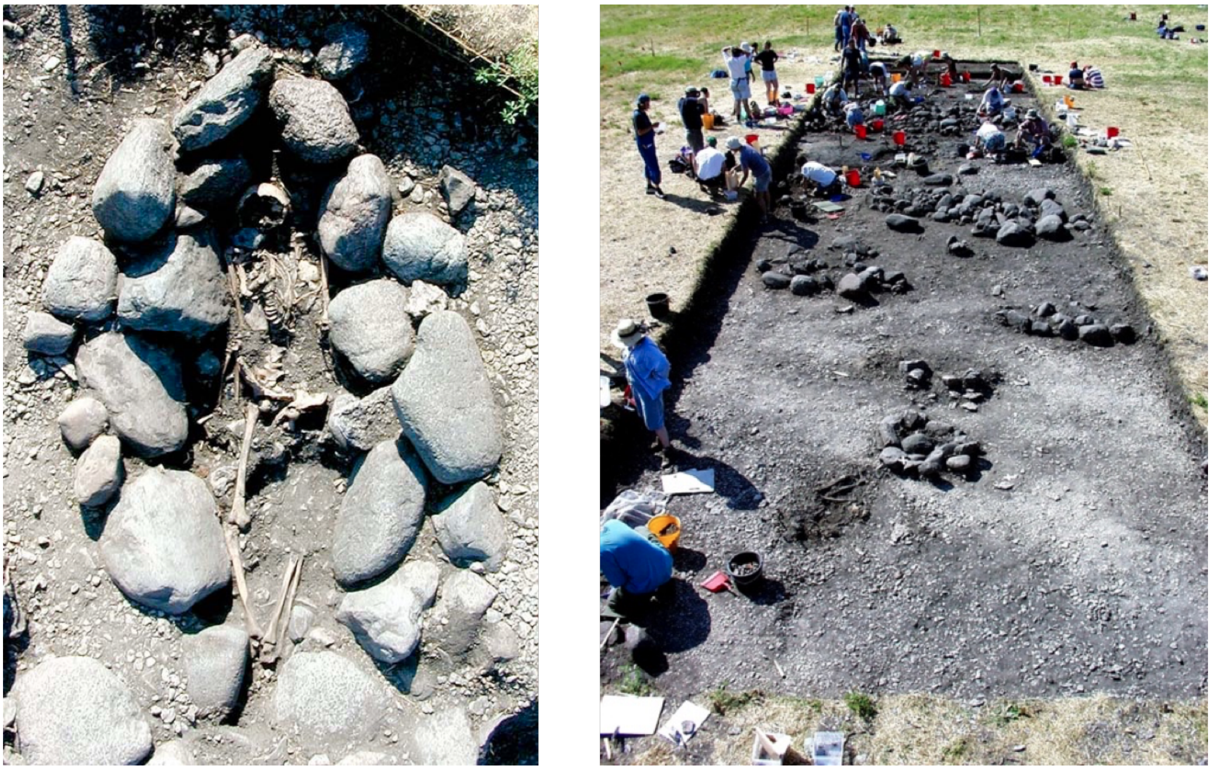

**Fig. S1.7**: Grave 8 (left) and the overview (right) of the Fröjel site it was excavated in. © Dan Carlsson

The excavations of the Fröjel site was initiated in the late 1980's and proceeded for many years under the management of associate professor Dan Carlsson, Centre for Baltic Sea studies, University of Gotland. The excavations revealed the remains of both a harbor and a place of commerce, with remnants of house structures from the 500's to the 1180's (Fig. S1.7). The archaeological finds included weight scales, balances, and silver coins, mainly Arabic and German, that indicated long distance trade. Other finds included raw materials and semi-manufactured objects of metal, bone, and antler, and imply the presence of handicrafts (combs and needles). Vast finds of animal bones indicate animal husbandry in the area, and finds of fish net sinkers indicate that the population were fishing in the area. A Viking Age grave field was also excavated in the area, and contained finds of jewelry and fine details of garments. It is also evident that a church was built at the site in the early 1000's[91,92]. The archaeological finds and the archaeological skeletal remains are stored at the County Museum Gotland.

Samples used for DNA analysis

VK56 Gotland_Frojel-001A98

VK57 Gotland_Frojel-03601
VK58 Gotland_Frojel-03604
VK60 Gotland_Frojel-00702
VK63 Gotland_Frojel-01499
VK64 Gotland_Frojel-03504
VK428 Gotland_Frojel-00287
VK429 Gotland_Frojel-01599
VK430 Gotland_Frojel-00502
VK431 Gotland_Frojel-00487A
VK432 Gotland_Frojel-00303
VK433 Gotland_Frojel-01798
VK434 Gotland_Frojel-01288
VK435 Gotland_Frojel-02500
VK437 Gotland_Frojel-02303
VK438 Gotland_Frojel-04498
VK439 Gotland_Frojel-02498
VK440 Gotland_Frojel-003A88
VK441 Gotland_Frojel-032A98
VK455 Gotland_Frojel-03401
VK456 Gotland_Frojel-02404
VK457 Gotland_Frojel-03299_1
VK458 Gotland_Frojel-03299_3
VK459 Gotland_Frojel-02198
VK460 Gotland_Frojel-04898
VK461 Gotland_Frojel-025A89
VK462 Gotland_Frojel-025B89
VK463 Gotland_Frojel-019A89
VK464 Gotland_Frojel-019B89

**Acknowledgements**

Thanks to Gotlands Museum for lending the Kopparsvik and Fröjel material.
Dentist Carolina Bertilsson, Gothenurg and Professor Peter Lingström, Department of Cardiology,

the Institute of Odontology, the Sahlgrenska Academy at University of Gothenburg. Dentist Björn Lundberg, Tandläkarpraktiken, Adelsgatan, Visby and Johanna Andersson, Gothenburg. Professor Kerstin Lidén, Department of Archaeology and Classical Studies, Stockholm University.

### Vikings in Medieval Russia

**Natalia Grigoreva (Institute for the history of material culture, Russian Academy of Sciences, Saint-Petersburg, Russia) and Vyacheslav Moiseyev (Peter the Great Museum of Anthropology and Ethnography (Kunstkamera), Russian Academy of Sciences)**

Scandinavian artifacts are frequently found in the settlements and burials between 9 and 11 centuries CE on the Northern territories occupied by early Russian state known as Old Rus'. These finds and historical literal sources namely the earliest Russian chronicle of Tale of Past Years according to which Vikings (Varyagi in Russian tradition) took part in the foundation of the Rus' state in 862 put forward a question about the role of Scandinavians in history and population processes in early Russia.

Regrettably pre-Christian burial tradition in Baltic area including Scandinavia was based on cremation which limits the wide use of skeletal remains in morphologic and genetic population studies. Most anthropological series referable to the problem are dated by period after the Christianization of the Rus' at the end of 10th century.

Among the sites series which can be closely related to Vikings are those from Shestovitsy, Chernigov region, Ukraine, Gnezdovo, Smolensk region, Russia, Staraya Ladoga (Leningrad region, Russia) and some others.

**Staraya Ladoga (before 1701 - Ladoga)**

Coordinates 59.999102, 32.295809

According to dendrochronological data Ladoga (in the saga «Aldeigja» or «Aldeigjuborg») was founded in 753. The place became the first residence of military leader Rurik possibly of Scandinavian origin in 862 before his moving to Novgorod and who found first Russian royal dynasty of Rurikovichi. In 11-13th centuries Staraya Ladoga functioned as trade outpost of the Novgorod Republic on the way from the Baltic Sea to Novgorod and then to Constantinople or to Caspian Sea.

It is generally believed that Medieval Ladoga being an important trade and handicraft center had complex population structure which included groups of different origin from all Baltic regions. The strongest Scandinavian influence in Staraya Ladoga is detected for the period from the middle of 9th to the middle of the 10th century. Apart of a large number of Scandinavian imports presence of two large log buildings with galleries in Ladoga settlement was reported. It was suggested that these buildings served as inns and trade-storages of Scandinavian merchants[93].

Unfortunately, early Scandinavian burials of 9-10 centuries found at Plakun site situated on the other bank of the Volkhov River opposite to Ladoga settlement consist of only cremations.

The only early medieval skeletal collection available for morphologic and genetic studies was excavated in 1938-1939 by V.I. Ravdonikas and G.P. Grozdilov on so called Zeml'yanoe Gorodische (Earth hillfort) of Staraya Ladoga. All burials in the cemetery were made in graves without mounds according to Christian tradition. Skeletons lay on their backs with their heads to the West. According to stratigraphy Ravdonikas suggested two periods of functioning of the cemetery namely 17-18 and 11-12 centuries. Several burials from the lower horizon was dated by $^{14}C$ as 880-1188. According to the analysis for strontium, the buried have differences from the local fauna. According to written sources in first half 11 century Ladoga area was given by Grand Prince of Rus' Yaroslav Mudry (Yaroslav the Wise) to his wife Swedish Princess Ingegerd Olofsdotter as a marriage gift, who in turn set her relative Earl Ragnvald Ulfsson as governor of the land. Scandinavian Kongesagaer (kings' sagas) testifies presence of Viking Christians in the military troops of Ragnvald Ulfsson for defense from local pagan tribes.

Collection of skeletons from earlier graves of Staraya Ladoga housed in the Peter the Great Museum of Anthropology and Ethnography (Kunstkamera) consists of 65 individuals. Craniometrical studies reveal that skulls of individuals from the southern part of the cemetery morphologically are closely related to series of Viking Age from Scandinavia[94] while those of Northern part possibly belong to admixed Slavic, Finnish and Scandinavian population. Later this finding was supported by results of integrative analysis of cranial metric and nonmetric traits[95].

Samples used for DNA analysis

VK14 Russia_Ladoga_5680-12

VK15 Russia_Ladoga_5680-16

VK16 Russia_Ladoga_5680-2

VK17 Russia_Ladoga_5680-17

VK18 Russia_Ladoga_5680-3

VK19 Russia_Ladoga_5757-1

VK20 Russia_Ladoga_5680-1
VK21 Russia_Ladoga_5680-18
VK22 Russia_Ladoga_5680-13
VK23 Russia_Ladoga_5680-9
VK218 Russia_Ladoga_5680-4
VK219 Russia_Ladoga_5680-10
VK220 Russia_Ladoga_5680-11
VK221 Russia_Ladoga_5757-14
VK408 Russia_Ladoga_5757-18
VK409 Russia_Ladoga_5680-14
VK410 Russia_Ladoga_5680-15

**Kurevanikha 2**

Coordinates: 58.783533, 36.162257

The Kurevanikha 2 medieval cemetery is located on Mologa River near Kurevanikha village, Ust'uzhna district, Vologda region. The cemetery was excavated by A.N. Bashenkin in 1990. The age of the burial place was estimated as 11-13 centuries. It was suggested that first Scandinavians appeared at the place in 8$^{th}$ century due to its important role on the way from Ladoga region to Volga trade way to Caspian province. Again, although this suggestion is supported by Scandinavian artifacts found in graves it cannot be proved by morphologic or genetic researches because of wide spread cremation tradition of before Christian period. Nevertheless, cranial metric studies reveal that Christian time series of skulls from Kurenvanikha 2 has close biological affinities with Scandinavian series of the Viking Age[94].

Samples used for DNA analysis
VK160 Russia_Kurevanikka_7283-3
VK161 Russia_Kurevanikka_7283-4

**Pskov**

Together with Gnezdovo and Staraya Ladoga Medieval Pskov was area with clear evidence of Scandinavian presence. The main part of Pskov Necropolis medieval time situated on the territory of

Okol’ny Gorod (Outskirts City) is occupied by kurgan burial grounds. While most of burials are very poor of artifacts few of them have rather rich kit of the Scandinavian things. Most of burials consist of cremations but also several burials with inhumations were found.

In 2003-2009 eight chamber burials with inhumations of 10-11 centuries were found near Starovoznesenskiy monastery (Ancient Pskov necropolis of 10$^{th}$ - beginning of 11$^{th}$ century). Originally these burials were covered by artificial mounds which were later destroyed during subsequent city construction. Nevertheless, these graves survived and were found during rescue archeological excavations. The group of burial chambers were also found in Kiev, Timerevo, Gnezdovo, Staraya Ladoga and some other. Basically, cameras are bigger than ordinary ground burial pits with their size up to 3.4 X 3.2 m. Their walls, ceiling and floor were made of logs. Basically, these burials were accompanied by numerous artifacts of Northern European and of Scandinavian origin.

It was suggested that chambers burial tradition in early Russia are of Scandinavian origin. The socio-cultural group which practiced this rite in Eastern Europe became part of the higher stratum of the Old-Russian population. With the adoption of Christianity that tradition subsided[96].

Samples used for DNA analysis

VK158 Russia_Pskov_7283-18

VK159 Russia_Pskov_7283-20

#### Gnezdovo, Russia

**Tamara Pushkina and Alexandra Buzhilova, Moscow State University**

The Gnezdovo complex of archeological monuments (main period of its existence covers the early $10^{th}$ - the early $11^{th}$ centuries) is situated about 13 km westward of present-day Smolensk. The complex occupies a vast territory on both banks of the Dnieper river and consists of two settlements and seven groups of burial mounds that surrounds settlements[97]. The complex includes 13 coin and coin-and-items hoards of the $10^{th}$ century. The hoards were found mainly in the territory of the Central settlement, the principal and the most explored settlement[98].

At the present time Gnezdovo settlement that belongs to the $10^{th}$-early $11^{th}$ century is designated as the only early town center in the Smolensk part of the Dnieper basin and the biggest monuments of the Old Russian state emergence in the territory of Eastern Europe. Many characteristics obtained in the course of archeological study results analysis brings Gnezdovo together with town centers that existed in Northern Europe in the late first-early second millennia AD, such as Birka, Hedeby etc.

The monument’s scope and its role in many respects were determined by Gnezdovo’s geographical position at the junction of Eastern Europe’s river systems that connect the Baltic with the Black Sea and the Caspian Sea.

Choice of the place for settlement made by Slavs and Scandinavians who came there practically simultaneously was determined by the landscape situation and favorable climatic conditions at the turn of the first and the second millennia: at that relatively dry-weather period the Dnieper flood plain was not drowned during high waters and was suitable for reclamation. Initially the settlement territory was inconsiderable and amounted to no more than 1-2 ha but as early as the first stage of the settlement existence the Central settlement, i.e. the fortified part of the settlement was formed. A small part of burials (about 1% of the collection) belongs to the initial stage of the settlement existence. This number of burials completely corresponds to the monument’s initially modest scope.

Heyday of Gnezdovo fell on the second half of the $10^{th}$ century: the settlement area increased and attained its maximum (about 30 ha) due to expansion of building-up within the flood plain limits and development of new considerable area along the right bank of the Dnieper above the flood plain and of inward lands.

The absolute majority of approximately 1000 of burial mounds explored in all 5 groups of mounds was built at the middle and the second part of the $10^{th}$ century. Rare burial mounds with cremations (such mounds were built near the eastern, north-western, and western boundaries of the settlement) are related to the earliest period of the settlement’s existence. Some mounds of the Forest mounds group and partially the Central mound group belong to such burials. The well-known mound L-13 is one of these mounds. L-13 mound gave the earliest for Gnezdovo find of Byzantine amphora with the Slavic inscription-graffito. In the middle and the second half of the $10^{th}$ century the other mound groups (the Dnieper, the Ol'shanskaya and the Right-Bank Ol'shanskaya groups) were formed. These late mounds are situated along the Dnieper bank downstream. At the same time the Forest and the Central mound groups continued to grow.

Single and paired burials performed in accordance with cremation rite under low mounds and so called big mounds with rather complicated cremation rite as well as single inhumations in graves and chambers under mounds belong to the late period of the settlement existence. Male burials in chambers accompanied with riding horses belong to the same period. By the end of the late period the total number of burial mounds attained 4500-5000. Cremation is predominant burial rite in Gnezdovo mounds (cremations comprise 52% of burials), inhumations compose about one third (31%) of burials in the collection of the explored burials. The rest 17% are so called empty mounds where remnants of burials have not been detected.

As judged by materials obtained in the mound necropolis, population of Gnezdovo attained approximately 800-1100 persons and this population distinguished itself by a stable demographic structure and obvious social differentiation.

Exploration of the settlement territory has demonstrated its variegated structure and availability of different zones: residential and manufacturing zones as well as areas related with Gnezdovo functioning as a river port. Traces of manufacturing activities represented by findings of tools, rejected products, intermediate products and raw material stocks have been noted on all explored plots, from the earliest period of the settlement existence and onwards.

At the heyday of the settlement no less than 5 manufacturing zones functioned in Gnezdovo. Findings related to processing of iron, nonferrous and precious metals are concentrated in these zones. Workshops produced adornments for Scandinavian and Slavic women, bridle and belt plates made in the Volga Bulgarian tradition and items specific for the local culture of long mounds. Topography of Oriental coin silver findings and Scandinavian antiquities witnesses their concentration in Gnezdovo and its immediate vicinity.

About 450 Oriental, Byzantine, and West European coins (the greater part of these coins is fragmented) have been found in burials and occupation layer. Dates of coin minting vary from the $6^{th}$ to the mid-$11^{th}$ centuries. 13 Gnezdovo hoardings contain over 1400 Oriental silver coins (intact and fragmented, with punches and riveted eyelets), two Byzantine nomismae, weigh—scales and plummets as well as various adornments. Adornments include 190 gold and silver items. There are items of Slavic, Scandinavian, and Oriental origin among these adornments.

If judged by concentration of Oriental silver Gnezdovo is comparable with such North European monument as Birka and Old Russian monuments: Kiev, Rurikovo gorodishche (i.e. a place of abandoned old fortified settlement), Novgorod (the earliest layers). So called Byzantine imported goods (glazed pottery, precious textiles etc.) comprise a considerable part of findings. Abundance of Scandinavian items (adornments, items of everyday use, amulets etc.) is one of the most expressive features of the Gnezdovo material culture. These findings are matched with data of burial rite analysis of which permits saying that no less than 25% of burials are Scandinavian ones.

Data of sporo-pollen, carpological, and osteological analysis bear witness that cultivated plants and pastoral farming played an important role in the settlement economy and in some extent provided for vital requirements of the settlement inhabitants. Inconsiderable area of arable land discovered in the western part of Gnezdovo complex beneath mounds of the Dnieper group and character of that time landscape bespeak of inability of the early town population to provide themselves with agricultural produce and had to get a considerable part of such products from without.

The latest burials of Gnezdovo necropolis are few and can be dated by the turn of the 10$^{th}$ and the 11$^{th}$ centuries. Archeological material indicate that a rather active life continued in Gnezdovo in the early 11$^{th}$ century, but Gnezdovo gradually lost its unique character of the urban center and became a feudal estate. The final stage connected with the town extinction and transfer of its functions to Smolensk described in chronicles. This fate brings Gnezdovo together with certain early town centers of North Europe. It is believed that these centers ceased to exist due to a complex of reasons. These reasons include a piecemeal termination of Islamic coins importation in the 70s of the 10$^{th}$ century, change of trade routes and a conflict between local elite and the central power of the emerging state[98].

Samples used for DNA analysis

VK222 Russia_Gnezdovo 60-95

VK223 Russia_Gnezdovo 75-140

VK224 Russia_Gnezdovo 78-249

VK252 Russia_Gnezdovo 78-258

VK253 Russia_Gnezdovo 78-262

VK254 Russia_Gnezdovo 81-287

VK255 Russia_Gnezdovo 81-292
VK272 Russia_Gnezdovo 77-241(g)
VK273 Russia_Gnezdovo 77-255
VK413 Russia_Gnezdovo 81-290
VK466 Russia_Gnezdovo 77-222
VK470 Russia_Gnezdovo 77-212

#### Sandomierz, Poland

**Monika Bajka, "Trzy Epoki" Archaeological Service and Marek Florek, Institute of Archaeology, Maria Curie-Sklodowska University in Lublin, Poland**

The inhumation cemetery is situated on the left bank of the Vistula, in the northern part of the Town Hill of Sandomierz, geo coordinates: N: 50°40’53,56’’; E: 21°44’57,08’’. The Town Hill occupies the edge of the Sandomierz Upland loess area, which towers circa 50 metres above the level of the Vistula. In the Medieval Age (since the end of the 10$^{th}$ century), Sandomierz was the main centre of the Sandomierz Region, and constituted the second centre (after Cracow) of the historical Lesser Poland. In the 12$^{th}$ century, Sandomierz, together with Cracow and Wroclaw, was called *sedes regni principalis*.

As recent hypotheses indicate, Sandomierz was established after the annexation of the northern part of the Lesser Poland (Sandomierz Region) to the State of the Piasts (the family ruling the tribe of Polanie from the GreaterPoland) in around the year 970. The new incomers from the Greater Poland (the centre of the Piast state) had the greatest influence on the creation of the town.

The cemetery encompasses the central and the northern part of the Town Hill. Because the site is in majority a built-up area, the site is known mainly from casual findings. In the years 2013-2015, in the northern part of the Town Hill, 9 graves were found. They represent the oldest part of the cemetery. Because of their shape (large, rectangular grave pits with walls covered with boards and wooden vaults, and with wooden coffins inside) some of them are associated with the “chamber graves” from Scandinavia, north-eastern Europe, northern and central Poland (Pomerania, Kuyavia, Greater Poland). The chamber of grave No. 9 was additionally surrounded with a circular palisade.

Some of the graves (including all the graves of chamber-like construction) were equipped with the following items: clay vessels, an iron axe, dress elements (horseshoe-shaped bronze buckles), silver and bronze decorations (silver rings, temple rings), sets for making fire (flint strike-a-lights, iron firesteels, knives etc)[99–101].

<u>Chronology</u>: Basing on the equipment (pottery, metal findings), the graves discovered in the years 2013-2015 should be dated to the time between the end of the 10th century and the middle of the 11th century. Radiocarbon dating of bones from the grave No. 8 was performed: 1070+/-35 BP, after calibration: 938-1022 cal AD, and from the grave No. 9: 1090+/-35 BP, after calibration: 888-1018 cal AD. This date seems to be the same like the one based on archaeological dating of the artifacts.

Elements indicating the foreign origin of some persons buried in the graves that were excavated in the years 2013-2015:

- the form of some graves: it resembles the form of the chamber graves; it generally does not occur in the Sandomierz Region and in the Lesser Poland
- the vessels found in the graves have no analogies in the local pottery from the 10th and 11th centuries, their forms refer to the pottery from the Greater Poland and central Poland
- horseshoe-shaped bronze buckles (from graves Nos. 7 and 9) which represent a garment element characteristic mainly of the region of the Baltic coast and north-eastern Europe (Rus)
- an axe from grave No. 7, which has the closest analogy to the finds from the cemeteries of the Varangian and Rus' cemeteries from Eastern Europe
- a key from grave No. 9, which was secondarily used as a firesteel; it has the closest analogy to the finds from Scandinavian and east European (Russian) sites.
- the results of the analysis of strontium content in the teeth of the persons buried in the cemetery

The furnishing of the graves is of high standard in comparison to other graves from the same period from Sandomierz and its neighborhood. The form of the graves and their furnishing indicate that they were created for the local elites of foreign origin, who had arrived here from other territories, probably from the Greater Poland, after the annexation of the Sandomierz Region to the State of Polanie (the State of the Piasts). The equipment elements (horseshoe-shaped bronze buckles, axe, key used as firesteels) and the form of the graves might indicate that the Varangians and Rus' people were also buried in the cemetery. They were at the service of the Piasts, e. g. as the members of the princely squads or as merchants.

Samples used for DNA analysis
VK494 Poland_Sandomierz 1/13

#### Cedynia, Poland

**Dariusz Błaszczyk, Institute of Archaeology, University of Warsaw**

Cedynia site 2 (Cedynia commune, Gryfino county, zachodniopomorskie voivodeship) is located on the Odra river (on the right bank) in Western Pomerania region. The cemetery was situated at the culmination and slope of a hill, ca. 200 m to the north-east of the fortified settlement (gród). In the cemetery 1308 inhumation and 9 cremation graves was unearthed. The cemetery was partially excavated. It was used from the end of the 11$^{th}$ c. and/or beginning of the 12$^{th}$ c. to the middle of the 14$^{th}$ c. Grave 558 was different from others graves in the cemetery with its grave equipment, large burial pit and traces of wooden funeral construction (a chamber?).

##### Grave 558

An elite grave – burial in a wooden chamber, male, *adultus*, oriented east-west[102,103]. The individual was facing east (a typical Christian orientation of that time) and buried in a supine position, on the back, arms along the body, legs strait. The deceased was buried in a wooden coffin which was placed in a chamber made of timber 2.8 (length) × 2.3 (width) × 1.2 (depth) m. The chamber was very poorly preserved, only traces of wood were visible. It was possibly built in a framework construction.

<u>Burial equipment</u>: Iron double edged sword (type α according to A. Nadolski[104]) at the left pelvic bone was discovered. The sword had a straight cross-guard, a relatively short handle and a lens-shaped pommel. Its total length was about 97 cm. On the blade under the cross-guard, the x-ray examination revealed a sign in a form of an outline of a face. This type of swords was used in the end of 11$^{th}$ c. and in the beginning of the 12$^{th}$ c. In addition, there were unearthed an iron knife at the left pelvic bone and a U-shaped belt fitting made of bronze plaque on lumbar vertebrae.

<u>Chronology</u>: Archaeological dating based on a typology of the sword: the end of the 11$^{th}$ c. or the first half of the 12$^{th}$ c. $^{14}$C date 1010±30 BP (Poz-75117, 973-1150 AD, 95.4%, a piece of a bone). The date is affected by the preservative covering the whole skeleton (Analyses curried out in Poznań Radiocarbon Laboratory, Poland).

<u>Additional information</u>: The skeleton was covered with rodent bones; they are probably the results of postdepositional processes. Over the burial stones were registered. However, probably they were not connected with the grave no. 558 but were possibly associated with the building of the nearby church[103]. In the cemetery one more burial with sword was found, male, grave no. 1120.

<u>Diet</u>: A sample from a rib produced the following isotope results: $\delta^{13}$C: -19,3, %C: 40,0, $\delta^{15}$N: 10,0, %N: 14,5, C:N: 3,2. It indicates that the diet of the studies individual was based on $C_3$ plants (cereals,

vegetables and fruit) with some amount of animal products (meat, dairy and eggs). He could also have in his diet a small share of freshwater fish. This type of diet was typical for the region and time period (Analyses carried out in Stable Isotope Facility, University of Bradford, UK).

Provenance: A sample from a second premolar P2 produced the following isotope result: $^{87}Sr/^{86}Sr$: 0,7110. This isotopic signature falls within the range of $^{87}Sr/^{86}Sr$ values occurring in the postglacial lowland areas of the southern shore of the Baltic Sea (0,7100-0,7124). It is very likely that the individual was of local origin - from Cedynia itself or from the West Pomerania region (Analyses curried out in Geochronology and Isotope Geochemistry Laboratory University of North Carolina, Chapel Hill, USA).

**Grave 435**

An ordinary grave – burial in a wooden coffin with no grave goods[105,106]. A male, *maturus*, oriented east-west, facing east (a typical Christian orientation of that time) and buried in a supine position, on the back, arms along the body, legs straight. This sample was taken for DNA analysis for comparative purposes.

Chronology: Middle Ages, 11$^{th}$ – 13$^{th}$ centuries

Diet: A sample from a rib produced the following isotope results: $\delta^{13}C$: -20,3, %C: 52,1, $\delta^{15}N$: 11,6, %N: 19,2, C:N: 3,2 (Analyses carried out in Stable Isotope Facility, University of Bradford, UK).

Samples used for DNA analysis

VK211 Poland_Cedynia gr. 435

VK212 Poland_Cedynia gr. 558

#### Czersk, Poland

**Dariusz Błaszczyk, Institute of Archaeology, University of Warsaw**

Czersk site 1 is situated on the Vistula river (on the left bank) in Masovia region (Góra Kalwaria commune, Piaseczno county, Masovia voivodeship). In Middle Ages Czersk was an important center of political and ecclesiastical power. A burial ground was located around a church. The cemetery was partially excavated yielding 797 burials. It was used from the end of the $11^{th}$ century or beginning of the $12^{th}$ century to the $13^{th}$ century[107].

**Grave 609**

An elite grave – male, *senilis* (about 60 years), oriented east-west, facing east (a typical Christian orientation of that time). The height of the individual was about 180 cm (according to M. Trotter and G. Gleser). Visible traces of degenerative changes (connected with the age) and wounds (broken and fused ribs) were observed on the skeleton. The deceased was buried in a wooden coffin with iron ferrules. He was buried in a supine position, on the back, right hand straight along the body, left arm bent in elbow and supported by hand on the pelvis, legs straight.

Burial equipment: Iron double edged sword (type α according to A. Nadolski[104]) at the left pelvic bone was unearthed. In addition, the following items were discovered: 2 bronze bowls (type VI according to T. Poklewski[108]), an iron spearhead bone (type V according to A. Nadolski[104]), a wooden bucket covered with decorated iron sheet, a golden ring, a thin, silver, spirally wound wire (alloy of silver with copper and gold).

Chronology: Archaeological dating based on a typology of the sword, the spearhead and bronze bowls: the end of the $11^{th}$ c. or the first half of the $12^{th}$ c. (most probably the beginning of the $12^{th}$ c.). $^{14}C$ date 1085±30 BP (Poz-68711, 894-1016 AD, 95.4%, a piece of a bone). The date is affected by the reservoir effect as a result of significant consumption of marine fish (proofed by N and C stable isotope analysis). After correction of $^{14}C$ it can be calibrated to 1050-1200 AD, 68% and dated probably to the turn of the $11^{th}$ - $12^{th}$ c (Analyses curried out in Poznań Radiocarbon Laboratory, Poland).

Additional information: The grave was situated near the church, according to some scholars grave 609 is interpreted as a burial of Magnus Haroldson, one of 3 sons of Harold II Godwinson, the last Anglosaxon king of England[109,110].

Diet: A sample from a rib produced the following isotope results: $\delta^{13}C$: -17,2, %C: 43,6, $\delta^{15}N$: 12,3, %N: 15,9, C:N: 3,2. It indicates that an important part of the diet of this individual was sea fish or anadromous fish (migrating, e.g. sturgeon). This type of diet was unusual for the region and time period and is very different from any known studied Polish populations (Analyses curried out in Stable Isotope Facility, University of Bradford, UK).

Provenance**:** Samples from a first molar M1 and a piece of rib produced the following isotope results: $^{87}Sr/^{86}Sr$: 0,7106 and 0,7114. Obtained results are ambiguous and not allow to determine whether the examined person was of local origin or was a newcomer. The $^{87}Sr/^{86}Sr$ value obtained for the first molar (complete mineralization to about 4.5 years of age) is found in both areas with post-glacial deposits as well as areas with marine carbonate rocks or clastic rocks such as loess. Such values occur in northern Poland and other parts of Europe (e.g. southern Scandinavia) but are rather not present in the territory of England. The difference in strontium values between the first molar (forming in early childhood) and a rib (reflecting the last two to four years of life) may indicate that the individual spent his adulthood (including the last years of his life) in a different place than his childhood. This would indicate that the man buried in the grave 609 changed his place of residence during his lifetime (Analyses curried out in Geochronology and Isotope Geochemistry Laboratory University of North Carolina, Chapel Hill, USA).

Samples used for DNA analysis
VK200 Poland_Czersk gr. 609

#### Kraków-Zakrzówek, Poland
#### Dariusz Błaszczyk, Institute of Archaeology, University of Warsaw

The cemetery was situated on the right bank of the Wisła river on the sandy dune in the foothills of the Twardowski Rocks. Cemetery at Kraków-Zakrzówek was a typical early Christian, not churchyard, inhumation, ‘flat’ burial ground with graves arranged in rows and oriented according to east-west axes. 75 certain and 22 probable burials as well as 2 clusters of bones was unearthed[111]. The cemetery was used from the beginning of the 11th to the beginning of the 13th century. One grave attracted special attention because of its extraordinary form. The grave no. 19 was situated in the center of the cemetery and from three sides surrounded by a ditch with traces of 9 postholes. Possibly, a ditch existed also on the fourth side but was destroyed by post-depositional processes. The length of the entire structure was about 4 m, and a width about 3.2 m. These traces indicate the existence on the surface of the grave a monumental timber structure in a form of ‘a house of the dead’ or a fence.

Grave 19 was different from others graves in the cemetery with its central location, surrounding ditch and grave equipment. This sample, however, wasn't sequenced due to poor DNA preservation.

**Grave 24**

A commoner's grave – A male *adultus* or *maturus* (about 40 years old) oriented east-west, facing west and buried in a supine position, right hand along the body, left hand bent with a hand on the pelvis, legs straight. The body height was 168 cm. No traces of a coffin were discovered.
Burial equipment: Whetstone.
Chronology: Middle Ages between the 11$^{th}$ and the beginning of the 13$^{th}$ c.
Provenance: A sample from a second premolar P2 produced the following isotope result: $^{87}Sr/^{86}Sr$: 0,7095. This isotopic signature falls within the range of $^{87}Sr/^{86}Sr$ values occurring in the areas with marine carbonate rocks or clastic rocks such as loess. This type of rock is a geological base in Kraków and its surroundings. It seems that the studied individual was of local origin. (Analyses curried out in Geochronology and Isotope Geochemistry Laboratory University of North Carolina, Chapel Hill, the USA).

Samples used for DNA analysis
VK210 Poland_Kraków gr. 24

**Acknowledgements**

Thanks for lending samples to the State Archeological Museum in Warsaw (Łukasz Stanaszek PhD), the Archaeological Museum in Kraków (Michał Zaitz MA) and the Regional Muzeum in Cedynia.

#### Bodzia, Poland

**Wieslaw Bogdanowicz, Museum & Institute of Zoology PAS, Warszawa, Poland**

Bodzia, located in the north-central part of Poland, approximately 40 km south-east of the city of Toruń, is one of the most fascinating archaeological discoveries dating back to the time of origin of the Polish state. The main discovery connected to this site relates to the remains of an early medieval cemetery found in 2007 during rescue excavations carried out by the Institute of Archeology and Ethnology of the Polish Academy of Sciences along the route of the A1 motorway. The discovery encompassed 50 chamber-like graves dated to the late 10$^{th}$ and early 11$^{th}$ centuries AD; along with 8 belonging to a 2$^{nd}$ phase (dating back to between the second half of the 11th century and the early 12$^{th}$ century). The unique character of this burial ground mainly lies in the fact that members of a small elite population have been buried there. Furthermore, almost all the dead are orientated N-S, in what is an uncommon feature for that time in Mediaeval Europe.

The layout of the cemetery has no analogies in Europe. It is formed of rows of graves with large burial pits placed in quadrangular burial spaces. The burial field is divided into rectangular sepulchral spaces, marked on the surface and arranged into 4 rows oriented along the east-west axis. Some of these plots are adjacent, especially those in the northern row with the shape of a trapezium narrowing down to the east. The others, located more to the south, are arranged in smaller clusters or individually, retaining the same orientation as the rest. The burials were located compactly and contiguously, in this way ensuring a clear delimitation of the cemetery boundaries.

Equally unique are the rich grave goods, which may be linked mainly with Scandinavia and Kievan Rus, but also with Southern and Western Europe. A characteristic feature of all the burials here is the bountiful presence of a range of items, including weapons (sword, langsax, spearhead, Khazarian-type pickaxe) – in the case of men, and numerous ornaments (rings, pendants, amulets, kaptorgas, necklaces, etc.) – in the case of women. There are abundant coins: 67 items from 58 graves. These relate to the Holy Roman Empire, England, the Premyslid State and Poland. These and many other features make the cemetery at Bodzia a very specific example highlighting Europe's past.

A study based on the strontium isotope 87Sr/86Sr[112] combined with genetic analyses[113] show that a part of the population buried at Bodzia was not local, but was probably of Scandinavian and/or Rus-Varangian origin. A special role is played here by the tomb of a young warrior (E864/I) buried together with three young women; one of them was placed below him, in what is a two-level burial pit. In his tomb, a ceremonial sword was folded, ornamented in the Mammen style. On the strap-end there is a bident – the tamga of Prince Sviatopolk the Accursed (1015-1019) – son of Vladimir the Great and husband of a daughter of Polish king Boleslav I (the Brave).

The cemetery extends back to the period in which the Polish State had its origins, and also relates to unknown episodes in that state formation, and to the emergence of elites in the early state under the Piasts. The aforementioned features (as well as others) ensure this discovery unique status where Europe's past is concerned.

Bone remains of five individuals have been studied (see the inventory of the burials and their contents after Sobkowiak-Tabaka[114]; with dating after Buko and Kara[115]):

E63 (♂) – adultus/early maturus, incomplete skeleton, pathological changes included dental caries, head of a Type 1 axe found near right femur; dated 978-1016 AD;

E864/I (♂) – adultus, skeleton with several injuries made by a sharp-edged tool with lack of traces of healing suggesting a violent death; weapons including an iron Petersen Type Z sword (with swords of this type mainly dated to the first half and middle of the $11^{th}$ century, i.e. the Late Viking Period[116]; dated 1010-1020 AD;

E58 (♂) – maturus, incomplete skeleton, pathological changes including lifetime loss of teeth, with weapons including a battle-knife of the langsax type, and tools an iron knife placed near the left arm, and another iron knife placed near the right femur;

E37 (♀) – early adultus, incomplete skeleton, with probable post-inflammatory changes visible on the left femur and a post-inflammation state also noted inside the right rib; iron bucket hoop near the feet; in a row of graves dated between 980/990-1030/1035;

E870 (♀) – adultus (20-25 years old), incomplete skeleton, no pathological changes noted, among findings are a few coins and a whorl made of Volhynian slate; dated 1017-1023 AD.

Samples used for DNA analysis

VK153 Poland_Bodzia E63

VK154 Poland_Bodzia E37

VK155 Poland_Bodzia E870

VK156 Poland_Bodzia E58

VK157 Poland_Bodzia E864/I

#### Orkney Islands, UK

**Julie Gibson, Ingrid Mainland, University of the Highlands and Islands, Orkney, UK**

**Newark, Deerness, Orkney (HY 5746 0413)**

Newark comprises a Viking Age and Medieval period chapel and cemetery located in the parish of Deerness on the Mainland of Orkney. The site was excavated in the late 1960s and early 1970s and although not fully published the osteological materials recovered have been subject to $^{14}C$ and isotopic analysis, providing a broad chronology for the cemetery and placing the human remains within the cultural traditions of Scandinavian Scotland[117,118].

The site at Newark stands in a sand-blown landscape at the side of a bay. The deposits remain to a maximum of c1.3m thick above glacial clay and had been truncated by the imposition of a grand post-medieval house. The cemetery has been eroding at least since the 1920's exposing over 100m of graveyard associated with a medieval chapel.

The chapel was excavated by Dr Don Brothwell between 1969 and 1972 in order to obtain a sequence of Norse skeletons; Burials continue to be exposed from time to time in the coastal section. The recent find of a Pictish Type 2 decorated cross slab on this site by Hugo Anderson Whymark, introduces the possibility of the presence of an earlier 8th century chapel (see https://sketchfab.com/models/6de93d22334a4b6da3098402e7e720b5).

Three skeletons were selected for analysis, based on preservation and availability. Of these, only one (VK205) has a secure date: 930+- 40, calibrated, but marine reservoir effect has been proposed by Barrett suggesting a date in early 11th century[117]. This individual is a female and was buried without grave goods.

Samples used for DNA analysis
VK204 Orkney_Newark for Brothwell
VK205 Orkney_Newark 68/12
VK206 Orkney_Newark 71(13)

**Buckquoy, Birsay, Orkney (HY 243 282)**

Buckquoy is a Late Iron Age ('Pictish') and Norse period settlement site located in the parish of Birsay on the Mainland of Orkney. The settlement has been a key site in discussion on Scandinavian

first contact[119], due to the admixture of Pictish and Scandinavian type material found within Pictish-style houses. Inhumations were associated with the structural remains, dating to both the Pre-Viking and the Viking occupation. A full account of the archaeology is to be found in Ritchie[120] with additional information on the dating of the skeletal remains in Ashmore[121].
Excavations undertaken in the early 1970's at Point of Buckquoy[120] revealed a somewhat disturbed inhumation inserted into the top of a multi-period settlement mound of the 8th to 10th centuries AD. Immediately beneath were the partial remains of a succession of Norse longhouses, overlying a Pictish multi-cellular dwelling. Identified by skeletal analysis as a male aged about 40, the burial (VK202) consisted of a simple scoop grave with the body thought to have been interred in a crouched position, laid on his right side and aligned S-N with his feet to the north. Grave goods consisted of a bronze ring-pin of the early 10$^{th}$ century, half of a silver penny of Edmund (940-6), an iron knife, a whetstone and an iron javelin-head. $^{14}$C dates are available but earlier than the artefactual assemblage, that accords with its stratigraphic relations. Brundle et al.[122] suggest the dated bone may be intrusive. VK201 was an earlier, Pictish, unaccompanied long cist burial of an adult male dated to 404 - 596 calAD[121].

Samples used for DNA analysis
VK201 Orkney_Buckquoy, sk M12
VK202 Orkney_Buckquoy, sk 7B

**Brough Road, Birsay, Orkney (HY 2466 2807)**
Archaeological remains from Brough Road are derived from excavation during the 1970s of a coastally eroding midden and associated structures dating to the Late Iron Age and Norse periods. A full account of the archaeology is to be found in Morris et al.[123] with additional information on the dating and of isotopic analysis of human remains in Ashmore[121] and Barrett et al.[124]. A pre-Viking long cist cemetery and later Viking or Norse inhumations were also recovered within this area, which overlooks the Brough of Birsay and Birsay village, important Pictish/Viking/Norse settlements and eccesiastical centre.

Three individuals were sampled from Area A Brough Road. The primary use of the site is reflected by cairn-burials, comprising two long cist graves underlying stone cairns. VK203 is a long cist grave unaccompanied inhumation containing only one body. Two dates from this individual have resulted in two widely spaced $^{14}C$ dates, 130-54 CalAD or 548-668 CalAD, but this individual represents in both cases a pre-Viking burial. Osteologically, this individual was probably a male and was elderly. The cairns were sealed by sand and midden layers into which were inserted two burials dating to Viking/Late Norse periods: VK208 - a rough cist grave cutting into the midden topping one of the cairns containing a disarticulated body of a probable male aged over 30 years. It was associated with artefacts iron artefacts, complete antler comb, hog-backed in form and of a Viking period date. Two dates from this individual indicate a later Pictish to Viking date-range, 650-980 CalAD or 890-1026 CalAD. On top of one the cairns (above) lay a flexed inhumation (VK207), not contained within a cist and with no grave finds. This was an adult, aged c. 30-35 years but sex could not be determined oseteologically. A $^{14}C$ date of 880 to 1160 Cal AD[121] was obtained for this individual.

Samples used for DNA analysis
VK203 Orkney_BY78, Ar. 1, sk 3
VK207 Orkney_BY78, Ar. 1, sk 1
VK208 Orkney_BY78, Ar. 1, sk 2

### Medieval skeletons from the Public Library Site in Trondheim, Norway

**Birgitte Skar and Lisa Mariann Strand, NTNU University Museum, Trondheim, Norway**

Seven skeletons originating from the Early and High medieval period in Norway, extending from 1030 to ca 1350 AD are part of the foundation of the genetic research material discussing demographics during the Viking Ages.

The medieval churchyard from which the skeletons originate was located at The Public Library Site (Folkebibliotekstomten) in the central part of the town Trondheim. After several minor excavations in the 19th and 20th century, the excavation of the St. Olavs churchyard located at this site was completed during a ten-month period between 1984-85[1].

The individuals that are included in the present analysis were between 11 and 50 years of age, representing both biological females and males. Each burial context shows distinct expressions of Christian ritualistic burial customs as they were inhumations, buried in an east-west direction and were either buried in plank coffins, trunk coffins or without coffin.

The seven individuals analysed in this study belong to the St. Olavs churchyard[125]. This part of the churchyard lasted for approximately 100 years. During this stage, the churchyard was extended and an increased amount of burials were found. In addition to the clearly Christian burial customs, burial rods and hazel sticks were present in some of the graves.

Five of these have been radiocarbon dated and display surprisingly old dates. Three of them except the sample from individual VK113 (SK223) are dated within the final Viking period, the two mentioned individuals, however, derive from the very beginning of the Medieval period. From the automated measure of stable carbon (**d**13 C) from the dating laboratory, it can be read that the stable carbon value is quite low. This indicates that the individuals have had a mixed diet potentially consisting primarily of marine resources. This is an issue that needs further investigation regarding stable carbon and nitrogen isotope values of the five dated individuals, which may again serve to clarify the obtained dates with reference to marine reservoir effect and thus shed light on the question of the dates being too old.

Samples used for DNA analysis

VK113 Norway_Trondheim_SK223

VK114 Norway_Trondheim_SK332

VK116 Norway_Trondheim_SK372

VK117 Norway_Trondheim_SK328
VK118 Norway_Trondheim_SK271
VK124 Norway_Trondheim_SK356
VK125 Norway_Trondheim_SK367

#### Italian Medieval sites

**Gabriele Scorrano (University of Copenhagen), Enrico Cappellini (University of Copenhagen), Pasquale Favia (University of Foggia), Italo M. Muntoni (Soprintendenza Archeologia)**

**San Lorenzo in Carminiano (Foggia, Italy)**

The medieval settlement of S. Lorenzo in Carminiano (then Carmignano) was the main village in Northern Apulia between Late Middle Ages and Modern Age. It is located in the central area of the "Tavoliere delle Puglie" plain, just outside the city of Foggia (Northern Apulia – Italy). The settlement topography is articulated into three subdivisions, bounded by ditches, with a northern trapezoidal one (enclosure I), probably surrounded by walls and extending over 7 hectares ca. (13$^{th}$ - 16$^{th}$ century), another in the north-western position (enclosure II), smaller in size and with a half-circular morphology, and a third southern one (enclosure III), elliptical and very broad (up to 15 ha). In the site only a small church dedicated to San Lorenzo has been found. Outside the church, along the bottom wall, various paving slabs have been discovered, both in cobwebs (USR 932-841) covered with a combustion ground, in tessellato, with stone tiles and brick sections and finally a wider lacer with brick remains[126]. Some poles' holes have also been found and they may perhaps refer to a late-medieval stage. The trench carried out in the area in front of the church identifies at least four phases: the first three referring to the Middle Ages, the latter probably to the 17$^{th}$-18$^{th}$ century[126].

The oldest traces of a funeral attendance are represented by a simple burial in the ground (t.2), hosting two individuals, and a third one.

The next occupation was set up directly on the cemeteries, obliterating the ditches, addressing a residential area of which two orthogonal sects have been identified so far in cobblestones tied to the ground. The bigger enclosure is the suburbium of settlement. In this area, some silos for cereal have been found, directly cut into the natural substrates of limestone clay and clay, without coating.

Samples used for DNA analysis

VK534 Italy_Foggia-869

VK535 Italy_Foggia-891

**Cancarro (Troia, Foggia, Italy)**

The church of Cancarro is situated at 3,5 km South-West from Troia (Foggia, Italy), at 430 m asl. The city of Troia was built on the ruins of the Roman town of Aecae in a strategic position on the Via

Traiana. The church was used between the 11$^{th}$ and the 13$^{th}$ century[127]. Next to the church archaeologists unearthed a cemetery with 54 well-preserved burials, often overlaying each other. Two graves host two individuals and five pits are ossuaries[127]. A minimum of 79 skeletons, mainly women, were confidently identified from re-used graves[127]. The cemetery was used between the late 11$^{th}$ century (Norman age) to the second half of the 13$^{th}$ century (Swabian-Angevin age), based on artifacts[127].

Samples used for DNA analysis

VK536 Italy_Foggia-1240

VK537 Italy_Foggia-1248

VK538 Italy_Foggia-1249

#### Balladoole, Isle of Man, UK
#### Allison Fox, Manx National Heritage

The Site: Around AD 900, a Viking was buried in an oak ship at Balladoole, Arbory in the south east of the Isle of Man. The site is a low hill, with a wide panorama over the surrounding landscape and seascape. There is artefactual evidence for Mesolithic and Neolithic use of the area, and structural evidence for Bronze Age burials and Iron Age occupation. It was during the excavation of the latter hillfort that the Viking burial was discovered[128].

The ship had been placed in a shallow pit, which disturbed several existing Christian burials. Boulders were placed around the hull of the ship to hold it in place, and the dead man and his goods placed within (Extended Data Fig. 1c). The grave goods included jewelry of outstanding workmanship, as well as simpler, practical objects, but no sword, indicating that his status was gained through trade. A low mound of earth and boulders was then raised over the ship and capped with the cremated remains of a horse, dog, pig, sheep or goat, ox and cat that had been sacrificed as part of the burial rite. A marker post was set in the top of the mound.

By the time the site was excavated in the 1940s, the timber of the ship had almost completely rotted away in the soil, leaving only 300 of the iron nails that once held it together. The ship was about 10.5 m in length and was a trading vessel built for sailing short distances and landing on beaches to buy and sell goods, ideal for trading around the Irish Sea.

The Viking: From the skeletal remains, the Viking was about 45 years old when he died, c. 175 cm tall, strong and muscular. His teeth were badly worn by chewing coarse food. During 2006/7, the face of the Viking was reconstructed by Dr Caroline Wilkinson and Caroline Needham, University of Dundee. Scientific analysis on his diet and origin was undertaken during the same period by a team led by Dr Leigh Symonds. Isotope analysis suggested that the Viking was quite likely to have come directly from Scandinavia rather than from around the site of his burial[129].

Samples used for DNA analysis

VK170 Isle-of-Man_Balladoole

#### Viking Age sites from Ireland

**Maeve Sikora (National Museum of Ireland) and Linzi Simpson (Trinity College Dublin)**

**Islandbridge, Dublin NMI 08e0693:001 (Ireland_08e693)**

This is the most recent discovery of a furnished Viking grave in Ireland. The grave, at Islandbridge in Dublin, is located close to the find spot of many Viking graves reported in the 19th century[130]. The human remains were discovered during an excavation in 2008 that took place as a response to the discovery of an iron sword and spearhead at the same location in 2004[131]. The in-situ human remains were discovered at a depth of 80-90cm below ground level but most of the skeleton had been heavily disturbed and did not survive. The individual was lying in a supine position, and based on the position of the in-situ vertebrae, was oriented approximately north-south with the head at the north. The legs appear to have been flexed, as indicated by the portion of the left femur which was in situ. Portions of the right scapula and clavicle, fragments of 7 right ribs, unsided fragments of radius and ulna, and fragments of vertebrae remains were recovered and have been identified by Barra Ó Donnabhain as those of a young adult male aged between 18 and 20 years at death[131]. Although the grave had been heavily disturbed, a copper alloy ringed pin and two copper alloy objects which probably represent a scale pan, were discovered associated with the remains. Copper staining on some of the rib bones suggests that these artefacts were originally placed on the torso. It is considered probable that the sword and spearhead found in the same area in 2004 were associated with this grave. Isotope analysis of the human remains uncovered indicate that this individual was non-local to Dublin and spent most of his early life in some region of Scandinavia before coming to Ireland[131]. The burial has not been radiocarbon dated but based on the artefact typology dates to the first half of the ninth century.

The main body of comparable material for this grave is found in the Islandbridge/Kilmainham/Inchicore area of Dublin. Graves from this area, the largest cemetery in the Viking west, have produced a rich array of artefacts with an impressive collection of weapons and trading material, among other objects. This grave was immediately north-west of the 'great gravel pit' that produced the rich collection of grave-goods in 1866[130].

Samples used for DNA analysis

VK546 Ireland_08E693

**Eyrephort, Co. Galway NMI 1947:55 (Ireland_EP55)**

This grave is one of the few graves of early Viking Age date in Ireland to be discovered outside of Dublin. It was discovered 1947 in west Co. Galway, in sand dunes in the townland of Eyrephort, close to the coast. It was not excavated at the time but a description of the burial was recorded based on the account of the finder who had uncovered and subsequently re-interred the remains. A full re-evaluation of the grave and contents has recently been published and the account here is based on this most recent analysis[130]. The human remains were analysed by Barra Ó Donnabhain and represent a young adult male, probably aged between 20 and 25 years at death and with a living stature of c.175cm[130]. The burial was extended and supine with the head to the south-west and the feet to the north-east. The grave contained an iron sword, spearhead and shield boss. Iron staining on the bones indicates that the sword probably lay along the left side, next to the arm. Both the spearhead and shield boss are of Scandinavian type and the sword is a double-edged sword of Petersen type K dating to the second half of the ninth century[130].

Samples used for DNA analysis

VK543 Ireland_EP55

**Finglas, Co. Dublin, NMI 04E0900 (Ireland_FG254)**

This grave was discovered during excavations in advance of construction works at Finglas, a suburb of Dublin, some 5.5km northwest of the city[132]. Finglas was formerly a village outside of Dublin, and probably developed from an early medieval monastic centre founded there. Although relatively little is recorded about the monastery, references to a scriptorium there in the ninth century suggest that it was considered a significant foundation[133]. This report is based on Kavanagh's account of the excavation and Sikora's publication on the finds[132,133]. The grave was located close to the church at Finglas, but outside of the boundary of the church grounds. It contained a single adult female who was buried with a pair of oval brooches, a casket decorated with bone plaques and an antler comb, only a portion of which survived. The skeleton was extended and supine, with the head to the south-west. It had been damaged by previous groundworks and the lower legs and feet had been heavily disturbed. The remains represented a woman aged between 25 and 35 at death. She had been buried wearing a pair of oval brooches, one at each shoulder. The casket and comb had been placed by her right side. Only one brooch survives intact while the second is very fragmentary. The brooches are single-shelled cast oval brooches of the Berdal type which have been gilded and inlaid with silver wire. The complete brooch is highly decorated with zoomorphic studs some of which are cast with the brooch and some of which were applied. These animals have eyes made of glass studs. This type of brooch dates to the mid-ninth century, and it is likely that the burial occurred around this time[133]. Finglas is not only unusual for the location of the burial and the high-status of some of the grave-goods. In Ireland, 78 of the 107 Viking graves known are those of males[130], and the discovery of a female grave is therefore highly significant. The location of the grave, clearly a non-Christian burial, adjacent to a Christian church is also interesting and as Harrison and Ó Floinn point out, is 'further evidence of the complex relationship between the 'pagan' Norse and 'Christian' Irish in the mid-ninth century'[130].

Samples used for DNA analysis
VK544 Ireland_FG254

**Ship Street Great**

**Licence no. 01E0722, (Ireland_SSG12)**

This Viking warrior grave was located on the western side of Ship Street Great, close to the junction with Stephen Street Lower. The removal of a deep cellar exposed the remains of a very truncated skeleton lying approximately 3m below the ground level, in the boulder clay. The partial skeleton was in a supine position with the head to the west and was lying in a shallow grave-cut. All that survived was the intact lower mandible, crushed skull fragments, vertebrae, some ribs, the shoulder blades, the right collar bone and part of the right arm. Analysis by Laureen Buckley, Osteologist, revealed that this was probably a male, aged between 25 and 29 who had suffered nutritional deficiency and had the beginnings of degenerative disease of the spine suggestive of a hard and strenuous lifestyle. Four small artefacts were found in the neck area, a small silver finger ring, a corroded iron object, a glass bead, and small twisted silver ring. A fragment of a pattern-welded sword was also found close to the skeleton but an examination of the surrounding boulder clay found no other features. A fragment of bone was sent for dating and this produced a 95% probability of the individual dating to between AD 665 and AD 865 (intercept date of AD 790) with a 68% probability of dating to between AD 690 and AD 775 (send to Centrum Voor IsotopenOnderzoek, Groningen). The date-range and mode of burial confirms this individual was likely to have been part of the hordes of Viking warriors which descended on Ireland from the late 8$^{th}$ century onwards. That the skeleton formed part of a larger group of burials was also indicated by the subsequent excavations in Golden

Lane by Edmond O'Donovan where four additional Viking skeletons (including a female) were found, one of which lay just 5m west of the Ship Street burial. These were interred in the environs of an Early Christian cemetery and were all of a similar date. Four skeletons with some with weaponry had been found previously 150 m to the north-east, at South Great George's Street, and isotope analysis of these suggest that two were reared in the Scandinavian region while the other two were reared closer to the Atlantic (in and around the British Isles), presumably in Viking colonies on the Western or Northern Scottish isles[130,134–136].

Samples used for DNA analysis
VK545 Ireland_SSG12

**Acknowledgements**
The authors are grateful to Mr John Kavanagh, excavator at Finglas, for providing information on his excavation, and also to Ms Laureen Buckley, Dr Denise Keating and Dr Barra Ó Donnabháin who analysed the remains from the sites mentioned above.

### Iceland


#### Hildur Gestsdóttir, Institute of Archaeology, Iceland


**Hofstaðir, Mývatnssveit**

The farm of Hofstaðir in Mývatnssveit in northern Iceland lies to the west of Lake Mývatn, and is bordered to the west by the river Laxá (Fig. S1.8). A Viking Age hall lies within the home-field of the Hofstaðir farm, west of the current farmhouses and up against a small scarp that demarcates the arable part of the home-field (the home-field boundary lies on top of the scarp). Excavations there, carried out under the direction of Gavin Lucas between 1996 and 2002, indicate that the hall was built in the middle of the $10^{th}$ century. The site of the church and cemetery at Hofstaðir is within the home-field of the modern farm, 80 m southwest of the Viking Age hall, up against the eastern edge of the old farm-mound, which was abandoned in the middle of the $20^{th}$ century. The cemetery sits on the eastern edge of the farm-mound.

Excavations of the site were carried out 2000-2004 and again 2010-2015. The excavations revealed at least two, possibly three phases of church structures in the center of an octagonal area bordered by a turf wall. Little remains of the earliest church, which appears to have been deliberately demolished, except post-foundations and remains of a trampled floor. Radiocarbon and tephrochronolgy dating indicate that this earliest church was established in the mid 10th century. The later church was built on the same spot as the earlier one, although the later church was slightly smaller. Tephrochronology indicates that this later structure was built before the 1300 eruption in Katla, which also seals all burials within the cemetery. The earliest graves clearly respect the oldest structure on the site, and recent radiocarbon dating of six of the skeletons from the site dated from 695-1148, although this range can be tightened in some instances as many of the burials clearly post-date the tephra from the 940 eruption in Veiðivötn.

A total of 170 skeletons were excavated at Hofstaðir, mostly from in situ burials, although there are four examples of redeposited graves. There is clear organization of the cemetery, as seen in other medieval cemeteries in Iceland. Females are mostly buried in the northern half, males mostly in the southern half and children up against the church; in particular, up against its southern wall. The burials are all inhumations. The grave-cuts are very tightly spaced with a lot of intercutting, especially in the area where the children are buried. All the burials are supine with the hands usually resting on the abdomen or alongside the body. About half of the adult burials were in simple wood-coffins of which nothing survived except wood-staining of the soil. The surviving depth of the burials ranged between 30 cm (in areas where there had been levelling of the land for agricultural purposes in the middle of

the 20th century) to about 80 cm (which represents the maximum depth of the burials while the cemetery was in use. Preservation in the cemetery was on average good and in most instances, quite consistent although there were a couple of locations where variations had clearly caused the creation of micro-environments within the cemetery. An example of this is the small porch which had been added to the later church on top of three graves located immediately west of it. This resulted in poorer preservation of the skeletal material in these graves than in the rest of the cemetery.
Bioarchaeological analysis of the skeletons from Hofstaðir have indicated a very high prevalence of inherited osteoarthritis, indicating that the people buried within the cemetery were closely biologically related[137,138].

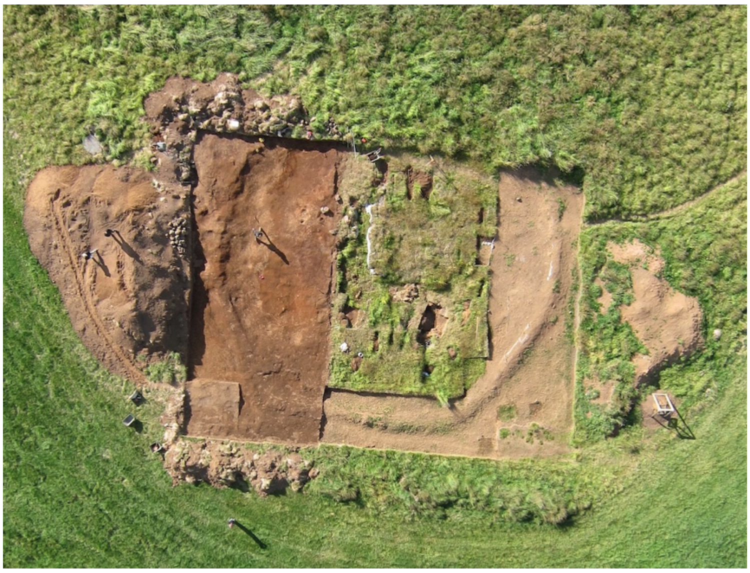

**Fig. S1.8**: Aerial view of the Hofstaðir archaeological site.

Samples used for DNA analysis
VK95 Iceland_HSM-A-127
VK98 Iceland_HSM-A-083
VK99 Iceland_HSM-A-104
VK101 Iceland_HSM-A-125
VK102 Iceland_HSM-A-128
VK110 Iceland_HSM-A-115S
VK111 Iceland_HSM-A-118
VK122 Iceland_HSM-A-114
VK123 Iceland_HSM-X-104

VK225 Iceland_HSM-A-108
VK226 Iceland_HSM-A-115N
VK227 Iceland_HSM-A-117
VK228 Iceland_HSM-A-120
VK230 Iceland_HSM-A-123

**Ingiríðarstaðir, Þegjandadalur**

Þegjandadalur is a largely abandoned valley in northern Iceland, with limited areas of pasture towards the mouth of the valley, to the north. Elevation above sea level rises from circa 40 m at the valley mouth to circa 180 m at the head of the valley. The valley runs broadly north – south for approximately 7 km, and is up to 1.6 km at its widest. The area of lowland is approximately 7.3 $km^2$. The valley walls rise steeply to eroded heathland in the east, at circa 400 m, and more gently to better vegetated heathland in the west at c. 230 m. At the southeastern limit of the valley a saddle gives access to Laxárdalur, at about 250 m above sea level. The grave field at Ingiríðarstaðir was discovered in 2008, as a result of the wider study of a complex and remarkably well preserved archaeological landscape in the valley of Þegjandadalur, in the county of Þingeyjarsysla in northeastern Iceland.

Excavation of the grave field was carried out between 2008-2015. The excavation revealed eight adult inhumation burials as well as the remains of seven horses in six burials, all of which had been disturbed in antiquity. In addition, a single undisturbed neonatal skeleton inhumation burial was recovered.

Radiocarbon and tephrochronological dating indicate a date of late $9^{th}$-early $11^{th}$ century AD.

Recent isotope analysis has demonstrated that the individuals buried at Ingiríðarstaðir were born in the region[139].

Samples used for DNA analysis
VK129 Iceland_ING08

**Hringsdalur, Arnarfirði**

In the summer of 2006 human remains and artefacts were discovered in an eroded sandbank on a small peninsula, Hreggnasi, which lies in the sea by the Hringsdalur valley in Arnarfjörður in the Westfjords in Iceland. This resulted an archaeological investigation of the area between 2006-2011.

The excavation revealed 4-5 burials, including one boat burial, which contained the disturbed remains of four adults, as well as a single undisturbed inhumation burial. A further 2-3 disturbed burials were located at the site.

Dating of the site has mainly been based on artefact typology, placing it in the 10$^{th}$ century AD[140].

Samples used for DNA analysis

VK127 Iceland_HDR08

VK128 Iceland_HDR111

#### Greenland

#### Jette Arneborg, The National Museum of Denmark

**Sandnes, Kilaarsarfi V51**

The Sandnes farm is lying close to the shore at the head of Ameralla fjord and has an associated church. This site is considered as the largest in the southern part of the Norse Western Settlement. The open valley behind the farm offered plenty of pasture while the nearby river supplied water for the farm. Today, the church is sanded up and flooded at high tide, clearly showing the dramatic changes of the landscape that has been happening since the first Norse settlers arrived. Ten ruins have been recorded on the Kilaarsarfik plain during the course of multiple expeditions. An unknown number of ruins have been lost to the fjord[141]. Church and churchyard were excavated in 1930 and 1932.

The discovered artefacts and radiocarbon dates indicate that the Sandnes farm was occupied from c. 1000 AD up until the time of the depopulation of the Western Settlement in the later part of the 14$^{th}$ century. Sr isotopic analysis of skeletons from the church yard from the later settlement period indicate that they were brought up in Greenland[142].

Samples used for DNA analysis

VK74 Greenland_V051_0928

VK75 Greenland_V051_KAL-0929 / skeleton 11

VK76 Greenland_V051_KAL-0957 / skeleton 27

VK77 Greenland_V051_KAL-0959 /skeleton 31

VK78 Greenland_V051_KAL-0960 / skeleton 30

VK196 Greenland_V051_KAL-0947 /skeleton 22

**Ujarassuit (Anavik) V7**

The Anavik farm is situated on a raised beach terrace at Ujarassuit fjord. The houses and the church lie spread over the large flat plateau where conditions for farming would have been excellent during the Viking Age. Nine house ruins have been recorded at the site, including one of the most well-preserved stone houses in the Western Settlement.

The main excavations of the site were carried out in 1932, when multiple buildings were identified. New excavations were conducted in 1982 when a small trench was opened in the churchyard to obtain human remains for δ13 C analysis. Radiocarbon dates and the positioning of the arms of the dead date the excavated burials to the $13^{th}$–$14^{th}$ century. The analysis of δ13 C indicated, that the diet of the analyzed individuals was mainly marine[143,144].

Samples used for DNA analysis
VK189 Greenland_V007_KAL-0992 / skeleton U04

**Narsarsuaq Ø149**

The Ø149 site is a Norse high-status farm with a connected church at Narsarsuaq in Uunartoq Fjord and was first recorded in 1921 by Poul Nørlund and identified as the remains of a Benedictine nunnery. The farm is located on a peninsula between Lichtenau Fjord and Uunartoq Fjord in the southern region of the Eastern Settlement close to the hot springs on the island of Uunartoq. 21 features, including the church, living houses, byres, stables and barns have been recorded on the site. The first archaeological excavations were conducted in 1945-46 and again in 1948 lead by C.L. Vebæk. Few Norse ruins were investigated such as the church and the churchyard, sections of the dwelling, a small stable and the stable/barn complex. The church belongs to the later phase of the settlement from about 1300 but has had one or more predecessors. The human remains from the grave yard were radiocarbon dated to c. $14^{th}$ century[143,145]. Sr isotopic analysis of skeletons from the church yard indicate that the buried were local Greenlanders[142].

Samples used for DNA analysis
VK190 Greenland_Ø149_KAL-0996 / skeleton 3, grave unit I
VK191 Greenland_Ø149_KAL-1000 /skeleton 7, grave unit I

**Ruin group Ø64**

The farm site of E64 is located in Igaliku Kujalleq, a small side branch of Igaliku fjord in the Norse Eastern Settlement. 12 features have been recorded on the site among which are a small church belonging to the group *landnam* churches that were established from the late $10^{th}$ century-around 1000. The church yard was excavated in 2007-08 led by Jette Arneborg. The excavated skeletons were radiocarbon dated within the period from late $10^{th}$ century to about 1200. Sr isotope analysis indicates that several of the buried were immigrants from Iceland[142].
X530, x532, x677 and x678 were all buried in the same grave.

Samples used for DNA analysis
VK1 Greenland_Ø64_KNK2655x677
VK6 Greenland_Ø64_KNK2655x678
VK9 Greenland_Ø64_KNK2655x530
VK11 Greenland_Ø64_KNK2655x532
VK186 Greenland_Ø64_KNK2655#78
VK187 Greenland_Ø64_KNK2655#72

**Brattahlid, Qassiarsuk, Ø29a**

The site is located on the Qassiarsuk plain in Tunulliarfik Fjord where 60 ruins are recovered and identified as the high-status farm Brattahlid, where one of the first Norse settlers such as Erik the Red settled with his family in the mid-AD 980s.

The northernmost farm at Qassiarsuk identified as the ruin group Ø29a, is thought to have been that of Erik the Red. According to Icelandic sagas Tjodhilde, the wife of Erik the Red had a church built on the farm around 1000, and the remains of a small church found in the beginning of the 1960s have been identified with Tjodhildes church mentioned in the sagas. Later radiocarbon dates of skeletons from the church yard indicate, that the small church was built at the time of settlement in the late 10$^{th}$ century. The church yard was taken out of use in the beginning of the 1200s and the church site relocated[143].

There was a sex bias in the arrangement of the graves in the churchyard. The southern side of the church was mostly for high status men while many of the women were buried on the north side. A few high-status women were also buried on the south side, whereas a few low-status men were buried on the north side, and studies show that there were clear differences of both stature and the teeth conditions between people buried on the south and north sides, likely reflecting the social structure of the Norse society[146].

A mass grave was also excavated on the south side of the church with 13 adult men and two boys of 10 and 17 years of age, respectively. This was a secondary grave since the bones of the skeletons did not lie in situ, indicating that the remains might have been moved to “Tjodhildes Church” from another grave or they may have died far away, and their bones were subsequently brought to Brattahlid for burial. Studies by Alexandersen and Prætorius[146] suggest that the individuals buried in the mass grave might have relatives. Sr isotopic analyses indicated that some of the buried at Tjodhildes church were immigrants from Iceland[142].

Samples F2, F3, F5, F6, F7, F8 and F9 are from the mass grave.

Samples used for DNA analysis
VK513 Greenland_Ø029a_KAL-1088 / skeleton F8
VK179 Greenland_Ø029a_KAL-1092 /skeleton F2
VK180 Greenland_Ø029a_KAL-1091 /skeleton F3
VK182 Greenland_Ø029a_KAL-1085 /skeleton F5
VK183 Greenland_Ø029a_KAL-1086 /skeleton F6
VK184 Greenland_Ø029a_KAL-1087 / skeleton F7
VK185 Greenland_Ø029a_KAL-1089 /skeleton F9
VK193 Greenland_Ø029a_KAL-1061 /skeleton 75

### The Church Site In Sandur, Sandoy, Faroe Islands

**Símun V. Arge, Tjóðsavnið - Faroe Islands National Museum**

**Settlement by the bay Sandsvágur**

According to the Færeyingasaga a man, Sniðálvur, lived in the village of Sandur in the Viking period. Local legends have it, that he lived at á Krossi by the church, and that he had fled from the Hebrides because of a charge of manslaughter.

Remains of ancient occupation have regularly been made visible by erosion of the coastline on both sides of the bay of Sandsvágur. Extensive Viking Period settlement have now been revealed by and north of the ancient church site, which have been dated back into the Viking Period. At the site of á Sondum settlement activity has been dated to $4^{th}$-$6^{th}$ century – the earliest dated proofs of human settlement in the islands even this has not been a continued settlement. The village of Sandur is the largest agricultural area in the islands, which may have been especially attractive for the landnámsman when he should choose a site to settle. Settlement historical investigations and place-name studies indicate three larger farms in the village already in the Viking Period[147].

**The church site in Sandur**

Archaeological excavations within the church as well as at the church site at Sandur, við Kirkjugarð, have often revealed remarkable finds. Prior to professional excavations took place the only coin hoard in the islands so far turned up in 1863–98 silver mints deriving from the Continent, Ireland, UK and Scandinavia, hidden by the end of the $11^{th}$ century. Archaeological investigations inside the church 1969-70 revealed five of the present church's predecessors, built in 1839. The first church at the site was a small, plain wooden stave-church apparently dating from the $11^{th}$ century. This church was replaced by a much larger church, Church 2; both churches had a Romanesque plan. The church-typology and the archaeological finds indicate that Church 2 was erected in late $12^{th}$ or early $13^{th}$ century. The archaeological excavation at the church in Sandur is of significant meaning concerning the development of the Faroese parish church.

Excavations in the extended area of the churchyard to the south of the church during the period 1970–2009 extensive settlement remains turned up in the 3000 m² large area. Rather than the proper habitation site this area is interpreted as an activity area with the remains of different activities. Also, a Viking Period burial ground turned up here, where 12 graves were found of which 7 have been investigated. The dead were buried with personal belongings indicating a high-status society in the $10^{th}$ century. The proper settlement area is to be found to the north of the church site where archaeology has verified deep stratified settlement layers along the eroding cliff edge from the period $8^{th}$-early $13^{th}$ century.

**Burials in Church 2**

In the nave of the second church at the site, Church 2 – the first one being a small wooden stave-church apparently dating from the $11^{th}$ century - 23 burials were found, obviously established in a planned manner below the wooden floor of the church, why this might have occurred during a restricted period of time (Extended Data Fig. 8c). Most of the dead were buried in wooden coffins, though a few were interred in the grave in grave clothes only. The coffins were of the trapezoid medieval type.

7 of the buried were adults and 16 were children. Of the seven adults, three were women and four were men, all over the age of 35. Child mortality in the medieval period was high and of the 16 children all but one died before their first year and most just a few months after birth. This child was approximately 4 years old when it died.

It is astonishing that both men, women and children are buried in the nave of the church. Usually only the priests were buried inside the church in the Middle Ages – at least outside the urban centers. The question arises whether this church may have had a special status which was different from a normal parish church. If the church was privately owned, it would have been reasonably to find members of the family of the owners amongst the buried within the church. Therefore, it is possible that those who are resting here are family members of the main farm here in Sandur, to which the church belonged.

The great calcoid content in the sand and the fact that they have been located inside the following churches protected by the rain causes that the outstanding preservation of the skeletons. This material is of special value to the understanding of the Medieval Faroese population.

The archaeological context makes the church site in Sandur one of the most interesting and promising historical sites in the Faroe Islands[147].

Samples used for DNA analysis

VK25 Sandoy_Church2_grave28

VK27 Sandoy_Church2_grave8

VK44 Sandoy_Church2_grave29

VK45 Sandoy_Church2_grave13

VK46 Sandoy_Church2_grave14
VK234 Sandoy_Church2_grave23
VK236 Sandoy_Church2_grave27
VK237 Sandoy_Church2_grave32
VK238 Sandoy_Church2_grave22
VK239 Sandoy_Church2_grave16
VK240 Sandoy_Church2_grave7
VK241 Sandoy_Church2_grave17
VK242 Sandoy_Church2_grave31
VK244 Sandoy_Church2_grave24
VK245 Sandoy_Church2_grave15
VK248 Sandoy_Church2_grave_B
VK24 Á Bønhúsfløtu_Nesi_Hvalba_grave1

#### Salme, Estonia

**Jüri Peets, Tallinn University, Estonia**

In autumn 2008 remains of human skeletons and ancient artefacts (sword fragments, rivets, gaming pieces, dice, etc.) were brought to light while digging an electrical cable trench for the lighting of a cycling track near the Salme borough on the island of Saaremaa in Estonia. Archaeological rescue excavations revealed that the finds came from a partly destroyed boat burial (Salme I). It was probably a rowing ship approximately 11.5 m long and ca 2 m wide. Fragments of three swords and other weapons, a lot of gaming pieces etc. were found as grave goods[148]. Skeletal remains of seven individuals were found, including some partial skeletons *in situ* (Fig. S1.9); also a lot bones of domestic animals, and the remains of goshawk (*Accipiter gentilis*) and sparrow hawk (*Accipiter nisus*) were found[149].

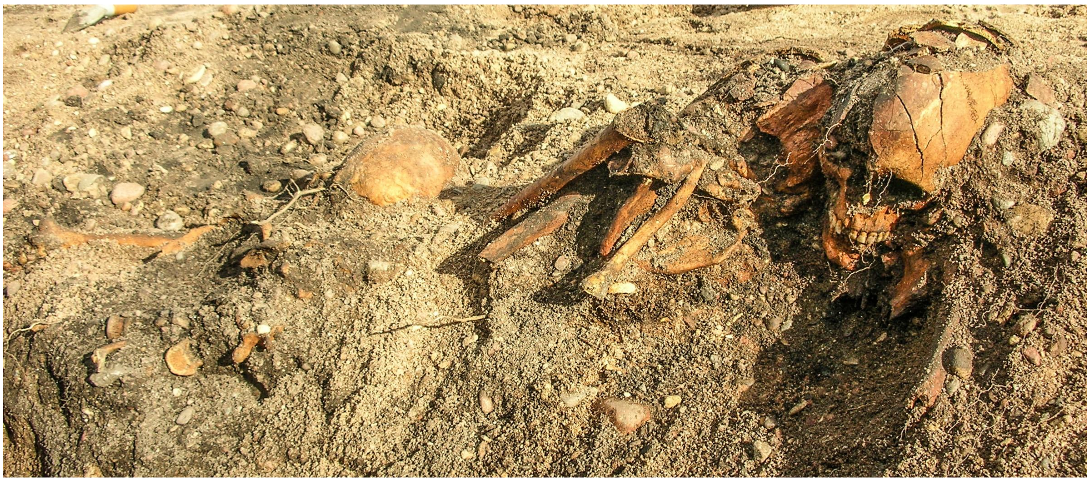

**Fig. S1.9**: Skeletal remains *in situ* in Salme I ship burial.

As a result of fieldwork during two summers (2010–2011) the second ship of Salme (Salme II) was discovered; practically the whole ship contour, which in the middle part could be observed to the height of 5–6 rivet rows, together with the remains of 34 warriors was unearthed[150,151]. The skeletons lay in three layers mostly in an area of about 3 x 4 m in the central part of the ship. In the two upper layers the perished warriors had been placed heads pointing NE, in the same direction as the ship. In the bottom layer they had been placed between the ribs, transverse with the longitudinal axle of the ship, heads pointing alternately east and west. The dead had been provided with rich grave goods, which mostly consisted of weapons, including more than 50 arrowheads, some spears and about 40 swords (whole and broken). At least five swords had hilts of gilded bronze, among them one ring-hilt sword with a blade of pattern welded steel. One sword had a blade ornamented with an inlay of golden wire and a handle decorated with garnets. Other items included nearly 300 gaming pieces of whale bone, antler combs, small padlocks, whetstones of schist, beads etc. The dead had been covered with shields with iron bosses. Besides the remains of meat animals, among osteological material of the Salme II ship some bones of goshawk and peregrine falcon (*Falco peregrinus*), and lots of mallard (*Anas platyrhynchos*) bones were recovered. At least five or six dogs had been buried in the ship, all in the top layer at the boards of the ship. The original length of the vessel, considering the proportions of the preserved remains, could have been 17–17.5 m or more[152]. Unlike the ship discovered in 2008, this ship may have moved also by sail[151].

Apparently the most important individual was the man (skeleton XII (L)) who was buried in the middle of the first row of the second layer in the mass burial of Salme II ship. He had a valuable ring sword (blade broken in several pieces), and warriors with shouldered swords placed at each side. Most likely the fallen warriors in the Salme I ship had been buried with as much homage and weapons as those in the second one. Apparently, they had been also covered with shields, of which, unfortunately, only shield rivets with domed heads have preserved. Artefact finds and $^{14}C$ analyses' results of samples of human bones suggest the origin of the burials in the same "event", that took place in about 750 AD. The "event" may have been an armed conflict and the perished warriors-seafarers were buried in two mass graves in the ships on the seashore. The shape of the weapons and other artefacts of both ships allow us to date them to the Late Vendel Period. Most of the finds belong to the Vendel Age period VII: 4 (c. 700–750 AD)[153] or earlier. An overwhelming part of human bones of both ships (4 samples of 7 persons from Salme I ship and 9 samples of 34 person from Salme II ship) were dated by $^{14}C$ (94,5%) to the 650–780 cal AD, but some samples, nevertheless, gave results up to 940 cal AD[148,150]. At the same time the earliest and the latest datings of samples from both ships are practically identical: the earliest is the dating of skull SaI: 3 from the Salme I ship, resulting in 1320±30 BP (Hela-1915), i.e. 650–780 cal AD and the latest of skeleton SaI: 4 that gave the result of 1200±30 BP (Beta-509634), i.e. 700–940 cal AD, whereas the respective datings of the samples from the Salme II ship are of skeleton XV (O), resulting in 1330±30 BP (Poz-109002), i.e. 649–767 cal AD and of skeleton XII (L), resulting in 1199±32 BP (Ua-50746), i.e. 700–940 cal AD, both skeletons originated from the second burial layer (Extended Data Fig. 1a). Most results of the analyses of animal bones remain within these limits. The reason for such great difference in the dating results is unclear. All analyzed samples were taken from materials without any later impact, i.e. all samples were taken from bones that were not disturbed after burial. Resolving this phenomenon is of utmost importance, considering that four skeletons, identified genetically as brothers, and the dating of their skeletons also differed considerably. In addition to the above mentioned skeleton XV (O), dated as the oldest (1330±30 BP), also skeletons of two other brothers were dated, i.e. skeleton XIV (N) 1295±32 BP (Ua-50747), and XXII (V) 1250±30 BP (Beta-509632). The skeleton XXVI (Ö), the fourth brother is currently not dated. Results of isotope analysis (study of $^{87}$Sr/$^{86}$Sr, $^{18}$O/$^{16}$O and $^{13}$C/$^{12}$C isotopes value in human dental enamel and bones) show that the warriors buried in the mass graves in the ships came from Central Sweden[30].

Samples used for DNA analysis

VK480 Estonia_Salme_II-VI(E)
VK481 Estonia_Salme_II-IV(F)
VK482 Estonia_Salme_II-XVI(P)
VK483 Estonia_Salme_II-XXII(V)
VK484 Estonia_Salme_II-XVII(Q)
VK485 Estonia_Salme_II-XV(O)
VK486 Estonia_Salme_II-VII(G)
VK487 Estonia_Salme_II-I(A)
VK488 Estonia_Salme_II-VIII(H)
VK489 Estonia_Salme_II-XXV(Ä)
VK490 Estonia_Salme_II-XIV(N)
VK491 Estonia_Salme_II-XXIV(Õ)
VK492 Estonia_Salme_II-II(B)
VK493 Estonia_Salme_II-XXXIV(Š)
VK495 Estonia_Salme_II-III(C)
VK496 Estonia_Salme_II-XXIII(W)
VK497 Estonia_Salme_II-XXVI(Ö)
VK498 Estonia_Salme_II-XXXII(Z)
VK504 Estonia_Salme_I-1
VK505 Estonia_Salme_I-2
VK506 Estonia_Salme_I-3
VK507 Estonia_Salme_I-4
VK508 Estonia_Salme_I-5

VK509 Estonia_Salme_I-6
VK510 Estonia_Salme_I-7
VK511 Estonia_Salme_II-XXVIII(X)
VK512 Estonia_Salme_II-XXVII(Ü)
VK549 Estonia_Salme_II-X(J)
VK550 Estonia_Salme_II-V(D)
VK551 Estonia_Salme_II-XXI(U)
VK552 Estonia_Salme_II-XI(K)
VK553 Estonia_Salme_II-XIII(M)
VK554 Estonia_Salme_II-XII(L)
VK555 Estonia_Salme_II-IX(I)

#### Glyn, Llanbedrgoch, Anglesey (Wales)

**Mark Redknap, National Museum Cardiff**

Excavations by Amgueddfa Cymru – National Museum Wales at Glyn, Llanbedrgoch between 1994 – 2001, in 2004 and in 2012 uncovered a small early medieval settlement which by the late ninth century had developed into a large entrepôt and regional centre about one hectare in size serving the kingdom of Gwynedd. During the second half of the ninth century a stone rampart was built to provide additional security[154]. In the second half of the ninth century rectangular timber buildings with ground-fast posts were remodelled, having sunken floors and sill-beam construction resting on low stone walls. A rich array of ninth/tenth-century artefacts reflect an Insular mix of local, Anglo-Saxon and Irish styles, as well as contact with the Scandinavian world (Hiberno-Norse hack-silver, fragments of Kufic silver coin, lead weights of different types, some capped with decorative metalwork, ringed pins and a fragment of oval brooch).

The excavations have provided evidence for the organised use of space for activities such as iron working, baking, bronze smithing, waste disposal, sleeping, cooking and burying. An unexpected bonus was the discovery in 1998 and 1999 of the remains of five people buried in a seemingly haphazard manner in the upper fill of the ditch outside the rampart. This raised questions about not only the identities of those buried so unceremoniously, but also the settlement's fate, who occupied it during the tenth century and their status. The preliminary osteological report identified a young adult female, a young adult male, a mature adult male and two adolescents, and radiocarbon dates pointed to their deaths occurring in the mid- to second half of the tenth century, burial possibly later in the early eleventh century[154].

The irregular disposition and orientation of the bodies and the observation that the wrists of one adult male (burial 3) appeared to have been tied behind the back, suggested a non-normative burial rite given that Christian burial within cemeteries was, by now, standard for the local population. Radiocarbon dates, stratification and the absence of late-tenth or early eleventh-century coins suggested that they had been buried during a final phase of occupation of the site in the tenth century[154].

During facial reconstruction work it was suggested that some of these features pointed to a genetic relationship between the skulls, either familial or more probably because the individuals originate from a small gene pool (Caroline Wilkinson, *in litt.*). Consequently it was considered possible that they represented local islanders, perhaps the victims of a Viking attack on the Anglesey[154]. This view is no longer held[155].

Since 2004, the human remains from Llanbedrgoch have undergone a complete osteological reassessment by Professor Alice Roberts in order to establish the demographic profile and health status of the individuals. This has refined the estimated age at death, and included a further inhumation found in 2012, just north of the 1998-99 inhumations.

Multiple isotope analysis for palaeodietary reconstruction and to identify the childhood place of origin of these individuals has now been conducted on burials 1-7 by Katie Hemer as part of her PhD, the analytical work being undertaken at the NERC Isotope Geosciences Laboratory in Nottingham following the award of a grant from NERC. Strontium results provide the most compelling evidence in the case of the Llanbedrgoch individuals because the signatures are distinctive. Full report on these burials is in preparation (Hemer, Roberts and Redknap, in prep.).

Juvenile burial 4 (context 737) sampled for DNA analysis, provided two radiocarbon dates: Beta-150715: 1030+/-40BP, Cal AD 960-1040; OxA-33786, 1055+/- 40 (2016) Cal AD 892-1031. Stable isotope analysis (strontium and oxygen) of this juvenile (9.5-14 years old) by Dr Katie Hemer suggests that he may have grown up between the ages of three and seven in parts of mid-Wales or on the border of England, while the other adult male (burial 3, with his wrists tied) in the same burial had strontium values consistent with old or radiogenic geologies found in parts of Scotland, Denmark or further afield in southwest Norway, and oxygen values within the range expected for the British Isles. None of the extra-mural burials grew up on Anglesey, and they appear to have grown up in places with diverse geologies and climate[155].

The archaeological context of this extra-mural cluster suggests that they may have been treated in death as incomers, outcasts because they were not local and perceived to be non-Christian, perhaps instilling fear or suspicion in the minds of those responsible for their burial. Bearing in mind the raids from Man in the 970s and 980s, some may have been hostages or slaves of the Welsh, if not free traders. These results are still under discussion.

Samples used for DNA analysis

VK171 Wales_Cardiff_burial#4_GL99
