## Supplementary for "Population genomics of the Viking world"

### SUPPLEMENTARY INFORMATION

#### Supplementary Note 2. Ancient DNA laboratory procedures and sample selection

##### Ancient DNA extraction

All aDNA laboratory procedures were conducted in the special aDNA clean-room facilities at Centre for GeoGenetics, Natural History Museum, University of Copenhagen according to strict aDNA standards[156,157]. The overwhelming majority of ancient samples were petrous bones and teeth (Supplementary Table 1). To maximize the yield of endogenous DNA from ancient human samples we targeted the cementum (the hard and relatively well preserved outer layer of the tooth roots)[158] or the otic capsule of the petrous bones[159].

The drilled bone sample (ranging from 100 to 400 mg) was briefly digested in digestion buffer (4.65 ml 0.5 M EDTA, 50 µl recombinant Proteinase K, 50 µl 100x TE and 250 µl 10% N-Laurylsarcosyl) for 45 minutes at 40˚C (pre-digestion step)[158]. After this, the samples were centrifuged for 2 minutes at 2000g and the resulting supernatant was removed. To the bone material, an identical digestion buffer was added for a full 24-hour digestion at 40˚C. After this digestion step, the samples were centrifuged for 5 minutes at 2000g and the remaining undigested pellets were stored for later re-extraction. Silica-powder-based DNA extraction protocol was used to extract the aDNA from 2 ml digested solution. The water based suspension of silica was prepared by mixing 6g of SiO2 with 50 ml H2O. After 1 hour of sedimentation, the top 48 ml supernatant was transferred to a new 50 ml tube. This was followed by another 5-hour sedimentation after which the top 43 ml of the supernatant was discarded and the silica was re-suspended and activated by 60 µl 37% HCL. To each of 2 ml digested sample, 20 mL of the binding buffer (19.54 ml Qiagen buffer PB, 360 µl 5M sodium acetate, 100 µl 5M sodium chloride) and 100 µl silica suspension was added and adjusted to pH 4-5 with 37% HCl[160]. This was followed by a 1-hour incubation at room temperature after which the supernatant was removed by a brief centrifugation step for 2 minutes at 2000×g. The silica was re-suspended in 1 ml binding buffer, transferred to 2ml Eppendorf tubes, centrifuged and washed twice with 80% ice-cold ethanol. The DNA was released from silica particles by 70 µl Qiagen EB buffer. With each round of extractions, negative controls were used.

#### NGS library preparation and sequencing of ancient samples

From 20 µl double-stranded DNA extracts blunt-end DNA libraries were prepared using Illumina-specific adapters and NEBNext DNA Sample Pre Master Mix Set 2 (E6070) kit according to the manufacturer's instructions with few below-mentioned modifications:

The end-repair step was conducted in 25 µl reactions using 20 µl of DNA extract. This was incubated for 20 mins at 12°C and 15 mins at 37°C, purified using PB buffer with Qiagen MinElute spin columns, and eluted in 17 µl EB buffer. This was followed by ligation (25 µl reactions) of the Illumina-specific sequencing adapters prepared according to Meyer and Kircher 2010[161]. This step was carried out for 15 mins at 20°C and the resulting DNA-adapter complex was purified using PB buffer and Qiagen MinElute columns, before eluting in 20 µl EB Buffer. The last, adapter fill-in reaction was conducted in 30 µl volume and incubated for 20 mins at 65°C followed by 20 mins at 80°C to inactivate the Bst enzyme. qPCR was conducted using SYBR green MIX (Roche) according to manufacturer's instructions and the same forward and reverse primers used for the subsequent index PCR step in order to calculate the total amount of DNA library in each sample and assess the optimal number of PCR cycles required for DNA library amplification at the subsequent step. After that, the 12µl DNA library was index-amplified in a 50 µl PCR reaction by mixing with 25 µl 2X Kapa U+, 1 µl of each primer (10 µM, inPE forward primer + indexed reverse primer) and 11 µl $H_2O$. The PCR thermocycling conditions were 45s at 98°C, followed by number of cycles (based on qPCR values) of 15s at 98°C, 30s at 65°C and 30s at 72°C, and a final 60s elongation step at 72°C. The amplified DNA library was purified with PB buffer using Qiagen MinElute columns and eluted in 50 µl EB buffer. Negative library controls, based on EB as well as negative extraction controls were included for each batch. To quantify the amount of the purified DNA libraries, Agilent Bioanalyzer 2100 was used. The library pools were sequenced (80 bp single read) on Illumina HiSeq 2500 machines at the Danish National High-throughput DNA Sequencing Centre. The base-calling and sequence sorting by sample-specific indices were produced by the Sequencing Centre using CASAVA v1.8.2.

#### Sample selection

A total of 528 ancient human remains were processed and screened (c. 15 million sequences per sample) for contamination levels and endogenous human DNA content. After removing contaminated and poorly preserved samples the number reduced to 442. Many of the remaining samples were prioritized based on their endogenous human DNA content and relevance to the project and were sequenced deeper resulting in 376 samples between 0.1 and 11.7X coverage, of which 216 were above

1X. A total of 64,786,513,002 DNA reads from 442 different samples were generated for this study, out of which 23,149,730,287 uniquely mapped to the human reference genome (Supplementary Table 2).

#### Supplementary Note 3 - Data quality assessment, contamination, error profiles and sex determination

AdapterRemoval v2.1.3[162] was used for removing Illumina adapter sequences and stretches of Ns at both ends of the ancient DNA reads, keeping only sequences with a minimum length of 30 bp. We mapped the adapter-free sequences against the human reference genome build 37 using BWA v0.7.10 aligner[163] with the seed (-l parameter) disabled for higher sensitivity of ancient DNA reads[164]. DNA sequences were processed with samtools v1.3.1[163] and only aligned sequences with mapping quality of at least 30 were kept. Picard v1.127 (http://broadinstitute.github.io/picard) was used to sort the reads and remove duplicates. DNA libraries were combined at sample level and realigned using GATK v3.3.0[165] with Mills and 1000G gold standard indels. At the end, realigned bams had the md-tag updated and extended BAQs calculated using samtools calmd. Read depth and coverage were determined using pysam (http://code.google.com/p/pysam/) and BEDtools[166]. The mapping statistics for the ancient samples are summarized in Supplementary Table 2.

DNA damage is one of the most characteristic features of ancient DNA molecules and is usually manifested in the form of single or double stranded breaks resulting in fragmented DNA molecules (usually <100 bp) and single-stranded overhangs, as well as cytosine deamination towards the 5' end of DNA fragments leading to the typical C→T transitions in the sequenced DNA reads.
We have used mapDamage v2.0 to obtain approximate bayesian estimates of damage parameters[167]. Three main parameters were assessed: (i) the C→T transitions rates at the first position of the 5' end of DNA reads, (ii) $\lambda$, the proportion of nucleotides in single-stranded overhang regions, and (iii) $\delta s$, which shows the estimated C→T transition rate for the single-stranded overhang segments.

Of the 442 ancient samples, the C → T transition rates at the first position of the 5' end of the DNA fragments ranged from 1.5% to 42.4% when comparing with the human reference genome (Figure S3.1). Both the lambda ($\lambda$) and DeltaS ($\delta s$) parameters show a significant deviation from zero, indicating that the bulk of the recovered DNA molecules were damaged and degraded which suggested that the majority of DNA molecules were of ancient origin.
The DNA damage parameters of each ancient sample from this study is presented in Supplementary Table 4.

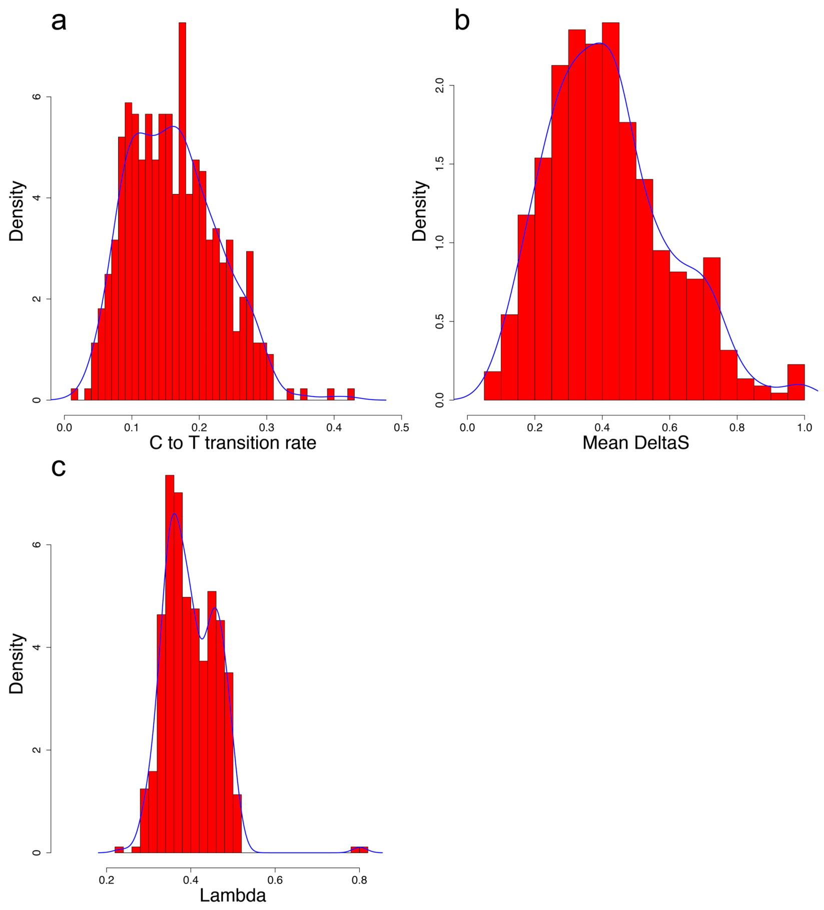


**Fig. S3.1**: Distribution of the ancient samples according to the DNA damage levels. (a) C→T transition rates; (b) transition rates in single-stranded overhangs; (c) fraction of bases in single stranded overhangs.

#### Contamination

**ContamMix:** This method relies on the reconstructed consensus mtDNA sequences of ancient samples; thus, the contamination should not exceed 50% for this method to work. The details of ContamMix is described elsewhere[168]. It also assumes that the sequenced mtDNA reads originate from a mixture of the reconstructed consensus sequence and 311 whole mitochondrial genomes, representing possible contaminant sequences from present-day worldwide populations. By comparing the mapping affinity of each ancient mtDNA read to the reconstructed consensus sequence and to the 311 possible contaminants, ContamMix reports the total fraction of non-endogenous reads as the contamination rate with a Bayesian estimate of the posterior probability of the contamination proportion.

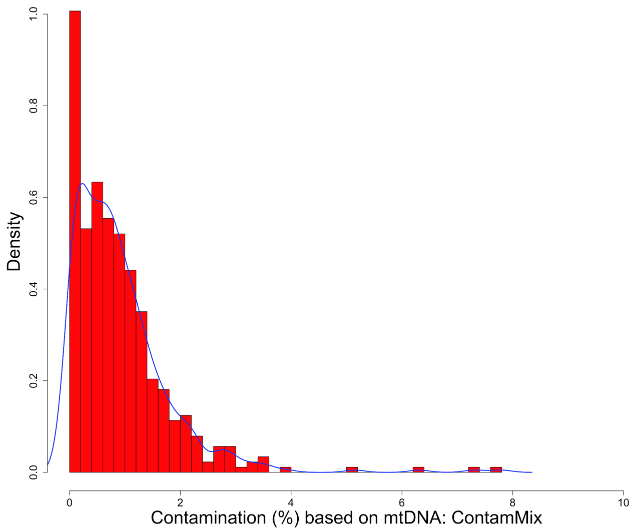


**Fig. S3.2**: Distribution of the ancient samples according to the contamination levels. All ancient samples have >9X sequencing depth (n=442) on mtDNA. The four samples with more than 5% contamination on the plot did not show significant contamination based on Schmutzi, therefore were kept for downstream analysis.

For this approach, we aligned all trimmed reads from ancient samples to the human mitochondrial reference genome: revised Cambridge Reference Genome (rCRS) with the same parameters as for the whole genome mapping. The sequencing depth of coverage for the mtDNA ranged 9-538X, the details are provided in the Supplementary Table 2. To construct the endogenous mtDNA sequence of ancient samples required for ContamMix we have used an in-house perl script. mtDNA sites with base quality <20 and reads with mapping quality <30 were discarded. Only SNPs and sites with at least 3X coverage were considered for consensus calling. At each mtDNA position a base was called only if it was observed in at least 70% of the reads covering that site. The distribution of the ancient samples based on contamination levels assessed by ContamMix are shown in Figure S3.2. The contamination estimates are presented in Supplementary Table 4.

**Schmutzi**: The amount of present-day human contamination was also estimated in each sample with Schmutzi[169]. This estimate was performed without the inclusion of the predicted contaminant in the database of putative present-day human contaminant mitogenomes used by Schmutzi (option "--notusepredC"). Unlike ContamMix, it does not estimate a parameter for error but instead uses input mis-incorporation rates due to deamination and uses base quality scores. Schmutzi also assumes a single contaminant source unlike ContamMix.

Both methods were applied to our dataset and only samples with contamination levels greater than 5% detected with both methods were removed.

Schmutzi was also used to infer the endogenous consensus sequences by mitigating the impact of deamination-induced nucleotide mis-incorporations and the presence of present-day human contamination. The former is achieved by incorporating the rates of mis-incorporations into a Bayesian model that considers every possible 4 base for the endogenous and the contaminant. Each aDNA fragment is assigned a probability of being endogenous given the contamination prior provided above. This probability is computed using the distribution of fragment lengths and the rate of base mis-incorporations due to deamination. The most likely base is produced along with a per-base error probability on a PHRED scale.

**Male X-chromosome based contamination analysis**: Since male individuals carry only 1 X chromosome inherited from the mother, any heterozygous position of that chromosome (outside the pseudoautosomal regions) would be either due to sequencing errors or contamination. In case of sequencing errors, the heterozygous positions are expected to be randomly distributed across the chromosome, while in case of contamination heterozygous positions should be restricted to mainly diagnostic positions (sites on the X chromosome that are known to be polymorphic). The difference of mismatch rates between these positions and adjacent sites indicates the contamination levels. This method is described in more detail in previous work[170] using the package ANGSD[171]. For this analysis, we removed the pseudoautosomal regions of the X chromosome and used mapping quality ≥30 and base quality ≥20 filters. The reported values are based on the maximum likelihood estimator using the unbiased sampling-based approach, i.e. “Method1” in ANGSD (Figure S3.3).

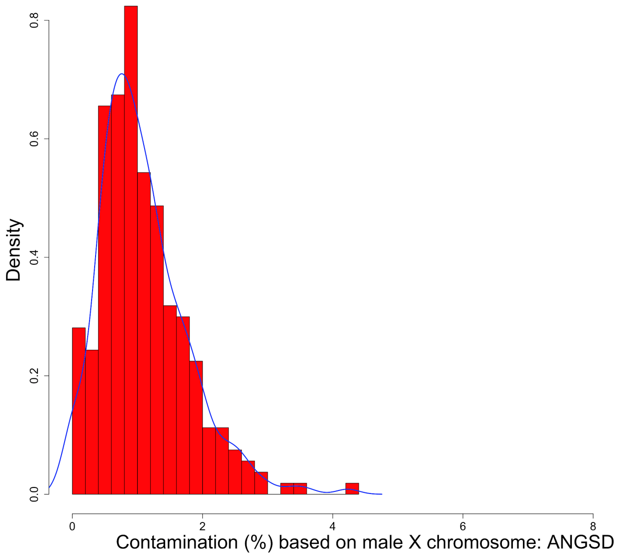


**Fig. S3.3**: Distribution of the ancient samples according to the X-chromosome based contamination levels. Only male individuals with >0.1X average genomic depth of coverage were used for the analysis (n=267).

#### Sex determination

The sex of ancient individuals was determined by calculating the Rγ parameter, which reflects the ratio of the fraction of Y chromosome reads to the fraction mapped to both X and Y chromosomes[172]. According to this method, individuals with Rγ values above 0.077 are considered as male, while the ones lower than 0.016 as female.

We assessed the sex of all the ancient samples in this study, regardless of the sequencing depth. We identified 141 females and 296 males (Supplementary Table 5) while the biological sex for the rest of the 5 samples were not identified (4 of which due to low coverage). This strong male bias was expected, since in many of the famous sites such as Dorset-UK (executed c. 50 males) and Salme-Estonia (2 ship burials of male warriors), only male individuals were buried. Moreover, whenever feasible (with all other factors being equal - e.g. age of the sample or preservation), males were prioritised due to the extra genetic information carried in the Y-chromosome. Interestingly, one of the ancient samples with unidentified biological sex (VK204) had c. 1X average genomic coverage. The inability of this method to identify the sex of this individual may indicate that this person was affected by some form of Klinefelter syndrome and had one of the non-usual karyotypes (e.g. XXY or XXXY). Given the relatively high rate of occurrence of this syndrome with roughly 1 in 576 males[173], it is not unlikely to observe one case in such large number of ancient samples.

#### Mitochondrial DNA analysis

For mtDNA haplogroup assignment we used the mtDNA consensus sequences created by Schmutzi. The mitochondrial haplogroups of the ancient Viking Age individuals were assigned using haplogrep[174].

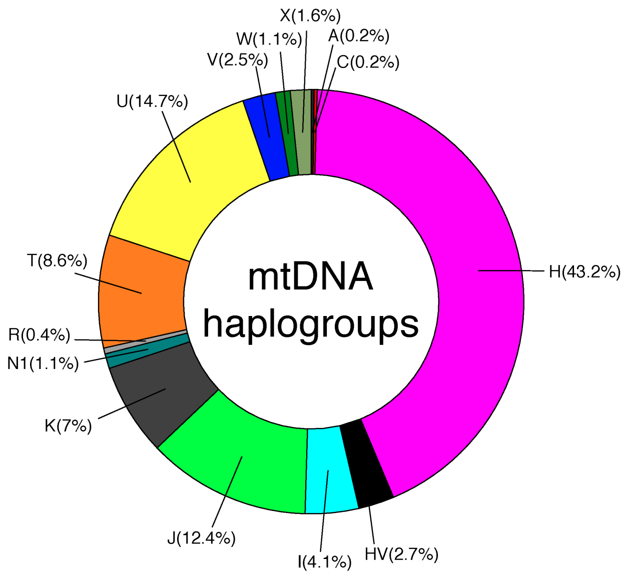


**Fig. S3.4**: The distribution of mtDNA haplogroups across the 442 ancient samples from this study. The haplogroup frequencies should be interpreted with caution since there are a few related individuals in this study especially from Faroe Islands and Salme boat burials (see the genetic relatedness section) which were not removed for the frequency estimation of the whole dataset.

The overall distribution of the mtDNA haplogroups of the ancient Viking age samples (Figure S3.4) is quite similar to the diversity of mtDNA lineages found in modern northern European populations. Namely, the frequently encountered (>5%) major mtDNA haplogroup were Hg H (43%), followed by Hgs U, J, T and K (http://www.eupedia.com/europe/european_mtdna_haplogroups_frequency.shtml). Interestingly, we found two individuals VK548 (a female individual from Norway; Nord-Trondelag 3705) and VK160 (a male individual from Russia; Kurevanikka_7283-3) who had mtDNA haplogroups commonly found in Asian populations: haplogroups A12a and C4a1a, respectively. The VK548 sample was previously suggested to have had the haplogroup A4b based on mtDNA HVR1 data[57].

The haplogroup frequencies in Figure S3.5 should be interpreted with caution since there are few related individuals in this study especially from Faroe Islands and Salme boat burials (see Supplementary Note 4) which were not removed for the frequency estimation of the whole dataset. The aligned whole mtDNA sequences of Viking age samples (roughly dating 1000 year ago) were used as an input for BEAST v1.8.4[175] to uncover the trajectory of the effective female population size (Ne) through time using the Bayesian skyline plot (BSP) method.

The MCMC chains were run for 10E8 states and sampled every 10E4 states with the first 10E6 states discarded as burn-in. We used the CIPRES open-access server for phylogenetics studies[176] to run the BEAST analysis. We checked the output data for convergence to a stationary distribution and sufficient effective sample size estimates using Tracer v1.7[177]. We used the GTR model with gamma plus invariant sites and a strict clock with a normal prior of 2.2E-8/site/year as the mean value with standard deviation of 2.2E-9. The resulting trees were annotated with TreeAnnotator 1.8.4 (BEAST package) and visualised by FigTree (http://tree.bio.ed.ac.uk/software/figtree/).

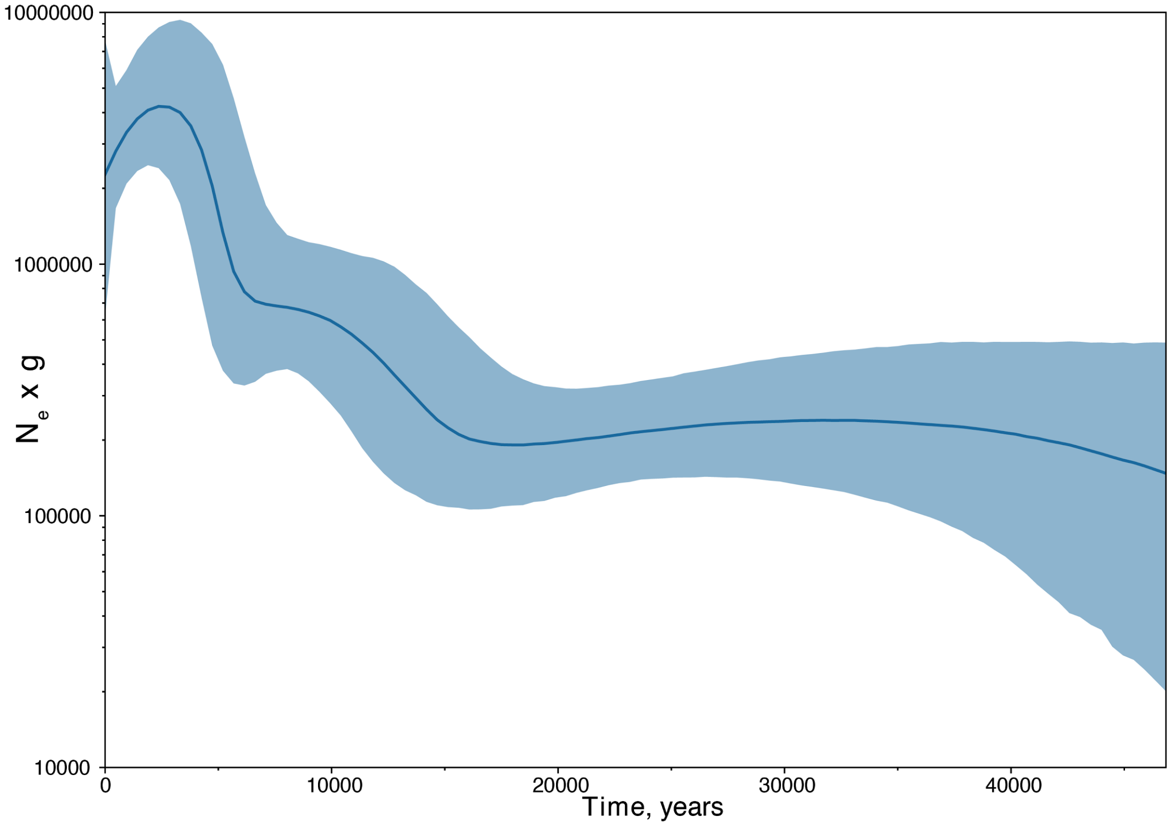


**Figure S3.5**: Bayesian skyline plot based on the ancient samples. Values on the y axis represent the effective female population size ($N_e$) x generation time (g).

A Bayesian Skyline plot analysis is shown in Figure S3.5. Even though the 95% confidence intervals are large, it shows an increase in female population size for the last 10-15 thousand years with a mean female population size of c. 100,000 assuming a generation time of 25 years. The increase in female population size after the Last Glacial Maximum (LGM) is in accordance with previously published data[178].

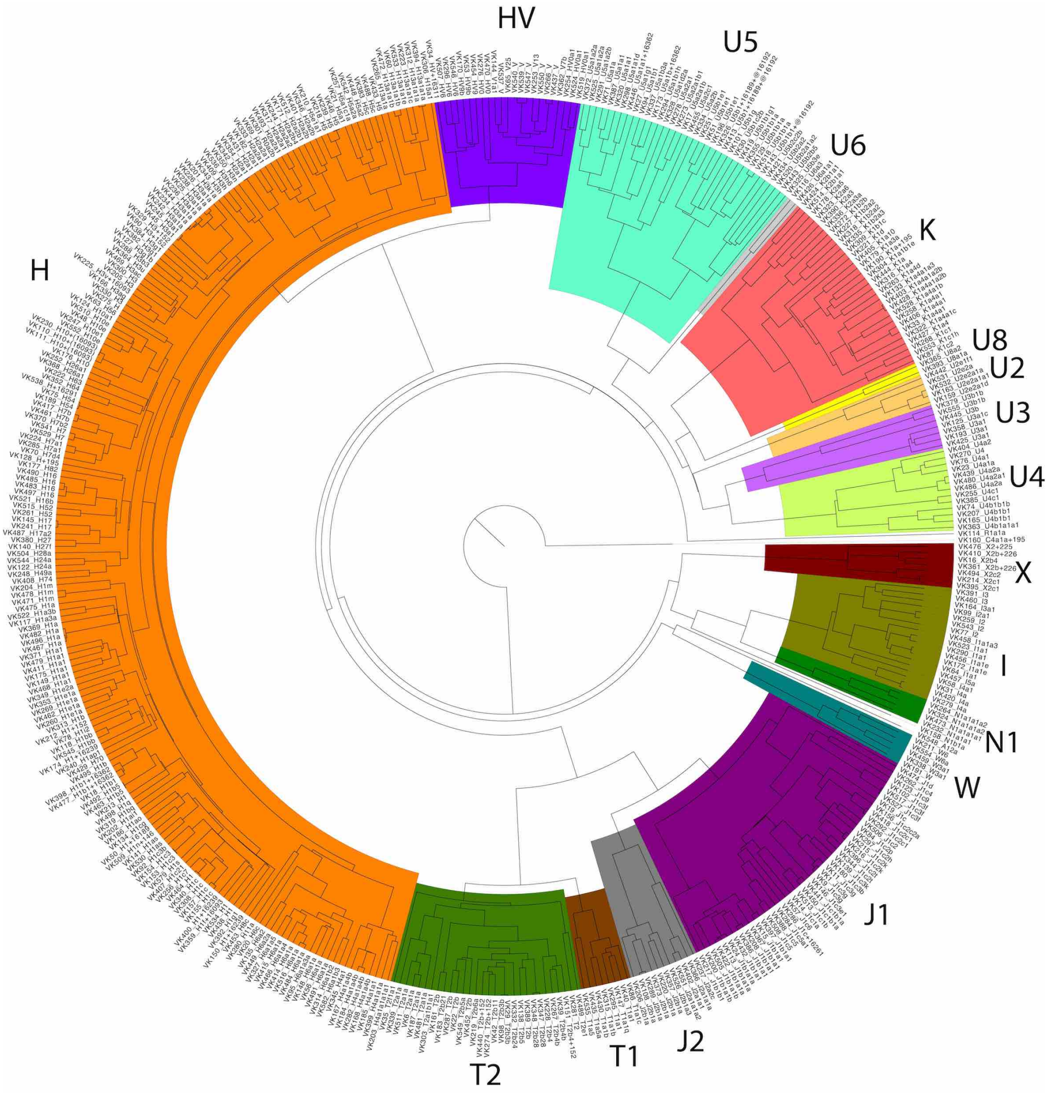


**Figure S3.6**: Phylogenetic tree of ancient samples from this study based on whole mtDNA sequences. The tip labels indicate the ancient sample IDs and mtDNA haplogroups.

A Bayesian phylogenetic tree of mtDNA sequences of all 442 ancient samples is presented in Figure S3.6.

#### Y chromosome haplogroup analysis

We assigned Y chromosome haplogroups to the ancient male samples with Yleaf v2[179], restricting our analysis to 26,083 biallelic SNPs from the ISOGG (International Society of Genetic Genealogy) 2019 database (https://isogg.org/tree/ISOGG_YDNA_SNP_Index.html) confidently placed within the Y-chromosome tree (i.e. excluding those annotated with '~', of uncertain placement, to avoid inconsistencies in the determination of haplogroups). The distribution of the Y chromosome haplogroups is presented in Figure S3.7.

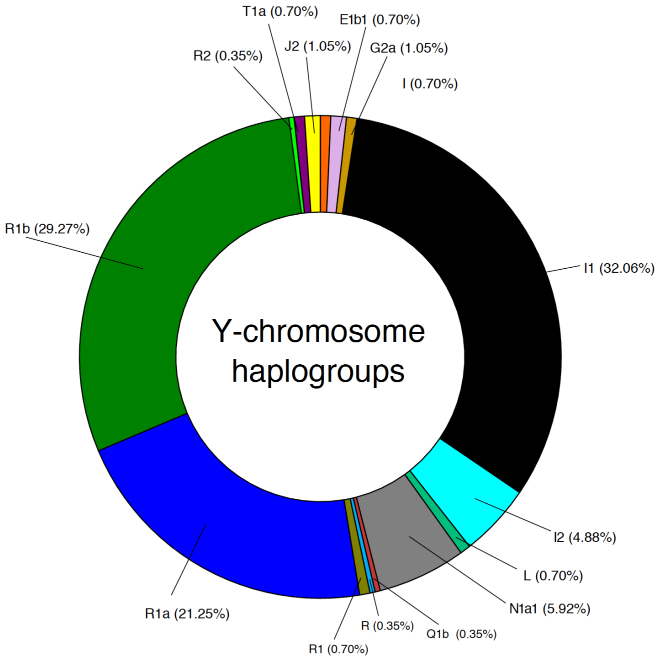


**Fig. S3.7**: Distribution of Y chromosome haplogroups in 276 ancient (mostly Viking Age) male individuals.

The assigned haplogroups are provided in the Supplementary Table 5. As in the case of mitochondrial DNA, the overall distribution profile of the Y chromosomal haplogroups in the Viking Age samples was similar to that of the modern North European populations. The most frequently encountered male lineages were the haplogroups I1, R1b and R1a.

Among the ancient samples, two individuals were derived haplogroups were identified as E1b1b1-M35.1, which are frequently encountered in modern southern Europe, Middle East and North Africa[180]. Interestingly, the individuals carrying these haplogroups had much less Scandinavian ancestry compared to the most samples inferred from haplotype based analysis. A similar pattern was also observed for less frequent haplogroups in our ancient dataset, such as G (n=3), J (n=3) and T (n=2),, indicating a possible non-Scandinavian male genetic component in the Viking Age Northern Europe. Interestingly, individuals carrying these haplogroups were from the later Viking Age (10$^{th}$ century and younger), which might indicate some male gene influx into the Viking population during the Viking period. Worth mentioning, that due to the small sample size of the rare haplogroups, these differences might be of stochastic nature therefore the results based on uniparental markers should be interpreted with caution.

**pathPhynder Analysis**: The highly degraded nature of aDNA data poses considerable challenges for combining ancient and modern samples for phylogenetic analyses. Typical analyses of ancient Y-chromosome data focus on a curated set of known markers (ISOGG), however, many more markers exist and will continuously be generated as more sequencing studies emerge. By making use of this additional Y-chromosome variation, we can increase the probability of overlap of ancient DNA reads with branch-defining markers and use this information to place ancient samples into more detailed phylogenies.

Using the pathPhynder workflow (https://github.com/ruidlpm/pathPhynder), we first assigned 56,246 variants from the 1000 Genomes Project to the branches of the Y-chromosome tree[181]. Next, for each aDNA sample, we generated a pileup at these informative sites, excluding C/T and G/A sites covered by a single read to minimise the impact of post-mortem deamination (‘conservative’ mode). Next, for each ancient sample, we determined the number of ancestral and derived alleles at each branch and traversed the 1000 Genomes Y-chromosome tree, using this information to map ancient samples to their most likely place in the phylogeny (Figure S3.8).

In the 1000 Genomes Project Y-chromosome phylogeny, haplogroup N is split into two main clades, one composed of mostly East Asian individuals (CHB) and one of Finnish individuals (FIN). The Viking and Early Viking samples from Estonia and Sweden are positioned within the FIN clade, more specifically at the branch defined by the marker VL29/CTS2929-N1a1a1a1a1a or at branches downstream of it. A large study of Y-chromosomes revealed that the VL29 lineage occurs at highest frequencies in present-day Estonians, Latvians, Lithuanians, Finns and other northern European populations[182].

Haplogroup I1, previously described as showing the signature of a star-like expansion (Poznik 2016), occurs in present-day Scandinavians at a frequency of 25–35%[183]. In the 1000 Genomes Project, this clade is mostly composed by FIN, GBR and CEU populations. I1 is the most well represented haplogroup in our ancient dataset, and most regions contained individuals belonging to this lineage (especially Estonia, Russia, Denmark and Greenland), with the exception of the Isle of Man, Ireland and Italy, however these regions have comparatively small sample sizes which may not allow the detection of this haplogroup. The ancient samples of the present-study are mainly distributed in two main clades, I1a1b1-L22, which accounts for 71% of the I1 haplotypes in a Y-chromosome survey of Finland[184], and I1a2a-S246. Of particular interest, the clade I1a2a1a1d1a-S247 is especially well represented in Estonian samples, and is found mostly in present-day Finnish and Northern Scandinavian groups.

While it likely that many I1 lineages dispersed into the British Isles from Scandinavia during the Viking period, it is also probable that earlier Anglo-Saxon migrations also played a role, as suggested by the East to West decreasing frequency cline of I lineages[185] and the presence of the I1 lineage in an individual from this period excavated in England[186]).

Within haplogroup I2, two Viking samples from Sweden and Ukraine, fall within the L621-I2a1a2b and its downstream I2a1a2b1-CTS10936 clade, together with a FIN individual. This lineage has a mostly Eastern European distribution[187], occurring more frequently in Slavic peoples. Other I2 ancient samples fall within the clade defined by I2a1b1-M223, with highest frequencies in Germanic and Scandinavian peoples[183]. A subset of these individuals from Sweden, Orkney and England was positioned in the I2a1b1a2b-Z161 branch, a lineage present in North Europe, especially in Denmark, Germany, the Netherlands, and England.

In terms of haplogroup R, only a single R2-M9710 individual was found in Iceland (VK123), and was positioned together with mostly South Asian populations of the 1000 Genomes project, while all other ancient samples belong to R1 subclades.

Within Europe, the subclades of R1, R1a-M420 and R1b-M343, reach higher frequencies in the East and West, respectively. In broad terms, the distribution of R1 lineages in our ancient dataset is consistent with this, with the Isle of Man and Italy carrying exclusively R1b derived lineages, and Ukraine and Estonia R1a derived lineages.

Focusing on R1a, two major lineages exist, one more common in Central and South Asian populations (R1a1a1b2-Z93), and the other in Europe (R1a1a1b1a-Z282)[188] (Underhill 2015). As expected, the R1a derived ancient samples here analysed fall within the European Z282 clade, and are positioned near Great British (GBR), European (CEU) and Finnish (FIN) individuals in the Y chromosome tree, rather than with the South Asian individuals in the Z93 clade. Many of the ancient Norwegian and

Swedish samples were determined to be derived for the R1a1a1b1a3a-S221/Z284 marker, of nearly exclusive distribution within Scandinavian populations and occurring at approximately 20% in Norway[188].

In terms of R1b lineages, the R1b1a1b1b-Z2105 and R1b1a1b1a-L11 clades have distinct distributions, with the first being typically associated with the Yamnaya[160,189,190] and distributed today around the Caucasus and Volga-Uralic regions[187], and the latter, more commonly found in Late Neolithic, Bronze Age and later periods in Central and Western Europe. A derived status at R1b1a1b1b-Z2105 was found in a single sample from Italy (VK535). The vast majority of other R1b samples belonged instead to the R1b1a1b1a-L11 clade, and were mostly distributed within its two main subclades: P312/S116-R1b1a1b1a1a2 and U106/M405-R1b1a1b1a1a1[191], of western and eastern distribution relative to the Rhine river basin[192].

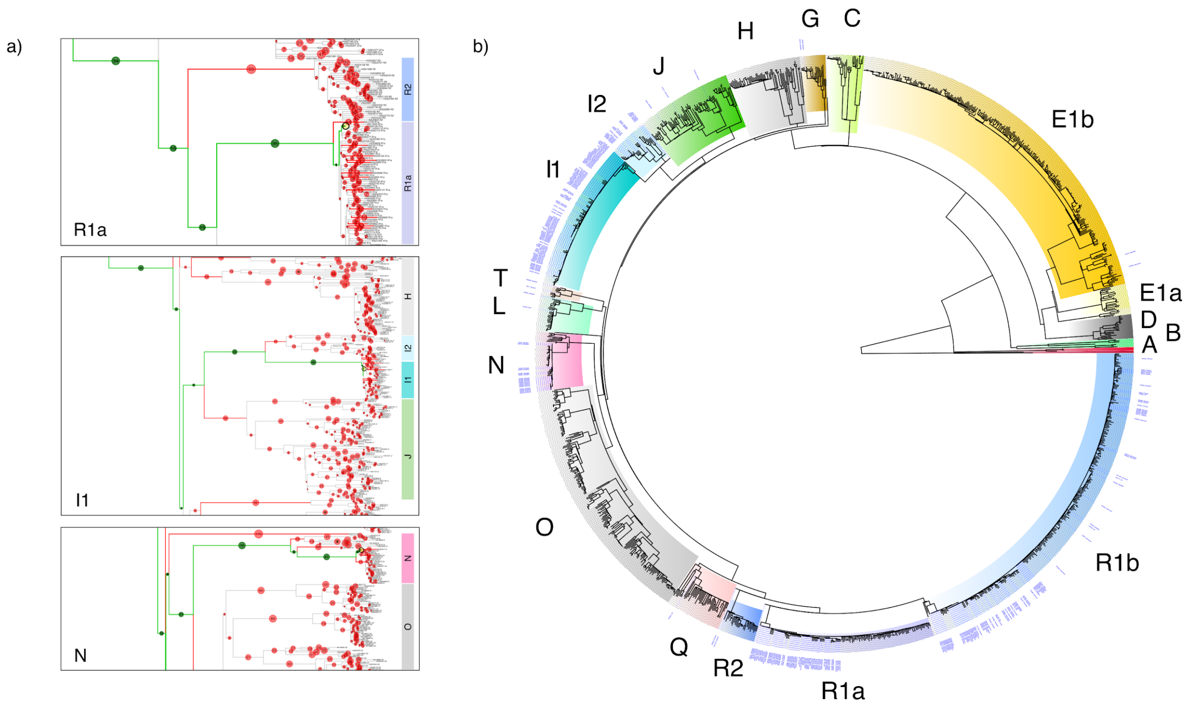


**Fig. S3.8**: Placing ancient DNA samples within the 1000 Genomes Y-chromosome tree: a) Examples of the identification of the best path within the Y-chromosome tree for 3 representative ancient samples. The phylogeny is traversed and the number of derived (green circles) and ancestral (red circles) markers present in a given ancient sample are evaluated in order to identify the best path (green branches) and position (yellow circle) where this individual can be placed. b) 1000 Genomes Y-chromosome phylogeny with 284 ancient samples (labelled in blue) added to the most likely place of the tree according to the number of derived and ancestral markers carried at each branch.

### Supplementary Note 4 - Kinship analysis

#### Introduction

Since many of the ancient samples were sampled from the same archaeological sites, it seemed possible that there would be some level of genetic relatedness between certain pairs of individuals across our dataset. To assess to what extent this was the case we used two different methods implemented in two programs NgsRelate[193] and READ[194] that have previously shown to be relatively robust for low coverage NGS data. In particular, we used NgsRelate to detect family relationship between all pairs of samples and double checked all detected first and second degree relationships with READ.

#### Methods

**READ (Relationship Estimation from Ancient DNA):** Since the genetic relatedness analysis can be difficult with low coverage data, we have used the program READ (in addition to the NgsRelate, see below section) which was shown to successfully work with ancient NGS data as low as 0.1X sequencing depth for determining up to second degree relationships[194]. This method estimates the proportion of non-matching alleles for each pair of individuals and uses the same statistic from unrelated individuals for normalization purposes. Based on the normalised proportions of shared alleles the degree of genetic relatedness between pairs of individuals is assigned as first degree (parent-offspring or siblings), second degree (half-siblings or nephew/niece – uncle/aunt or grandparent-grandchild), unrelated or identical. All the sites in the vcf files were assumed as homozygous (by only taking the majority allele for each polymorphic site) and low frequency alleles (<1%) were removed from the resulting plink files with –maf 0.01.

**NgsRelate:** We also used the program NgsRelate to infer relatedness[193]. NGSrelate is a maximum-likelihood based program that for a pair of individuals estimates the three coefficients, $k_0$, $k_1$ and $k_2$, from genotype likelihoods (GLs) instead of called genotypes and through the GLs it takes into account the uncertainty of the true genotypes, which is inherent to low-depth data. The coefficients, $k_0$, $k_1$ and $k_2$, denote the proportions of the genome where the pair of analyzed individuals share 0, 1 and 2 alleles identical by descent, respectively. Importantly, they can be used to infer relatedness, e.g. the expected values for a full sibling pair is $k_0$=0.25, $k_1$=0.5 and $k_2$=0.25, and in general, the more related a pair of individuals is the lower $k_0$ is expected to be.

Since it has not been shown that NgsRelate works for sequencing data of ultra low depth, we only included the 376 samples with sequencing depth above 0.1X in the analyses. From these we estimated GLs and allele frequencies with ANGSD[171] using the SAMtools GL model (-gl 1). For this we only used reads with MapQ<30 and bases with baseQ<20 and excluded sites where the minor allele frequency across the analysed samples was below 0.05. To minimize potential issues caused by high error rates caused by the samples being ancient, we only estimated GLs and allele frequencies for the autosomal sites where 1000 Genomes CEU population has a minor allele frequency of 0.05 and are not transitions, and we used the minor and major alleles from this CEU population as input to ANGSD (-doMajorMinor 3). Across all samples this resulted in a dataset, GLs and allele frequencies, from 1,752,719 sites.

We applied NgsRelate to the dataset and used accelerated EM algorithm to obtain maximum likelihood estimates (-m 1). As a stopping criterion for the EM algorithm, we used a log-likelihood difference of less than $10^{-6}$ between two consecutive EM-steps (-t 1e-06), and set the maximum number of steps to 10000 (-i 10000). To be able to assess whether the EM algorithm converged, we ran ten NgsRelate analyses with different starting seeds. For each pair of samples in the dataset, we used the estimates from the analysis run with the highest likelihood and for all pairs the likelihood difference among the top 5 runs were less than 0.15, suggesting that convergence was reached.

**PRIMUS:** The pedigree reconstructions based on the kinship coefficients were conducted using PRIMUS - Pedigree Reconstruction and Identification of a Maximum Unrelated Set[195].

#### Results

The results of kinship analysis of all pairs based on the NgsRelate analysis are depicted in Figure S4.1 and the estimated relatedness coefficients for pairs estimated to have $k_0<0.875$ are listed in Table S4.1. The results for first and second degree relationships obtained by READ were identical with that of NgsRelate.

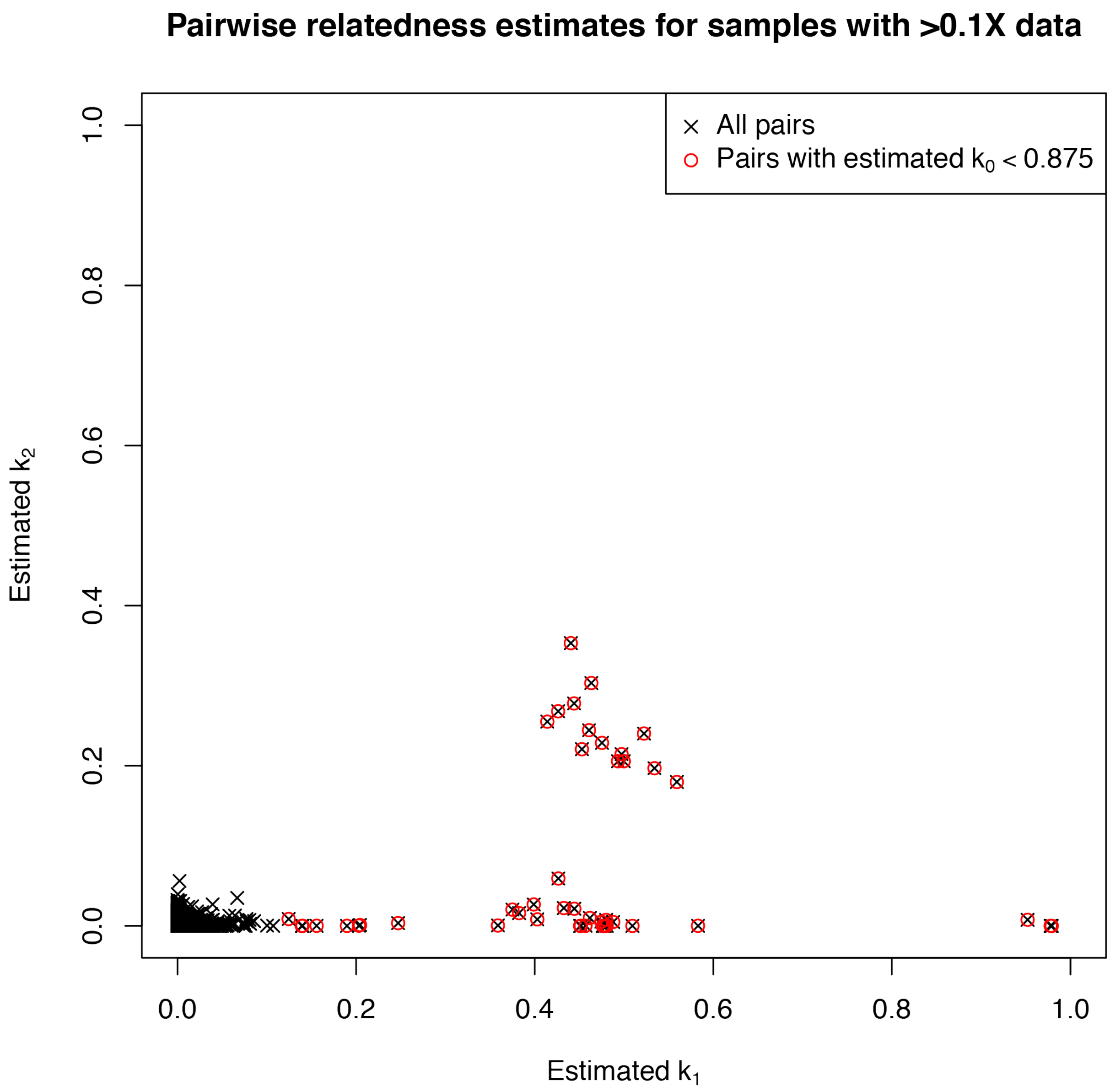


**Fig. S4.1:** Estimated relatedness coefficients for all pairs of samples with more than 0.1X data. The estimates for each pair is shown with black crosses and the ones with estimated $k_0$ below 0.875 are marked with a red circle.

These results suggest that a large number of individuals are closely related in our ancient dataset, in particular the individuals from the Faroe Islands, Iceland and Salme ship burial.

**Table S4.1:** Estimated relatedness coefficients for pairs of ancient individuals with more than 0.1X data with $k_0$ below 0.875.

| **IDs** | | **Origin** | | **Relatedness estimates** | | | |
|---|---|---|---|---|---|---|---|
| **Sample 1** | Sample 2 | Sample 1 | Sample 2 | # of sites | k0 | k1 | k2 |
| **VK46** | VK245 | Faroe Islands | Faroe Islands | 1144972 | 0.491 | 0.509 | 0.000 |
| **VK46** | VK45 | Faroe Islands | Faroe Islands | 189451 | 0.574 | 0.399 | 0.027 |
| **VK154** | VK156 | Poland | Poland | 1023563 | 0.522 | 0.478 | 0.000 |
| **VK230** | VK110 | Iceland | Iceland | 1331502 | 0.306 | 0.426 | 0.268 |
| **VK230** | VK111 | Iceland | Iceland | 1036610 | 0.021 | 0.979 | 0.000 |
| **VK279** | VK144 | Denmark_Fyn | Oxford | 250669 | 0.528 | 0.462 | 0.010 |
| **VK25** | VK236 | Faroe Islands | Faroe Islands | 1136739 | 0.296 | 0.475 | 0.229 |
| **VK25** | VK238 | Faroe Islands | Faroe Islands | 1117599 | 0.641 | 0.359 | 0.001 |
| **VK25** | VK234 | Faroe Islands | Faroe Islands | 791012 | 0.301 | 0.493 | 0.206 |
| **VK25** | VK242 | Faroe Islands | Faroe Islands | 702498 | 0.507 | 0.488 | 0.005 |
| **VK25** | VK44 | Faroe Islands | Faroe Islands | 567811 | 0.513 | 0.480 | 0.007 |
| **VK179** | VK183 | Greenland | Greenland | 640061 | 0.860 | 0.140 | 0.000 |
| **VK110** | VK111 | Iceland | Iceland | 962379 | 0.022 | 0.978 | 0.000 |
| **VK483** | VK497 | Salme | Salme | 1130328 | 0.206 | 0.440 | 0.353 |
| **VK483** | VK490 | Salme | Salme | 942642 | 0.288 | 0.497 | 0.214 |
| **VK483** | VK485 | Salme | Salme | 795096 | 0.261 | 0.559 | 0.180 |
| **VK237** | VK244 | Faroe Islands | Faroe Islands | 645056 | 0.867 | 0.124 | 0.009 |
| **VK539** | VK540 | Shestovitsa | Shestovitsa | 853398 | 0.331 | 0.414 | 0.255 |
| **VK497** | VK490 | Salme | Salme | 876836 | 0.295 | 0.500 | 0.206 |
| **VK497** | VK485 | Salme | Salme | 739597 | 0.327 | 0.453 | 0.221 |
| **VK236** | VK238 | Faroe Islands | Faroe Islands | 1034784 | 0.549 | 0.451 | 0.000 |
| **VK236** | VK234 | Faroe Islands | Faroe Islands | 734873 | 0.269 | 0.534 | 0.197 |
| **VK236** | VK242 | Faroe Islands | Faroe Islands | 655170 | 0.602 | 0.382 | 0.016 |
| **VK236** | VK44 | Faroe Islands | Faroe Islands | 529445 | 0.543 | 0.457 | 0.000 |
| **VK342** | VK527 | Öand | Uppsala | 909447 | 0.861 | 0.139 | 0.000 |
| **VK342** | VK354 | Öland | Öland | 682763 | 0.548 | 0.452 | 0.000 |

| | | | | | | | |
|---|---|---|---|---|---|---|---|
| **VK238** | VK234 | Faroe Islands | Faroe Islands | 718915 | 0.545 | 0.433 | 0.022 |
| **VK238** | VK242 | Faroe Islands | Faroe Islands | 640593 | 0.524 | 0.476 | 0.000 |
| **VK238** | VK44 | Faroe Islands | Faroe Islands | 515690 | 0.519 | 0.481 | 0.000 |
| **VK406** | VK33 | Skara | Skara | 805059 | 0.040 | 0.952 | 0.008 |
| **VK168** | VK167 | Oxford | Oxford | 746748 | 0.520 | 0.475 | 0.005 |
| **VK527** | VK517 | Uppsala | Uppsala | 816579 | 0.233 | 0.463 | 0.303 |
| **VK555** | VK490 | Salme | Salme | 770828 | 0.844 | 0.156 | 0.000 |
| **VK245** | VK240 | Faroe Islands | Faroe Islands | 636199 | 0.534 | 0.444 | 0.021 |
| **VK245** | VK45 | Faroe Islands | Faroe Islands | 132706 | 0.278 | 0.444 | 0.278 |
| **VK157** | VK155 | Poland | Poland | 248596 | 0.237 | 0.522 | 0.240 |
| **VK490** | VK485 | Salme | Salme | 619501 | 0.295 | 0.461 | 0.244 |
| **VK187** | VK183 | Greenland | Greenland | 460045 | 0.797 | 0.203 | 0.000 |
| **VK241** | VK244 | Faroe Islands | Faroe Islands | 436861 | 0.749 | 0.247 | 0.003 |
| **VK240** | VK45 | Faroe Islands | Faroe Islands | 107208 | 0.514 | 0.426 | 0.059 |
| **VK234** | VK242 | Faroe Islands | Faroe Islands | 454640 | 0.605 | 0.375 | 0.020 |
| **VK234** | VK44 | Faroe Islands | Faroe Islands | 367567 | 0.523 | 0.476 | 0.000 |
| **VK19** | VK408 | Ladoga | Ladoga | 141229 | 0.417 | 0.583 | 0.000 |
| **VK242** | VK44 | Faroe Islands | Faroe Islands | 329339 | 0.589 | 0.403 | 0.008 |

In the Salme ship burial 4 male siblings (VK483, VK485, VK490, VK497), i.e. brothers were discovered (Figure S4.2a), which was additionally supported by the identical mtDNA and Y chromosome profiles (Supplementary Table 5). It is worth mentioning here that the slight differences in Y chromosomal haplogroup assignments in this case are likely due to the presence of ancient DNA damaged sites or insufficient genomic coverage. As one would expect, the four brothers were buried relatively close to each other in the same layer of the Salme II ship burial. In Iceland, we identified a female individual (VK111) with her two children (one male VK110 and one female VK230, Figure S4.2b).

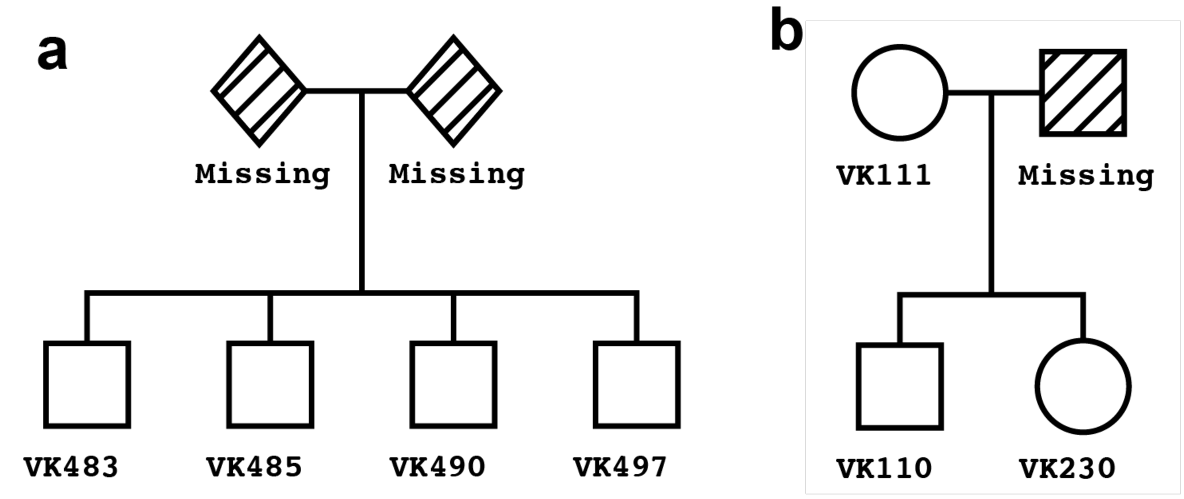


**Fig. S4.2**: Genealogy networks for individuals from Salme (a) and Iceland (b).

The largest number of related individuals were identified in Faroe Islands. Three family groups were identified in the Church burial in Faroe Islands, however, only two groups are analysed here due to the relative closeness of family members within the two groups. Even though we could not see significant degree of relationship between the groups, we cannot exclude the genetic ties between these groups due to missing individuals in our dataset.

Family-1 included 5 infants grave22 (VK238), grave31 (VK242), grave23 (VK234), grave27 (VK236) and grave28 (VK25). The possible four pedigree networks are shown in Extended Data Fig. 8.

The second family included 4 related individuals from the Faroe Islands (Extended Data Fig. 8). The most "parsimonious" network likely reflects the true genetic relationship between the individuals also considering the grave locations in the churchyard, i.e. two infants in graves 13 (VK45) and 15 (VK245) buried close to their (likely) grandparents in graves 7 (VK240) and 14 (VK46).

Among the results from the genetic relatedness analysis it is worth mentioning two exciting cases that were unexpected. We have identified a $2^{nd}$ degree related pair of male individuals (i.e. half-brothers, nephew-uncle or grandson-grandfather) from across the North Sea, one sample (VK279) excavated in Denmark (site Galgedil on Fyn) while his half-brother (VK144) in UK (site Oxford). Similarly, another pair of individuals that are estimated to be third or fourth degree relatives was found in Sweden, one of them excavated on the island of Öland (VK342) while the other one was from Uppsala (VK527).

### **Supplementary Note 5 - Imputation**

Genotype imputations were conducted for 298 ancient samples (289 from this study + 9 from the study by Krzewińska et al.[196]) that had a sequencing depth greater than 0.5X. Since these ancient samples were sequenced at low depth of coverage, we used Beagle v4.1[197] for imputations based on the genotype likelihood data, which was first estimated by GATK v3.7.0 UnifiedGenotyper. To generate the genotype data we only called biallelic sites present in the 1000G dataset and only the observed alleles (--genotyping_mode GENOTYPE_GIVEN_ALLELES). The resulting VCF files were filtered by setting genotype likelihoods to 0 for all three genotypes (e.g. hom ref, het and hom alt) for sites with potential deamination (C>T and G>A) as described by Martiniano et al.[198]. Following this, the per-individual vcfs were merged using bcftools-v1.3.1. The combined VCF were then split into 15,000 markers each and imputed separately using beagle-4.0 using the 1000G phase3 map included with beagle (*.phase3.v5a.snps.vcf.gz and plink.chr*.GRCh37.map) with input through the genotype likelihood option. Run time for imputing using beagle was approximately 280,000 core hours.

#### Supplementary Note 6 - Merge with existing panels

To assess the genetic relationship between the ancient Viking samples and other populations (both modern and ancient) we merged the whole-genome shotgun data of the Vikings with two different SNP array datasets of modern worldwide and ancient populations. Since most ancient samples from this study were low coverage, to obtain the genotypes of the ancient samples (mostly Viking age) the "samtools mpileup" command was used followed by a single read sampling of the majority allele for each of the sites present in the relevant reference dataset, with mapping and base quality ≥30.

**Scandinavian panel - SNP array of European populations enriched for Scandinavians**: A new reference panel was constructed based on 3 published datasets for which we conducted quality control (QC) analysis: the EGAD00010000632 dataset from Leslie et al.[199] (UK dataset), the EGAD00010000124 dataset from Genetic Analysis of Psoriasis Consortium & the Wellcome Trust Case Control Consortium 2[200] (IRE dataset) and the EGAD00000000120 dataset from The International Multiple Sclerosis Genetics Consortium & The Wellcome Trust Case Control Consortium 2[201] (EU dataset). The UK dataset was genotyped on the Human1-2M-DuoCustom SNPchip and contained 1,115,428 sites and 2,912 individuals. The IRE and EU datasets were genotyped on the Human670-QuadCustom SNPchip with 580,030 sites and included 2,622 and 11,376 individuals, respectively. The UK dataset consist of only UK individuals, the IRE dataset has both UK and Irish individuals and the EU dataset contains individuals from Australia, Belgium, Denmark, Finland, France, Germany, Italy, New Zealand, N. Ireland, Norway, Poland, Spain, Sweden, the UK and the US.

The datasets were converted from their original genotype (.gen) file format to binary plink files using GTOOL and plink[202], the GTOOL default cut-off of 0.9 for genotype probabilities was used, this included all autosomal + the X and Y chromosomes. The datasets included a list of both SNPs and individuals that passed QC which we used, leaving 870,170 markers and 2,578 individuals for the UK dataset, 535,475 markers and 2,178 individuals for the IRE dataset and 475,806 markers and 10,299 individuals for the EU dataset.

All the datasets were checked to be on the TOP strand as specified by the SNPchip manifest. Then the datasets were put on the plus strand using plink and strand files (detailing for each SNPchip which strand TOP refers to for each individual site, from: http://www.well.ox.ac.uk/~wrayner/strand/). Then using liftOver[203] the datasets were lifted from hg18 to hg19. Sites that deviated from Hardy-Weinberg equilibrium were filtered away, using a cutoff P-value of $1x10^{-6}$. The three datasets were then merged and the sites, were none of the alleles agreed with the hg19 reference sequence, were removed along with duplicated sites, based on genomic position. Non-autosomal sites were removed. SNPs in the MHC/HLA region on chromosome 6 were removed, due to the fact that it is hard to map to this region due to the high degree of recombination and because of observed big differences in frequencies between the datasets for markers in this region. The CHB (Han Chinese) and YRI (Yoruba) populations from the 1000 Genomes project phase 3 database were merged to this panel as outgroups.

#### Human Origins panel - Affymetrix Human Origins SNP array panel of worldwide populations

This dataset of modern individuals consists of 2180 individuals from 213 worldwide populations[204,205]. We extracted autosomal genotypes at a subset of 593,102 SNPs that were also included in the "1240K" capture panel[206].

#### Ancient panels

We constructed datasets for population genetic analyses by merging the newly sequenced Viking Age individuals as well as other previously published ancient individuals[20,159,160,186,189,190,198,206–228] with the two modern reference panels described above. Ancient individuals were represented with "pseudo-haploid" genotypes, obtained by randomly sampling an allele passing filters (mapping quality ≥ 30 and base quality ≥ 30), further requiring that it matched one of the two alleles observed in the reference panel (Supplementary Table 3). For high coverage ancient and modern individuals, we used diploid genotypes obtained using samtools / bcftools as previously described.

#### 1000 Genomes dataset

Five European populations from the 1000 Genomes project phase 3 database were used along with CHB (Han Chinese) and YRI (Yoruba) populations as outliers to assess the genome wide allele frequencies for various SNPs associated with pigmentation phenotypes and lactose intolerance. The five European groups included all the individuals from IBS (Spanish), TSI (Tuscan), CEU (Utah Residents with Northern and Western European Ancestry), GBR (British) and FIN (Finnish) datasets.

### Supplementary Note 7 - Latent ancestry modelling

Based on the ancient pseudohaploid individuals from the "HO 1240K" panel (see Supplementary Note 6) we ran ADMIXTURE[229] by thinning the dataset for linkage disequilibrium using plink with recommended settings (--indep-pairwise 50 10 0.1). This dataset contained 1169 samples for 171769 markers for the autosomal chromosomes. We did 50 replicates with different seeds for k=2 to k=20. For each replicate, we calculated the cross-validation error, with the distribution shown as boxplots in Figure S7.1. We used pong[230] to identify the best run for each K and pong was also used to identify similar components between different Ks. See Figure S7.2 for a visualization for K=2 to K=6. In Extended Data Fig. 2 we chose the 517 most relevant individuals representing 60 different populations and visualized this for K=2 to K=5.

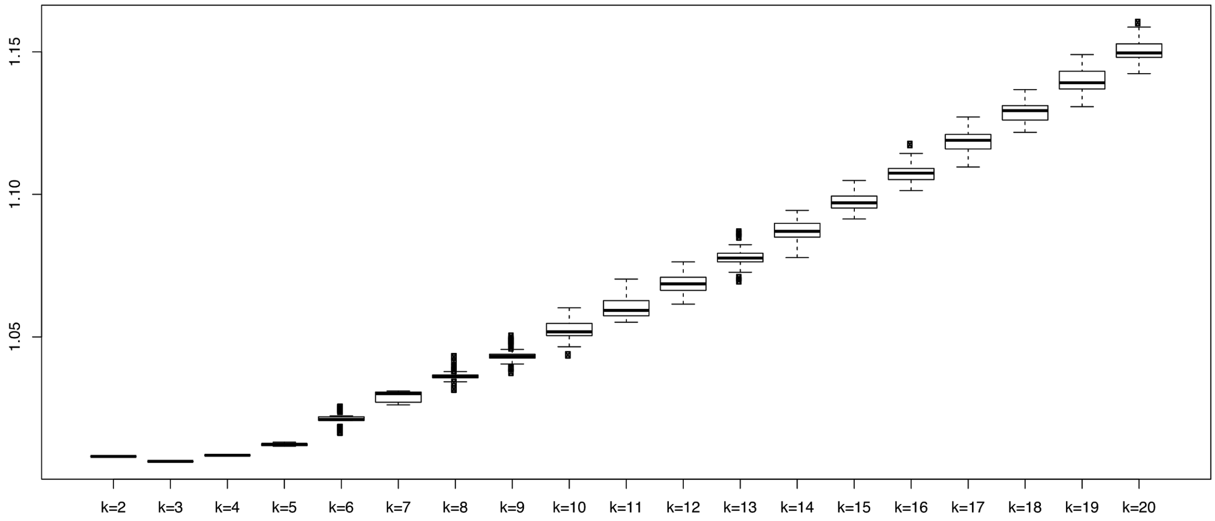


**Fig. S7.1**: Boxplots of the cross-validation error for 50 replicates of admixture runs for K=2 to K=20 with different seed values.

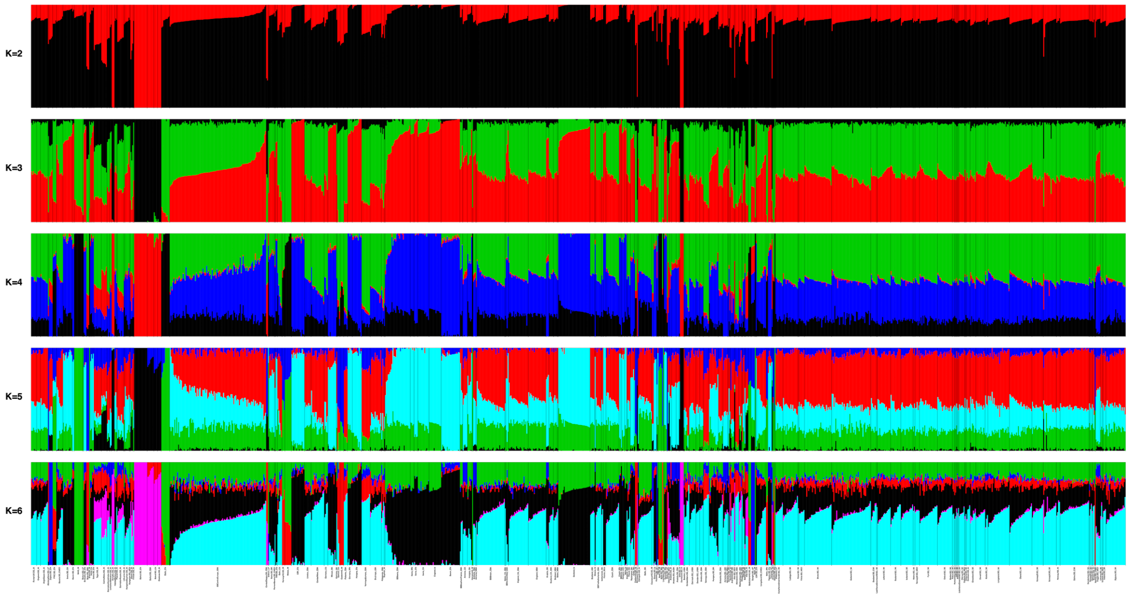


**Fig. S7.2**: Model-based clustering analysis of 1169 individuals: K=2 to K=6. This dataset includes 815 published and 354 ancient samples from this study (mostly Viking Age). See the Extended Data Fig. 2 for a subset of most relevant populations from this dataset**.**

#### Supplementary Note 8 - Genetic clustering

The large number of ancient individuals included in the analysis panels facilitates genetic clustering using the ancient individuals themselves, rather than projecting them on axes of variation inferred from modern populations. We carried this out using multi-dimensional scaling (MDS) on a distance matrix obtained from pairwise identity-by-state (IBS) sharing between individuals.

##### Batch effects

Genetic clustering using genetic similarity among the ancient individuals themselves provides a more accurate picture of their genetic differentiation than projecting them on axes of variation inferred from modern populations. However, a potential pitfall of this approach is that it is more susceptible to biases due to batch effects between sets of individuals, e.g. those with differences in sample processing. The dataset used in this study includes large numbers of individuals, compiled from heterogenous sample origins (e.g. shotgun sequencing vs in-solution capture), both of which are expected to exaggerate this potential issue.

To investigate this issue, we first assessed dimensions inferred from genetic clustering of 1,265 ancient West Eurasians using multi-dimensional scaling (MDS) on a pairwise identity-by-state (IBS) sharing matrix. While the first two dimensions differentiate samples with respect to their genetic ancestry, we observe a possible batch effect separating capture and shotgun samples along dimension 3 (Figs. S8.1 and S8.2).

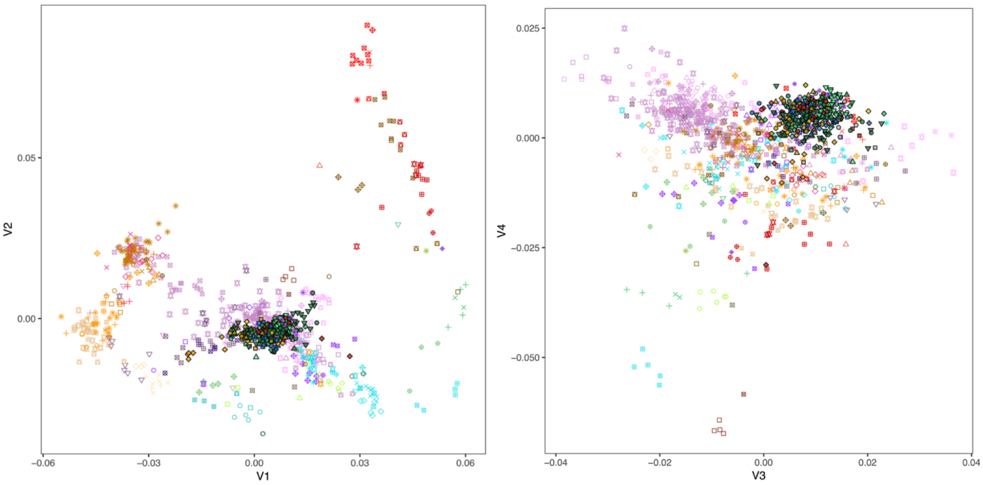


**Fig. S8.1:** MDS on 1,265 ancient West Eurasian individuals, showing dimensions 1 and 2 (left) and 3 and 4 (right). Plot symbols indicate population grouping as used throughout the study.

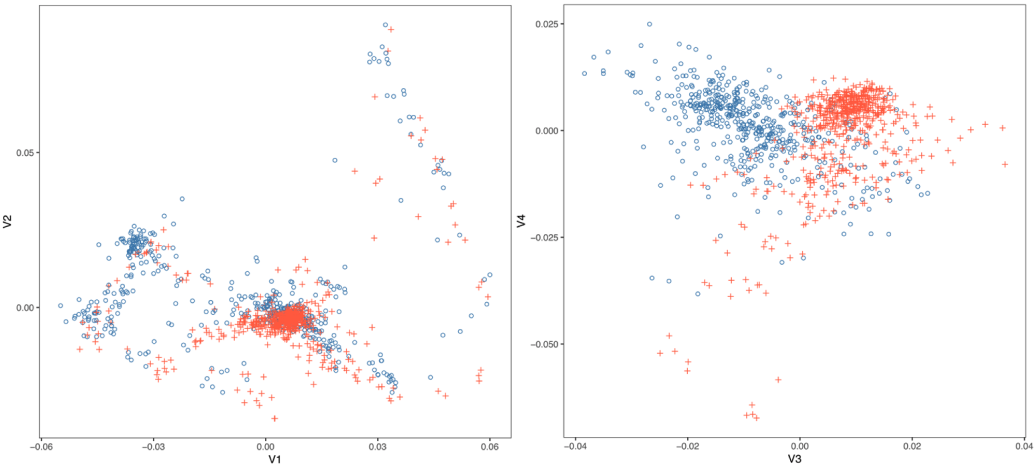


**Fig. S8.2:** MDS as in Fig. S8.2, but with plot symbols indicating samples processing using shotgun sequencing (red crosses) or in-solution capture (blue circles).

To further investigate, we included data from a total of 36 previously published individuals which were generated using both shotgun and capture approaches. We then carried out principal component analysis (PCA) on different subsets of individuals and investigated the loadings of each SNP along the inferred components, using the algorithms implemented in the GCTA package[231]. Performing

PCA on the full dataset of 1,265 individuals as well as the shotgun/capture paired individuals recovers structure that closely resembles the MDS results (Fig. S8.3).

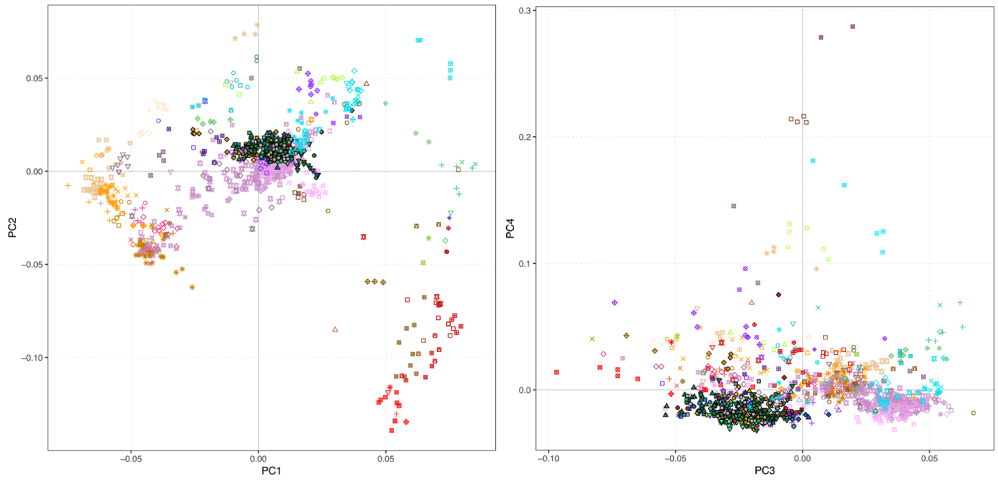


**Fig. S8.3:** PCA on 1,265 ancient West Eurasian individuals and shotgun/capture pairs, showing dimensions 1 and 2 (left) and 3 and 4 (right).

We confirm batch effects using the shotgun/capture pairs, which show clear and consistent separation along PC3 as well as a subtler effect along PC2 (Fig. S8.4).

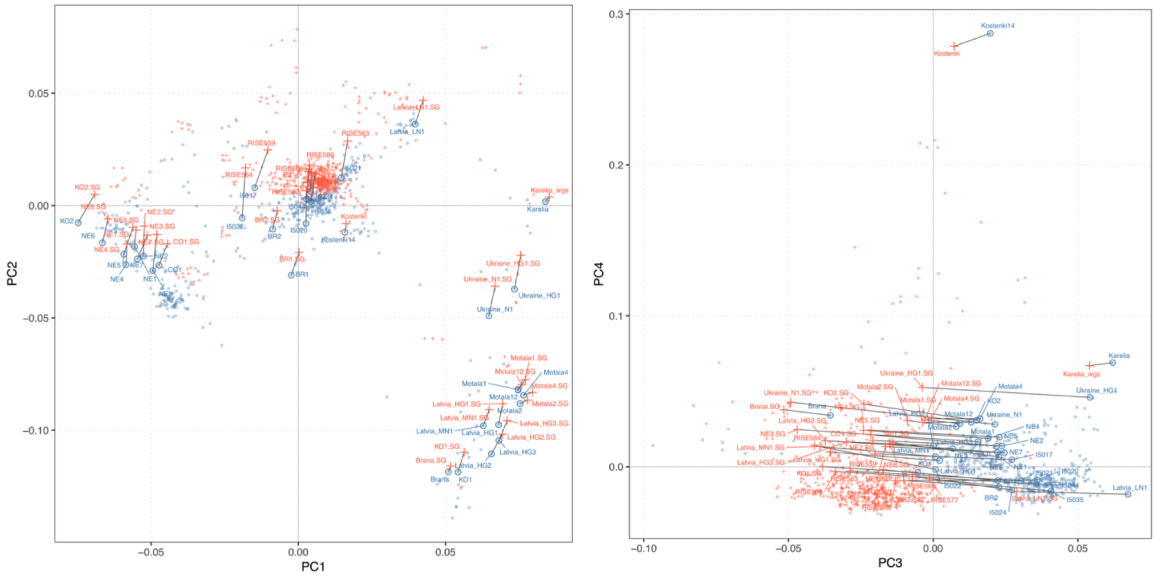


**Fig. S8.4:** PCA as in Fig. S8.4, but with plot symbols indicating sample processing using shotgun sequencing (red crosses) or in-solution capture (blue circles). Pairs of individuals with data from both approaches are joined with black lines and indicated by sample name.

However, inspection of the distribution of SNPs with high loadings along PC3 reveals a more complex pattern, with batch effects appearing to be confounded with a biological effect due to SNPs within genomic regions previously reported to show evidence for recent positive selection in West Eurasians [206] (Fig. S8.5).

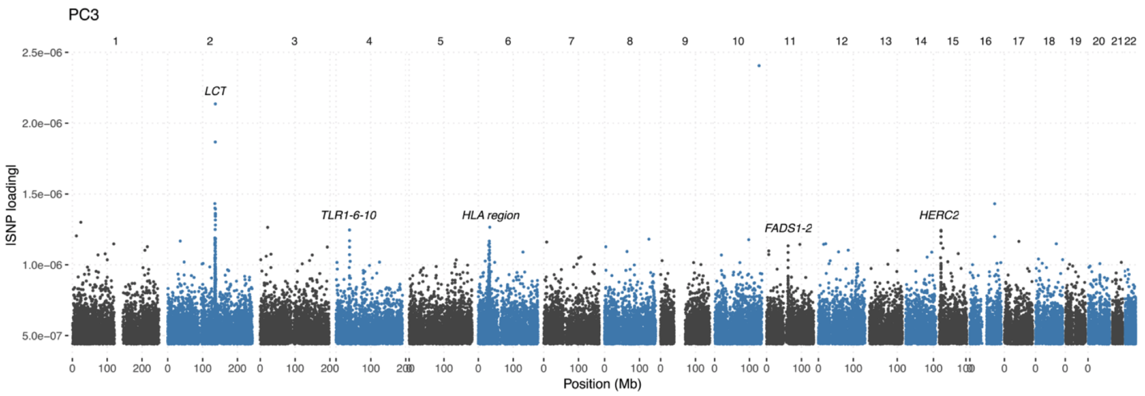


**Fig. S8.5**: Manhattan plot of SNP loadings along PC3. Peaks of high SNP loadings overlapping genes previously reported as targets of recent positive selection in Western Eurasians are indicated.

The observed confounding is not entirely surprising as the investigated ancestry and age groups are not evenly distributed across the two sample processing groups. The bulk of the shotgun data originates from Viking Age and later individuals from Northern Europe, in contrast to a large fraction of earlier Bronze Age individuals from Western and Southern Europe in the capture data. We therefore repeated the PCA analysis on a subset excluding any Iron Age or later individuals. Similar to the analysis with all individuals, we find evidence for shotgun/capture batch effects along PC3, but without strong effects of individual gene regions under selection as seen before (Fig. S8.6, S8.7)

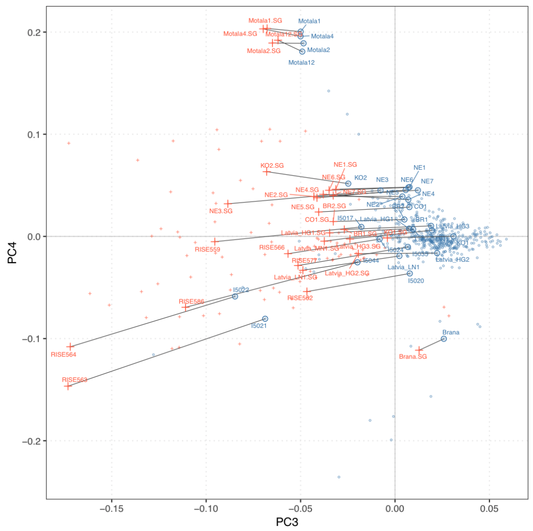


**Fig. S8.6:** PCA on a subset of pre-Iron Age individuals, with plot symbols indicating samples processing using shotgun sequencing (red crosses) or in-solution capture (blue circles). Pairs of individuals with data from both approaches are joined with black lines and indicated by sample name.

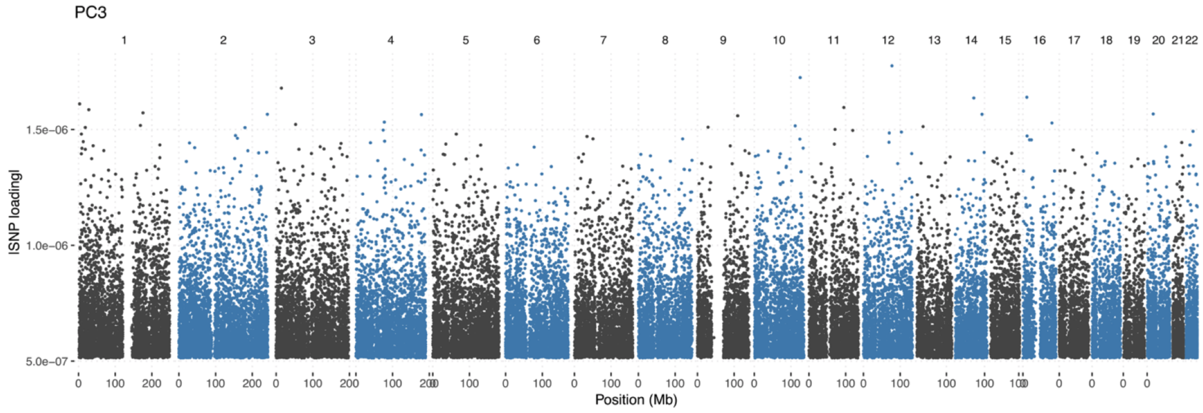


**Fig. S8.7:** Manhattan plot of SNP loadings along PC3 for subset of pre-Iron Age individuals.

Based on these results, we tested two subsets of SNPs for batch effect removal. For each of the PCA analyses above (Fig. S8.6. and Fig. S8.8), we identified the subset of SNPs with the highest loadings along PC3 (absolute value > 1 standard deviation from the mean), and removed those from the dataset. Results for PCA on the full set of individuals using the two SNP subsets are shown in Fig. S8.8.

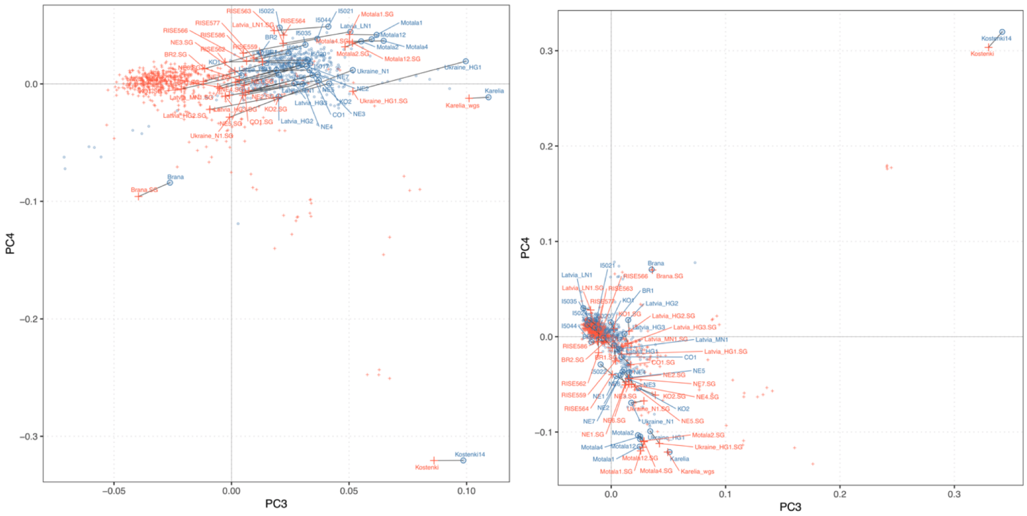


**Fig. S8.8:** Biplot of PC3 and PC4 from a PCA on all individuals, using two different subsets of SNPs for batch effect correction: SNPs with high loadings in pre-Iron Age individual dataset (left) or in the full dataset (right).

We find that batch effects are reduced but still visible along PC3 removing the SNPs identified using the pre-Iron Age individuals (Fig. S8.8 left), whereas they are removed from PC3 and PC4 when excluding the set of SNPs identified in the full set (Fig. S8.8. left). Investigating the loadings along PC3 in the pre-Iron Age corrected SNP set revealed that effects of genomic regions under selection are still present and even amplified compared to the full set of SNPs (Fig. S8.9).

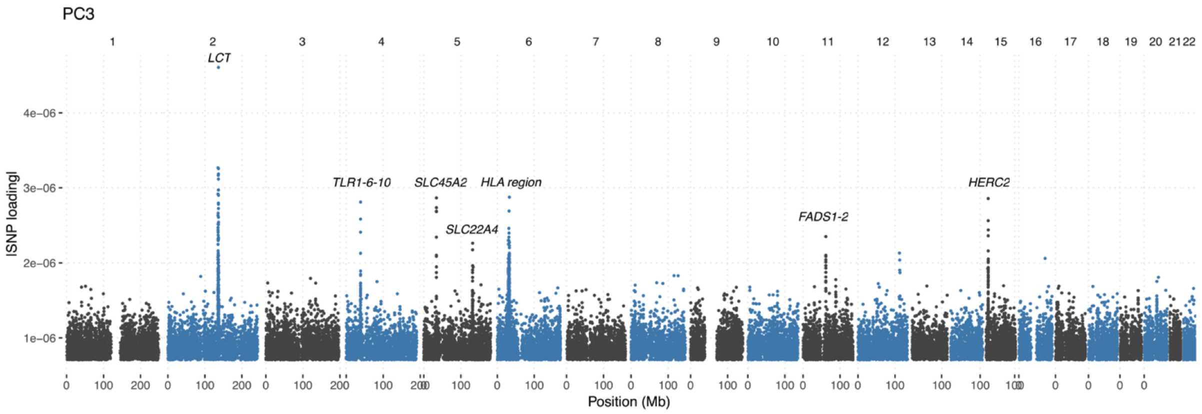


**Fig. S8.9:** Manhattan plot of SNP loadings along PC3 for full set of individuals, using batch correction by removing SNPs identified in the pre-Iron Age individuals.

In summary, we find that both differences in sample processing and genuine biological differences due to regions affected by recent positive selection impact genetic affinities between individuals in subtle and complex ways. The larger sample sizes available of Western Eurasians from the Bronze Age onwards, combined with their greater genetic homogeneity compared to earlier individuals has facilitated the detection of these previously hidden effects. As neither of the two effects is desirable in analyses of population history, we restricted all analyses of population history which use both shotgun and capture individuals to the subset of SNPs identified using the full dataset (i.e. all outliers in Figure S8.5).

#### Clustering analysis

We combined the batch-corrected MDS with further dimensionality reduction using uniform manifold approximation and projection (UMAP), an approach that has recently been shown to effectively visualize population structure among individuals across multiple scales[1]. Results of this analysis are shown in Figure 2 and Figure S8.10. The projection using IBS-UMAP (using the first 14 dimensions from the MDS) enhances local clustering of ancient individuals with a similar genetic ancestry, without loss of global relationships among the different clusters. Furthermore, genetic clusters not evident from the first two dimensions of the MDS are easily recognizable, exemplified by the clear separation of Iberian and British Neolithic individuals in IBS-UMAP (Fig 2).

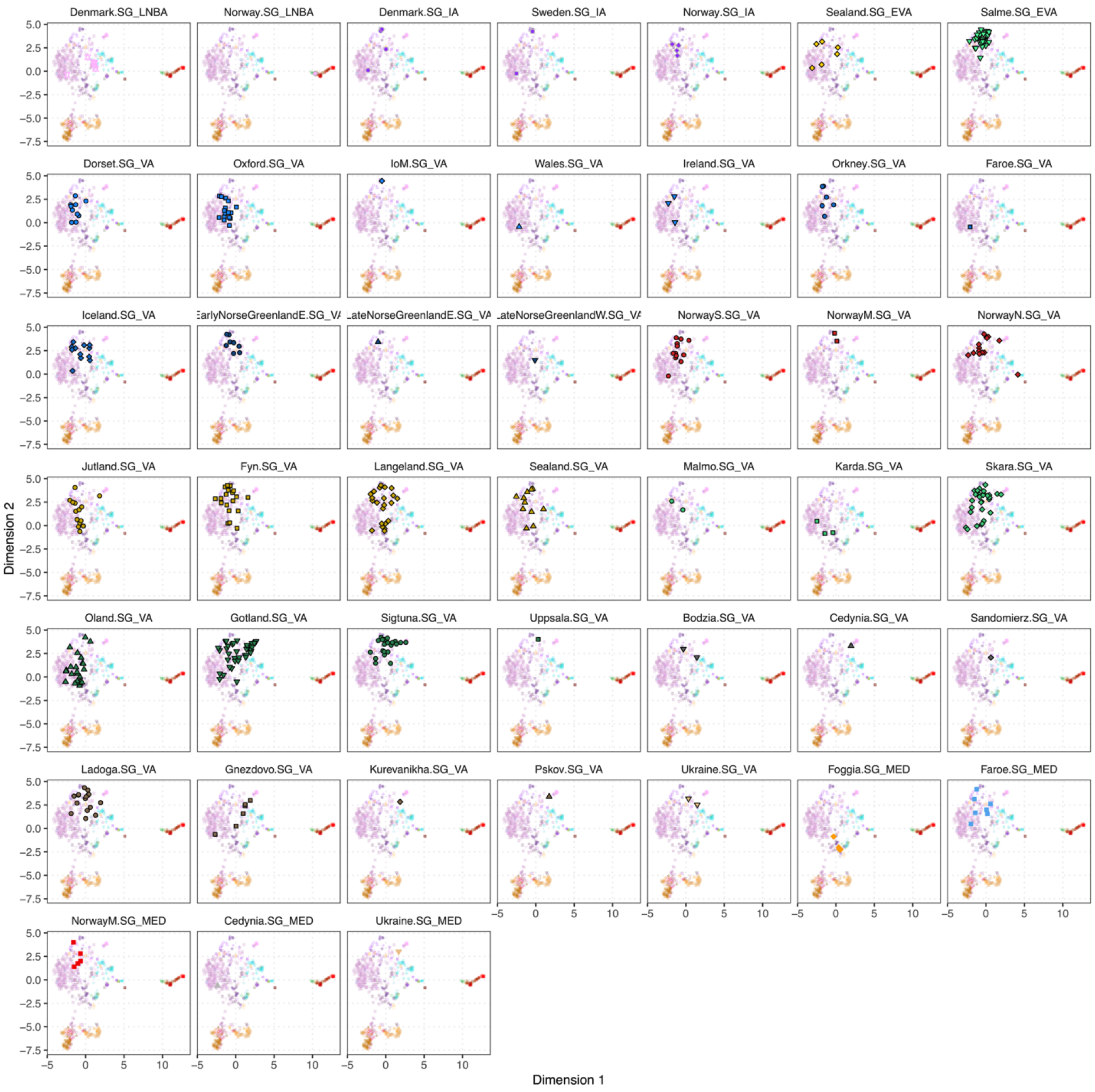


**Fig. S8.10:** Genetic clustering of study individuals using IBS-UMAP. Each panel highlights position of individuals from indicated groups over the full set of individuals.

### Supplementary Note 9 – Population genetics

#### Ancestry modelling using qpAdm

We performed estimation of ancestry proportions for ancient groups using qpAdm[4]. Each ancient group was initially modelled following previous studies of ancient Europeans, using three deep ancestral lineages: West Eurasian hunter-gatherers (represented by Loschbour); Anatolian farmers (represented by individuals from Barcin); and Steppe pastoralists (represented by individuals associated with the Yamnaya culture). Ancestry proportion estimation using qpAdm is based on $f_4$-statistics of the from $f_4$ (X,O1;O2,O3), where X is either the source or target population, and O1/O2/O3 are triplets of outgroups to the source/target groups. As such, they are susceptible to batch effects described above, e.g. if both source/target groups and the set of outgroups are composed from different sample processing schemes. To minimize batch effects and/or biases due to ancient DNA damage or SNP ascertainment, we conducted this analysis on a dataset restricted to shotgun sequenced individuals, using 1,485,845 transversion-only sites that were found polymorphic and with a minor allele count ≥ 5 in an outgroup population (YRI) in the 1000 Genomes Project. For all analyses we used a set of nine outgroups

KwazuluNatal.SG_N
Ust.SG_UP
Yana.SG_UP
Sunghir.SG_UP
Bichon.SG_LP
Zagros.SG_EN
DevilsCave.SG_N
Baikal.SG_EN
Alaska.SG_LP

Results of these analyses are shown in Extended Data Fig. 6 and Supplementary Table 7. While the three source groups (Loschbour.SG_M, Barcin.SG_EN, Yamnaya.SG_EBA) provide a good fit for most ancient Europeans, a number of them reject the fit at $p < 0.01$. We attempted to further improve the fit of those groups by testing each of the following additional putative source groups in turn in a four-way model.

CaucasusHG.SG_M
Armenia.SG_MLBA
Anatolia.SG_MLBA
Botai.SG_EBA
XiongNu.SG_IA
SarmatianRussia.SG_IA
Saami.SG_IA
Tagar.SG_IA

All target groups except Oland.SG_VA were successfully modelled this way ($p \geq 0.01$), and can be broadly grouped into two patterns. For northern and eastern Viking Age groups (NorwayN.SG_VA, Salme.SG_VA, Sigtuna.SG_VA, Ladoga.SG_VA), source groups with east Eurasian ancestry (XiongNu.SG_IA, Tagar.SG_IA, Saami.SG_IA, SarmatianRussia.SG_IA) provide good model fits, with estimated ancestry proportions reflecting the overall east Eurasian ancestry proportions. We note that among the fitting source groups are three Iron Age individuals with Saami ancestry from Levänluhta in Finland (Saami.SG_IA), which have recently documented the extension of Saami acnetry further south than the present-day distribution[232,233]. While the present analysis does not allow the distinction of the source groups, the fact that we find individuals with Saami-related ancestry among the Norwegian Viking Age groups suggests gene flow with Saami groups as a likely contributor to these signals. A different pattern is found for Karda.SG_VA and Foggia.SG_MED, which can only be fit as models including eastern source groups rich in Caucasus-related ancestry (CaucasusHG.SG_M, Armenia.SG_MLBA, Anatolia.SG_MLBA). Similar ancestry compositions have been previously documented in Greek Bronze Age individuals[220] and contemporary southern Italians[234]. However, the presence of such ancestry in Kärda in Southern Sweden is more surprising, suggesting either descendants of a remnant group from an earlier expansion of southeastern groups (e.g. during the migration period) or ongoing contacts during the Viking Age.

#### Baltic ancestry in Gotland

Genetic clustering using IBS-UMAP suggested genetic affinities of some Viking Age individuals with Bronze Age individuals from the Baltic. To further test these, we quantified excess allele sharing of Viking Age individuals with Baltic BA compared to early Viking Age individuals from Salme using $f_4$ statistics. We find that many individuals from the island of Gotland share a significant excess of alleles with Baltic BA (Extended Data Fig. 4), consistent with other evidence of this site being a trading post with contacts across the Baltic Sea.

#### Genetic diversity

We quantified genetic diversity for ancient groups using "conditional nucleotide diversity" as previously described[5]. Briefly, we selected SNPs that were polymorphic in an outgroup population (YRI from the 1000 Genomes Project) and with a minor allele count >= 5, and calculated average pairwise differences between individuals. Results of this analysis are shown in Fig. S9.1.

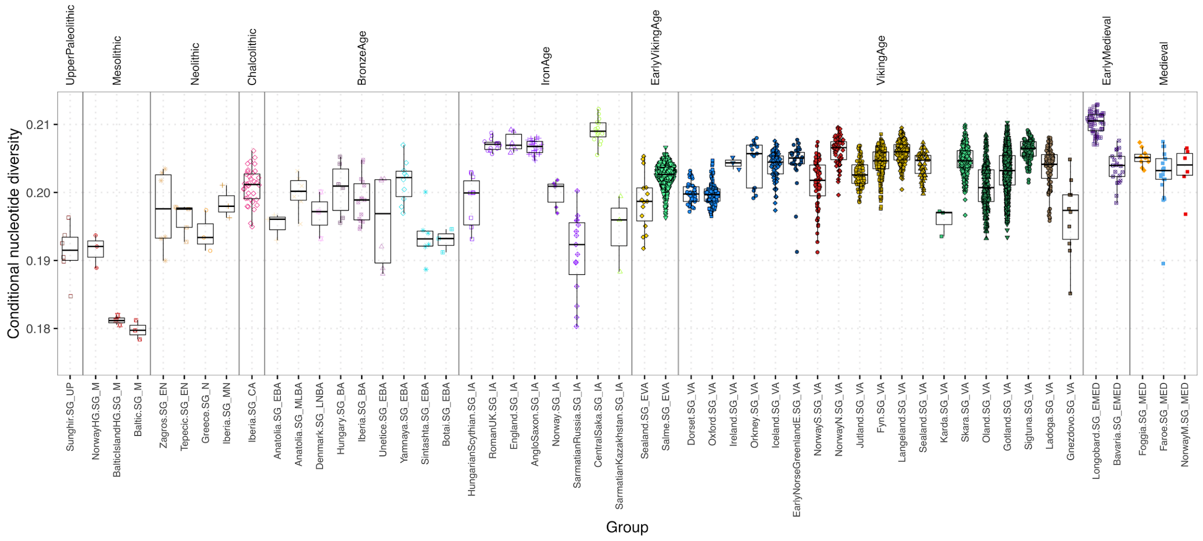


**Fig. S9.1:** Conditional nucleotide diversity of various VA groups from this study and comparative ancient populations.

#### Supplementary Note 10 – Identity-by-descent

We inferred genomic segments shared via identity-by-descent (IBD) within the context of a reference panel of 1,464 present-day Europeans, using IBDseq. Genetic clustering by MDS on a distance matrix obtained from pairwise IBD sharing and UMAP revealed fine-scale population structure among Viking Age individuals invisible from allele-frequency-based IBS analyses (Figs. 3 and S10.1).

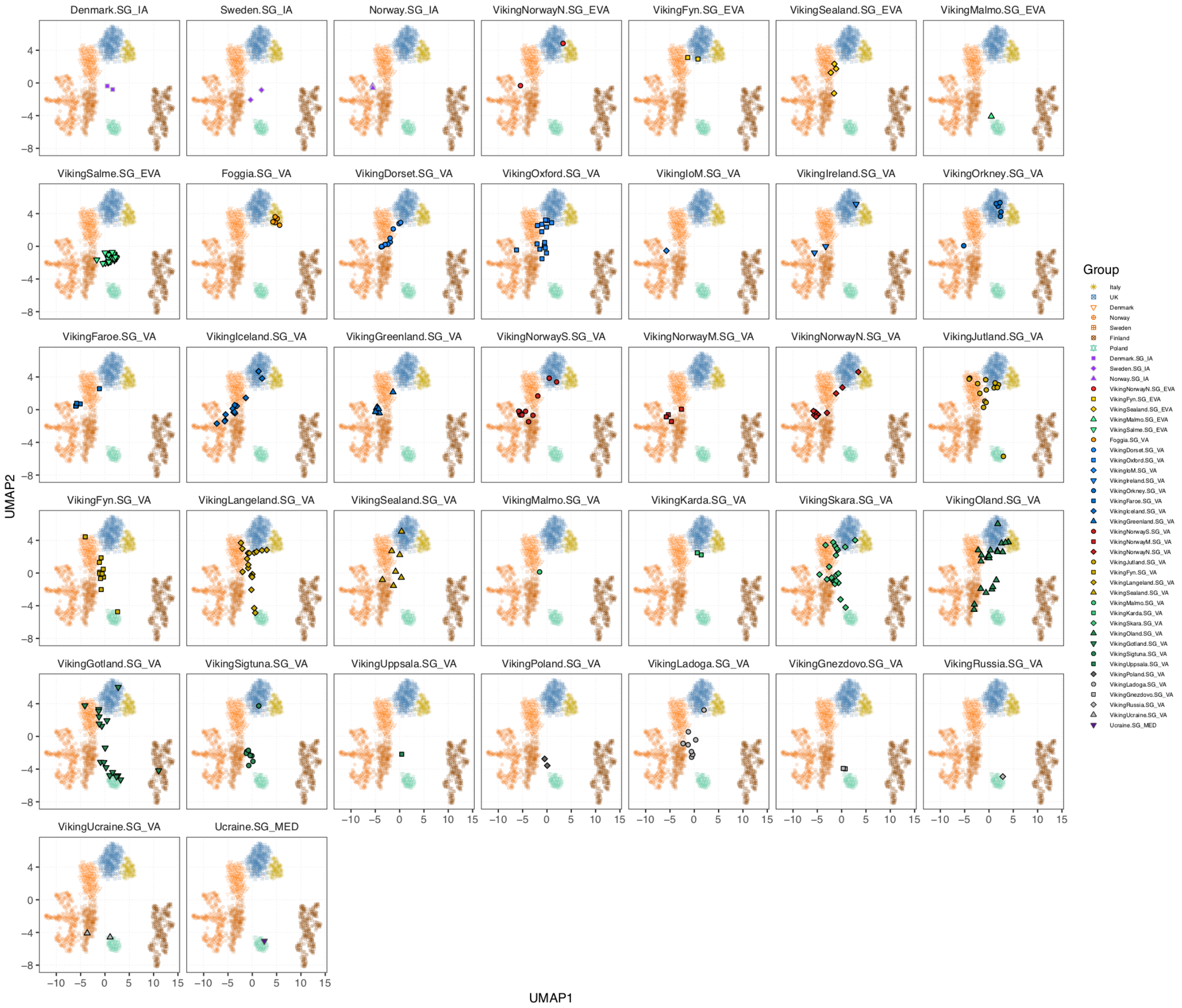


**Fig. S10.1**: Genetic clustering of imputed Viking Age individuals using IBD-UMAP.

To investigate signatures of inbreeding we analyzed distributions of genomic segments homozygous-by-descent (HBD). We find overall low levels of homozygosity among Viking Age individuals, including those from more remote locations like Greenland (Figs. S10.2 and S10.3).

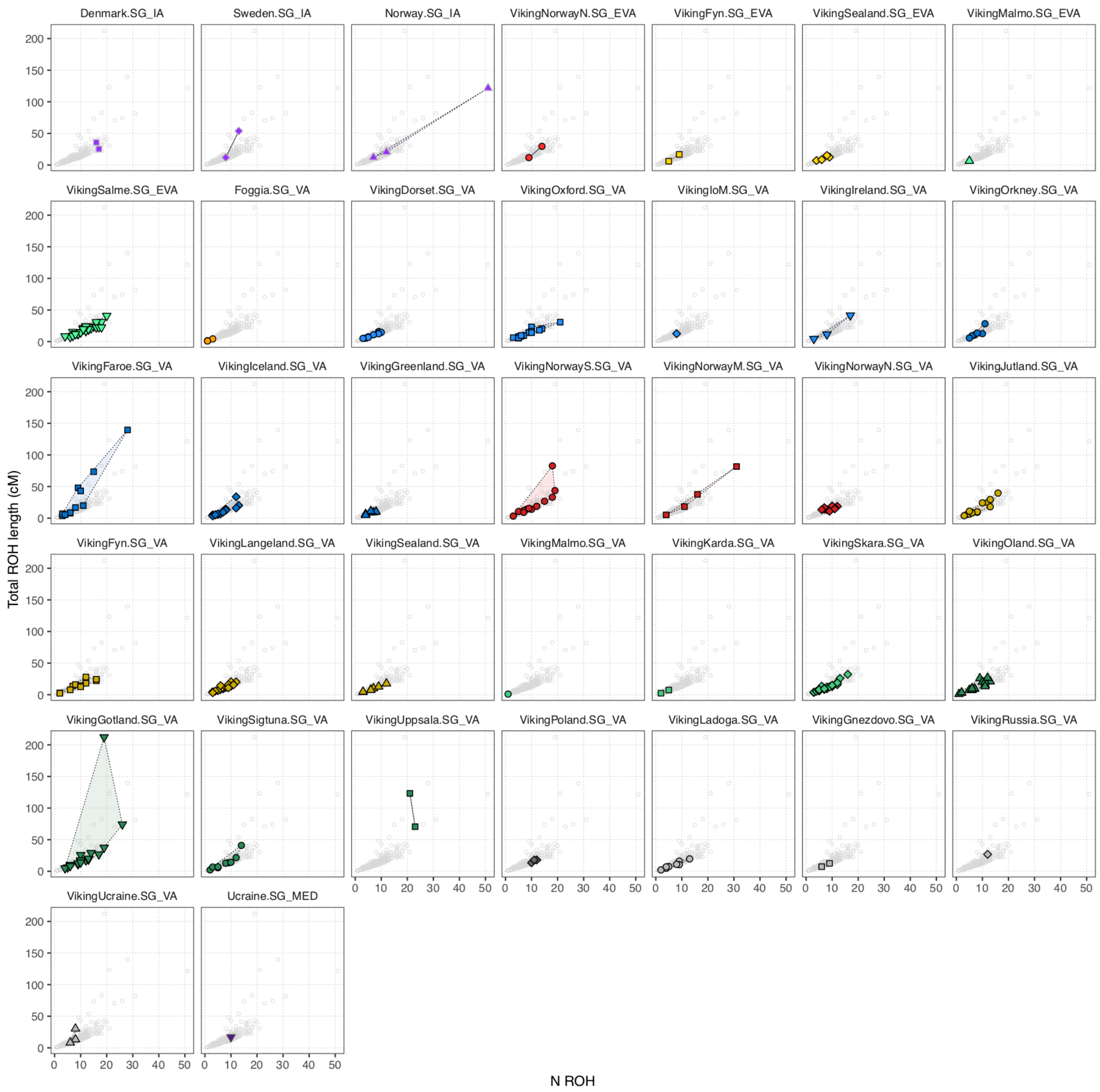


**Fig. S10.2**: Number and total length of genomic segments homozygous-by-descent in Viking Age individuals.

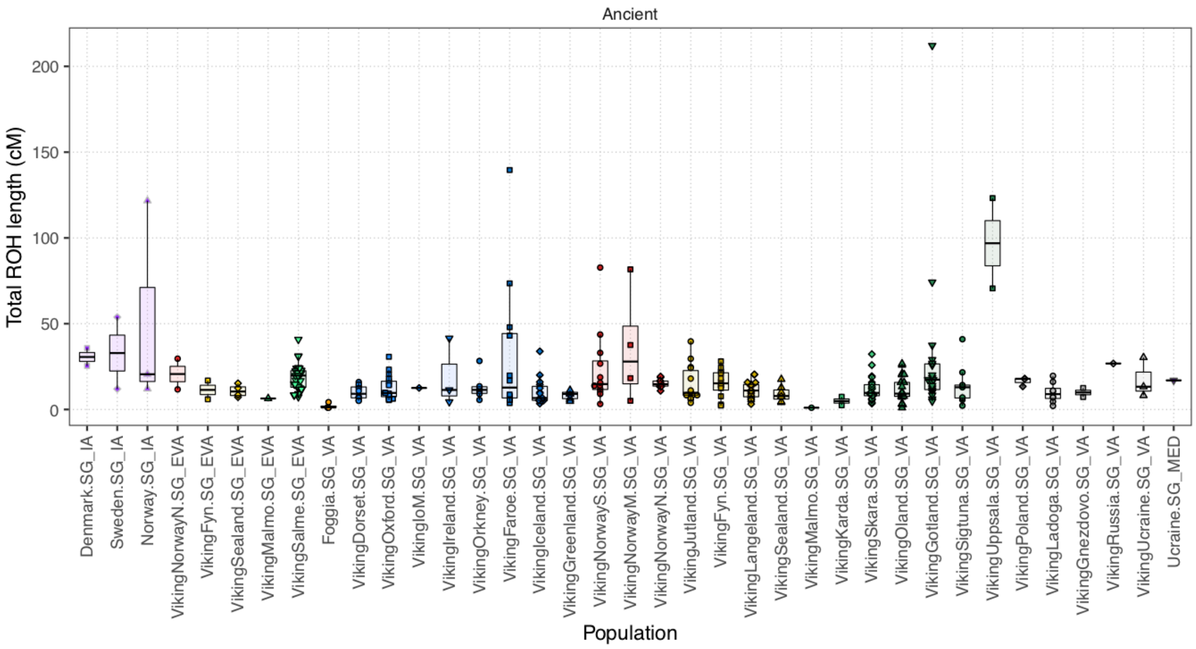


**Fig. S10.3**: Distributions of total length of genomic segments homozygous-by-descent in Viking Age individuals.

### Supplementary Note 11 - Chromosome Painting

When examining the sequence data for the 319 ancient Viking genomes sequenced to 1X or higher, we found that standard analyses were inadequate in extracting population structure within Scandinavia. We therefore used a fine-scale method called Chromosome Painting[235] to extract maximum signal from the ancient DNA.

To understand our analysis, it is important to define the terminology:

- "Recipient": an individual whose genetics we wish to describe.
- "Donor individuals": a set of individuals who we will "paint" the recipient from, that together form a "Panel".
- "Paint": Use a model, ChromoPainter [235], to estimate the total amount of genome that each donor is the closest relative to the recipient in the panel.
- "Palette": a grouping of the donors into clusters, to obtain the amount of genome that the recipient has a closest relative from each donor population.
- "Surrogate population": a population that we will use to represent a named population of interest.

Briefly, we will use ancient DNA to create a set of K high-quality surrogate populations that are compared to a palette of M donor populations, with M>K to allow us to retain as many individuals as possible. We will then be able to describe each modern or ancient individual by painting them against the donor populations (to form a palette vector of length M) and then describing them as a mixture of the K surrogate populations' palettes. This procedure is a refinement of that used by the GLOBETROTTER method [236].

This approach has the advantage that it is robust to batch effects of modern and ancient samples, because we focus on comparing all individuals to only one panel. The chromosome painting method contains an error rate parameter that is unique to each sample. Therefore, batch effects will be treated as noise, which will be higher in the non-reference batch, and should not produce systematic bias.

#### Chromosome Painting Procedure:

1. **Unsupervised Ancient Sample Analysis.** We first tried ChromoPainter and FineSTRUCTURE [235] in its default, unsupervised approach. However, unsupervised, there

was not clear population structure in this data for the clustering method to identify strong structure (Figure S12.1).

2. **Create Modern Reference.** Create a modern reference panel using 1873 modern individuals sampled from Northern Europe, using the standard FineSTRUCTURE pipeline:
    a. Apply ChromoPainter to paint all modern individuals using the remaining individuals as donors;
    b. Cluster with FineSTRUCTURE;
    c. Assign geographical meaning to the clusters using the known labels;
    d. Call the resulting clustering the "Modern Reference Panel", which consists of 23 Modern Surrogate populations and 23 Modern Donor populations.
3. **Create Ancient Reference**. Create an "ancient reference panel" using the modern reference panel:
    a. Apply ChromoPainter to paint all ancient individuals using the "Modern Population Palette";
    b. Create a supervised "Ancient Population Palette" consisting of populations that: A: "represent" a modern ancestry direction, or B: are "best associated with" a modern ancestry direction, using an iterated Mixture Model;
    c. Create an "Ancient Population Surrogate" for each modern population, consisting of the individuals that "represent" each modern population. For K=7 modern populations, this results in a matrix of K=7 rows (surrogate populations) and 2K=14 columns (donor palette populations) which captures the ancient population structure.
4. **Infer Ancestry.** Learn about population structure in either modern individuals or ancient individuals by painting them with respect to the "ancient population panel" and fitting them as a mixture using the "ancient population surrogates".

After the panels are constructed, there are additional steps to be performed:

5. **Check Ancient Reference**. We perform a range of checks and sensitivity analyses to ensure that the inference procedure performs as expected.
6. **Principal Components Analysis.** As a sensitivity and triangulation exercise, we also consider the PCA of the ancient population panel as well as an all-vs-all ChromoPainter analysis including modern and ancient populations.
7. **Interpretation**. We then interpret the ancestry results.

#### Ancient Sample Analysis

We used fs2.0.8 (www.paintmychromosomes.com) using the protocol described with the software [235]) to paint the 319 individuals whose ancient DNA will form the "ancient sample",. Each is painted using all other ancient samples to create the "coancestry matrix", which is the number of independent segments or "chunks" or genome for which individual j (columns) is the closest match to individual I (rows). This allowed us to identify related individuals who share more and longer chunks than do unrelated individuals; after removal of one of each related pair we were left with 255 unrelated ancient individuals who will be used to create the "ancient panel". The coancestry matrix is then used by FineSTRUCTURE to perform unsupervised clustering, as shown in Figure S11.1.

Whilst there is population structure visible in these results, it is insufficient to clearly define useful population labels. This is because a) there are relatively few ancient samples, b) the ancient populations come from very similar populations, and c) many individuals are not in fact representatives of ancient population groups.

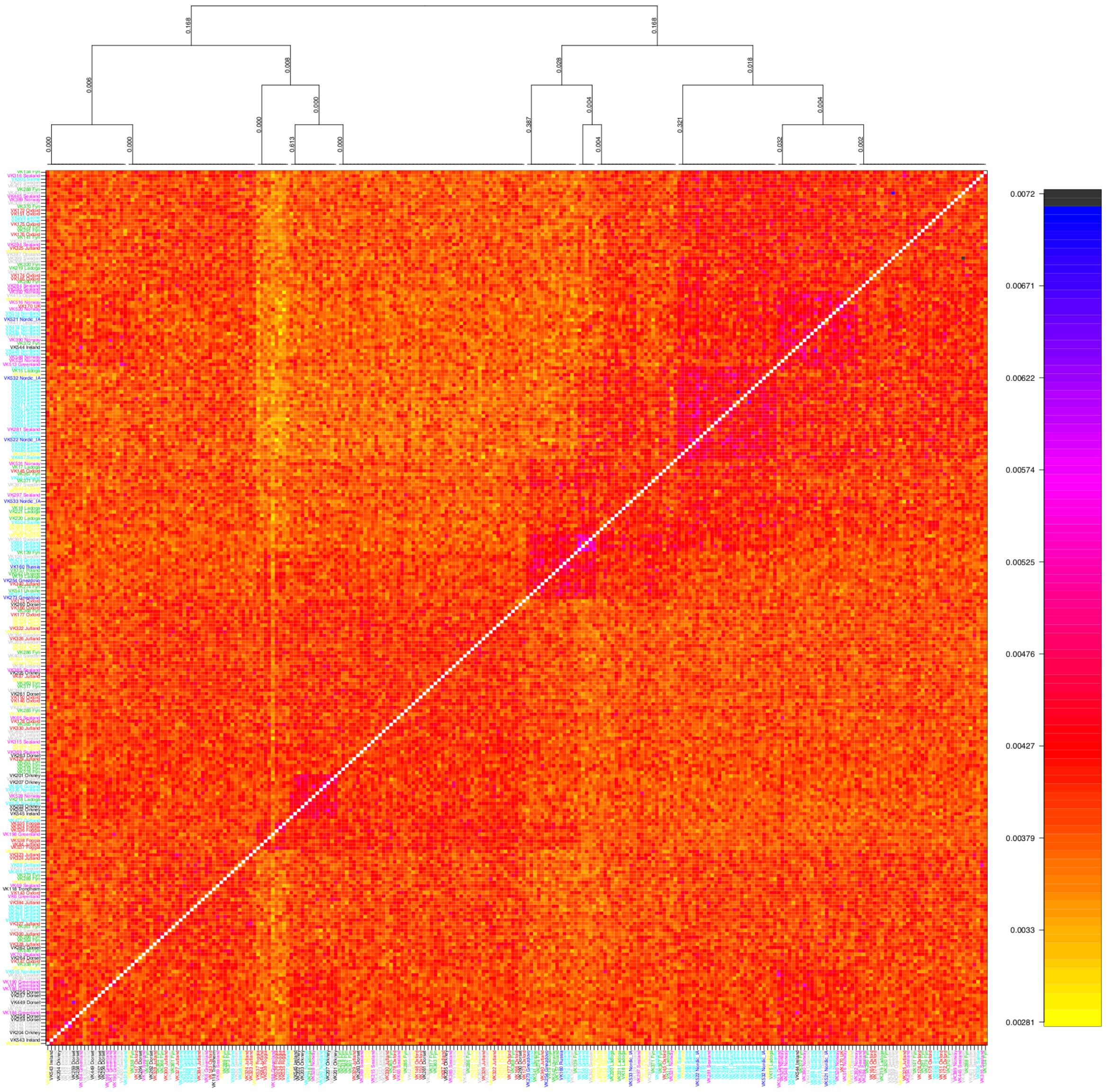


**Fig. S11.1**: Unsupervised ChromoPainter and FineSTRUCTURE results for 255 unrelated ancient individuals. The tree shows the clustering and relationship between clusters as inferred by FineSTRUCTURE, whilst the heatmap shows the Coancestry Matrix output by ChromoPainter.

#### Modern References

We used fs2.0.8 (www.paintmychromosomes.com) using the protocol described with the software [235] to paint 1675 modern individuals primarily from across Europe (UK, Italy, Poland, Denmark, Sweden, Finland, Norway, as well as China and Africa) who together form the "modern sample". FineSTRUCTURE identified 40 populations which after removal of small populations and merging of the Chinese (CHB) and African (YRI) sub-populations created 23 modern populations consisting of the 1554 unrelated individuals who could be associated with a modern population label. These together will form the "modern reference panel". Painting against this "Modern Population Palette" is done by simply summing the contribution from each individual in each donor population.

These results are shown in Figure S11.2. The clustering is strong and perfectly stratified by population label. Each population is characterized by receiving higher ancestry from its own population, implying that each represents a unique aspect of genetic drift.

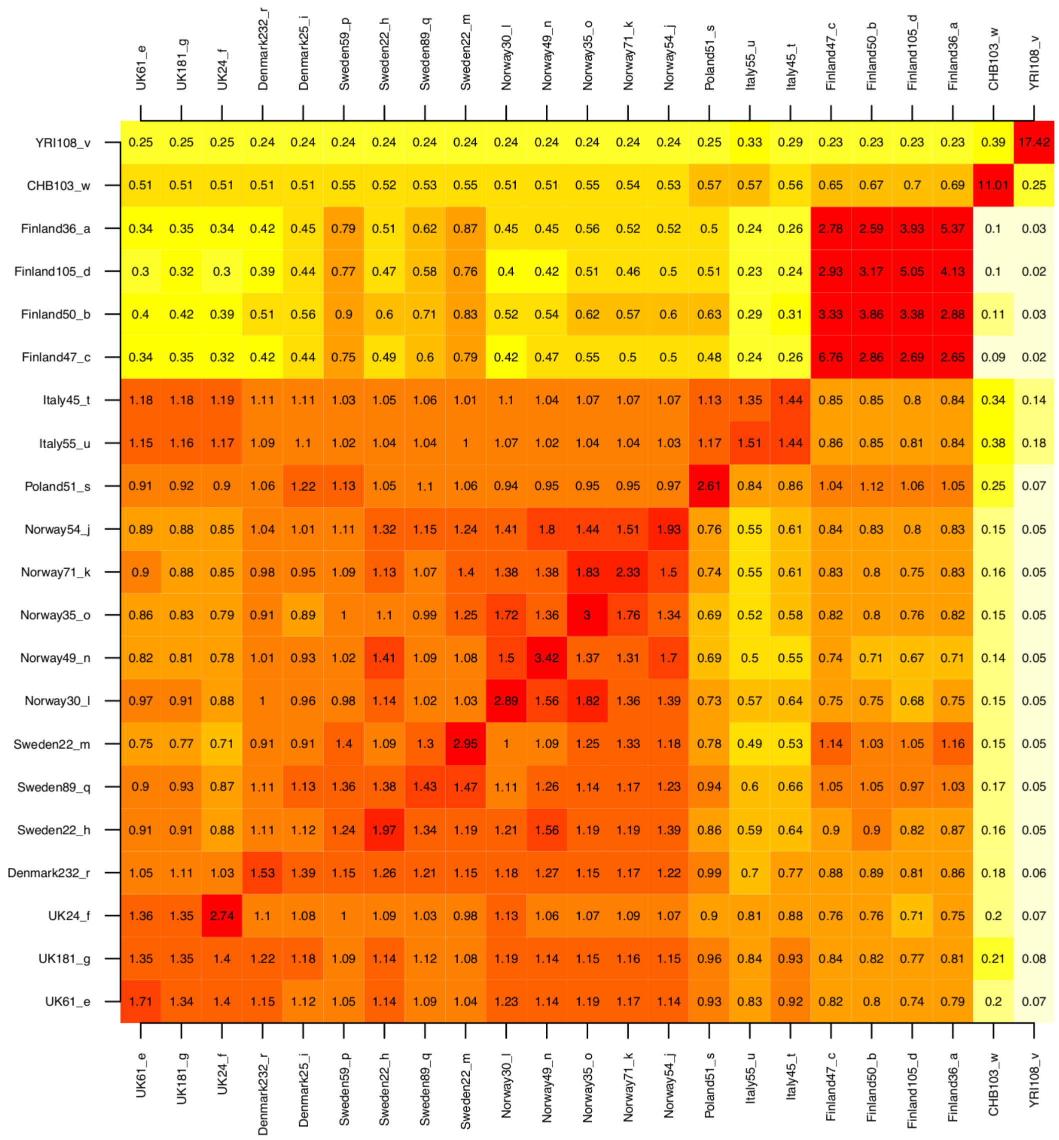


**Fig. S11.2**: Unsupervised ChromoPainter and FineSTRUCTURE results for 1554 unrelated modern individuals, where all donor individuals in the same population (i.e. columns) have been summed to create a "population donor palette", which is averaged over all individuals in the same "recipient population" (i.e. rows). The tree shows the relationship between the populations as inferred by FineSTRUCTURE.

#### Ancient References

This stage is more involved. We first:

**a. Apply ChromoPainter to paint all ancient individuals using the "Modern Population Palette".**

We now use ChromoPainter v2 to paint each ancient sample against each modern sample. Unlike above, where fs2 automatically learns parameters (which measures recombination rate and which measures error rate), we have to manually learn these using ChromoPainter v2's Expectation-Maximisation procedure, as performed in the GLOBETROTTER method [236]. We then rerun ChromoPainter v2 with these parameters fixed for all individuals. We obtain a painting for each ancient sample in the ancient panel, described as a K=23 long palette as shown in Figure S11.3.

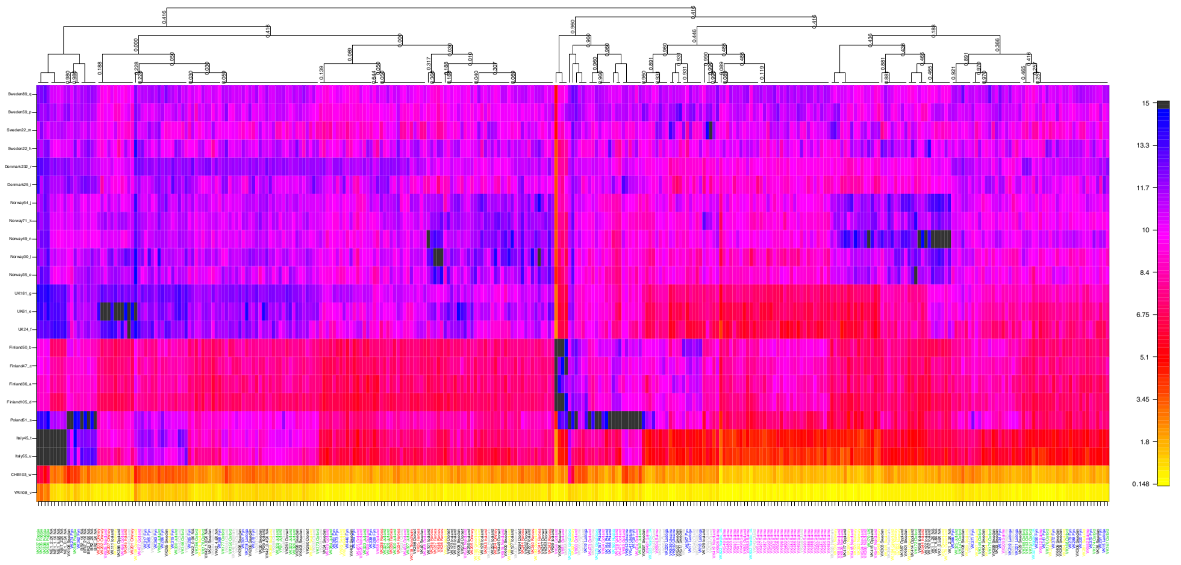

**Fig. S11.3**: Ancient panel painted against the Modern Population Palette. The palette is ordered as in Figure S11.2, whilst the ancient individuals are ordered according to the FineSTRUCTURE clustering.

There are three advantages of using the modern palette, i.e. using Figure S11.3 over Figure S11.1:

1. Because the donor populations are much larger, there is reduced statistical noise in the painting.

2. The donor populations are readily interpretable.
3. The ancient individuals naturally separate into populations that share a lot of drift with specific modern populations – i.e. they are "representative" of them, and those that don't.

Further – and perhaps surprisingly – most modern European populations (at the level of the 7 labels, not the 23 inferred populations) looks to be effectively represented.

In terms of difficult to assign populations, Denmark is one. The individuals which best match a Denmark population also well-match the "UK181_g" population. The UK populations contain individuals from the PoBI study and so we can confirm that these are English and hence contains a high proportion of Anglo-Saxon ancestry. The "UK61_e" population which is well-matched by the ancient Orkney individuals contains individuals from Scotland and Northern Ireland, whilst "UK_24f" contains individuals from Wales. Another difficult population is Sweden, for which ancient individuals often well-associate with significant Finland or Norwegian ancestry.

We exploit these labels by creating an initial labelling of the ancient populations. To do this we normalise the ancient-vs-modern coancestry by a) calculating the amount of ancestry received per donor individual from the 7 labelled countries; b) normalising each donor label to have mean 1. Figure S11.4 shows how this normalisation changes the matrix. This normalisation is chosen to move from a representation that asks "which populations are important for an individual" to one that asks "which individuals are important for a population"?

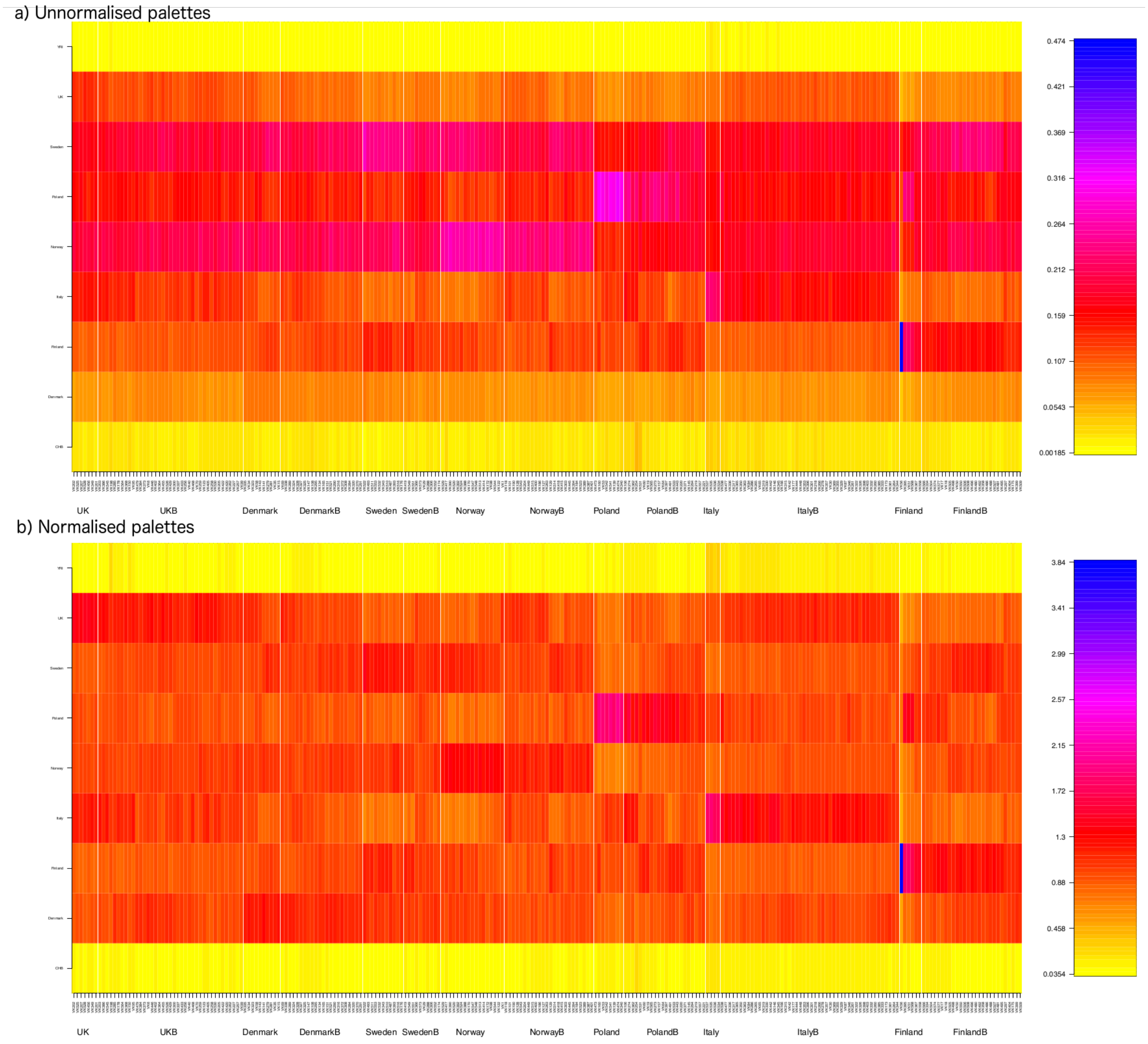


**Fig. S11.4:** a) the ancient individuals represented by their palettes, in unnormalised form, for which each column represents the importance of each population to a specific individual. b) the normalised coancestry matrix on which selection of individuals is performed, for which each row represents the importance that each individual has to a specific population.

For each of the 7 European labels, we then use the following criteria to select "potential representative individuals":

a. Individuals score population X more than any other population in the normalised coancestry;
b. Additionally, they are in the top $x_k$ for population $k$, where $x$ is chosen to represent the first change-point in the density of scores (Figure S11.5).

The remaining individuals are then assigned to their best-matching population as "potential non-representative donors".

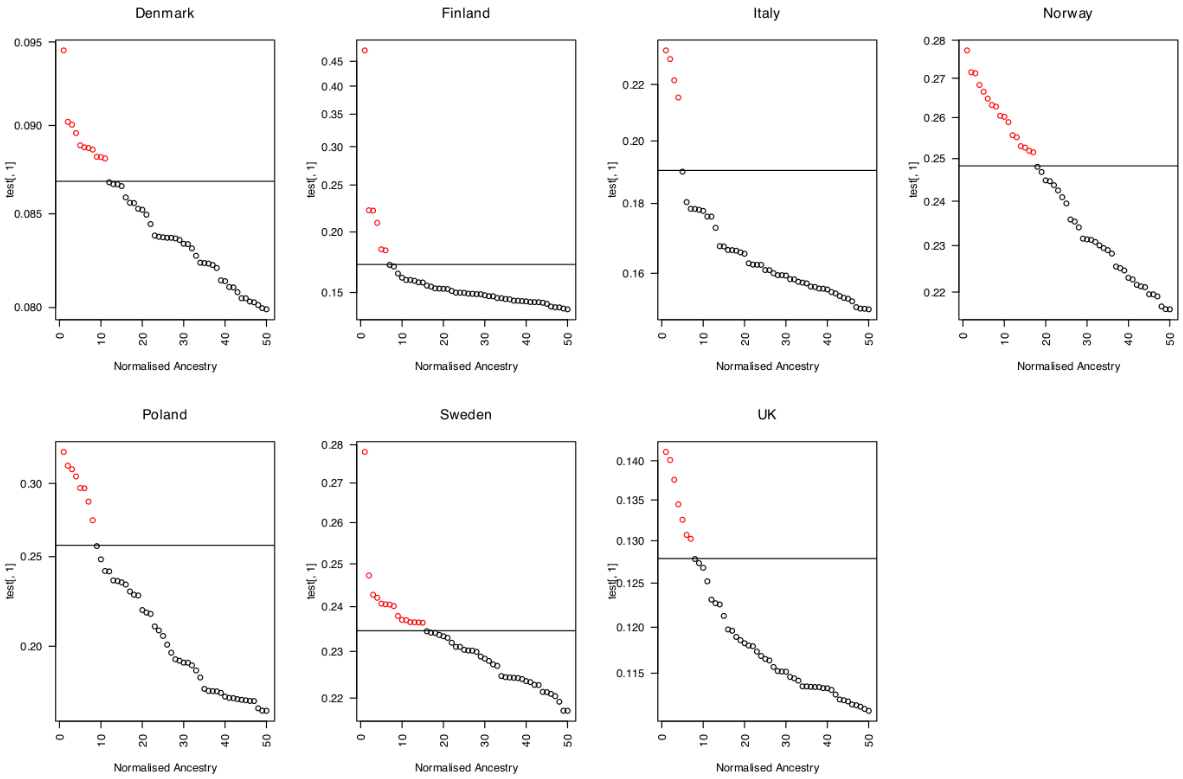


**Fig. S11.5**: Thresholds for defining the most important individuals for each modern population. All individuals above the line are chosen as representative, whilst those below may be assigned to their best scoring population as "non-representative donors".

#### Ancient Population Palette

We are now in the position of having a well-chosen set of populations. The next task is to create an ancient population palette that can be used as a reference. This stage follows the procedure used in GLOBETROTTER [236].

First we repaint each ancient individual using the 14 donor populations, this time "leaving out" themselves as donors from their own population, and one random individual from each of the other populations [236].. This is done three times: once to learn the painting parameters $N_e$ and $\mu$, a second time to learn a genome-wide prior for the donor palette, conditional on the parameters; and a final time using a shared $N_e$, $\mu$, and an individual-specific donor-prior, to obtain a high-quality palette. For each of the 7 surrogate populations (defined by the "representative individuals" above) we learn the average amount of genome received from each donor individual in each donor population. This K by

M=2K matrix is the "ancient population palette" for the ancient population panel. The results from this are shown in Figure S11.6, for both the ancient panel and the modern panel.

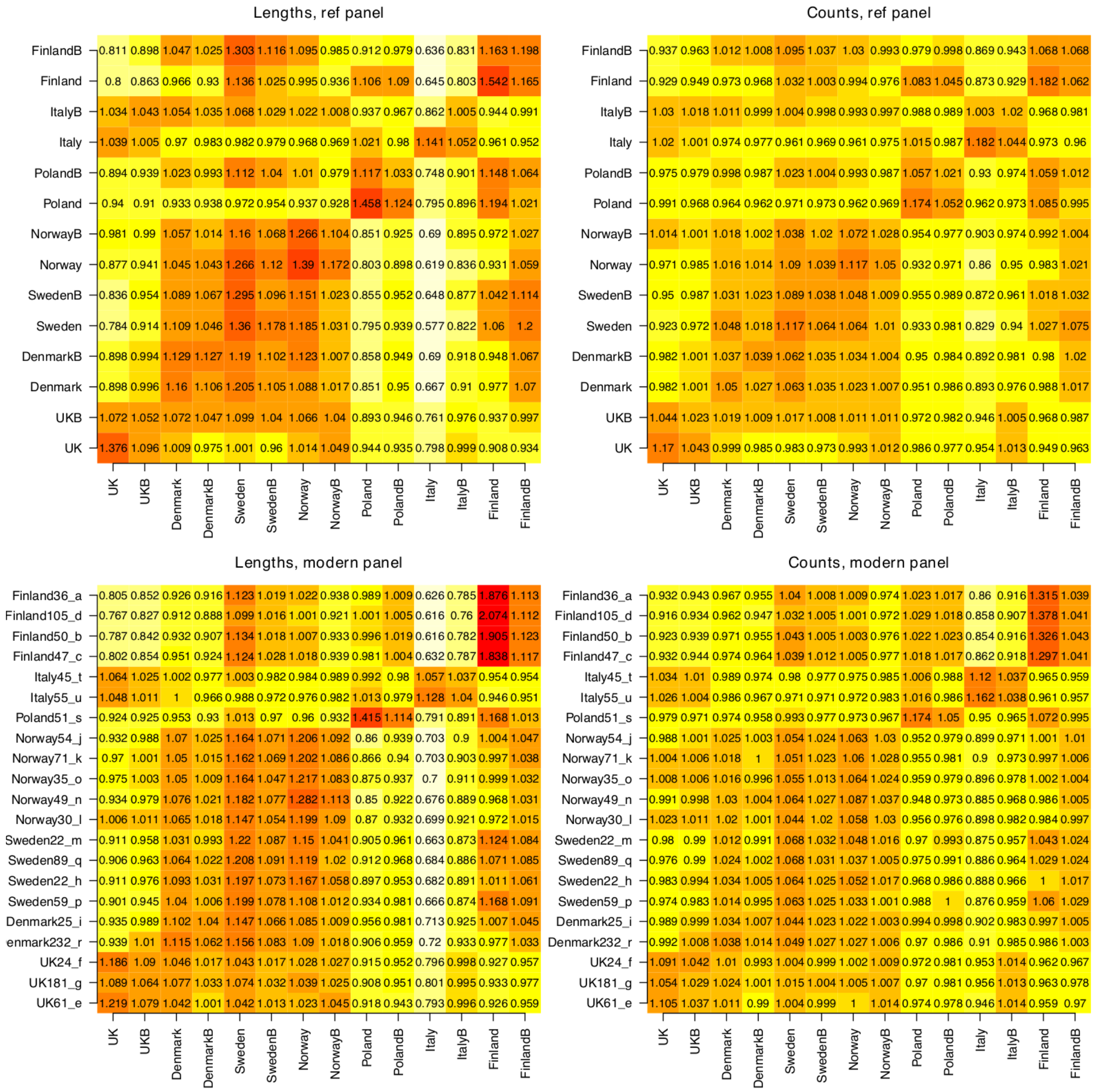


**Fig. S11.6**: Ancient DNA painting palette (top) and modern panel painted against the ancient panel (bottom). We show both the total genome shared (lengths) and inferred number of shared recombination events (counts). We use the lengths data for onward analysis. The plots show the average of each score, per donor individual, normalised to have mean value of 1. The rows show the average for each recipient individual in a population, when painted against the populations in the columns.

The importance of the leave-one-out procedure is that now the individuals used in the donor populations (who cannot share genome with themselves) are now exchangeable with the individuals in the modern populations. That is because they are all painted against exactly $n_k - 1$ donor individuals, where $n_k$ is the number of individuals in population $k$, regardless of whether the individual itself is in the panel.

#### Ancestry learning

The target of inference is an admixture profile for each modern and ancient individual. We use the first stage of the admixture estimation method as implemented in the software GLOBETROTTER [236]. To quantify uncertainty, we resample with replacement the per-chromosome palettes for each individual and reapply admixture estimation. We report the average and confidence interval over 100 random samplings as performed by [236]. Supplementary Table 6 gives the per-individual estimates of ancestry.

Given that we have performed imputation and the ancient genomes have variable coverage, it is important to test how this will effect inference. We first check whether sequence depth is driving any broad scale painting structures. Figure S11.7 shows that the results of the Unsupervised Ancient Sample Analysis are not associated with sequence depth.

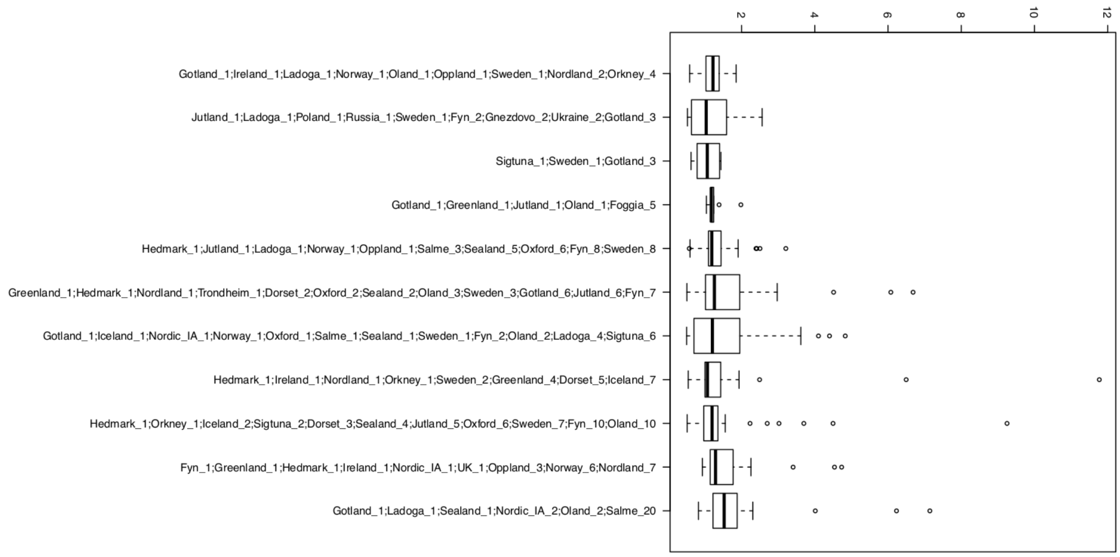


**Fig. S11.7**: The labels of the individuals in the unsupervised cluster membership (see Unsupervised Ancient Sample Analysis) showing the sequence depth of members. There is no association between membership and sequence depth.

Next we examined the fine-scale effects of sequence depth by downsampling two high quality genomes, VK1 (Greenland East Settlement; sequence depth 11.8) and VK50 (Gottland; sequence depth 6.2), at the raw read level, to an average sequence depth of 1,2, or 4. We repeat the analysis as if this individual were only available at the specified sequence depth. Figure S11.8 shows how the inference changes as a function of sequence depth, along with the individual bootstrap samples. In general, there is surprisingly little variation as a function of sequence depth, with some evidence that 1X is biased towards a more mixed solution, but not dramatically worse than the variation induced by changing depth. There is no evidence that a) wrong ancestry components are added, or b) major changes in ancestry can occur. All changes are within Scandinavian ancestries and the correct components are always recovered.

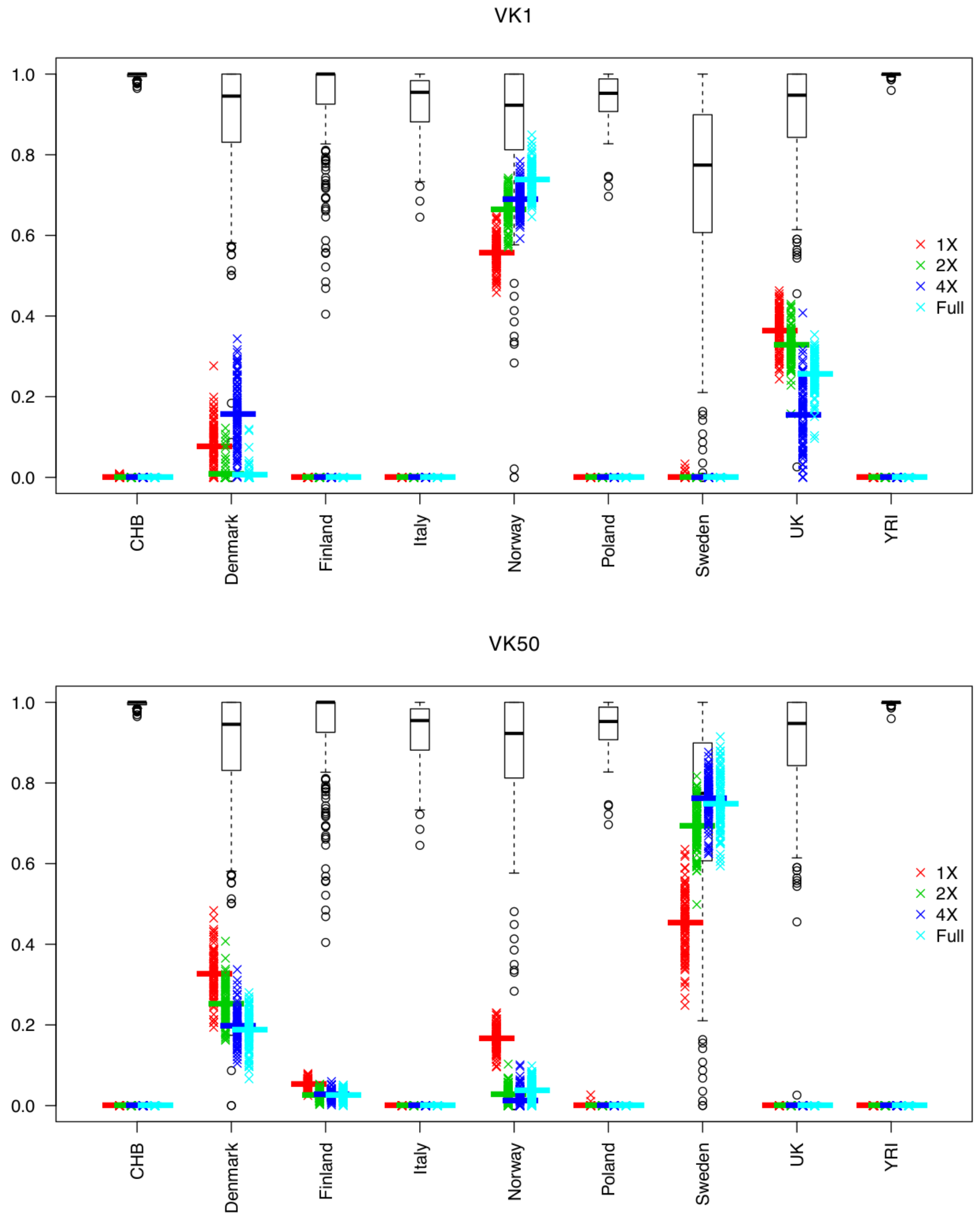


**Fig. S11.8**: Top: VK1, Bottom: VK50. The mean results along with 100 bootstrap resamples of the painting are shown for ancestry reconstruction from the 7 populations. Results are shown for each sequence depth (crosses of different colours: red/blue/green/cyan). Also shown is the distribution of each component within the specified population (black boxplots/circles); e.g. the "Denmark" boxplot shows the distribution of Denmark ancestry for individuals from the Denmark population.

#### Spatio-temporal Regression Model

To assess the spatio-temporal structure in the data, we use a simple regression model:

$$a_{ik} = \alpha_{jk} t_i + \beta_{jk} x_i + \gamma_{jk} y_i + \varepsilon_{ijk},$$

where $a_{ik}$ is the amount of ancestry individual $i$ possesses from population $j$, $t_i$ is the "age category" of the individual (1=iron age, 2=early Viking, 3= Viking, 4=Medieval) and $(x_i, y_i)$ are the longitude and latitude of the death location of the individual. We choose to analyse the samples by region, hence restricting to a subset of individuals $i \in P_r$ found in that region $r$ (UK, Scandinavia, whole Europe including UK and Scandinavia). The results of the inference are a time-effect $\alpha_{jk}$, a longitude effect $\beta_{jk}$ and a latitude effect $\gamma_{jk}$.

We perform time-tests in Scandinavia and Europe, and lat-long tests in Scandinavia, UK, Europe for a total of 8 tests, each run for 7 ancestries.

**Table S11.2**: Regression results for spatio-temporal structure in ancestries. Each row shows the value for a region (named in the columns) of $\alpha_{jk}$ (for age), $\beta_{jk}$ (for longitude) and $\gamma_{jk}$ (for latitude) in the model described in Spatio-temporal Regression Model. Significance thresholds: *: $p < 0.05$, **: $p < 0.005$, ***: $p < 0.0005$.

| | **Scandinavia Age** | **Scandinavia Lat** | **Scandinavia Long** | **Europe Age** | **Europe Lat** | **Europe Long** | **UK Lat** | **UK Long** |
|---|---|---|---|---|---|---|---|---|
| **UK** | 0.106 | 0.008 | -0.004 | 0.07* | 0.008* | -0.004* | **0.06*** ** | -0.027 |
| **Denmark** | **0.193** ** | **-0.022*** ** | -0.003 | 0.073* | -0.012* | 2.4e-4* | -0.022 | **0.061** ** |
| **Sweden** | **-0.613*** ** | -0.001 | 0.001 | **-0.374*** ** | **0.001*** ** | **0.003*** ** | -0.005 | 0.016 |
| **Norway** | **-0.173** ** | **0.037*** ** | **-0.04*** ** | 0.047 | 0.018 | -0.005 | -0.017 | **-0.068** ** |
| **Poland** | **0.188*** ** | -0.007 | **0.018** ** | 0.069* | -0.003* | 0.003* | -0.001 | 0.003 |
| **Italy** | **0.136** ** | **-0.015*** ** | -0.002 | **0.129*** ** | **-0.015*** ** | **0.001*** ** | -0.012 | 0.012 |
| **Finland** | 0.163* | -0.002 | **0.025*** ** | -0.014 | 0.003 | 0.003 | -0.003 | 0.004 |

Such regression analyses are dependent on the sample locations and choice of geographical region. They are therefore intended as a guide to formalise the spatio-temporal relationships that are clear by-eye in the maps smoothing ancestry estimates. They are helpful to understand what the inferred ancestry components "mean". The within-Scandinavia results make it clear that the group called "Sweden" represents a historical population that once existed in Sweden, replaced by more southern population/s containing more continental European ancestry. Similarly, Norwegian ancestry has declined but is still higher in the North-West of Scandinavia (i.e. Norway). Italian and Danish ancestry both increase over time and are higher in the south of Scandinavia, consistent with a migration flow. Similarly, the UK analysis shows that the population labelled "UK" was at the time of sampling predominant in the north, with "Danish"-like ancestry found in the East and Norwegian in the West.

#### Principal Components Analysis of Painting

We perform a Principal Components Analysis of the covariance of the painting which (if the data were treated as containing no linkage-disequilibrium) is equivalent under some theoretical conditions [235]. Specifically, we formed an $N_{M+A}$ by M matrix of the ancient and modern combined palette painting, calculate the $N_{M+A}$ by $N_{M+A}$ matrix of covariance, and compute the eigenvalue decomposition using the function "eigen" in R. Figure S11.9 shows the results of this analysis.

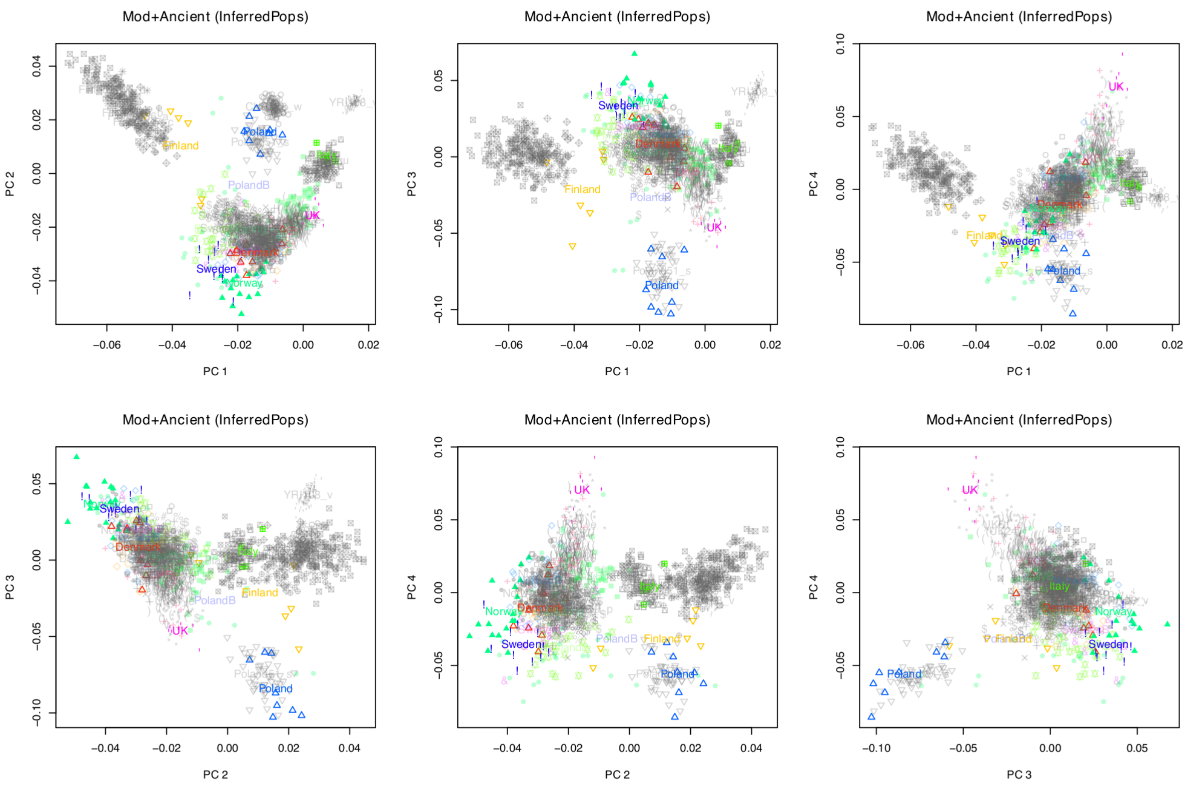


**Fig. S11.9**: PCA of the ancient and modern samples using the ancient palette, showing different PCs. Modern individuals are grey and the *K=7* ancient panel surrogate populations are shown in strong colors, whilst the remaining *M-K=7* ancient populations are shown in faded colors.

Several features are clear from Figure S11.9. Firstly, modern Finnish individuals are not like ancient Finnish individuals, modern individuals have ancestry of a population not in the reference; most likely Steppe/Russian ancestry, as Chinese are in the reference and do not share this direction. Ancient Swedes and Norwegians are more extreme than modern individuals in PC2 and 4. Ancient UK individuals were more extreme than Modern UK individuals in PC3 and 4. Ancient Danish individuals look rather similar to modern individuals from all over Scandinavia.

By using a supervised ancient panel, we have removed recent drift from the signal, which would have affected modern Scandinavians and Finnish populations especially. This is in general a desirable feature but it is important to check that it has not affected inference. As a sensitivity analysis, we therefore repeated the analysis but this time using the modern and ancient samples together in “all-vs-all mode”, using the same label annotations as above. This leads to a $N_{M+A}$ by $N_{M+A}$“coancestry matrix” which we normalise [235] in the standard way (zero mean rows, standardize variance), before removing African and Chinese individuals from the rows, so that they are not projected but that their ancestry is represented. We then use the R function "svd" to perform a Singular Value Decomposition (theoretically and practically equivalent to computing the Principal Components). This is shown in Figure S11.10.

As expected, variation in the large modern dataset dominates the picture. Further, the early PCs are dominated by within-population structure. Whilst PC1 describes similarity with Africa for high values, PC2 describes Finnish ancestry, PC3 variation within Finnish ancestry, PC4 describes Norwegian ancestry and PC5 describes variation within Norway at one end of its range. PC8 describes the first batch effect between modern and ancient samples; that it is a relatively small effect compared to the population structure is additional information that the imputation procedure has not biased the inference.

The story for Modern-vs-ancient Finnish ancestry is consistent, with ancient Finns looking much less extreme than the moderns. Conversely, ancient Norwegians look like less-drifted modern Norwegians; the Danish admixture seen through the use of ancient DNA is hard to detect because of the extreme drift within Norway that has occurred since the admixture event. PC4 vs PC5 is the most important plot for the ancient DNA story: Sweden and the UK (along with Poland, Italy and to an extent also Norway) are visibly extremes of a distribution the same "genes-mirror-geography"[4] that was seen in the Ancient-palette analysis. PC1 vs PC2 tells the same story – and stronger, since this is a high variance-explained PC - for the UK, Poland and Italy.

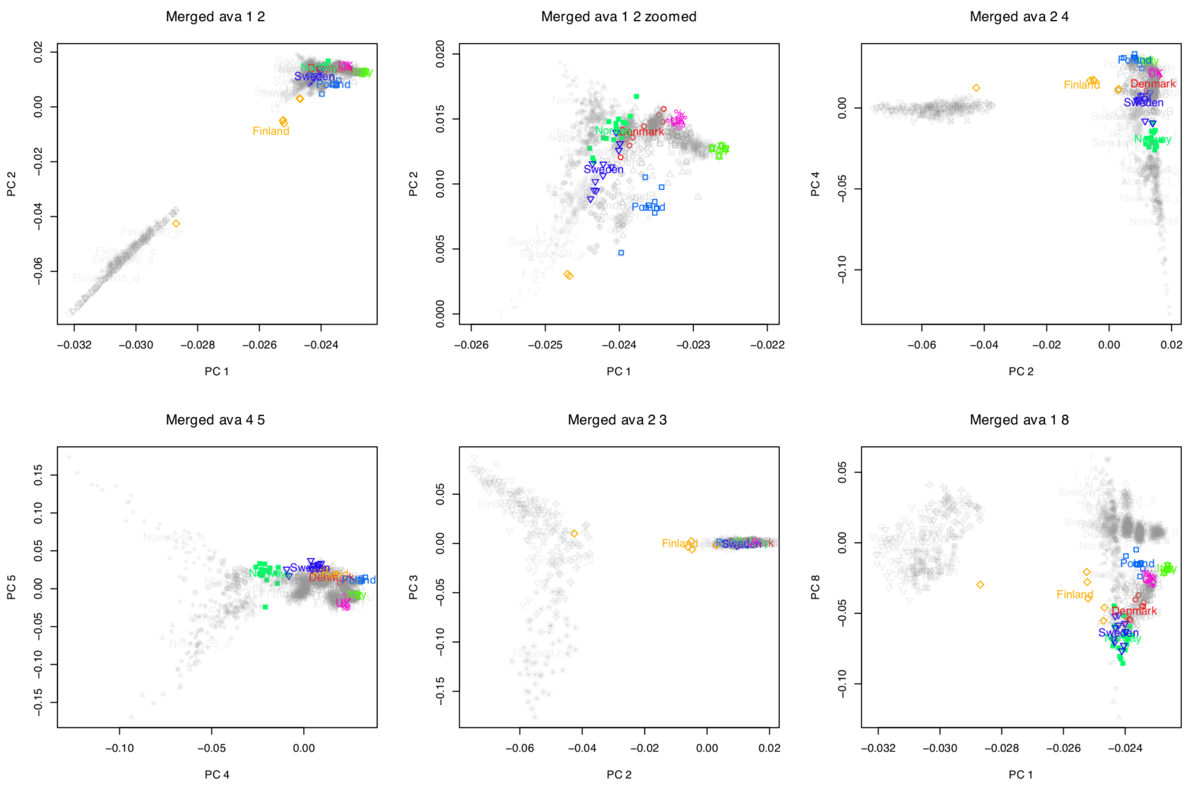


**Fig. S11.10**: PCA of the ancient and modern samples using no palette (see text), showing selected PCs to illustrate the salient features. Modern individuals are grey and the $K=7$ ancient panel surrogate populations are shown in strong colors.

PCA should be carefully interpreted as it can easily be misleading[3,5]; for example, individuals from very different populations can be mapped to the same location in some PCs if they do not differ in their variation in the directions being displayed. The PCs are also dominated by variance components that represent a large number of individuals (e.g. modern variation), or a few individuals that share a large fraction of the variance (e.g. Finnish). Despite these problems, the PC analysis provides support for the inferences described through the chromosome painting admixture analysis.

#### Interpretation: Inference of historical ancestry sharing using Chromosome Painting

The first summary of the data to consider is the population averages of ancestry by population, commonly called the confusion matrix. This is shown in Figure S11.11, which as a sensitivity analysis includes an analysis based on “chunk counts” as used by FineSTRUCTURE as well as the total amount of genome shared, called chunk lengths, on which all other inference is based.

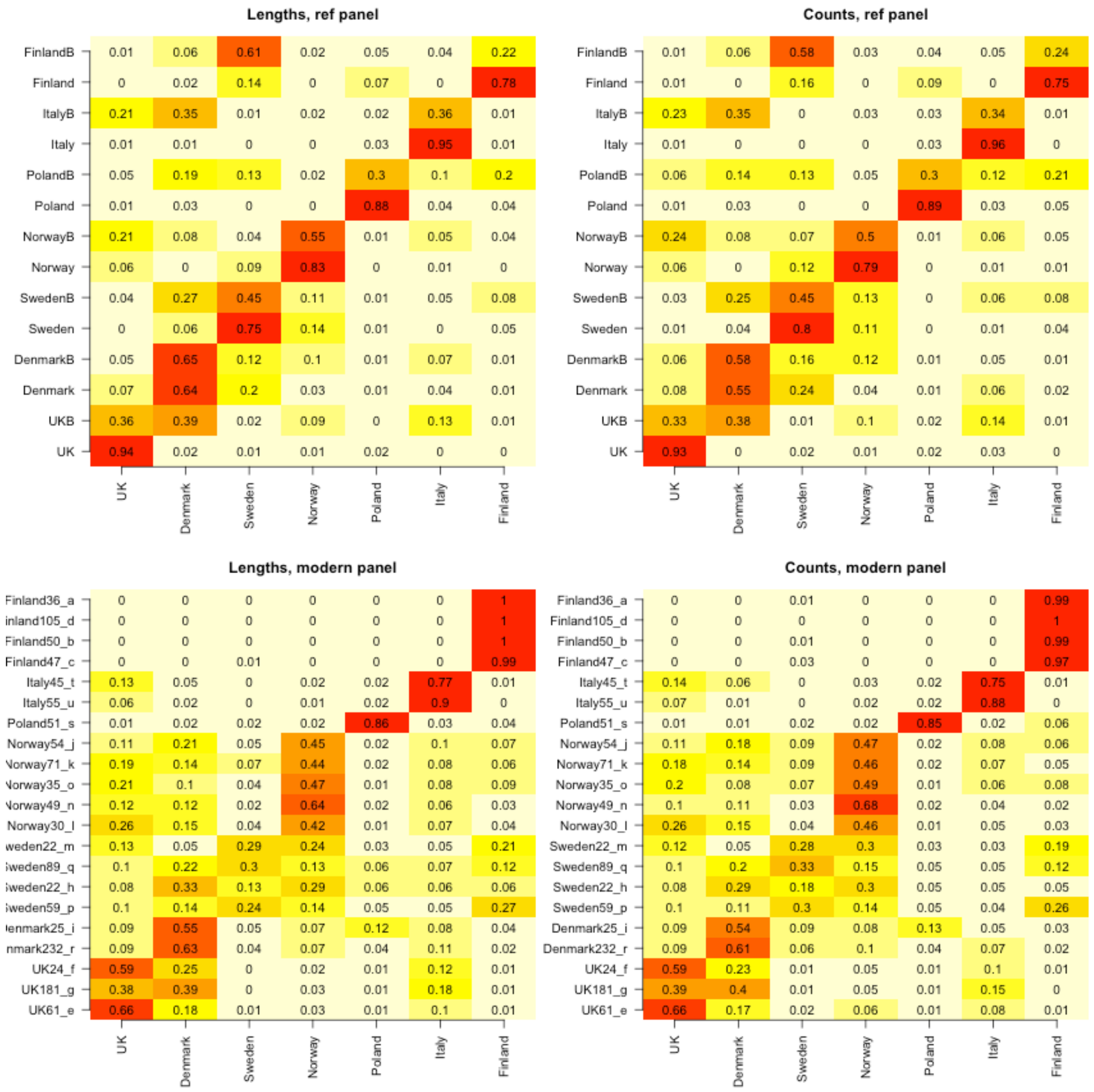


**Fig. S11.11**: Left: results based on inference using the total length of genome shared with individuals from each donor population. Right: results based on inference using the total number of "chunks" (most recent recombination events). Top: ancient panel. Bottom: modern panel.

The general structure of this inference is:

a) We can recover the representative donors' reference populations well.
b) Lengths perform slightly better than counts for this problem.
c) The populations labelled "B", who are "non-representative donors", are always more of a mixture than the "representative donors".
d) Norwegian ancestry is found primarily in Scandinavia, with small amounts in UKB.

e) Swedish ancestry is found primarily in Sweden and Scandinavia, though is also found in Finland and PolandB.
f) UK ancestry is found primarily in the UK but also in the ItalyB and NorwayB populations.
g) Denmark ancestry is found in nearly every B population though is lowest in NorwayB and FinlandB.
h) Poland ancestry is restricted to Poland and Finland.
i) Italy ancestry is found in all B populations and no representative populations.
j) Finland ancestry is found in Finland, PolandB and SwedenB only.

These are inferred populations that were labelled based on which ancestral population "was closest to them", rather than which they were closest to. Empirically this is symmetric for all cases except FinlandB, who are more related to Sweden than Finland on average.

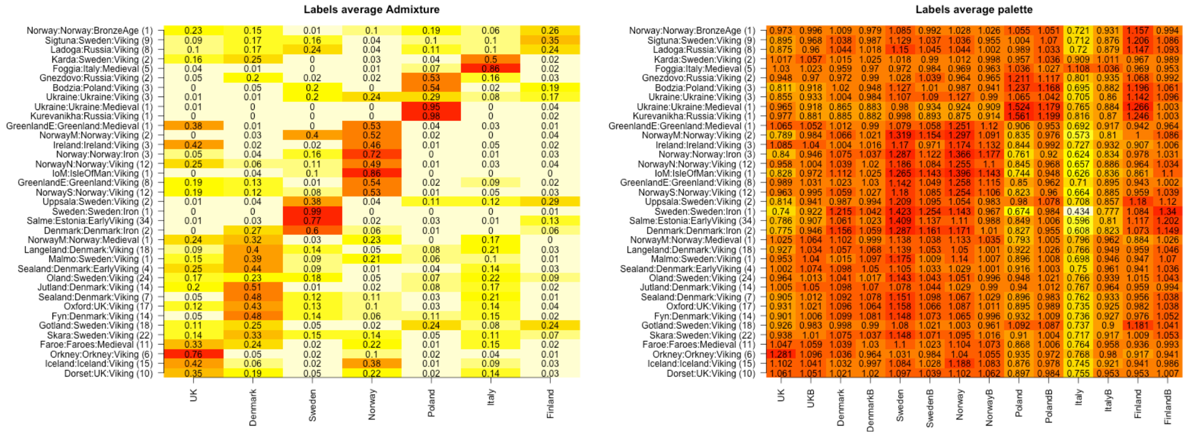


**Fig. S11.12**: Results presented in terms of the labels, which are shown in "Region:Country:Age (sample size)" format. Left: Inferred ancestry. Right: Average painting palettes.

Figure S11.12 shows the same results as Figure S11.11 (for admixture) and Figure S11.6 (for palettes) but presented by labels, that is, the region, country and age of each set of samples. It must be remembered that the individuals concerned were found in heterogeneous locations and where there are multiple individuals, may be heterogeneous genetically. To explore diversity of sites explicitly, we devised a measure of diversity that could characterise highly diverse vs homogeneous sample labels, shown in Extended Data Fig. 5.

For the purpose of characterising diversity there isn't a ready off-the-shelf measure. The data consist of a diversity of probability vectors. We quantify their diversity by computing the average Kullback-Leibler (KL) Divergence for each individual label from the average of that label:

$$D(A^{(l)}) = \frac{1}{n_l}\sum_{i=1}^{n_l} KL(A_i^{(l)} \,||\, p^{(l)})$$

where $A^{(l)}$ is the $n_l$ by $K$ matrix of ancestry estimates in label $l$, $p^{(l)}$ is the length $K$ vector of average ancestries in that label, and $KL(Q \,||\, P) = \sum_{k=1}^{K} q_k log_2\left(\frac{q_k}{p_k}\right)$. This therefore measures a population to be "diverse" if there is a large deviation of individual ancestry estimates away from the average ancestry in that population.

We confirm that this score is well-calibrated using simulations. We simulated from a hierarchical dirichlet setup where we make a random "population mean" $\alpha \sim Dirichlet(rep(\alpha_0, K))$ and then sample individuals $x_i \sim Dirichlet(\alpha)$. This is shown in Figure S11.13 (right), with a search of good choices of $\alpha_0$(left) being guided by the fit to the data under a normal distribution approximation. This simulation confirms that the diversity measure is appropriate for the data.

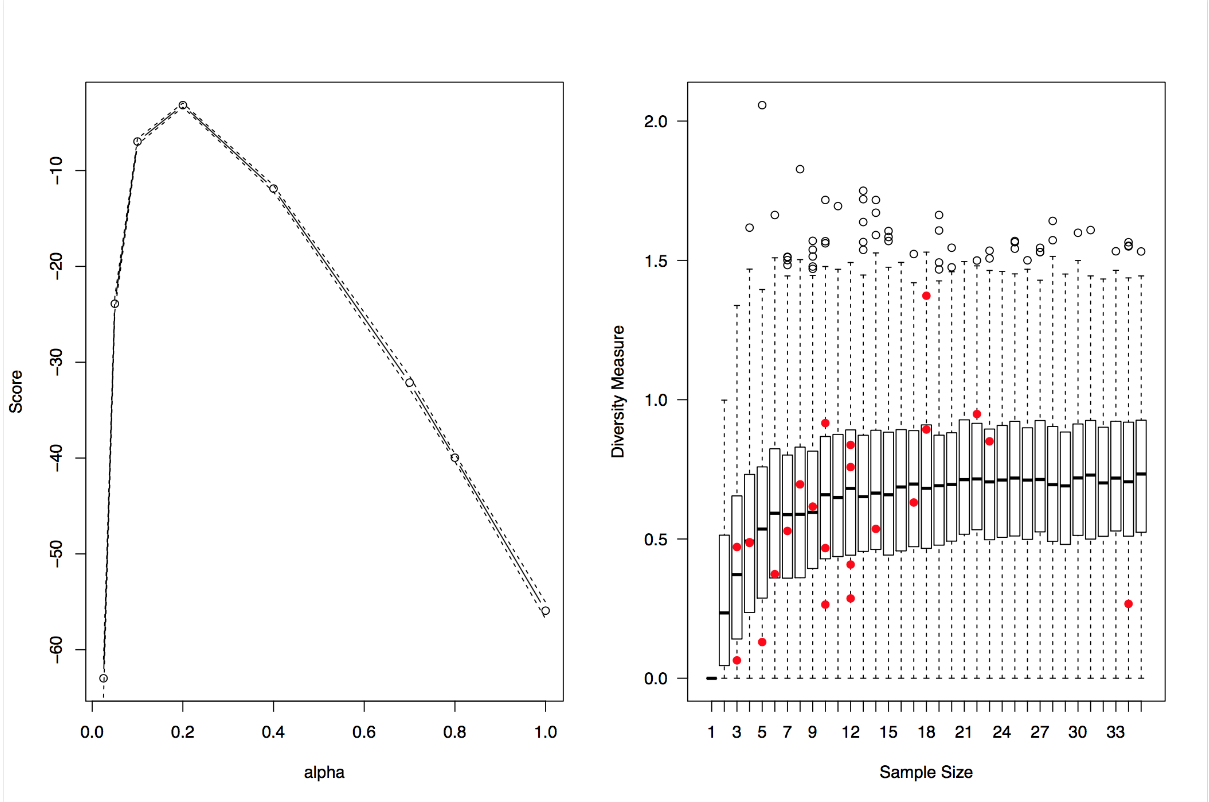


**Fig. S11.13**: Simulated diversity measures for labels. Left: quality of fit of simulations to the data as a function of a tuning parameter $\alpha_0$ (see text). Right: simulated data (boxplots) and observed data (red dots) as a function of sample size, to check model fit. Note that the distribution is testing model fit, not statistical significance of diversity.

This exploration of diversity raises several points. Firstly, Bodzia (which is associated with our 'Poland' population), Foggia (associated with our 'Italy' population), and Salme (which is associated with out 'Sweden' population) are relatively homogeneous. Dorset is also homogeneous, containing primarily 'Norway' or 'UK' ancestry.

Extreme diversity is seen in Gotland, a well-known trading location. Other locations such as Skara and North Norway are diverse due to the presence of UK, Danish, Swedish and Norway ancestry. The overall picture is of high-diversity - and high heterogeneity - in Scandinavia, brought in from outside. Sampling must be considered for the peripheral regions, which were sampled to target Viking cultural burial.

#### Modern Interpretation: Inference of historical ancestry sharing using Chromosome Painting

The estimates in Figure S11.11 of ancestry in modern populations from historical populations is a mean, and therefore does not separate ancestry contributions from recent admixture from that which is typical of the population. The median is not affected by some individuals with relatively recent ancestry, but instead produces estimates that do not sum to 1. To estimate ancestry of populations, we therefore consider the *spatial median* of the individual ancestry estimates using the R package "ICSNP"[237], which is a multivariate extension of the median that preserves the sum.

We estimate confidence intervals by resampling individuals with replacement within each population and recomputing the spatial median. We report the 95% confidence range (2.5% and 97.5% quantiles).

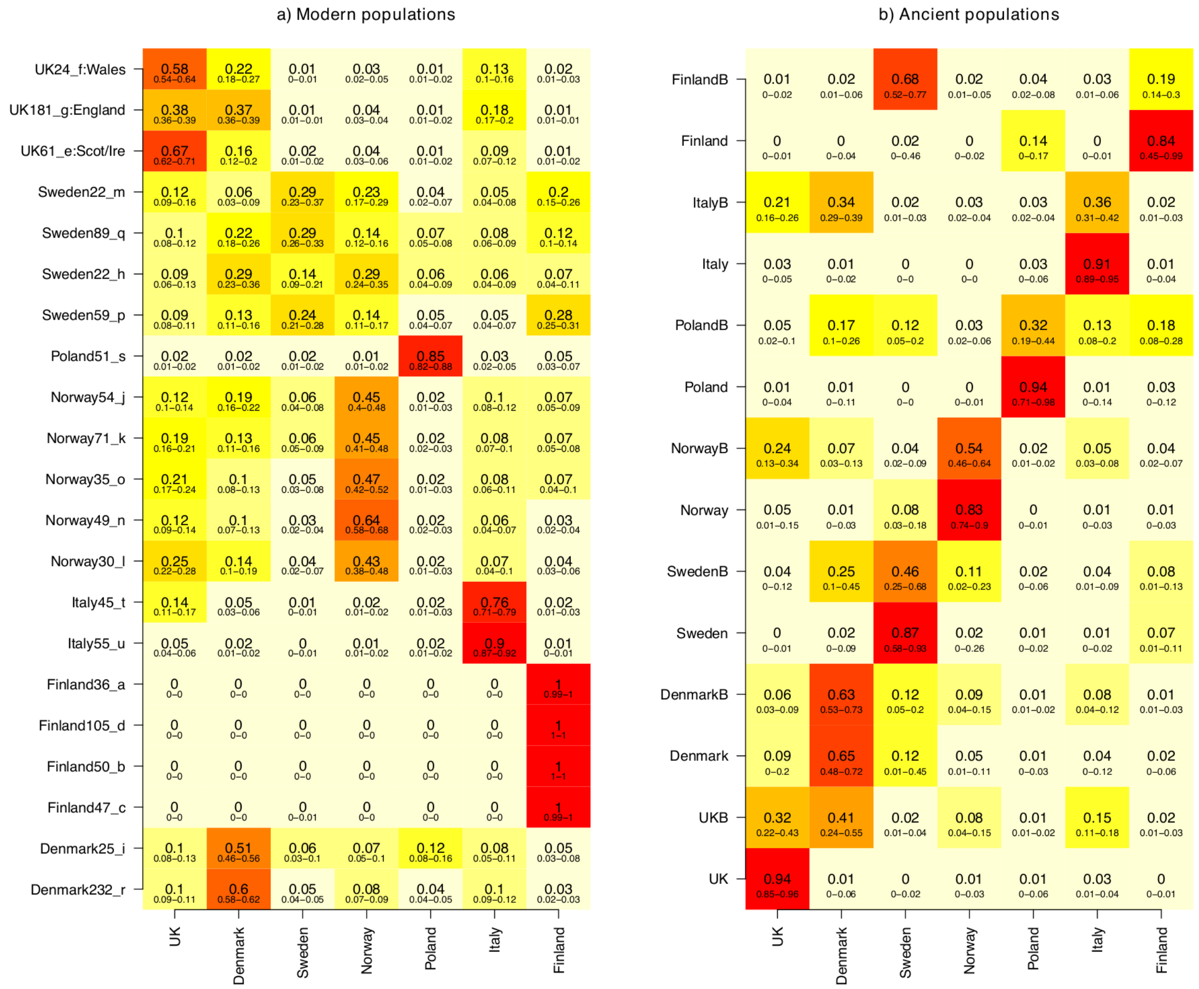


**Fig. S11.14**: Population-based admixture estimates based on the spatial median, as described in the text. a) estimates of the proportion of ancient populations in modern populations. b) estimates of the proportion of ancient populations in ancient samples.

Figure S11.15 shows our best estimates for modern populations, and the ancient sample groupings, based on this spatial median approach. The ancient reference populations are all seen as broadly their own population. The ancient non-reference populations may not be meaningful as they are not expected to be homogenous. However, the modern populations are relatively homogeneous and the spatial median should represent the typical individual from these populations.

From this, we can see the spread of ancestry during and around the Viking era:

- UK populations have all received high ‘Denmark’ ancestry. Although Anglo-Saxon and Danish Viking ancestry are hard to distinguish, Viking-era Danes have too much “Sweden” ancestry to have contributed more than around 6% ancestry into England, whereas they could plausibly have contributed all (up to 16%) of the Scottish and Irish signal. Anglo-Saxon samples are needed to explore this further.

- The 'Norway' ancestry signal in the UK cannot be explained via the Danish or Anglo-Saxon contribution. These fractions (4% in England, Scotland, and Ireland, 3% in Wales) likely correspond to the Norwegian Viking legacy in Britain.
- Modern UK individuals contain around 9-18% 'Italian' ancestry, plausibly associated with the Normans and associated increase in population movement during that era. This is a two-way process, with high fractions of the 'UK-like' ancestry in a sub-population of Italians.
- Modern Norwegians are structured by their proportion of 'UK' and 'Danish' ancestry.
- Modern Swedes are structured by a 'Finland'-like group, a 'Denmark'+'Norway' group, and a 'UK' group.
- Modern Danes are structured into high and low 'Polish' ancestry groups, both with similar amounts of 'Norway' and 'UK' ancestry, suggesting that these admixtures occurred earlier. Indeed, the ancient panel implies that this process started in the Viking era, where the high confidence interval is explained by high inter-individual variation.

To understand the biases involved in the use of spatial.median we plot the estimates from this procedure compared to the individual data in Figure S11.15. From this it is clear that small estimates are biased upwards from the sample median for each ancestry, due to the constraint that ancestries should sum to 1. This has been accounted for in the discussion above.

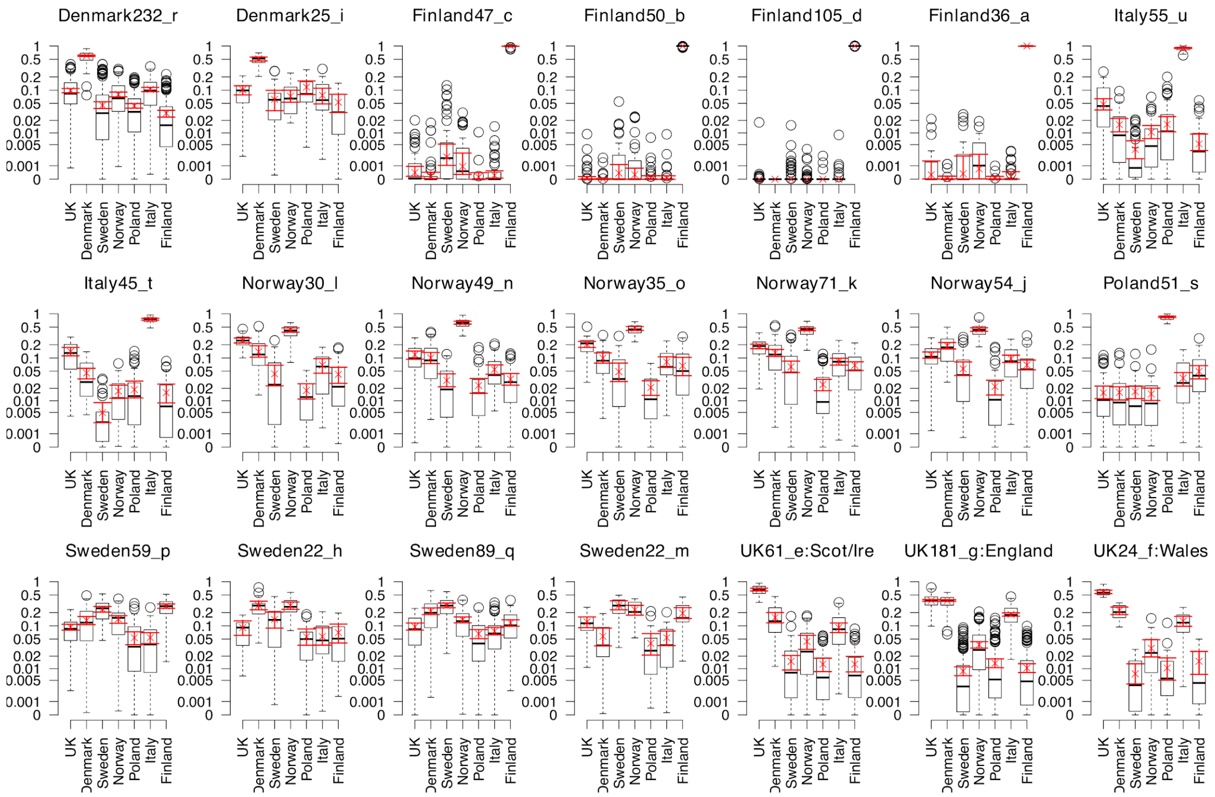


**Fig. S11.15**: Spatial median estimates (red; crosses for best estimate with range shown) and individual estimates (black; reported as a boxplot) for each modern population using the 7 ancient reference populations. Black horizontal bars denote the median for each sample.

#### Evidence for Pictish Genomes

Our interpretation for the Orkney samples can be summarised as follows. Firstly, they represent "native British" ancestry, rather than an unusual type of Scandinavian ancestry. Secondly, that this "British" ancestry was found in Britain before the Anglo-Saxon migrations. Finally, that in Orkney, these individuals would have descended from Pictish populations. The evidence for this interpretation is:

1. In Figure S11.10 showing modern and ancient samples painted together and analysed using PCA without any supervision, the 'UK' cluster is outlying far from Scandinavia in PC1-2 and PC4-5, and PC3 is describing drift inside modern Finland.
2. In Figure S11.9 showing modern and ancient samples painted together and analysed using PCA after projection into our supervised clusters, the 'UK' cluster is further outlying. In PC1-

3 it clusters with nothing but is closest to ‘Italy’ and ‘Poland’. In PC4 it is an extremal point opposite Poland and Finland.

3. In Figure S11.3 showing ancient individuals painted against modern populations, the Orkney individuals best match modern Scottish populations (UK61_e), and show no particular affinity to modern Scandinavian ancestry. In Figure S11.4 showing the importance of modern populations for ancient samples, the modern UK populations do not well describe this variation.
4. In Figure S11.6a-b showing the painting of ancient population clusters against one another, the UK population is rarely used to describe Scandinavians (low values in the ‘UK’ column), whereas the Italians do. Conversely, the ‘UK’ population does use an excess of Norway ancestry, implying a degree of contemporary admixture into that population (likely due to low sample sizes of unadmixed individuals).
5. In Figure S11.6c-d showing the painting of modern population clusters against ancients, the Irish and Scottish (UK61_e) and Welsh (UK24_f) populations receive the most ‘UK’ ancestry. This is followed by Italy, and then some Norwegian subpopulations into which British is known to have been taken.
6. In Figure S11.11 showing the inferred ancestry of ancient populations, ‘UK’ ancestry is little seen in other main clusters, and appears in known admixtures including ‘NorwayB’ and ‘ItalyB’.
7. In Figure S11.12 showing the inferred ancestry of ancient sampling locations, the ‘UK’ ancestry is found at highest proportions in Orkney, Iceland, Ireland, Dorset in England, and Medieval Faroe Islands.
8. Conversely, of the 10 individuals with more than 80% ‘UK’ ancestry (Supplementary Table 6) only 4 are in Orkney and 1 in Ireland. The rest are in Scandinavia during the Viking period: Norway (VK386, VK525, VK528) and Sweden (VK456, VK405). These populations are sampled more heavily and the overall rate of ‘UK’ ancestry in those locations is low. This is consistent with the migration of individuals with ‘UK’ ancestry into Scandinavia, rather than a population present in Scandinavia.
9. In Figure S11.14-15 showing modern admixture estimates, the “UK” fraction is ordered: Scotland and Ireland, Wales, England, Norway30_l, rest of Norway, Northern Italy, Denmark and Sweden. This shows that this ancestry is located in the UK today, though widely

distributed throughout Europe. Note that we have not shown whether the signal in Italy is due to post-Viking era admixture, or a signal of ancient population structure.

Therefore 'UK' represents a group from which modern British and Irish people all receive an ancestry component. This information together implies that within the sampling frame of our data, they are proxying the 'Briton' component in UK ancestry; that is, a pre-Roman genetic component present across the UK. Given they were found in Orkney, this makes it very likely that they were descended from a Pictish population.

Modern genetic variation within the UK[199] sees variation between 'native Briton' populations Wales, Scotland, Cornwall and Ireland as large compared to that within the more 'Anglo-Saxon' English. This is despite subsequent gene flow into those populations from English-like populations. We have not attempted to disentangle modern genetic drift from historically distinct populations. Roman-era period people in England, Wales, Ireland and Scotland may not have been genetically close to these Orkney individuals, but our results show that they have a shared genetic component as they represent the same direction of variation.

#### Supplementary Note 12 - Spatiotemporal patterns of ancestry in the Viking World

We used ordinary kriging implemented in the function 'idw' of the R package *gstat*, to interpolate the proportion of each ancient genome that was attributed by our *fineStructure* analysis to one of the pre-defined ancestry groups: 'UK', 'Denmark', 'Norway', 'Sweden', 'Italy', 'Poland' and 'Finland'.

We chose to plot a Europe-wide map - including Greenland - (Figure S12.1) and smaller maps of particular regions of interest: Scandinavia (Figures S12.2-S12.4), the British Isles (Figure S12.5) and the Baltic region (Figure S12.6). For Europe-wide maps, we used a grid size of 0.2° x 0.2°. For all other maps, we used a grid size of 0.1° x 0.13°. As our densest temporal sampling was for Scandinavia, we made separate Scandinavian maps for the Iron Age (Figure S12.2), the Early Viking Age (Figure S12.3) and the Viking Age (Figure S12.4). For all other maps, we combined samples across time periods, but the vast majority of our samples were from the Viking Age, so our signal was dominated by this particular period: we only have Viking Age samples from the British Isles, while for the Baltic region we only have Viking and Early Viking Age samples.

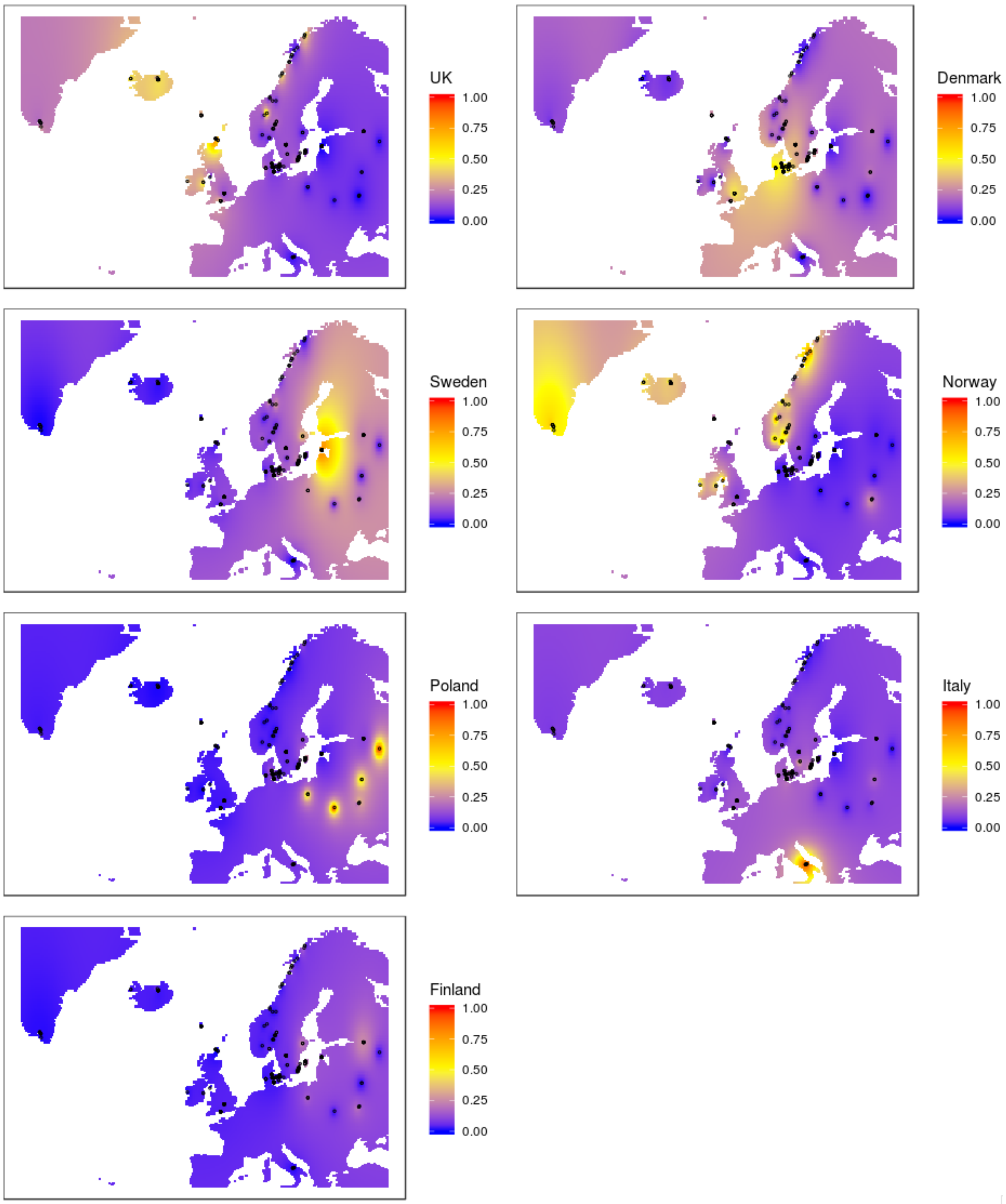


**Fig. S12.1:** Maps of interpolated *fineStructure* ancestries for Europe and Greenland, combining Iron Age, Early Viking Age and Viking Age samples.

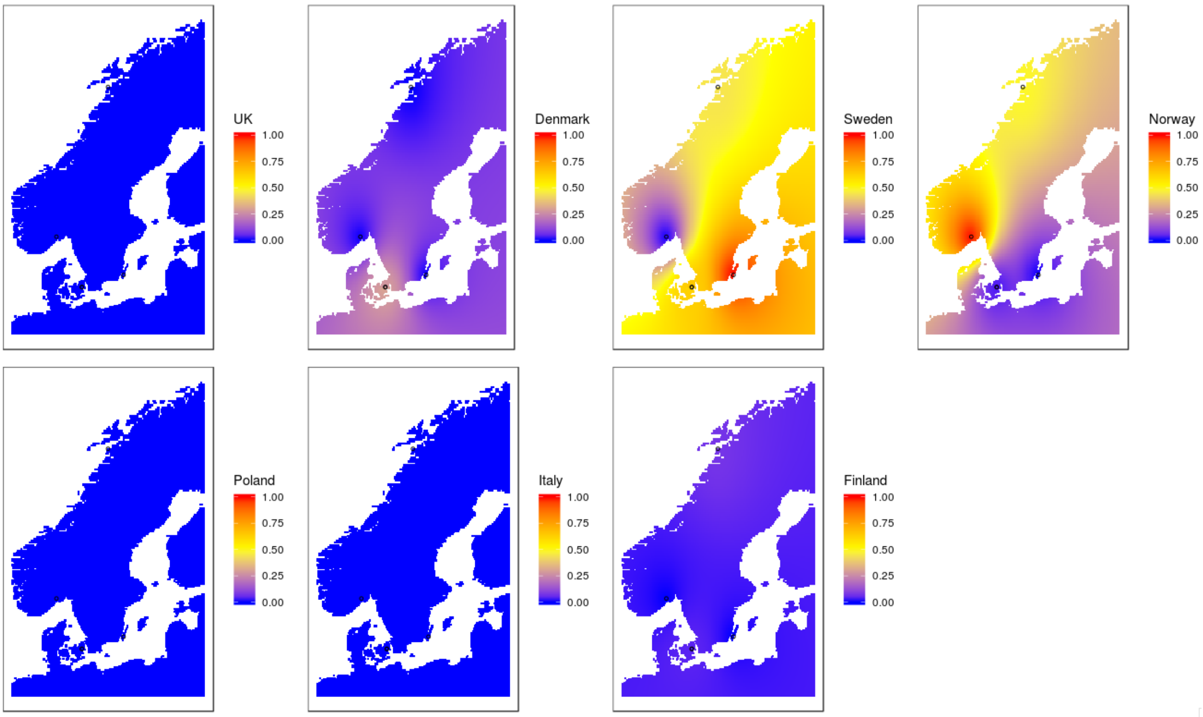


**Fig. S12.2:** Maps of interpolated *fineStructure* ancestries for Scandinavian Iron Age samples.

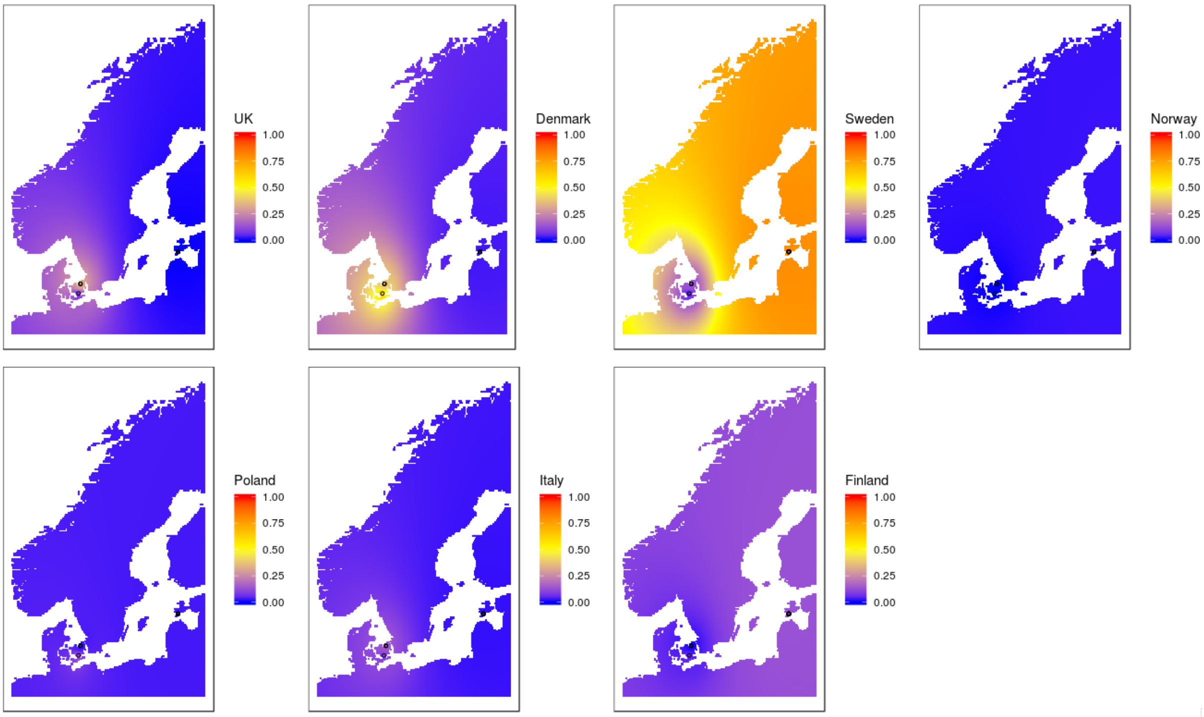


**Fig. S12.3:** Maps of interpolated *fineStructure* ancestries for Scandinavian Early Viking Age samples.

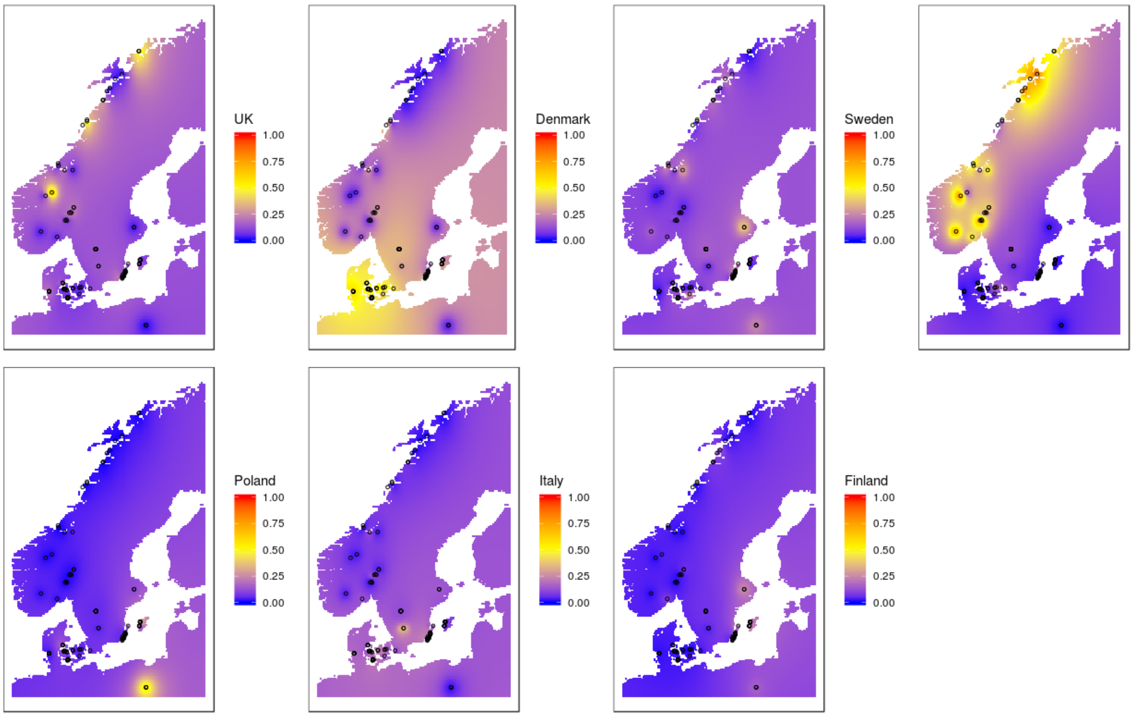


**Fig. S12.4:** Maps of interpolated *fineStructure* ancestries for Scandinavian Viking Age samples.

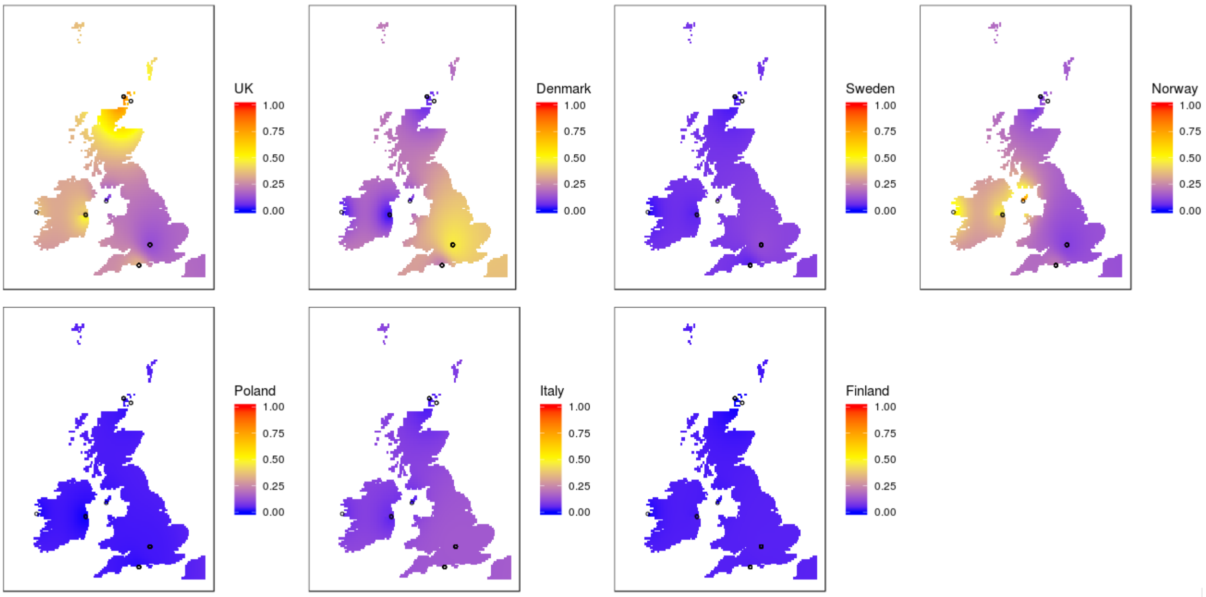


**Fig. S12.5:** Maps of interpolated *fineStructure* ancestries for Viking Age samples from the British Isles.

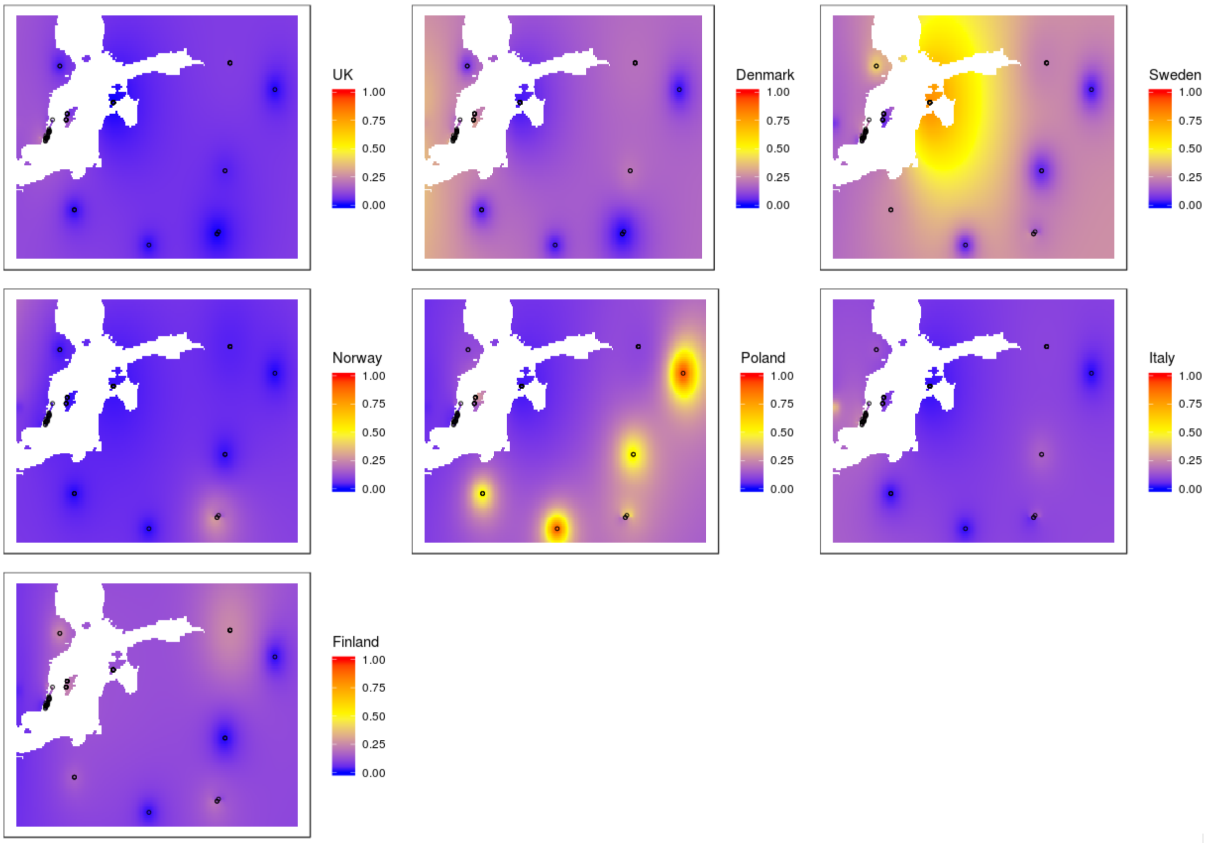


**Fig. S12.6:** Maps of interpolated *fineStructure* ancestries for Early Viking and Viking Age samples from the Baltic region.

### Supplementary Note 13 - Lactase persistence and pigmentation SNPs

#### Lactase persistence

To investigate the level of lactase persistence in the ancient population we estimated the derived ‘A’ allele frequency of the SNP rs4988235 known to affect expression of the lactase LCT gene. The ancestral “G” allele is responsible for lactase intolerance in adult Europeans[238]. Since most of the ancient samples were sequenced at low depth of coverage, the ANGSD software package was used to estimate the allele frequencies of the ancient population based on the genotype likelihood data. We used the five European populations (CEU – Northern European, FIN – Fins, GBR – British, TSI – Italy, IBS – Spain) and two outgroups (Yoruba – YRI; Chinese – CHB) from the 1000 Genomes Project as comparative groups as well as the modern Danish population from the IPSYCH case-cohort study[239]. The results are presented in Figure 5.

#### Pigmentation

Having a large number of ancient individuals allowed us to assess the frequencies of SNPs responsible for pigmentation phenotypes of the ancient dataset (Vikings) at population level. For this analysis we have used ancient individuals from the whole Vikin Age period (i.e. Early Viking Age + Viking Age) from Scandinavia (n=262).

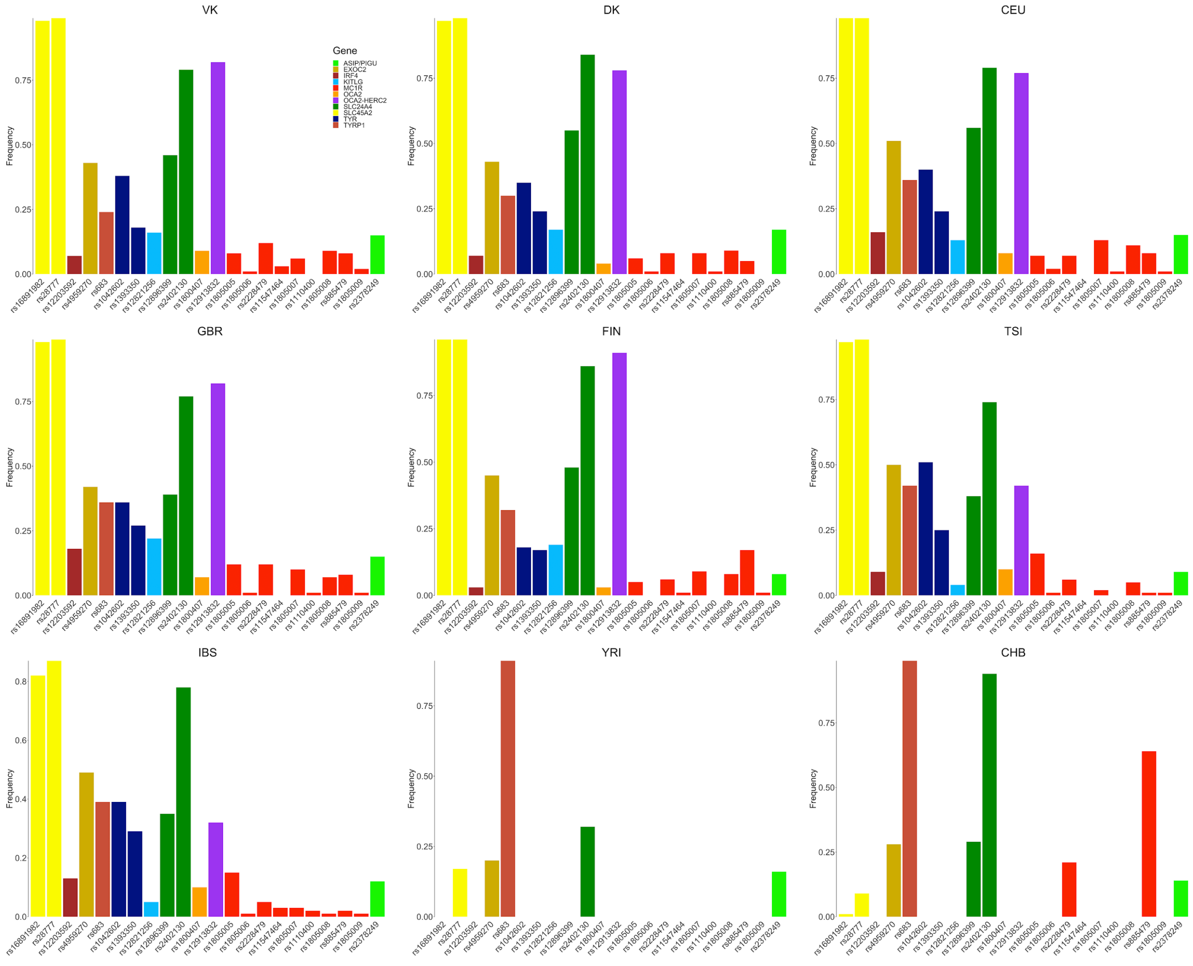


**Fig. S13.1**: The frequencies of derived SNPs with strongest influence on human pigmentation phenotype in Viking ("VK") and comparative groups. The rs IDs of the SNPs are indicated on the X axis. The genes that contain these SNPs are mentioned in the figure legend. Eight additional populations are also included for comparison: CEU, FIN, GBR, TSI, IBS; YRI; CHB from the 1000 Genomes project and "DK" representing the modern Danish population from IPSYCH case-cohort study.

The SNPs with strongest association with lighter hair and eye pigmentation phenotypes such as the ones in HERC2, OCA2 and TYR genes in humans are elevated in the Viking population, and the profile of allele frequency distribution is close to the present-day Northern European population represented here by the "CEU" (1000 Genomes Project) and the modern Danish population ("DK") from IPSYCH case-cohort study. The frequencies of informative SNPs associated with pigmentation are presented in Figure S13.1.

This suggests that the genetic profile of pigmentation SNPs we observe in northern Europeans today had been largely formed at the onset of the Viking period.

We have applied the HIrisPlex[240] model to predict the hair and eye colour for two ancient individuals with highest average sequencing depth of coverage, i.e. VK1 and VK42. We used imputed genotypes for both individuals.

**Table S13.1**: The list of SNPs used for the HIrisPlex model for the VK1 and VK42 samples.

| Chromosome | Pos | SNP ID | Gene | Ref | Alt | VK1 | VK42 |
|---|---|---|---|---|---|---|---|
| **5** | 33951693 | rs16891982 | SLC45A2 | C | G | G/G | G/G |
| **5** | 33958959 | rs28777 | SLC45A2 | C | A | A/A | A/A |
| **6** | 396321 | rs12203592 | IRF4 | C | T | C/C | C/C |
| **6** | 457748 | rs4959270 | EXOC2-LOC105374875 | C | A | C/C | C/A |
| **9** | 12709305 | rs683 | TYRP1 | C | A | A/A | A/A |
| **11** | 88911696 | rs1042602 | TYR | C | A | C/A | C/A |
| **11** | 89011046 | rs1393350 | TYR | G | A | G/G | G/G |
| **12** | 89328335 | rs12821256 | KITLG | T | C | T/T | T/T |
| **14** | 92773663 | rs12896399 | SLC24A4-LOC105370627 | G | T | G/G | T/T |
| **14** | 92801203 | rs2402130 | SLC24A4 | G | A | A/A | A/A |
| **15** | 28230318 | rs1800407 | OCA2 | C | T | C/C | C/C |
| **15** | 28365618 | rs12913832 | HERC2 | A | G | G/G | A/A |
| **16** | 89985844 | rs1805005 | MC1R | G | T | G/G | G/G |
| **16** | 89985918 | rs1805006 | MC1R | C | A | C/C | C/C |
| **16** | 89985940 | rs2228479 | MC1R | G | A | G/A | G/G |
| **16** | 89986091 | rs11547464 | MC1R | G | A | G/G | G/G |

| | | | | | | | |
|---|---|---|---|---|---|---|---|
| **16** | 89986117 | rs1805007 | MC1R | C | T | C/C | C/C |
| **16** | 89986130 | rs1110400 | MC1R | T | C | T/T | T/T |
| **16** | 89986144 | rs1805008 | MC1R | C | T | C/C | C/C |
| **16** | 89986154 | rs885479 | MC1R | G | A | G/A | G/G |
| **16** | 89986546 | rs1805009 | MC1R | G | C | G/G | G/G |
| **20** | 33218090 | rs2378249 | ASIP-PIGU | G | A | G/A | A/A |

Table S13.1 summarizes the genotypes of each of the 22 SNPs used for phenotype analysis. For VK1 individual: the estimated probability of having blue eyes was 0.85, while the hair color probabilities were blond (0.63), brown (0.29), red (0.01) and black (0.07). For VK42 individual: the estimated probability of having brown eyes was 0.98, while the hair color probabilities were blond (0.15), brown (0.6), red (0.001) and black (0.25).

### Supplementary Note 14 - Finding signatures of selection in Europe in the past ten millennia

#### Introduction

We aimed to find SNPs whose allele frequencies changed significantly in the last 10,000 years, using our ancient human genomes to look at the frequencies of alleles in the past. We combined our VA and IA genomes with previously published present-day, Bronze Age, Neolithic and Mesolithic sequence data typed at the Human Origins array (see Supplementary Note 6). We filtered for genomes that were younger than 10,000 BP and that were located within a bounding box encompassing the European continent: 30 < latitude < 75 and -15 < longitude < 45. We then used neoscan in Ohana[241,242] to scan for variants whose allele frequencies were strongly associated with time, after controlling for genome-wide changes in ancestry that might have also occurred over time. We only analyzed sites with a minor allele frequency > 1%.

#### Methods

Briefly, Ohana works by modelling the allele frequencies across a specific number (K) of ancestry components via a multivariate Gaussian distribution. Each genome is modeled as a mixture of 1 or more components. For more details, see Cheng et al.[241,242]. After genome-wide ancestry component estimation via qpas, the neoscan method implemented in Ohana[243] tests a model in which the frequency of an allele at a site is determined by the genome-wide-estimated ancestry components against a model in which the frequency is also influenced by a free parameter (alpha) that determines the dependence of this frequency on the time at which a genome was sampled. Thus, at each SNP, neoscan computes a log-likelihood ratio that is equal to 2 * (best local log likelihood - global log likelihood). The global log likelihood is:

$$H_0 : \ln(L_j) = \sum_i^I \left( g_{ij} \cdot \ln\left( \sum_k^K \left( q_{ik} \cdot f_{kj}^{\text{est}} \right) \right) + (2 - g_{ij}) \cdot \ln\left( \sum_k^K \left( q_{ik} \cdot \left( 1 - f_{kj}^{\text{est}} \right) \right) \right) \right)$$

In turn, the local log likelihood is:

$$H_a : \ln(L_j) = \sum_i^I \left( g_{ij} \cdot \ln\left( \sum_k^K \left( q_{ik} \cdot f_{kj}^{\text{alt}_i} \right) \right) + (2 - g_{ij}) \cdot \ln\left( \sum_k^K \left( q_{ik} \cdot \left( 1 - f_{kj}^{\text{alt}_i} \right) \right) \right) \right)$$
$$f_{kj}^{\text{alt}_i} = \min\left( 1,\ \max\left( 0,\ f_{kj}^{\text{est}} + \alpha \cdot \frac{(t_i - t_{\text{ave}})}{t_{\text{ave}}} \right) \right)$$

The notation here follows from Cheng et al.[241]. I is the total number of sites. $g_{ij}$ is the called genotype for individual i at SNP j, where 0 is homozygous major, 1 is heterozygous, and 2 is homozygous minor. $q_{ik}$ is the estimated genome-wide proportion of ancestry from component k in individual i. $f_{kj}$ is the estimated frequency in component k at site j. The local log likelihood is maximized with respect to the parameter alpha, which interacts with the time of sampling of individual i ($t_i$) normalized by the average of all the sampling times ($t_{ave}$). This means that SNPs that are not well-modeled by the genome-wide mixture of ancestries, and whose deviations from this mixture depend strongly on time, will have large likelihood ratios.

Sampling times were set as 2000 before present (BP) for all Iron Age samples, 1200 BP for all Early Viking Age samples, 1000 BP for all Viking Age and Early Norse samples, 750 BP for all Medieval and Norse samples, and 500 BP for all Late Medieval samples. The sampling times for all other ancient published samples were obtained from the midpoint of their radiocarbon date range. In our final scan, we used K=3, but also verified that our top candidate SNPs were largely robust to the choice of K, by repeating the analysis with higher K, ranging from 4 to 7. The resulting likelihood ratios for all genotyped SNPs in the autosomes are displayed in Figure S14.1.

We call the above described scan the ‘general’ scan (Figure S14.2, Table S14.1), but we also performed two additional scans, focusing on either ‘ancient’ positive selection (older than 4000 BP, Figure S14.5, Table S14.2) or ‘recent’ positive selection (younger than 4000 BP, Figure S14.8, Table S14.3). For the recent scan, we set all times older than 4000 BP to be equal to 4000 BP, so that ancient selection would not affect the local log-likelihood. Conversely, for the ancient scan, we set all times younger than 4000 BP to be equal to 4000 BP.

For each analysis, we selected candidate SNPs whose log-likelihood ratio was larger than the 99.9% empirical quantile of the genome-wide distribution of log-likelihood ratios, and that had at least two nearby SNPs (in a +/- 500kb region) that had a ratio score larger than the same quantile. We note, however, that because we are working with a dataset intersected with the Human Origins SNPs, the highest scoring SNP in a given region may not be the causal one, as the latter may have not been selected for SNP capture. We plotted allele frequency time series for candidates SNPs (Figures S14.3, S14.6, S14.9 for the “general”, “ancient” and “recent” scans, respectively), grouping ancient genomes into five different periods: a period ranging from 10,000 to 8,000 BP, one ranging from 8,000 to 6,000 BP, one ranging from 6,000 BP to 4,000 BP, one ranging from 4,000 BP to 2,000 BP, one ranging from 2,000 BP to the present (excluding present-day samples) and one containing present-day samples only. We used Jeffreys priors to obtain 95% Bayesian credible intervals for the allele frequency of each time period. For each candidate SNP, we also plotted the scores in the local region surrounding the SNP (Figure S14.4, S14.7, S14.10 for the "general", "ancient" and "recent" scans, respectively). Finally, we visually verified whether any of the top selection candidates could appear as SNPs due to sequencing or mapping errors in the gnomAD browser[244].

#### Results

In the general scan, we mostly find previously described candidate regions for positive selection in Europe: the LCT/MCM6 region, the TLR region, the HLA region, SLC45A2 and SLC22A4, all of which have been previously reported to be under selection in Europe[160,206].

In the "ancient" selection scan, we find several new candidates for selection, including a region overlapping COL27A1, DNFB31 and AKNA. The region is centered around AKNA, which codes for a transcription factor that regulates CD40 and its ligand. It is expressed by B and T lymphocytes, natural killer cells and dendritic cells, and plays an important role in the secondary immune response[245]. Another candidate region includes the DCC gene, implicated in colorectal cancer[246]. A third new candidate region overlaps the DFFB and CEP104 genes. DFFB codes for a nuclease involved in apoptosis[247] while CEP104 codes for a protein involved in ciliary structural integrity and may be involved in Joubert syndrome[248]. In this scan, we also find a strong candidate for selection in a region overlapping the CXCR4 gene, though this one is close to the LCT/MCM6 region and shows similar allele frequency dynamics in time, so we cannot discard it may be part of the same selective event.

In the "recent" selection scan, we again recover the LCT/MCM6 region (with the highest score by far), the TLR region and the HLA region, suggesting that positive selection on these regions has either begun (for the LCT/MCM6 region) or persisted (for the other two regions) in recent times. However, the SLC45A2 region is not recovered in this scan, in agreement with the previously inferred history of allele frequency change in this region[160,206].

### Tables

**Table S14.1:** Top candidate SNPs from "general" neoscan temporal scan with K=3. SNPs are listed if they had a log-likelihood ratio score than the 99.9% quantile of the empirical distribution of log-likelihood ratios, and at least two neighboring SNPs (+/- 500kb) with a score larger than the same quantile.

| CHR | POSITION (hg19) | LIKELIHOOD RATIO | GENES (+/- 200 kb) |
|---|---|---|---|
| **2** | 136557319 | 66.053 | R3HDM1,UBXN4,LCT,MCM6,DARS |
| **6** | 31348164 | 31.690 | POU5F1,HCG27,HLA-C,HLA-B,MICA,MICB,MCCD1,ATP6V1G2-DDX39B,DDX39B,ATP6V1G2,NFKBIL1,LTA,TNF |
| **4** | 38859799 | 24.450 | KLF3,TLR10,TLR1,TLR6,FAM114A1,TMEM156,KLHL5 |
| **5** | 131586598 | 18.257 | PDLIM4,SLC22A4,SLC22A5,C5orf56,IL3,CSF2,P4HA2 |
| **5** | 33954511 | 11.715 | ADAMTS12,RXFP3,SLC45A2,AMACR,C1QTNF3 |

**Table S14.2:** Top candidate SNPs from "ancient" neoscan temporal scan with K=3. SNPs are listed if they had a log-likelihood ratio score than the 99.9% quantile of the empirical distribution of log-likelihood ratios, and at least two neighboring SNPs (+/- 500kb) with a score larger than the same quantile.

| CHR | POSITION (hg19) | LIKELIHOOD RATIO | GENES (+/- 200 kb) | NOTES |
|---|---|---|---|---|
| **9** | 117102046 | 41.088 | COL27A1,ORM1,ORM2,AKNA,DFNB31 | |
| **2** | 136976255 | 39.16 | CXCR4 | Near LCT/MCM6 region |
| **18** | 50686991 | 30.029 | DCC | |
| **1** | 3776166 | 27.294 | TP73,CCDC27,SMIM1,LRRC47,CEP104,DFFB,C1orf174 | |
| **5** | 131586598 | 26.997 | PDLIM4,SLC22A4,SLC22A5,C5orf56,IL3,CSF2,P4HA2 | |

| | | | |
|---|---|---|---|
| **6** | 29732302 | 23.505 | GABBR1,OR2H2,MOG,ZFP57,HLA-F,HLA-G,HLA-A |
| **1** | 61115790 | 21.933 | N/A |
| **5** | 33958910 | 20.607 | ADAMTS12,RXFP3,SLC45A2,AMACR,C1QTNF3 |
| **16** | 83508437 | 18.912 | CDH13 |
| **4** | 38886293 | 18.04 | KLF3,TLR10,TLR1,TLR6,FAM114A1,TMEM156,KLHL5 |
| **1** | 163837511 | 17.854 | N/A |

**Table S14.3:** Top candidate SNPs from "recent" neoscan temporal scan with K=3. SNPs are listed if they had a log-likelihood ratio score than the 99.9% quantile of the empirical distribution of log-likelihood ratios, and at least two neighboring SNPs (+/- 500kb) with a score larger than the same quantile.

| CHR | POSITION (hg19) | LIKELIHOOD RATIO | GENES (+/- 200 kb) | NOTES |
|---|---|---|---|---|
| **2** | 135631400 | 52.517 | TMEM163,ACMSD,CCNT2,MAP3K19,RAB3GAP1 | Near LCT/MCM6 region |
| **6** | 31348164 | 17.39 | POU5F1,HCG27,HLA-C,HLA-B,MICA,MICB,MCCD1,ATP6V1G2-DDX39B,DDX39B,ATP6V1G2,NFKBIL1,LTA,TNF | |
| **4** | 38859799 | 12.357 | KLF3,TLR10,TLR1,TLR6,FAM114A1,TMEM156,KLHL5 | |
| **8** | 140922006 | 11.382 | TRAPPC9,C8orf17 | May not be a true SNP: no rs ID |
| **13** | 34067497 | 8.713 | STARD13 | May not be a true SNP: no rs ID |

### Figures

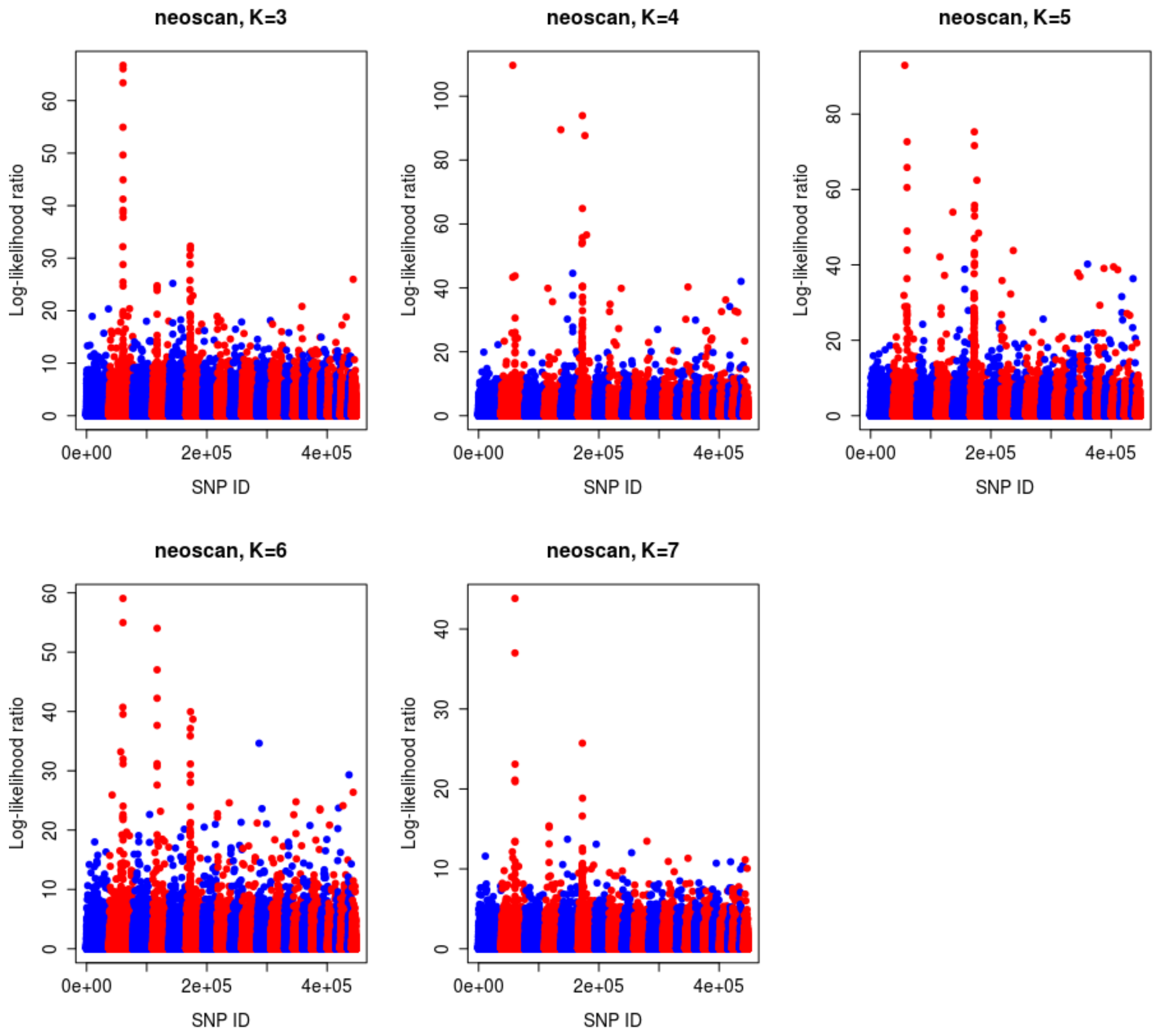


**Fig. S14.1:** Manhattan plots of Ohana neoscan with a latent ancestry fitting ranging from K=3 to K=7.

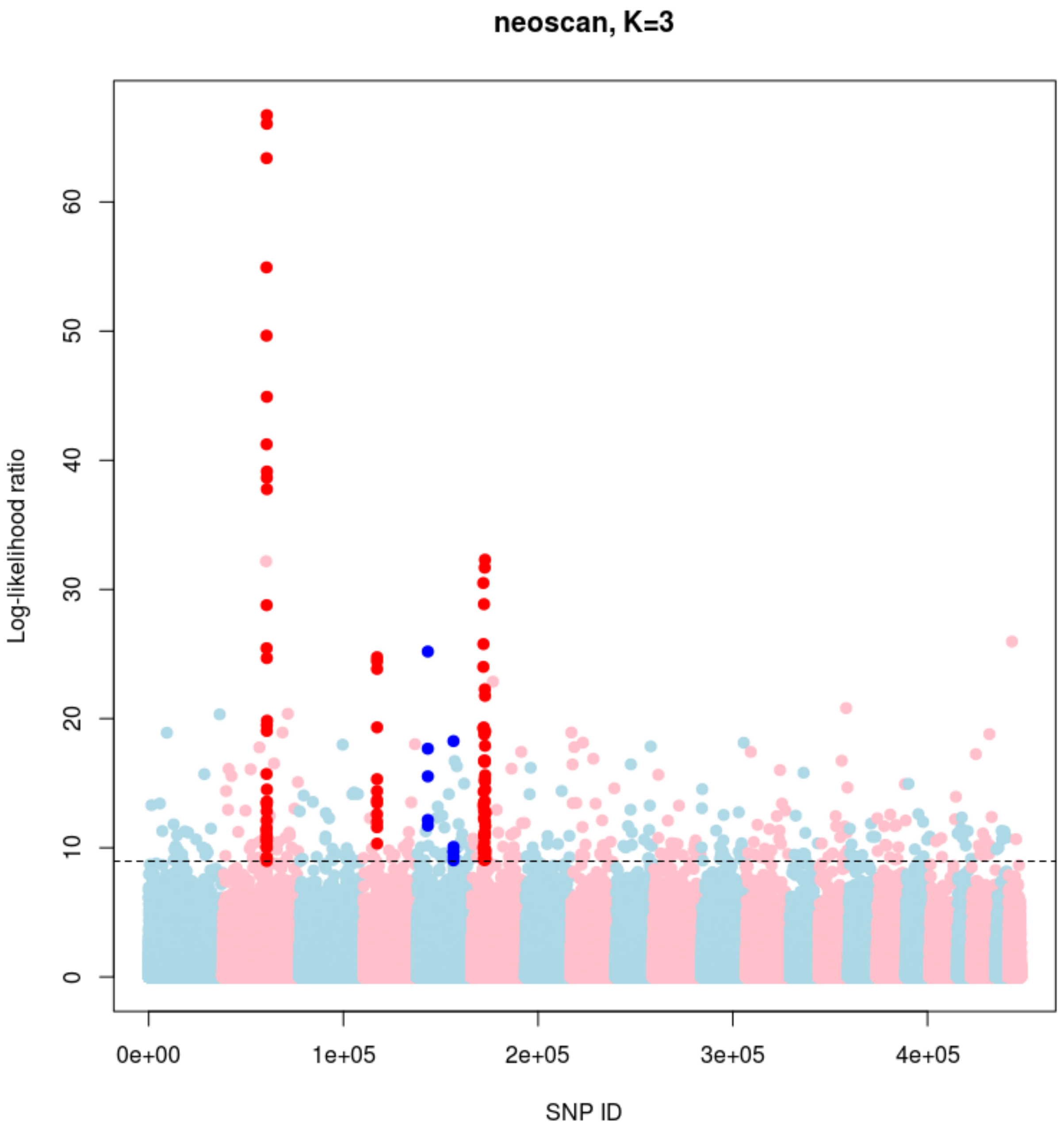


**Fig. S14.2:** Manhattan plot of Ohana "general" neoscan (K=3) looking for SNPs whose allele frequencies were strongly associated with time over the entire 10,000 BP period, after accounting for genome-wide changes in ancestry over time. The highlighted SNPs have a score larger than the 99.9% quantile of the empirical distribution of log-likelihood ratio, and have at least two neighboring SNPs (+/- 500kb) with a score larger than the same quantile.

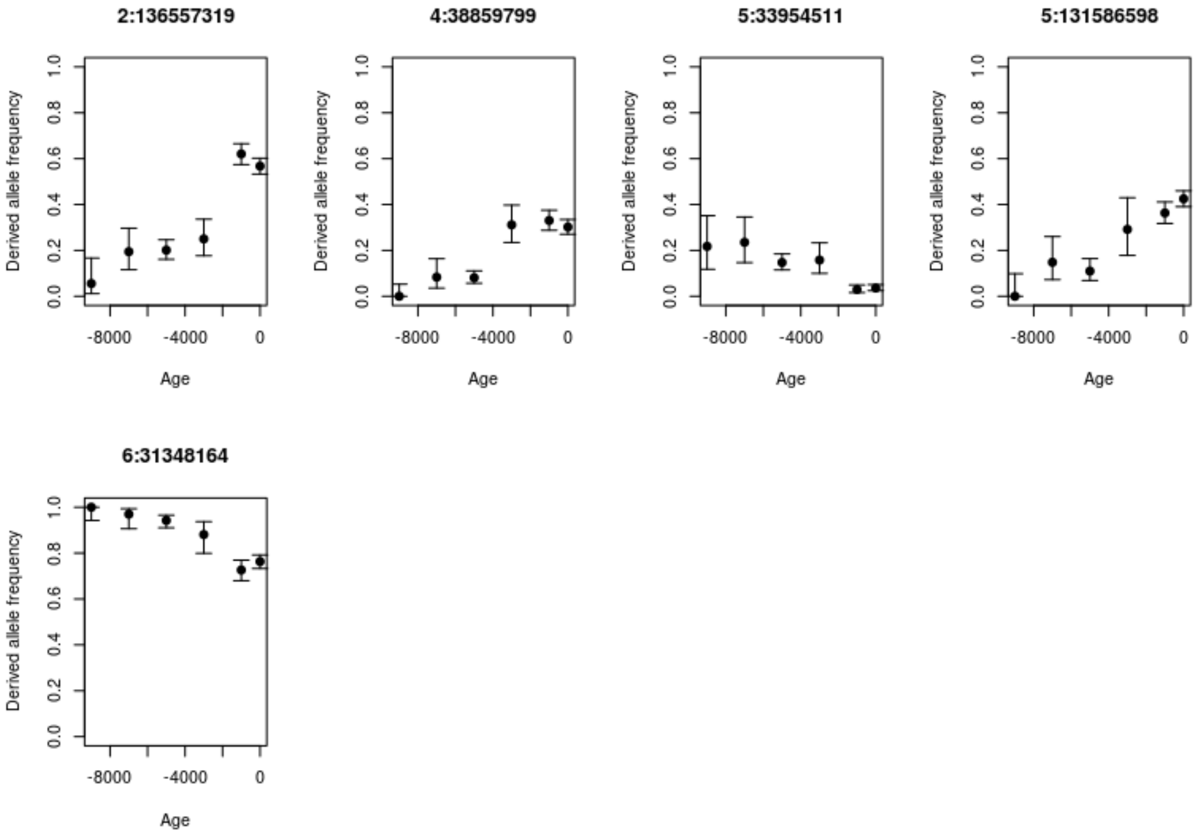


**Fig. S14.3:** Derived allele frequencies of top candidate SNPs from “general” scan with K=3, as a function of age, after aggregating ages of ancient samples into 2,000-year bins. 95% Bayesian credible intervals (error bars) were computed using a Jeffreys prior.

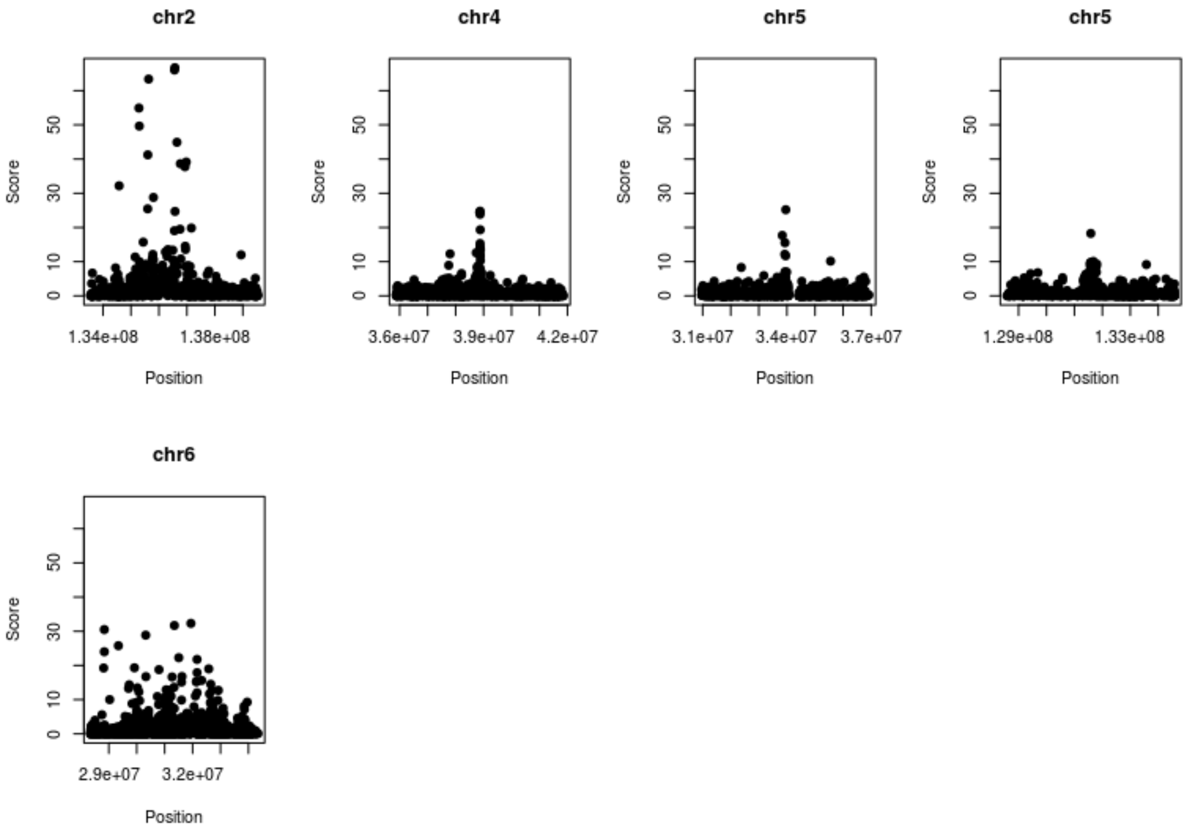


**Fig. S14.4:** Zoomed-in plots of regions (+/- 3 Mb) surrounding the candidate SNPs from the “general” scan.

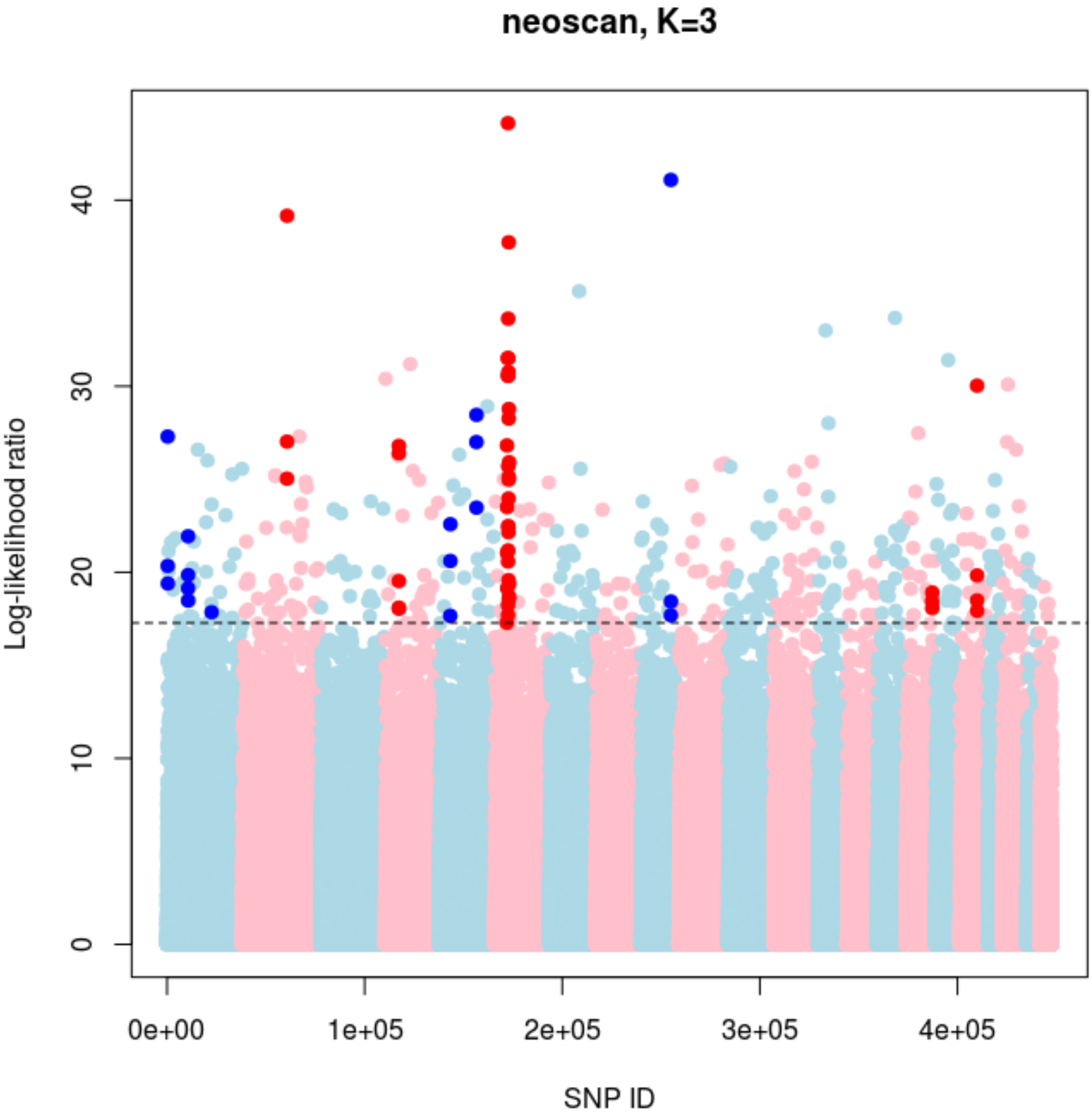


**Fig. S14.5:** Manhattan plot of Ohana ‘ancient’ neoscan (K=3) looking for SNPs whose allele frequencies were strongly associated with time after 4,000 BP, after accounting for genome-wide changes in ancestry over time. The highlighted SNPs have a score larger than the 99.9% quantile of the empirical distribution of log-likelihood ratio, and have at least two neighboring SNPs (+/- 500kb) with a score larger than the same quantile.

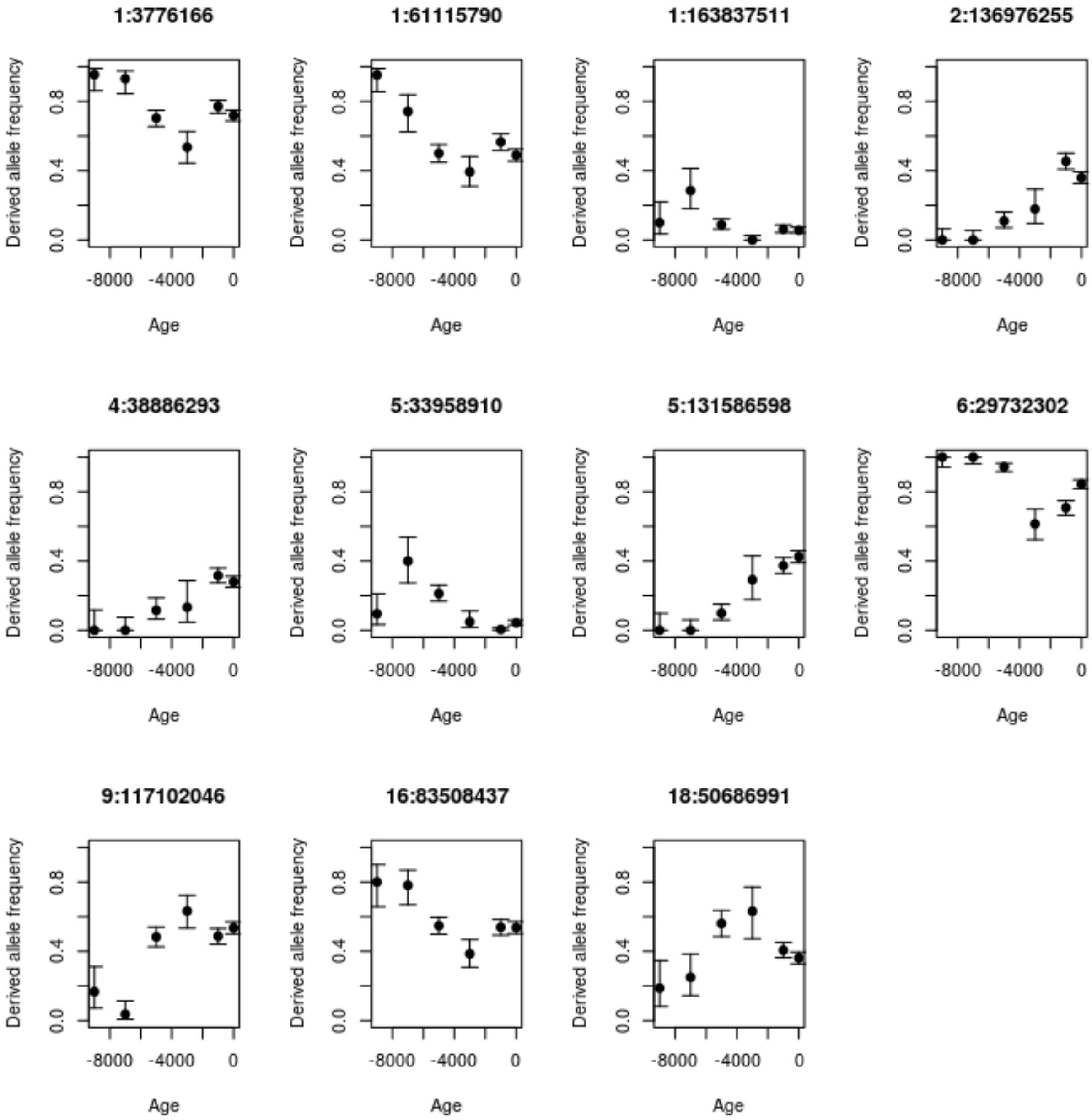


**Fig. S14.6:** Derived allele frequencies of top candidate SNPs from 'ancient' scan, as a function of age, after aggregating ages of ancient samples into 2,000-year bins. 95% Bayesian credible intervals (error bars) were computed using a Jeffreys prior.

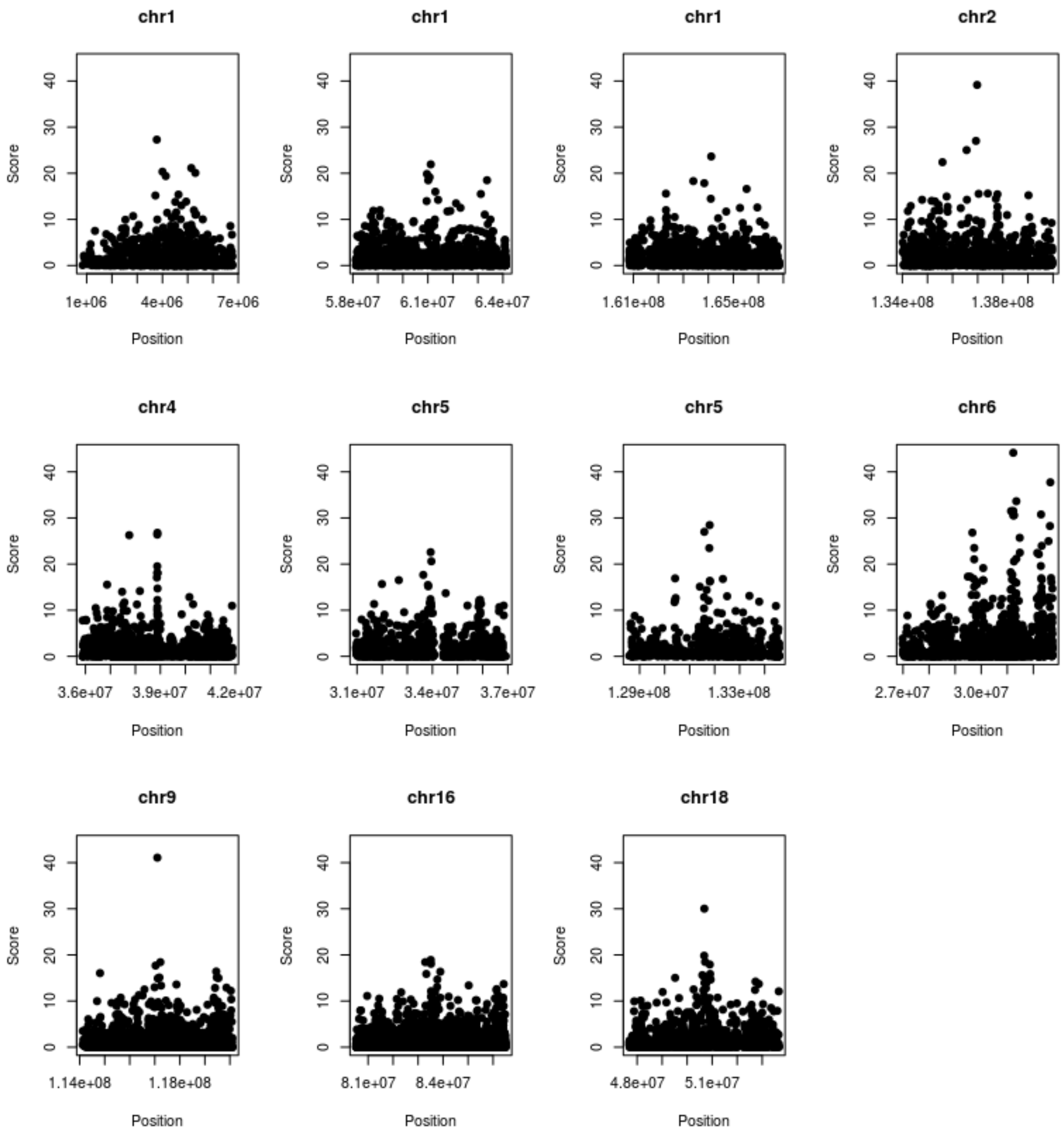


**Fig. S14.7:** Zoomed-in plots of regions (+/- 300 kb) surrounding the candidate SNPs from the 'ancient' scan.

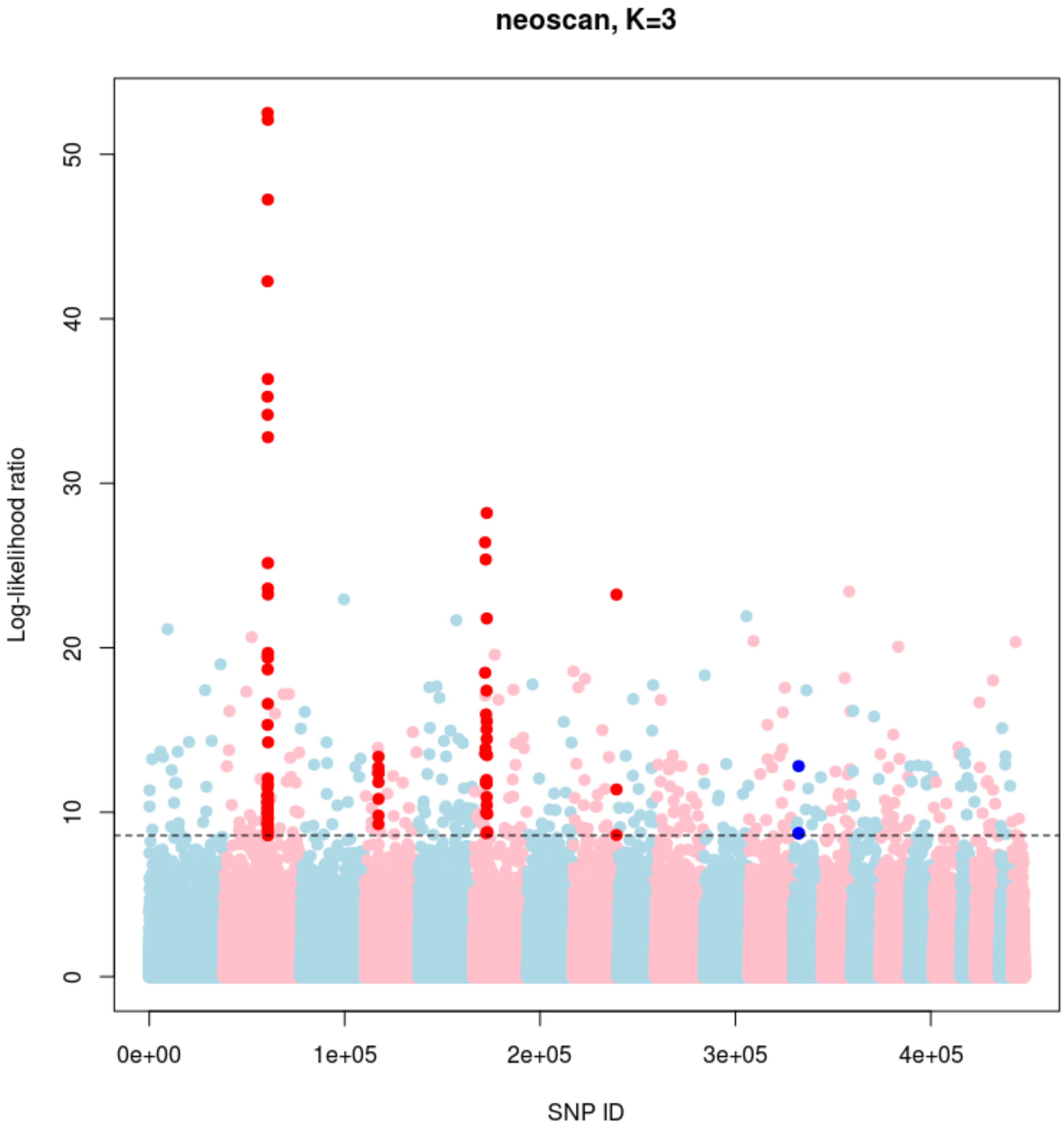


**Fig. S14.8:** Manhattan plot of Ohana 'recent' neoscan (K=3) looking for SNPs whose allele frequencies were strongly associated with time after 4,000 BP, after accounting for genome-wide changes in ancestry over time. The highlighted SNPs have a score larger than the 99.9% quantile of the empirical distribution of log-likelihood ratio, and have at least two neighboring SNPs (+/- 500kb) with a score larger than the same quantile.

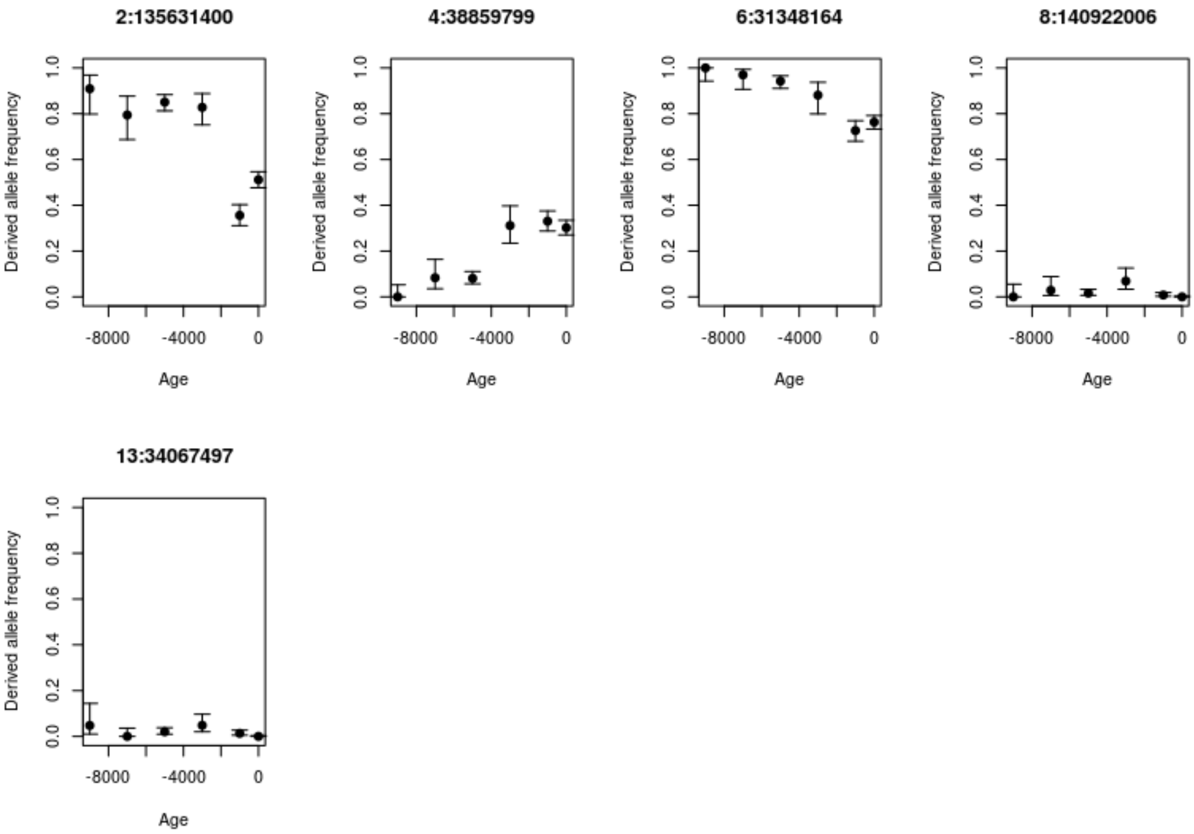


**Fig. S14.9:** Derived allele frequencies of top candidate SNPs from ‘recent’ scan, as a function of age, after aggregating ages of ancient samples into 2,000-year bins. 95% Bayesian credible intervals (error bars) were computed using a Jeffreys prior.

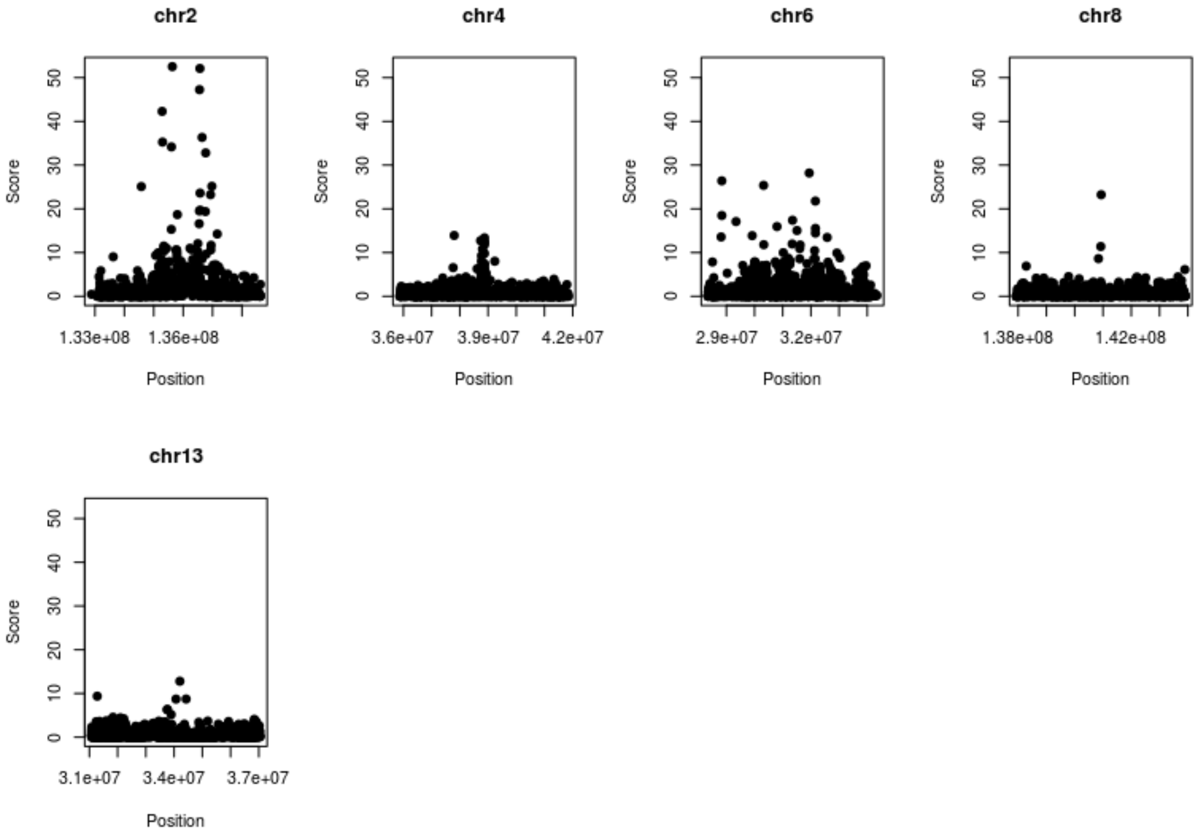


**Fig. S14.10:** Zoomed-in plots of regions (+/- 3 Mb) surrounding the candidate SNPs from the 'recent' scan.

### Supplementary Note 15

#### Tracking the evolution of complex traits in Scandinavia

We wanted to examine whether we could identify signals of recent population differentiation of complex traits by comparing genotypes of Viking Age samples excavated in Scandinavia (i.e. Denmark, Sweden and Norway) with those of a present-day Scandinavian population. We chose to focus on traits for which summary statistics from well-powered genome-wide association studies (GWAS) were available.

#### Samples & genotyping

For comparison with the Viking Age samples we used imputed genotypes from subjects born in Denmark between 1981-2011 from the IPSYCH case-cohort study[239]. To minimize potential bias from sample source and genotyping platform (the IPSYCH samples are genotyped on Illumina PsychArray and imputed with SHAPEIT3[249] and IMPUTE2[250] using 1000 Genomes as reference) we filtered both datasets on markers' imputation info (>0.98) and minor allele frequency (>0.1) before merging the datasets on c. 1.3M SNP markers present in both datasets. We then further filtered the merged dataset to include only samples and markers with >0.98 genotype yield.

#### Principal component analysis

To prevent the large present-day Danish sample from dominating the weights of the principal components, we used a subset of samples to estimate principal components, and then the rest of the sample was projected onto these components. We pruned genotypes reiteratively with respect to LD ($R2<0.1$ within a window of 100 adjacent SNPs) to a set of 21,013 uncorrelated autosomal SNPs and used all unrelated Viking age samples and a subset of unrelated IPSYCH samples enriched for ethnic diversity (≈1,000 random population samples with both parents born in Denmark and further ≈3,000 samples with both parents born outside Denmark, including ≈2,000 with both parents born outside Europe) and derived 25 ancestry-sensitive principal components (PC) with SMARTPCA[251].

#### Polygenic score analysis

We downloaded summary statistics from the Genome wide association study ATLAS webpage (https://atlas.ctglab.nl)[252], from studies of 16 disease- and anthropometric traits (excluding those related to cognition) published in 2017 or later with SNP heritability estimated at >0.1, sample size of >100,000, and >100 identified genome-wide significant loci. We calculated polygenic risk scores based on independent ($R^2$<0.1 within 10Mb range) genome-wide significant allelic effects and standardized them to a unit representing the standard deviation of the mean of their distribution. We then removed outliers (anyone with a value for any of the 25 PCs falling more than 4 standard deviations away from the group mean) reiteratively from within each ancestry group (treating the Scandinavian Viking age samples as one ancestry group), and subsequently tested for difference in PRS distribution between Viking age samples and Danish ancestry IPSYCH random population samples using a linear regression model correcting for sex and the 25 principal components. The analysis was done in R (version 3.5.0) and we use the *ggplot* function of the *ggplot2* package, and the *forest* function of the *metafor* package to plot the results in figure S15.1 and Extended Data Fig. 5.

#### Results

A plot of the first two PCs (Figure S15.1) shows clustering of the present-day IPSYCH random sample according to ancestry (as determined by parents' country/region of birth), and that the Viking age samples (N = 148) cluster together with the Danish ancestry IPSYCH random sample (N = 20,551). PRS for three of the 16 traits showed difference between PRS of Viking age samples and the Danish ancestry IPSYCH random population sample (Extended Data Fig. 5); these were PRS for black hair colour and standing height from GWAS of the UK biobank (N≈385,000), and for schizophrenia from a GWAS meta-analysis of the Psychiatric Genomics Consortium (N=105,318). The difference in PRS for height and schizophrenia between the Viking Age and present-day Danish random sample did however not remain significant after taking into account the number of tests.

To test whether the observed difference in PRS for black hair colour was driven by a few large effect loci, or otherwise dependent on the estimated allelic effects from the respective GWA studies, we performed a binomial test of the number of risk alleles found in higher frequency in the Viking Age sample (n=65) and the present-day sample (n=41), respectively, which showed a significant difference from a 50/50 distribution (P = 0.025). To further test whether this difference could be explained by the frequency distribution of the risk alleles, we performed a permutation test, in which we replaced the actual risk alleles for black hair colour 1,000 times with randomly drawn alleles matched on ancestral allele frequency. In 17 of the 1,000 permutations the distribution of risk alleles (according to whether the frequency was higher in the Viking Age or present-day sample) deviated as much, or more, from a 50/50 distribution, than we had observed with the actual risk alleles (adjusted P = 0.017). Hence, we conclude that the observed PRS difference is neither explained by a few large effect loci, nor by a general tendency of alleles with the same frequency distribution as the risk alleles to be found at differing frequencies between the Viking age and present-day sample. Thus, it appears that frequencies of established common alleles affecting hair colour have significantly changed in the Danish population since the Viking Age, whereas we do not observe any significant change for alleles affecting other common anthropometric traits and a few complex disorders. At the moment, we cannot conclude whether this difference is due to selection acting on these alleles between the Viking Age and the present time, or to some other factors, or whether a similar change in allele frequencies affecting hair colour has occurred in the other Scandinavian populations.

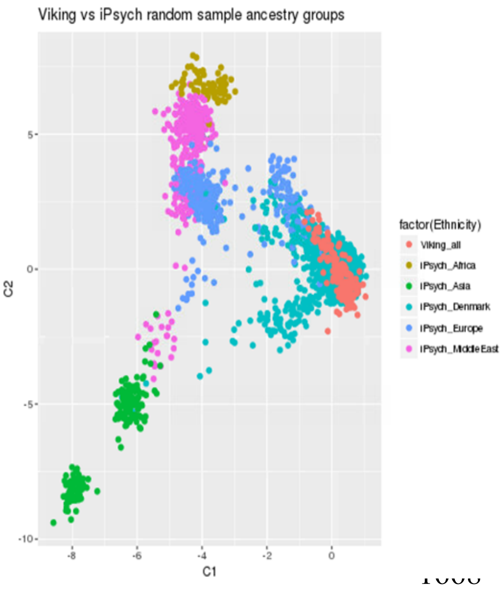


**Fig. S15.1**: A cluster plot of the two first principal components across Viking Age samples (red) and IPSYCH randomly drawn Danish population sample grouped by parents' birthplace (only using samples where both parents are born either in Denmark or one of the other respective regions). The Viking age samples fall mostly within the cluster of Danish ancestry.
